# MicroCOSM: a model of social and structural drivers of HIV and interventions to reduce HIV incidence in high-risk populations in South Africa

**DOI:** 10.1101/310763

**Authors:** Leigh F. Johnson, Mmamapudi Kubjane, Haroon Moolla

## Abstract

**Background and objectives:** South Africa has one of the highest HIV incidence rates in the world. Although much research has focused on developing biomedical strategies to reduce HIV incidence, there has been less investment in prevention strategies that address the social drivers of HIV spread. Understanding the social determinants of HIV is closely related to understanding high-risk populations (‘key populations’), since many of the factors that place these key populations at high HIV risk are social and behavioural rather than biological.

Mathematical models have an important role to play in evaluating the potential impact of new HIV prevention and treatment strategies. However, most of the mathematical modelling studies that have been published to date have evaluated biomedical HIV prevention strategies, and relatively few models have been developed to understand the role of social determinants or interventions that address these social drivers. In addition, many of the mathematical models that have been developed are relatively simple deterministic models, which are not well suited to simulating the complex causal pathways that link many of the social drivers to HIV incidence. The frequency-dependent assumption implicit in most deterministic models also leads to under-estimation of the contribution of high-risk groups to the incidence of HIV.

Agent-based models (ABMs) overcome many of the limitations of deterministic models, although at the expense of greater computational burden. This study presents an ABM of HIV in South Africa, developed to characterize the key social drivers of HIV in South Africa and the groups that are at the highest risk of HIV. The objective of this report is to provide a technical description of the model and to explain how the model has been calibrated to South African data sources; future publications will assess the drivers of HIV transmission in South Africa in more detail.

**Methods:** The model is an extension of a previously-published ABM of HIV and other sexually transmitted infections (STIs) in South Africa. This model simulates a representative sample of the South African population, starting from 1985, with an initial sample size of 20 000. The population changes in size as a result of births and deaths. Each individual is assigned a date of birth, sex and race (demographic characteristics). This in turn affects the assignment of socio-economic variables. Each individual is assigned a level of educational attainment, which is dynamically updated as youth progress through school and tertiary education, with rates of progression and drop-out depending on the individual’s demographic characteristics. Each individual is also assigned to an urban or rural location, with rates of movement between urban and rural areas depending on demographic characteristics and educational attainment.

The model assigns to each individual a number of healthcare access variables that determine their HIV and pregnancy risk. These include their ‘condom preference’ (a measure of the extent to which they wish to use condoms and are able to access condoms), use of hormonal contraception and sterilization, use of pre-exposure prophylaxis (PrEP), male circumcision, HIV testing history and uptake of antiretroviral treatment (ART). Access to these healthcare services changes over time, and is also assumed to depend on demographic and socioeconomic variables, as well as on the individual’s health status.

Sexual behaviour is simulated by assigning to each individual an indicator of their propensity for concurrent partnerships (‘high risk’ individuals are defined as individuals who have a propensity for concurrent partnerships or commercial sex). Each individual is also assigned a sexual preference, which can change over their life course. Three types of relationship are modelled: sex worker-client contacts, short-term (non-marital) relationships and long-term (marital or cohabiting) relationships. Individuals are assumed to enter into short-term relationships at rates that depend on their risk group and demographic characteristics. Each time a new short-term partner is acquired, the individual is linked to another individual in the population, with the probability of linkage depending on the individual’s sexual preference and preference for individuals of the relevant age, risk group, race, location and educational attainment. Individuals marry their short-term partners at rates that depend on their demographic characteristics. Frequencies of sex are assumed to depend on demographic characteristics and relationship type, and migrant couples are assumed to have reduced coital frequency. Probabilities of condom use also depend on demographic characteristics and relationship type, and are assumed to be strongly associated with levels of educational attainment.

Women’s risk of falling pregnant is assumed to depend on their sexual behaviour, natural fertility level, contraceptive usage and breastfeeding status. Adoption and discontinuation of hormonal contraception is assumed to depend on demographic characteristics, sexual behaviour and past pregnancy and contraceptive experience. Girls who fall pregnant while in school are assumed to be less likely to complete their schooling than those who do not fall pregnant.

Probabilities of HIV transmission per act of sex are assumed to depend on several biological factors, including the viral load of the HIV-positive partner, whether the HIV-positive partner is on ART, the presence of other STIs, the type of contraceptive used, the age and sex of the susceptible partner, male circumcision, the type of relationship, and the use of new HIV prevention methods such as PrEP. If an individual acquires HIV, they are assigned a CD4 count and viral load, both of which change dynamically over the course of HIV infection. The HIV mortality risk is determined by the individual’s CD4 count. HIV-positive individuals are diagnosed at rates that depend on their demographic characteristics and CD4 count, and if they disclose their HIV status to their sexual partners after diagnosis, this is assumed to lead to increased rates of condom use. Assumptions about HIV transmission probabilities have been set in such a way that the model matches the observed trends in HIV prevalence, by age and sex, in national South African antenatal and household surveys.

The model also simulates male incarceration. Rates of incarceration are assumed to depend on men’s demographic characteristics and educational attainment, and are also assumed to be higher in men who have previously been incarcerated.

**Results and conclusions:** The model matches reasonably closely the observed levels of HIV prevalence in South Africa by age and sex, as well as the observed changes in HIV prevalence over time. The model also matches observed patterns of HIV prevalence by educational attainment, by urban-rural location and by history of recent migration. Estimates of HIV prevalence in key populations (sex workers, MSM and prisoners) are roughly consistent with surveys. The model has also been calibrated to match total numbers of HIV tests and male circumcision operations performed in South Africa. The model estimates of levels of HIV diagnosis and ART coverage are consistent with the Thembisa model, an HIV model that has been calibrated to South African HIV testing and ART data.

Although many of the phenomena simulated in the MicroCOSM model have been simulated in previously-published HIV models, MicroCOSM is the first model that systematically describes all of these phenomena in a fully integrated model. This makes it possible to use the model to describe complex interactions between socio-economic and behavioural factors, and their influence on disease and health-seeking behaviour. It also provides a framework for understanding socio-economic and racial inequality in health outcomes in South Africa, and for assessing the potential impact of strategies to reduce these inequalities.

## 1. Introduction

South Africa has one of the highest levels of HIV prevalence in the world [1]. Although there has been much success in reducing rates of mother-to-child transmission of HIV [2, 3] and AIDS mortality in South Africa [4, 5], adult HIV incidence rates remain unacceptably high [6]. Much HIV prevention research has focused on biomedical interventions, which address the proximal determinants of HIV transmission. However, there has been less focus on the social and structural drivers of HIV, i.e. the more distal determinants of HIV spread. Some of the commonly considered social factors include high levels of migration, rapid urbanization, intimate partner violence, gender inequality, low levels of education, income inequality, high cost of lobola and associated late entry into marriage, high levels of incarceration, homophobia, HIV stigma, alcohol and substance use, and social norms around family planning [7]. New interventions may be important in addressing these social drivers. For example, cash transfer interventions may be able to reduce gender inequality [8] and community mobilization interventions may be effective in preventing violence against women [9].

Understanding the social and structural determinants of HIV is closely related to understanding high-risk populations (or ‘key populations’). Some of the commonly-identified high risk groups include sex workers, men who have sex with men (MSM), people who inject drugs, young women, migrants, prisoners and individuals in serodiscordant relationships [7]. In some cases, the elevated risks of HIV in these key populations are due to biological factors; for example, the high HIV prevalence in MSM is partly due to the higher risk of HIV transmission in acts of anal intercourse than in acts of vaginal intercourse [10]. In many cases, though, social and structural drivers are also important in explaining the elevated HIV risk. For example, the high levels of HIV prevalence in migrants may be partly attributable to the social environments in urban informal settlements, through which migrants often pass. Regardless of the reasons for the high HIV incidence rates in key populations, it is important that interventions be tailored to these high risk groups.

Mathematical models have played an important role in evaluating the role of HIV prevention and treatment strategies, but relatively few models have been used to help understand the role of social drivers of HIV [11]. There is a need for such models to quantify the effect of social drivers on risk behaviours and HIV incidence, as the effects are often difficult to measure empirically. For example, several models have been developed to simulate the potential effects of circular migration on the spread of HIV in South Africa [12–14]. Models have also been used to show how high rates of male incarceration can change the structure of sexual networks [15]. In addition, models have shown that high rates of HIV incidence in sex workers may be partly due to unsafe work environments and criminalization of sex work, which are associated with client sexual violence and police harassment [16].

The majority of mathematical models that have been developed to simulate HIV interventions are deterministic models [11], which divide populations into compartments of individuals who are assumed to be homogeneous in their behaviour and HIV risk. An obvious limitation of this modelling approach is that it is not possible to simulate individual-level variation in behaviours and biological determinants of HIV transmission. Another important limitation is that most deterministic models of HIV rely on the frequency-dependent assumption, i.e. the assumption that an individual’s risk of acquiring HIV is proportional to the prevalence of HIV in the population [17]. In more realistic network models, the risk of acquiring HIV depends only on whether the individual’s current partner(s) have HIV. Frequency-dependent models tend to under-estimate the importance of high risk groups in driving HIV transmission [18], and this is a major drawback when evaluating the relative efficiency of different targeting strategies. Another disadvantage of deterministic models is that they are less flexible in modelling the complex causal pathways that link social drivers and HIV incidence. For example, men who perpetrate intimate partner violence (IPV) are also typically men who have multiple partners [19], and thus women who are exposed to IPV might be at an increased risk of HIV not because of the direct effect of IPV but because of the indirect effect of their partners’ greater engagement in HIV risk behaviour. Deterministic, frequency-dependent models are less well suited to modelling such correlations in behaviour because they do not assign individuals to specific partners.

Agent-based models (ABMs), also commonly referred to as microsimulation models, are able to overcome many of the limitations of deterministic models [20]. These models simulate each individual in the population as a separate unit, assigning different characteristics to each individual and thus overcoming the assumption of homogeneity that is implicit in deterministic models. Most ABMs are network models that do not rely on the frequency-dependent assumption, making them more realistic in modelling the effect of interventions in high-risk groups. In addition, ABMs are more appropriate for simulating network effects and complex causal pathways [20], which are not easily understood when dealing with homogeneous model compartments. However, ABMs require substantially greater computing power and computing time than traditional deterministic models. In addition, their results have stochastic variation associated with them, and this can cause imprecision in model estimates.

MicroCOSM (Microsimulation for the Control of South African Morbidity and Mortality) is an ABM developed to simulate the epidemiology of various diseases and the social processes that drive these diseases in South Africa. The model is an extension of a microsimulation model that was originally developed to simulate HIV and other sexually transmitted infections (STIs) in South Africa [18]. This report describes the extensions that have been made to the model for the purpose of modelling social drivers of HIV and key populations. Although the focus of this report is limited to HIV, the model structure is highly flexible, and work is currently in progress to include the modelling of other diseases such as tuberculosis, diabetes and cervical cancer. Some components of the model are described in detail in other publications. For example, the modelling of IPV and its association with HIV is described in a separate publication [21]. Because IPV was not found to be a major driver of HIV transmission, the model described here does not include IPV or IPV interventions. The modelling of STIs other than HIV is also described in more detail elsewhere [18].

This report is intended to serve as a technical description of the MicroCOSM model and its calibration to South African data sources. Future publications will present more detailed evaluations of the role of different socials drivers and the importance of interventions in key populations.

## 2. Overview of methodology

The model described in this report is an extension of a previously-described network model of HIV and other STIs in South Africa [18]. This model simulates a representative sample of the South African population, starting from 1985, with an initial sample size of 20 000. The population changes in size as a result of births and deaths. Each individual is assigned a date of birth, sex and race. The key variables that are defined for each individual in the model are summarized in Figure 2.1.

**Figure 2.1:**
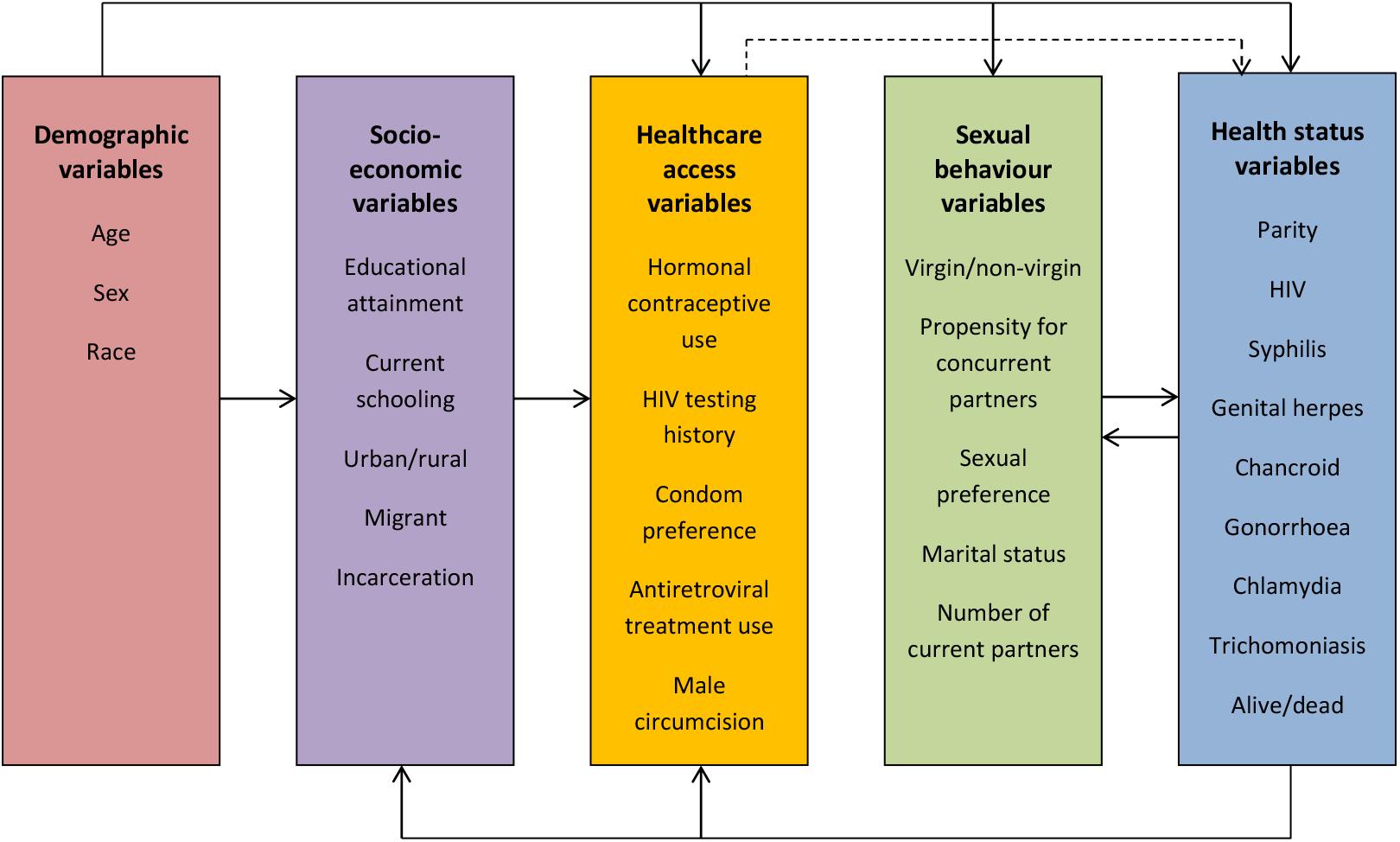
Variables defined for each individual in the MicroCOSM model. Arrows represent assumed causal relationships.

A number of socio-economic variables are defined for each individual. These include educational attainment (highest grade completed, including an additional category for tertiary qualifications) and whether currently in school or a tertiary institution. Individuals are also assigned to urban or rural locations. These socioeconomic variables are modelled dynamically; individuals are assumed to progress through successive grades while they remain in school and can also migrate between urban and rural areas. Probabilities of migration between urban and rural areas are assumed to depend on age, sex, race, marital status and level of education. In addition to these ‘permanent’ migrations, the model also simulates trips from urban to rural areas (and vice versa), of short durations, in the case of couples who are geographically separated.

The model assigns to each individual a number of healthcare access variables that determine their HIV and pregnancy risk. These include their ‘condom preference’ (a measure of the extent to which they wish to use condoms and are able to access condoms), use of hormonal contraception and sterilization, use of pre-exposure prophylaxis, male circumcision, HIV testing history and uptake of antiretroviral treatment. Access to these healthcare services changes over time, and is also assumed to depend on demographic and socioeconomic variables, as well as on the individual’s health status. Another important variable related to HIV testing history is the individual’s disclosure of their HIV status to sexual partners; individuals who have disclosed that they are HIV-positive to their sexual partners are assumed to be more likely to use condoms.

Several sexual behaviour characteristics are simulated. Individuals are classified as either having a propensity for concurrent partnerships (‘high risk’) or as being serially monogamous (‘low risk’). Individuals are also assigned a sexual preference, which can change over the life course. Three types of partnerships are modelled: short-term relationships (not married or cohabiting), long-term relationships (marital or cohabiting), and contacts between sex workers and their clients. Although commercial sex is not modelled in MSM, the model makes allowance for a fourth relationship category in MSM, namely casual sex (once-off sexual encounters). At weekly time steps, individuals can acquire a new partner, end their existing partnerships or marry their short-term partner. Acquisition of a new partner involves randomly sampling from the pool of available partners in the simulated population, with selection of partners based on assumed preferences for partners of the same age, race, educational attainment, risk group and location (urban or rural). Assumptions about coital frequencies and condom use differ according to the type of relationship, and also depend on factors such as age, educational attainment and race.

The model simulates male incarceration. Each man is assigned an incarceration status, which indicates whether he is out of prison, awaiting trial while in prison, or sentenced and in prison. Men who have previously been incarcerated are assumed to be at an increased risk of re-arrest when compared to men with the same characteristics who have not previously been arrested. Rates of arrest are also assumed to depend on age, educational attainment and race.

Women’s risk of falling pregnant is assumed to depend on a number of factors, including their natural fertility, their number of sexual partners, their use of condoms and other contraceptives, whether they are currently breastfeeding, their HIV status and stage of HIV disease, and their age and race. Women who fall pregnant are assigned a date of conception, a date of delivery and a date of weaning (if they breastfeed). Children who are born to HIV-positive mothers have a risk of acquiring HIV, either perinatally or postnatally, with the extent of this transmission risk changing over time, depending on maternal access to antiretroviral prophylaxis and replacement feeding.

Sexual HIV transmission risks are assumed to depend on a host of behavioural and biological factors. The biological factors affecting the transmission risk per sex act include the type of sex act, the presence of other STIs, the age and sex of the susceptible individual, the HIV viral load of the HIV-positive partner, use of hormonal contraception, pregnancy, male circumcision, and use of other prevention methods (PrEP and condoms). The behavioural factors include many of the behavioural variables referred to previously, including the number of sexual partners, coital frequency, type of relationship and patterns of sexual mixing (which determine the partner’s risk of HIV infection). Because of the substantial uncertainty that exists regarding the probabilities of HIV transmission per sex act, we assign prior distributions to represent the uncertainty regarding the average transmission probabilities that apply in heterosexual intercourse, for different partnership types. We randomly sample from these prior distributions and identify the HIV transmission parameters that give the best fit to South African HIV prevalence data.

Although the model simulation begins in 1985, HIV is only introduced into the population in 1990, by randomly assigning a proportion of individuals in the high risk group an HIV-positive status in 1990. This is partly so that there is a 5-year ‘burn-in’ period, in which the social and behavioural conditions affecting HIV spread can reach their ‘steady-state’ levels prior to introducing HIV. It is also partly because the extent of the stochastic variation in the model outputs is reduced when the HIV epidemic is initialized in 1990 rather than 1985. Most model results are therefore only presented from 1990 onward.

Each adult who acquires HIV is randomly assigned an initial CD4 count and viral load, and these are updated at weekly time steps. Changes in CD4 count and viral load are mutually dependent, and depend also on the individual’s age and sex. Rates of HIV-related mortality depend on the individual’s CD4 count, and the incidence of opportunistic infections is also CD4-dependent. The model allows for HIV testing through a number of different testing modalities, including ‘general’ HIV testing (mostly self-initiated facility-based testing), antenatal testing, testing of patients with opportunistic infections, testing in STI clinics, testing following partner notification, testing prior to medical male circumcision, and testing in prisons. Once individuals have been diagnosed HIV-positive, they can start ART, their rate of ART initiation depending on ART eligibility criteria and ART availability, both of which change over time. The model also allows for ART interruptions and re-initiation of ART.

To each individual we assign an ID number. For each individual we also record the ID of their primary and secondary partners (and ID of commercial or casual sex contacts when relevant). This allows us to link individuals in a sexual network and thus determine the transmission of HIV and other STIs.

## 3. Demographic and socio-economic variables

### 3.1 Initial population size by age, sex and race

The simulation starts with a population of 20 000 individuals, in mid-1985. This is intended to be a representative sample of the South African population, and the profile of the starting population – by age, sex and race – is thus sampled from national estimates of the starting population in 1985. The starting population profile is taken from the ASSA2008 AIDS and Demographic model [22], a model developed by the Actuarial Society of South Africa (ASSA). The ASS2008 model estimates a population of 32.3 million in 1985, and the initial population of 20 000 individuals is thus a 0.062% sample of the national total.

The initial population profile is taken from the ‘full’ version of the ASSA2008 model, which stratifies the population by population group (race). The ASSA2008 model estimates that of the starting population, 73.7% is black, 14.0% is white, 9.5% is ‘coloured’ (a term widely used in South Africa to refer to individuals of mixed race) and 2.8% is Asian. The Asian population is small and most surveys lack sufficiently large sample sizes to estimate parameters for the Asian population with any degree of precision. We have therefore not modelled Asians as a separate race group. However, because the Asian population is close to the white population in socio-economic characteristics [23], urbanization [24] and HIV prevalence [25], we have grouped together Asian and white populations for the purpose of assigning the initial population to different race groups. This means that 16.8% of the initial population is assigned to the ‘white/Asian’ group in the model.

### 3.2 Non-HIV mortality rates

Non-HIV mortality rates are specified by age, sex and race, and are assumed to change over time. The assumed non-HIV mortality rates are the same as those in the ASSA2008 full model [22], which estimates non-HIV mortality rates through calibration of the model to recorded death statistics.

### 3.3 Contraception

For the purpose of the sections that follow, the term ‘contraception’ is used to refer to either injectable contraception, oral contraception, or female sterilization. Although condoms are considered in the model, assumptions about their use are described elsewhere (see section 4.7). The term ‘hormonal contraception’ is used here to refer to either injectable or oral contraception. The model does not consider hormonal implants, as these were introduced only in 2014 and uptake has not yet reached high levels [26, 27].

#### 3.3.1 Assumptions about prevalence of contraception use in 1985

At the start of the simulation, in 1985, all women are assigned an initial contraceptive usage status. Women who are younger than 15, older than 49, or who are pregnant at the start of the simulation, are assumed not to be using any contraceptive method. For women aged 15-49, who are not pregnant, we assign initial probabilities of any modern contraceptive usage in such a way that the overall patterns of contraceptive usage match those in the 1987-89 DHS, as reported by Burgard [28]. The 1987-89 DHS excluded women who had never been pregnant and who had never been in a marital/cohabiting relationship. For these women, we relied on data from other studies to determine relative rates of contraceptive usage.

Based on the data presented by Burgard [28], rates of contraceptive usage are specified separately for each race and for each of three age groups (15-24, 25-34 and 35-49). The baseline rates (shown in Table 3.3.1) are those for women who are sexually active, cohabiting/married, whose highest educational attainment is grade 10, and who have had a previous child.

**Table 3.3.1:**
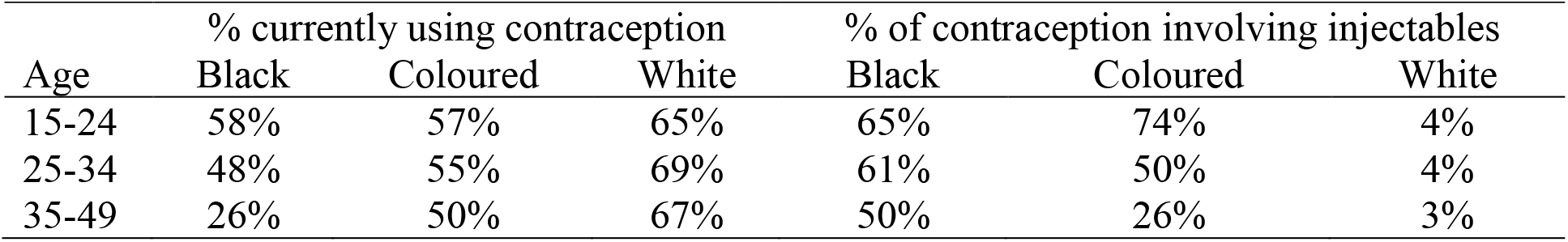
Proportion of sexually active, married/cohabiting, ever-pregnant women, with grade 10, who are assumed to be using modern contraception, in 1985

| Age | % currently using contraception |  |  | % of contraception involving injectables |  |  |
| --- | --- | --- | --- | --- | --- | --- |
|  | Black | Coloured | White | Black | Coloured | White |
| 15-24 | 58% | 57% | 65% | 65% | 74% | 4% |
| 25-34 | 48% | 55% | 69% | 61% | 50% | 4% |
| 35-49 | 26% | 50% | 67% | 50% | 26% | 3% |

The effect of marital status on contraceptive usage appears to depend on women’s birth history, and odds ratios for the association between marriage, previous birth and contraceptive usage are therefore specified as interactions (Table 3.3.2). The odds ratios in the first row are set to 1, since this corresponds to the baseline category in Table 3.3.1. The odds ratios in the second and third rows are estimated based on logistic regression models fitted by Burgard [28]. However, as the Burgard analysis excluded women who had never been married and never had a child, it cannot be used to estimate the odds ratios in the final row. MacPhail *et al* [29] report that in sexually active women aged 15-24, marital status was not significantly associated with modern contraceptive usage after controlling for other factors (such as sexual activity and previous children). It is therefore assumed that the odds ratio for unmarried, never-pregnant women is the same as that for married, never-pregnant women.

**Table 3.3.2:** Assumed effects of marital status and prior pregnancy on the odds of contraceptive usage in sexually experienced women (specified as odds ratios)

| Marital status | Gravidity | Population group |  |  |
| --- | --- | --- | --- | --- |
|  |  | Black | Coloured | White |
| Married/cohabiting | Ever pregnant | 1 | 1 | 1 |
|  | Never pregnant | 0.3 | 0.3 | 0.6 |
| Unmarried, not cohabiting | Ever pregnant | 1.7 | 0.85 | 0.5 |
|  | Never pregnant | 0.3 | 0.3 | 0.6 |

Odds ratios are also specified to represent the effect of not being sexually active, and the effect of educational attainment. MacPhail *et al* [29] report that in sexually active women aged 15-24, women who were sexually active in the last month were 1.5 times more likely to use contraception than women who had been sexually active in the last 12 months but not in the last month. The odds ratio for sexual inactivity is therefore set to 0.67 (1/1.5), if women are sexually experienced. The odds ratio per additional grade passed is set at 1.10, based on the logistic regression model fitted by Burgard [28].

It is assumed that in women who are not sexually experienced, contraceptive usage is rare. Data from the 1998 DHS [30] are summarized in Table 3.3.3. Based on these data, we assume that the rate of contraceptive usage in women aged 15-49, who are not yet sexually experienced, is 3%. (Due to a lack of more detailed data, this proportion is assumed to be the same for all virgins, regardless of age, race, education, etc.) This is consistent with a South African qualitative study, in which very few teenagers who used family planning services reported having visited family planning clinics prior to starting sexual activity [31]. It is also consistent with a study of users of injectable contraceptives in 10 South African clinics, of whom only 0.2% reported that they were not yet sexually experienced [32].

**Table 3.3.3:** Rates of contraceptive usage in women aged 15-19 in 1998 DHS

|  | n | % using modern<br>contraception |
| --- | --- | --- |
| All women aged 15-19 | 2249 | 28.5% |
| Less Sexually active in last 4 weeks | 412 | 64.4% |
| Less Ever sexually active, but not in last 4 weeks | 605 | 54.7% |
| = Never sexually active | 1232 | 3.7% |

At baseline, those women who are assigned to use of contraception are assigned to one of three methods: injectable contraception, the pill (oral contraception), or female sterilization. (Only a small fraction of women reported using condoms for the purposes of contraception in the 1987-89 DHS [33], and in the interests of simplicity, we therefore do not consider this in setting the baseline contraceptive usage.) Table 3.3.1 shows the assumed fractions of contraceptive users (in the baseline category) who are assumed to be using injectables. As before, these assumptions have been set in such a way that the model matches the results of the 1987-89 DHS, as reported by Burgard [28]. Consistent with other studies, these data suggest a high rate of injectable contraception use among black South African women [34–36], and relatively low rates among white women. The baseline rates in Table 3.3.1 are adjusted by an odds ratio of 0.55 in the case of women who have never been pregnant, and by an odds ratio of 0.90 per additional grade passed, based on the logistic regression models fitted by Burgard.

The use of female sterilization is strongly related to age and previous childbearing. In women who are assigned to contraception use at baseline, but who are not assigned to use of injectable contraception, it is assumed that a certain proportion has been sterilized. This proportion is set to zero in women who have never had children, and is assumed to depend on age among those women who have had children. Table 3.3.4 shows survey estimates of the fraction of women who are sterilized, expressed as a fraction of women who are either sterilized or using oral contraception. Although this information is not accessible in the case of the 1987-89 DHS, the data from two other surveys in the 1990s are shown. The Chimere-Dan study [34], which was conducted in the former Transkei (rural Eastern Cape) in 1994 suggests a lower fraction sterilized than reported in the 1998 DHS [30]. Since the latter survey is more nationally representative, we have used these data in setting the model assumptions about the fraction of non-injectable contraceptive users who are sterilized.

**Table 3.3.4:** Fraction of women using oral contraception and female sterilization, by age

| Age group | Using oral contraception (A) |  | Female sterilization (B) |  | Sterilization fraction B/(A+B) |  |
| --- | --- | --- | --- | --- | --- | --- |
|  | Chimere-Dan (Transkei, 1994)<sup>a</sup> | DHS (SA, 1998)<sup>b</sup> | Chimere-Dan (Transkei, 1994)<sup>a</sup> | DHS (SA, 1998)<sup>b</sup> | Chimere-Dan (Transkei, 1994) | DHS (SA, 1998) |
| 15-19 | 43.2% | 3.5% | 0.0% | 0.0% | 0.0% | 0.0% |
| 20-24 | 37.8% | 9.6% | 0.0% | 0.4% | 0.0% | 4.0% |
| 25-29 | 26.5% | 14.2% | 2.6% | 2.8% | 8.9% | 16.5% |
| 30-34 | 18.7% | 14.6% | 7.8% | 8.4% | 29.4% | 36.5% |
| 35-39 | 18.9% | 10.8% | 6.3% | 17.0% | 25.0% | 61.2% |
| 40-44 | 15.4% | 7.5% | 23.1% | 24.3% | 60.0% | 76.4% |
| 45-49 | 28.6% | 3.8% | 17.9% | 23.6% | 38.4% | 86.1% |

<sup>a</sup> Expressed as a fraction of all women using contraception. <sup>b</sup> Expressed as a fraction of all women.

Figure 3.3.1(a) shows the model estimates of the fraction of women using contraception in 1985, compared with the results of the 1987-89 DHS. The parameters in Table 3.3.1 have been set in such a way that there is reasonably close agreement between the survey and the model. Similarly, Figure 3.3.1(b) shows the fraction of women using contraception who choose to use injectable methods; the model assumptions have again been set to ensure reasonable consistency between the survey and the model.

**Figure 3.3.1:**
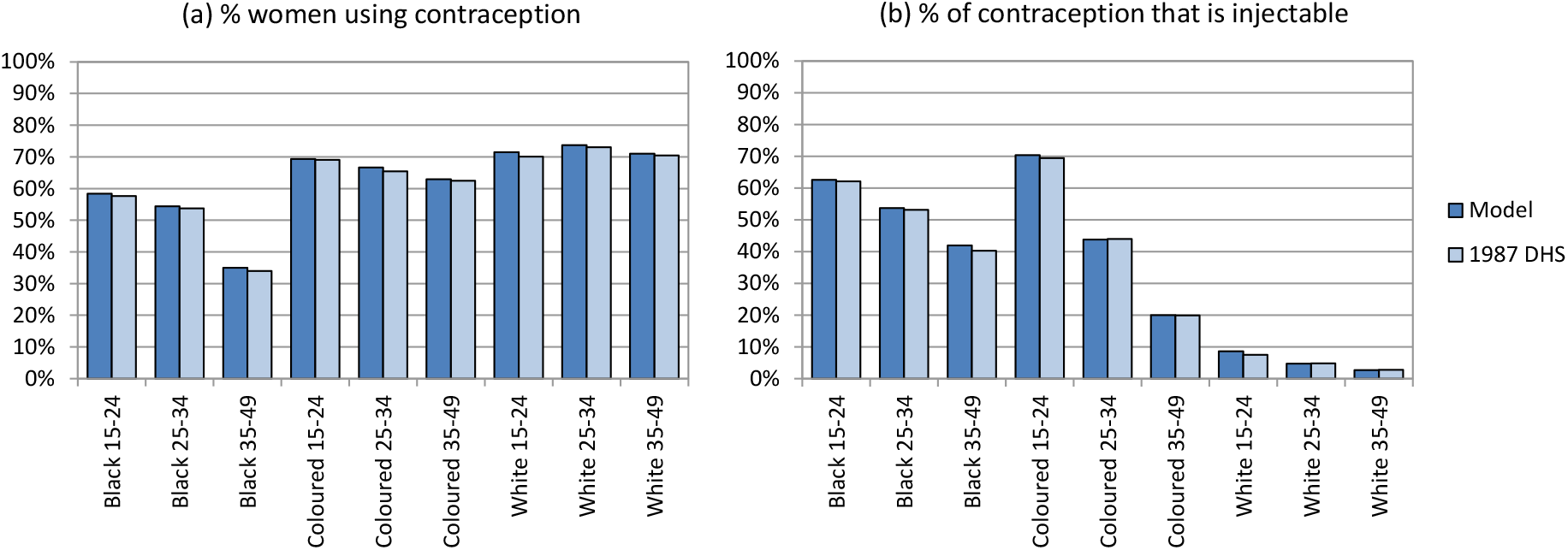
Model calibration to contraception prevalence data from the 1987-89 DHS. Model results are the average from 10 simulations. In both analyses, women were only included if either ever pregnant or ever married (but women who were currently pregnant were excluded). In panel (b), the analysis is further limited to the subset of women who reported using injectables, the pill or sterilization.

#### 3.3.2 Rates of sterilization

Women who have had a previous child, who are sexually active, and who are not currently pregnant or infertile, are assumed to get sterilized at a constant annual rate that depends on their current age. Table 3.3.5 shows the proportions sterilized in the 1998 and 2003 DHSs. The assumed annual rates of sterilization, also shown in the table, have been set so that the model matches the 1998 DHS results. These age-specific rates are multiplied by adjustment factors of 0.6 in black women, 2.8 in coloured women, and 2.8 in white women, based on observed racial differences in the prevalence of female sterilization in the 1998 and 2003 DHSs [37]. Substantially lower levels of sterilization have been measured in the recent 2016 DHS [27], suggesting a trend towards declining rates of female sterilization over time. The model therefore assumes that the rates of sterilization (expressed as a multiple of those in 1997) decline linearly from 1.0 in 1997 to 0.5 in 2007, then remain constant thereafter.

**Table 3.3.5:**
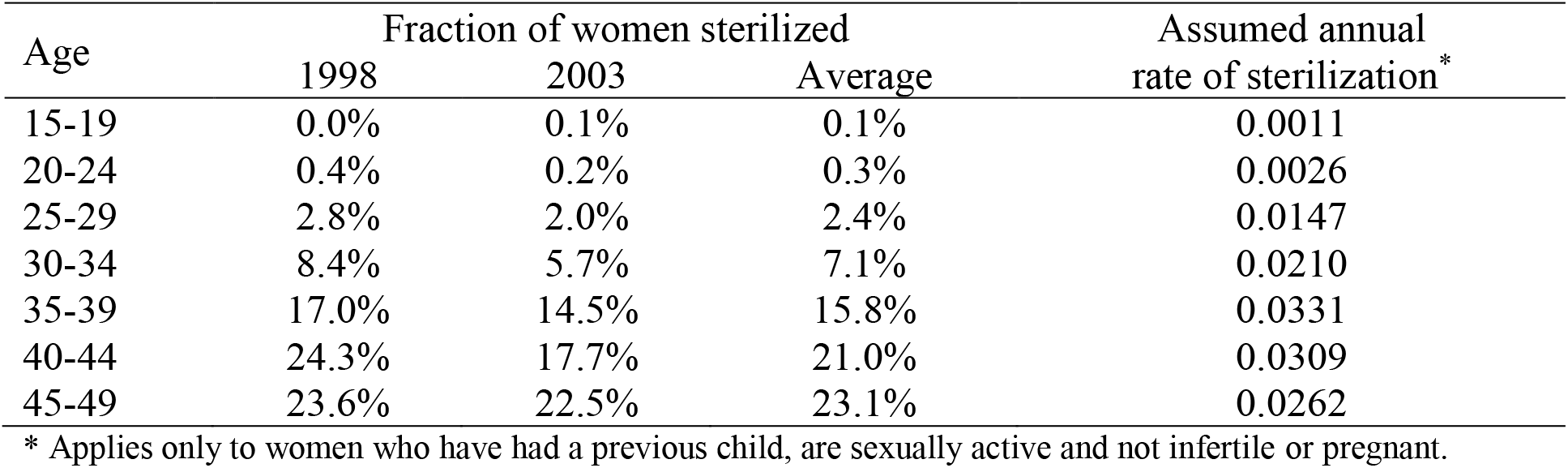
Rates of female sterilization

| Age | Fraction of women sterilized |  |  | Assumed annual rate of sterilization* |
| --- | --- | --- | --- | --- |
|  | 1998 | 2003 | Average |  |
| 15-19 | 0.0% | 0.1% | 0.1% | 0.0011 |
| 20-24 | 0.4% | 0.2% | 0.3% | 0.0026 |
| 25-29 | 2.8% | 2.0% | 2.4% | 0.0147 |
| 30-34 | 8.4% | 5.7% | 7.1% | 0.0210 |
| 35-39 | 17.0% | 14.5% | 15.8% | 0.0331 |
| 40-44 | 24.3% | 17.7% | 21.0% | 0.0309 |
| 45-49 | 23.6% | 22.5% | 23.1% | 0.0262 |

\* Applies only to women who have had a previous child, are sexually active and not infertile or pregnant.

#### 3.3.3 Initiation of hormonal contraception

The model allows for initiation of hormonal contraception (the pill or injectable methods) at a number of time points: at the start of a new relationship, on discontinuation of condom use within an existing relationship, on entry into commercial sex, after birth, or at other times.

##### 3.3.3.1 Initiation at the start of a new relationship

At the time of starting a new sexual relationship, women who were not previously using hormonal contraception are assumed to start using hormonal contraception with a given probability. In women aged 15-19, who were not previously pregnant but have previously used hormonal contraception, and who do not use condoms with their new partner, this probability is set to 0.40 for black women, 0.65 for coloured women and 0.65 for white women. These baseline values have been chosen such that the overall prevalence of contraceptive use by race is consistent with the data in the 1998 DHS (see Figure 3.3.5). The baseline values are adjusted as follows:

- For women who have had a **previous child**, the odds of initiation of hormonal contraception is multiplied by 2.5 in black and coloured women, and by 1.5 in white women. This is based on odds ratios of 2.8-4.8 for black and coloured women in the 1987-89 DHS (comparing women with 1 previous birth to women with no previous birth) and odds ratios of 1.4-1.6 in white women aged 15-34 [28]. The 1987-89 DHS may be biased because it excluded women who had neither been pregnant nor married. In another analysis of data from the Eastern Cape (a mainly black African population), the odds ratio was 1.4 when comparing 1-2 prior births to no prior births, and 3.1 when comparing 3-4 prior births to no prior births [38]. The assumption of a bigger effect of prior pregnancy on contraceptive uptake among coloured and black women when compared to white women is consistent with relatively low rates of teenage pregnancy among white South Africans [39] and the observation that many black South African women only begin to use hormonal contraception after their first birth [31, 40].
- The odds of initiating hormonal contraception are multiplied by ratios of 0.80, 1.00, 1.30, 1.50, 1.15 and 0.45 in the 20-24, 25-29, 30-34, 35-39, 40-44 and 45-49 **age groups** respectively. These odds ratios have been chosen to match the age pattern of contraceptive use observed in the 1998 DHS (see panel (a) of Figure 3.3.3).
- The previously-specified probabilities are assumed to apply in women whose highest **educational attainment** is grade 10. For each additional grade passed, the odds of initiating contraception is assumed to increase by a factor of 1.15. This is higher than the odds ratios estimated in various regression models fitted to South African data sources [28, 35, 38] because the regression models are applied to cross-sectional data, whereas we are interested in modelling the effect of education on the *incidence* of contraceptive use. The odds ratio of 1.15 has been chosen such that the model matches the pattern of contraceptive use by educational attainment in the 1998 DHS (see panel (b) of Figure 3.3.3).
- The previously specified baseline probabilities are assumed to apply to women who have average **fecundability** (fecundability value of 1). Since women who are less fertile are less likely to adopt contraception [41], the baseline probability of adopting hormonal contraception is assumed to be multiplied by the square root of the woman’s fecundability parameter. This means that a woman who is infertile is assumed not to adopt hormonal contraception.
- No adjustment is made in the case of women who have previously been **diagnosed HIV-positive**. Although some studies have shown that there is a substantially higher odds of contraceptive use in HIV-positive women than in HIV-negative women (and especially when comparing women on ART), much of this difference is explained by differences in condom use, and differences in hormonal contraceptive use appear less marked [42, 43]. This effect is usually attributed to more frequent clinic attendance, and advice given to HIV-positive women to make use of family planning services [42]. However, this increase in contraceptive use may be only temporary, as a number of studies suggest that fertility desires and pregnancy rates in HIV-positive women receiving ART increase at longer ART durations [44–46]. On the other hand, a study in KwaZulu-Natal found that HIV-positive women who had been diagnosed were more likely to be using non-barrier contraceptive methods than women who were undiagnosed, and that this difference remained even 4-7 years after ART initiation [47].
- If women use **condoms** with their new sexual partner, it is assumed that there is a reduced probability that they will also use hormonal contraception. Several studies have shown a significant negative association between condom use and use of non-barrier contraceptive methods. For example, in a study of women attending health services in Cape Town and Umtata, Morroni *et al* [48] found that the odds of condom use were reduced significantly in women who were using non-barrier contraception (aOR 0.52, 95% CI: 0.27-0.99). Although there have been a number of other South African studies, they are generally of limited value because they report only univariate associations between condoms and non-barrier methods. In studies of young, sexually active women, univariate odds ratios have been estimated at 0.13 [49] and 0.38 [50], but in other more heterogeneous samples higher odds ratios have been reported [42, 51–53]. The univariate odds ratios in the latter group of studies are likely to be biased, because many of the factors that affect condom use (e.g. age, education) also affect hormonal contraception use, so failure to control for these factors is likely to obscure the true negative association between condom use and hormonal contraception use. We have set the assumed odds ratio for the association between hormonal contraception use and condom use at 0.35, the average of the three odds ratios that we consider to be most reliable [48–50].
- There are assumed to be no differences in levels of condom use between **urban and rural** areas. Although Burgard [28] found that rural residence was associated with lower odds of contraceptive use in multivariate analysis, Kaufman [35] found that urban-rural differences in contraceptive use were not significant after controlling for partner absence (which is relatively more common for women living in rural areas, and which is already controlled for in our model).
- The model allows for changes in uptake of hormonal contraception over **time**, with the extent of these changes depending on age group. For all age groups other than the <25 age group, uptake of hormonal contraceptive use is assumed to have remained constant over time, as there has been relatively little shift in contraceptive use patterns at the older ages, between the 1998 and 2016 DHSs. However, in the <25 age group, uptake is assumed to have been constant only up to 1997. The odds of hormonal contraception uptake in 2005 are assumed to be 0.2 times those in 1997, with the odds ratio declining linearly from 1.0 in 1997 to 0.2 in 2005 (the odds ratio is assumed to remain constant at 0.2 after 2005). These assumptions were made in order to ensure reasonable model fit to the observed changes in contraceptive patterns over time (see Figure 3.3.2). The explanation for the reduction at the young ages is not clear, but may have to do with factors such as the passing of the Choice on Termination of Pregnancy Act in 1996 and the introduction of the Child Support Grant in 1998 [54], as well as increasing emphasis on condoms as a dual protection strategy.

**Figure 3.3.2:**
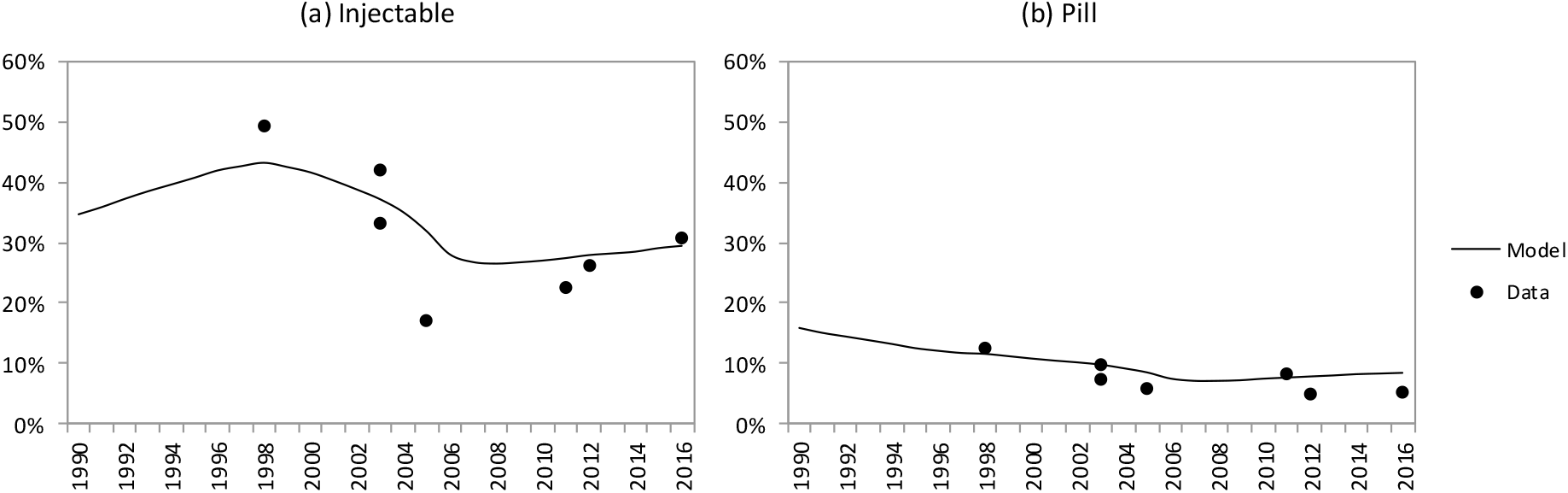
Contraception use among young women (15-24) who are sexually active. Data are from national South African surveys [27, 30, 36, 53, 68–70]. Model estimates represent the average from 100 simulations.

**Figure 3.3.3:**
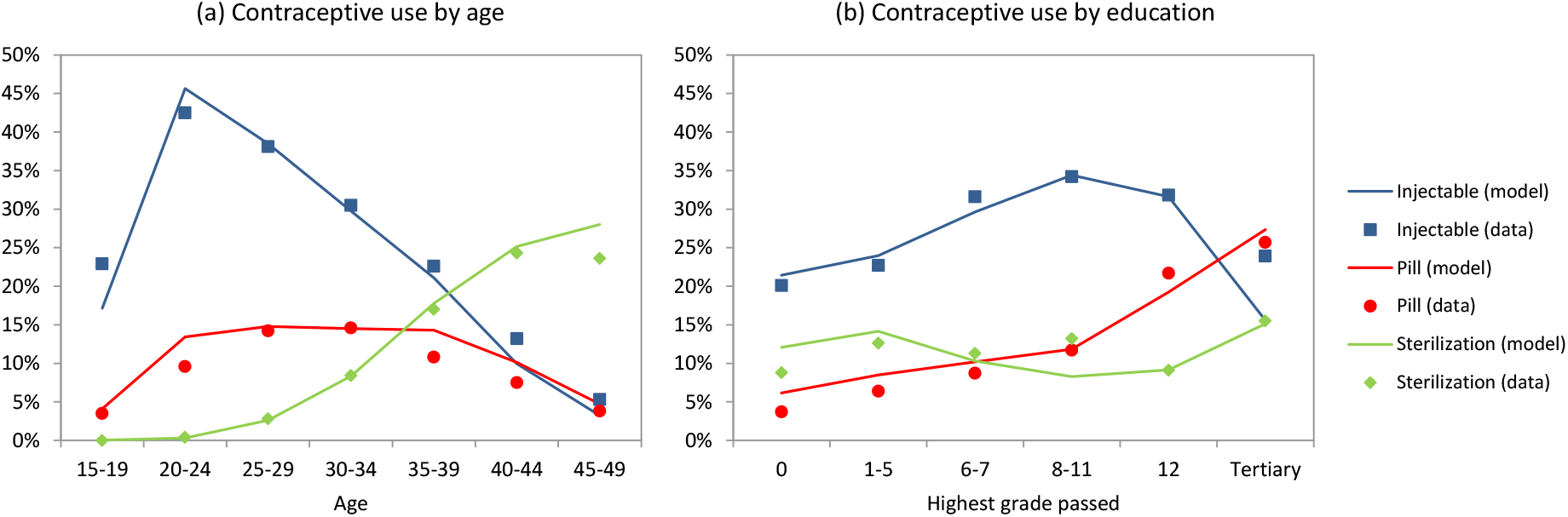
Levels of current contraceptive use in 1998, among all women. Data are from the 1998 DHS, and include all women (regardless of sexual activity or marital status). Model estimates represent the average from 100 simulations.

**Figure 3.3.4:**
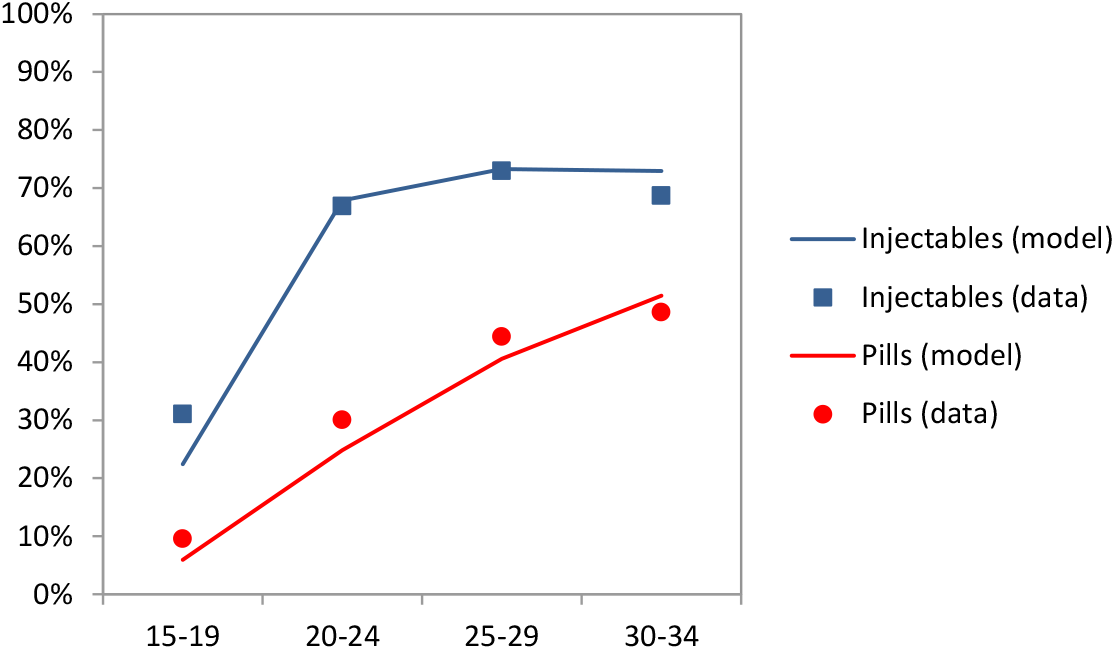
Proportion of women who have ever used hormonal contraception in 1998. Data are from the 1998 DHS. Model estimates represent the average from 100 simulations.

**Figure 3.3.5:**
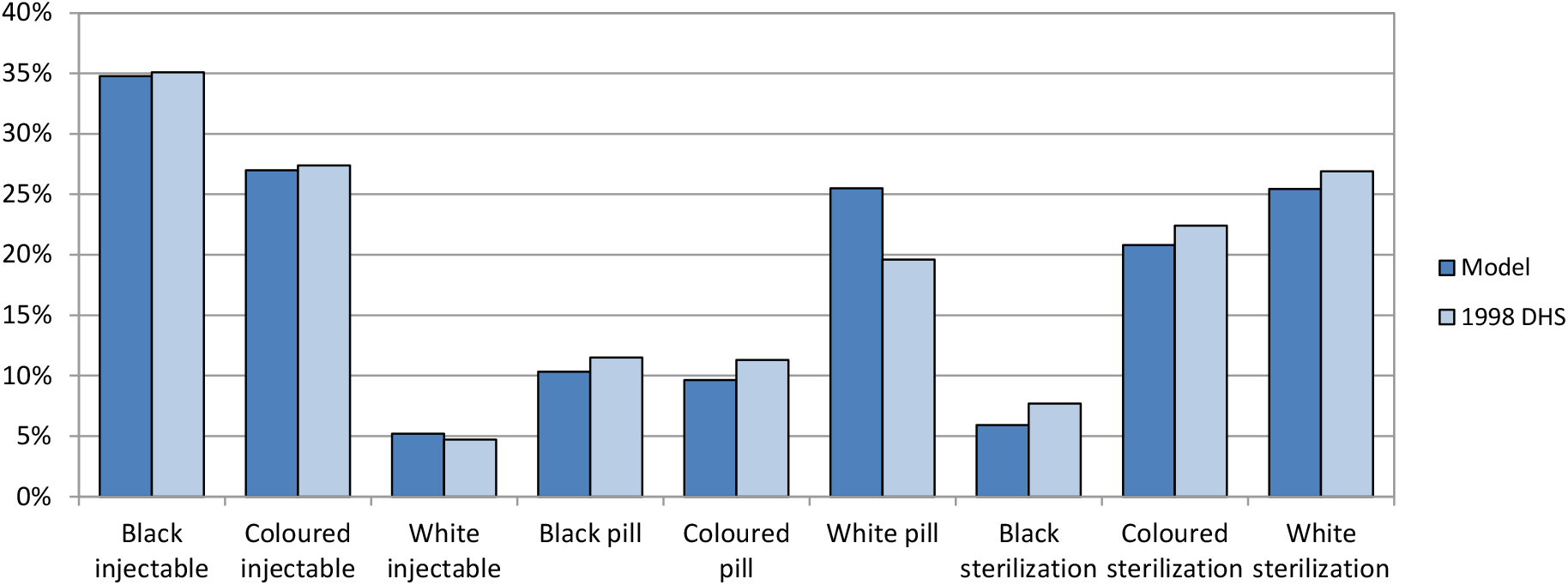
Levels of contraceptive use by race in 1998, among sexually active women. Data are from the 1998 DHS, and include only those women who were sexually active in the last 4 weeks. Model estimates represent the average from 100 simulations; women are only counted if they currently have a sexual partner.

##### 3.3.3.2 Initiation after discontinuation of condom use

If women do not initiate contraceptive use at the start of a new partnership because condoms are being used, we allow for the possibility that a woman may switch to using hormonal contraception at the time that condom use is discontinued (if condom use is discontinued). The probability of initiating hormonal contraception is calculated following a similar approach to that described before. If x is the probability that hormonal contraceptive use is initiated in a relationship that starts *without* condom use (specified previously), and *y* is the probability that hormonal contraceptive use is initiated in a relationship that starts *with* condom use (derived from the odds ratio of 0.35 specified previously), then the probability that hormonal contraceptive use is adopted at the time of condom discontinuation is (*x* – *y*)/(1 – *y*). (This is the conditional probability of adopting hormonal contraception when condoms are not used, given that hormonal contraception was not adopted when the relationship started.)

##### 3.3.3.3 Initiation after starting sex work

Women who enter commercial sex are assumed to have the same probability of starting hormonal contraception as women starting new relationships. For the sake of simplicity, we use the same probability that would apply if the woman was entering into a relationship in which condoms were *not* being used, because even though condom use among sex workers is high, the fraction of sex workers who report hormonal contraception use (51% in Johannesburg, 38% in Cape Town and 32% in Durban, in a recent survey [55]) is similar to the fraction of sexually active black women who reported using hormonal contraception at the time of the 1998 DHS (46%), when condom use was still relatively low [30].

##### 3.3.3.4 Initiation after birth or weaning

Women are also assumed to initiate contraception after birth or weaning with a given probability. It is assumed that if women do not breastfeed, the probability applies at the time of birth. If the woman breastfeeds, the probability applies at the earlier of (a) weaning or (b) six months after birth, in line with South African data showing that uptake of hormonal contraception after birth reaches its peak around 6 months postpartum [56], and in line with international guidance, which notes that breastfeeding is less effective as a contraceptive strategy after 6 months postpartum [41]. The baseline probability of hormonal contraceptive adoption, which applies to women aged 15-19 who are in a sexual relationship, is set at 0.4, based on estimates for postpartum women aged 12-21 in rural Limpopo [40]. Although other studies have recorded proportions of 65-85% [56–58], assuming these high rates would lead to modelled birth intervals longer than those observed (see Figure 3.3.6). The baseline probabilities are adjusted as follows:

- The **age** adjustment factors are set at 1.00 for 15-19, 0.85 for 20-24, 0.70 for 25-29, 0.6 for 30-34, 0.50 for 35-39, 0.40 for 40-44 and 0.30 for 45-49, based on evidence from rural South Africa, which shows that the probability of women using contraception after birth declines with increasing age [40].
- The same **education** adjustment factors are assumed as before, based on the findings of a KwaZulu-Natal study that assessed uptake of dual protection (condoms and non-barrier methods) by 14-weeks postpartum [43].
- The same **fecundability** adjustments as assumed previously are applied, i.e. assuming that women with lower levels of fecundability are less likely to start using hormonal contraception.
- As before, no effect of prior **HIV-positive diagnosis** is assumed (OR 1.0). Although a KwaZulu-Natal study [43] found higher rates of contraceptive uptake postpartum in HIV-positive women, a Cape Town study found no difference between HIV-positive and HIV-negative women in postpartum uptake of contraception [58], and an Eastern Cape study found a significantly lower adoption of pills and injectable methods after birth in HIV-positive women [57]. The higher adoption of hormonal contraception after birth in HIV-positive women in the KwaZulu-Natal study may be partly attributable to the shorter duration of breastfeeding in mothers who have been diagnosed positive, which is accounted for separately in the model (see section 3.4).
- The model allows for changes in hormonal contraceptive uptake over time. In young women (aged <25) the uptake of hormonal contraceptive use is assumed to have declined between 1997 and 2005 (the adjustments are the same as described previously).
- No adjustment is made in relation to **marital status**, since this does not appear to have a significant effect on the adoption of contraception postpartum [43].
- If the woman is **not in a sexual relationship**, the probability is set to zero.

**Figure 3.3.6:**
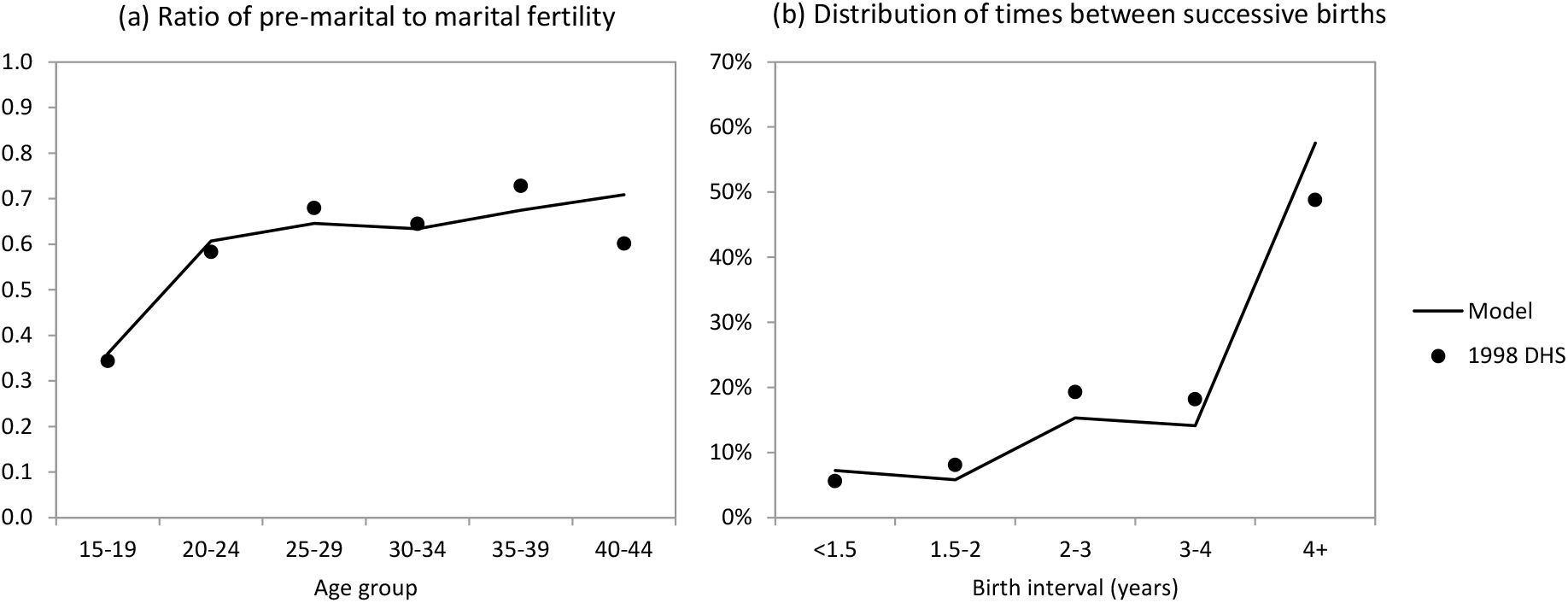
Fertility differences by marital status (a) and birth intervals (b) Data are from the 1998 DHS. Model estimates represent the average from 100 simulations. Ratios in panel (a) were calculated from the 1998 DHS by Tom Moultrie (personal communication).

##### 3.3.3.5 Initiation at other times

Women may adopt hormonal contraception at other times than the formation of a new relationship or following birth. For example, women who are in relationships might start to use hormonal contraception after being exposed to family planning awareness programmes, after family planning services become more accessible, or in response to household economic shocks that make it temporarily undesirable to have further children. Even women who are not sexually active may adopt hormonal contraception, for example to alleviate the symptoms of dysmenorrhoea (painful periods) or menorrhagia (abnormal uterine bleeding) [59]. The model therefore allows for the initiation of hormonal contraception at times other than the formation of a new partnership or following birth. However, there is a lack of information on the frequency with which women adopt hormonal contraception under these circumstances. As noted in section 3.3.1, the prevalence of contraceptive use in 15-24 year-old women who are virgins is low (around 3%), even though this is the age group in which the symptoms of dysmenorrhoea are usually most intense [59]. Based on this, it is arbitrarily assumed that the annual rate of starting hormonal contraception in women aged 15-19, who are not pregnant or sterilized, is 0.01 (excluding starting of contraception for the reasons mentioned previously). The same age adjustments as described previously are assumed to apply at older ages.

#### 3.3.4 Choice of hormonal method

##### 3.3.4.1 First-time hormonal contraceptive users

Among women initiating hormonal contraception for the first time, the probability of choosing injectable methods (over oral contraception) is assumed to depend on race. The probability is set to 0.65 for black women, 0.60 for coloured women and 0.12 for white women, based on fitting the model to 1998 DHS data (see Figure 3.3.5). The following adjustments are made to take into account other covariates:

- As with the initial assumptions in 1985, the odds of choosing injectable contraception (over oral) is multiplied by a factor of 1/0.55 in women who have **previously given birth** [28, 37]. The higher relative use of injectables in women who have previously given birth might be because it is common practice in South African postnatal services to offer injectable methods immediately after delivery [58].
- As before, it is assumed that the odds of using injectable contraception (over oral) are multiplied by a factor of 0.95 for each additional **grade completed**. This is consistent with the range of ORs (0.90-0.98) estimated by Burgard *et al* [28].
- Preference for injectable methods is assumed to **increase over the 1985-95 period**, consistent with a comparison of the 1987-89 DHS and the 1998 DHS, which found that among women using hormonal methods, the OR of injectable use in 1987-89 (relative to 1998) was 0.32 in 15-24 year olds, 0.44 in 25-34 year olds and 0.88 among 35-49 year olds [28]. The baseline probabilities specified previously are assumed to apply in 1995 and all subsequent years. In 1985, the OR for injectable use is multiplied by 0.4, and the ORs that apply in subsequent years are calculated by linearly interpolating between the OR of 0.4 in 1985 and the OR of 1.0 in 1995.
- No **age** effect is assumed.
- Although there is evidence from a study in Johannesburg to suggest that injectable methods are more commonly chosen among women on ART than other women [42], no other studies have confirmed this. It is therefore assumed that method preference is independent of **ART use and HIV status**.

##### 3.3.4.2 Previous users of hormonal contraception

In women who previously used contraception, the probability of adopting a hormonal contraceptive method different from that adopted previously is *p*_1_ × *p*_2_, where *p*_1_ is the probability of using that method if the woman was starting hormonal contraception for the first time (calculated as described previously), and *p*_2_ is set at 0.3. This value of 0.3 has been chosen such that the pattern of cumulative method use, by age, is consistent with data from the 1998 DHS (see Figure 3.3.4).

#### 3.3.5 Discontinuation of hormonal contraception

The model allows for discontinuation of contraception in a number of circumstances. Firstly, all women who were previously using contraception are assumed to stop taking contraception at the time of **falling pregnant** (i.e. in the event of contraceptive failure).

Secondly, women are assumed to discontinue contraceptive use at a constant monthly rate if they are **not currently sexually active**, and not in a relationship. MacPhail *et al* [29] found that among young women aged 15-24, who had been sexually active in the last 12 months, sexual activity in the last month was strongly associated with contraceptive use (aOR 1.5, 95% CI: 1.1-2.0). Similarly, Kaufman [35] found that among women who had ever been married, those who reported that their partner returned home once a month or less frequently were significantly less likely to report use of contraception (aOR 0.79, p < 0.001). Based on these results, we assume a monthly rate of method discontinuation of 0.2 if a woman is not sexually active.

Thirdly, women are assumed to discontinue hormonal contraception if they get **sterilized**.

Fourthly, women are assumed to discontinue contraceptive use at a constant monthly rate if they are currently in a relationship, either because they **desire children** or for **other reasons** (e.g. health reasons, or moving to an area where hormonal contraception is not accessible). We do not model temporary discontinuations in women who want to check that they are still menstruating; although 1998 DHS data suggest that this is relatively common in women using injectable contraception (occurring at a rate of about 0.04 per annum [30]), these interruptions would usually be of short duration and women would be at relatively low risk of falling pregnant during these interruptions (since it usually takes a number of months for the risk of conception to return to normal after injectable contraception is discontinued). The monthly rate of discontinuation in women who are in relationships due to desire for children or other reasons is set to 0.015 (this increases to 0.02 in the case of women who are geographically separated from their partner [35]). This is based on the 1998 DHS, in which 21.7% of women using a modern contraception method other than sterilization reported a break in contraceptive use over the last 12 months, and of these women, 74.8% reported the break was for reasons other than sexual inactivity or wanting to check that they were still menstruating. The implied annual rate of discontinuation is −ln(1 – 0.217 × 0.748) = 0.177, which is equivalent to a monthly rate of 0.015. This rate is slightly lower than estimates from other sources. In a 2006 survey of women attending health facilities in Cape Town and Umtata, Morroni *et al* [48] found that the median duration of hormonal contraception use was around 24 months (which is equivalent to a roughly 29% annual probability of contraceptive discontinuation, higher than the rate observed in the 1998 DHS). In another study of women starting injectable contraception in Gauteng, Beksinska *et al* [60] report that 28% of women discontinued contraception within the first year of use; however, lower rates of discontinuations might be expected at longer durations, and it is also important to note that the authors used a very broad definition of discontinuation (which included women who were more than 2 weeks late in returning for reinjections).

#### 3.3.6 Contraceptive efficacy

Table 3.3.6 summarizes estimates of the relative reduction in the incidence of pregnancy in women using oral contraception and injectable contraception, in African populations. Studies were only included in this analysis if they evaluated the protective effect of contraception in a multivariate model, treating contraception as a time-updated covariate; studies that examine only the effect of baseline contraception use on subsequent incidence of pregnancy [61, 62] are likely to under-estimate the protective effect of contraception. The studies consistently show that injectable contraception has greater efficacy (on average 80%) than oral contraception (56%). However, all three studies included in Table 3.3.6 are from the same group of authors, with substantial overlap in datasets, and the consistency between estimates is therefore not surprising. Contraceptive efficacy appears to be similar in HIV-negative and HIV-positive women. Although there is concern that progestin-based contraceptives may be less effective in women who are receiving efavirenz, this appears to be mainly the case for the implant [63], which we are not currently considering in our model.

**Table 3.3.6:** Percentage reduction in pregnancy incidence due to oral and injectable contraception in African populations

| Study | Location | Population | Contraceptive efficacy (95% CI) |  |
| --- | --- | --- | --- | --- |
|  |  |  | Injectables | Pills |
| Ngure *et al* [64] | 7 African countries | HIV-positive women | 75% (63-83%) | 49% (19-68%) |
|  |  | HIV-negative women | 76% (46-90%) | 36% (0-74%) |
| Reid *et al* [65] | SA, Kenya, Zambia | HIV-negative women | 87% (78-93%) | 69% (54-79%) |
| Pyra *et al* [66] | 7 African countries | HIV-positive women, not on ART | 80% (76-84%) | 64% (53-72%) |
|  |  | HIV-positive women, on ART | 82% (62-91%) | 63% (9-85%) |
| Average |  |  | 80% | 56% |

The average levels of contraceptive efficacy shown in Table 3.3.6 are likely to be under-estimates of the true efficacy parameters. In the American context, Trussell [41] argues that the incidence of pregnancy in women not using contraception cannot be meaningfully compared to that in women using contraception because there is a selection bias determining which women use contraception: women who are less fecund (or who are infertile) are less likely to be using contraception than women who are very fertile. Using estimates of pregnancy incidence from societies in which contraceptive use is rare as the denominator, Trussell estimates efficacy rates of 97% for injectable contraception and 95% for oral contraception. However, if efficacy was calculated using the actual pregnancy incidence rates in US women who report no contraception (around 40% per annum) in the denominator, efficacy estimates would reduce to 88% and 82% respectively. Since similar biases are likely to exist in the African context, it is appropriate to adjust the average efficacy parameters in Table 3.3.6. We set the assumed efficacy rates such that the implied method failure rates are half of those implied by the Table 3.3.6 proportions, i.e. at 90% for injectable contraception and at 78% for oral contraception. The efficacy of injectables may be lower in the South African context than in the US context because South African women are frequently late in returning for reinjections or are not provided with reinjections when they return [32], and often take short breaks in contraceptive use [60].

Condom efficacy is difficult to estimate. Although the studies in Table 3.3.6 suggest condoms have relatively low rates of contraceptive efficacy (15-33%) when compared to the hormonal methods, these analyses all controlled for some measure of unprotected coital frequency in the multivariate analysis, which would dilute the true protective effect of condoms. Trussell [41] also estimates that rates of contraceptive failure are substantially higher for condoms than for hormonal methods. However, Minnis *et al* [67] found that among Zimbabwean women, the incidence of pregnancy was reduced by 81% (95% CI: 74-86%) when women reported consistent condom use – an efficacy comparable to the 80% estimated for injectable methods in Table 3.3.6. In univariate analysis (before controlling for frequency of unprotected sex), Ngure *et al* [64] found that the incidence of pregnancy was similar in women using condoms and women using oral contraceptives. We therefore set the assumed contraceptive efficacy of condoms to 78%, the same as that assumed for oral contraceptives – though the Minnis data suggest a higher efficacy and the Trussell data suggest a lower efficacy.

Women who have been sterilized are assumed to experience a 99.5% reduction in their risk of pregnancy. Although there are no African studies to support this assumption, data from the US suggest efficacy rates of between 99.0 and 99.7%, depending on the comparison group [41].

#### 3.3.7 Calibration to survey data

Figure 3.3.2 shows that the model is roughly consistent with surveys of hormonal contraceptive usage among young sexually active women. Consistent with the findings of Burgard [28], the model estimates that there was a significant shift in method preference between the 1987-89 and 1998 DHSs. The model also estimates a marked reduction in the prevalence of hormonal contraception use in young women following the late 1990s. This is partly due to the assumed reduction in hormonal contraception use by women who use condoms consistently (which started to increase in the late 1990s), but more importantly, it is the result of the assumed reduction in the uptake of hormonal contraception use among young women. The model over-estimates the level of injectable contraceptive use reported in 2005. In the 2005 survey, respondents could only select one contraceptive method, and the absolute number of 15-24 year olds who reported no contraceptive method exceeded the absolute number of 15-49 year olds who reported no contraceptive method [68], suggesting errors in the tabulation of the results. The inconsistency between the model and the data in 2005 could therefore be due to biases or errors in the data.

Figure 3.3.3 shows the relationship between contraceptive use, age and educational attainment, for each of the three major contraceptive methods. In panel (a), there is close agreement between the modelled age pattern and that observed in the 1998 DHS. In panel (b), there is good agreement between the modelled effect of educational attainment on pill use and that measured in the 1998 DHS, and to a lesser extent there is consistency in the case of sterilization and injectable use. However, the model underestimates the level of injectable use among women with tertiary education.

Whereas Figure 3.3.3(a) shows the age pattern of *current* contraceptive use in 1998, Figure 3.3.4 shows how the proportion of women reporting having *ever* used hormonal contraceptive methods differs in relation to age. The model estimates for 1998 are in close agreement with the results of the 1998 DHS, suggesting that the model assumptions about patterns of switching between methods are reasonable. However, the model estimates for women aged 30-34 may be biased, as the model only simulates contraceptive use in 1985 and subsequent years, and some of the women aged 30-34 in 1998 may have been using methods prior to 1985 that were different from those used over the 1985-98 period (the bias would be more substantial in older age groups, and the comparison is therefore not shown for older age groups).

Figure 3.3.5 compares the modelled levels of contraceptive use by race in 1998 with those measured in the 1998 DHS. The model estimates of injectable use are consistent with the data. However, the model consistently under-estimates the rates of sterilization for all race groups, despite the consistency in Figure 3.3.3(a). Although it is concerning that the model is over-estimating rates of pill use among white women in 1998, a higher rate of pill use in white women was observed in the 2003 DHS (27.4%).

Although the previous figures show the model outputs in terms of contraceptive use, it is useful to validate the model by comparing modelled fertility patterns with those observed in surveys. Figure 3.3.6(a) shows the modelled ratio of fertility rates in unmarried women to those in married women over the period from mid-1995 to mid-1998, and compares these to the ratios estimated from the 1998 DHS data. As expected, the ratio is very low in the 15-19 age group because many of the women in this age group who are not married are not yet sexually active. The model results are roughly consistent with the survey results, even though the model makes no explicit assumption about the effect of marital status on contraceptive use. Figure 3.3.6(b) shows the distribution of times between the two most recent births, for women who gave birth in the five years before the 1998 DHS, comparing the survey and model estimates. The model estimates are roughly consistent with the DHS, although the model estimates slightly longer birth intervals than the DHS data suggest.

### 3.4 Fertility

Each woman in the model is assigned a risk of falling pregnant, based on her current sexual behaviour, her current breastfeeding, her use of contraception, her natural fertility level (‘fecundability’) and her HIV stage (if she is HIV positive). This section describes the approach we adopt in modelling each of these effects.

Firstly, we describe the modelling of ‘fecundability’. For all prepubescent girls, the fecundability parameter is set to zero. Similar to the approach in other mathematical models of fertility [71, 72], we assume that the age at which women become fecund is uniformly distributed between ages 12 and 20, and that the age at which women cease to be fecund is uniformly distributed between ages 35 and 50. On becoming fecund, a woman is assigned a fecundability parameter. This parameter is sampled from a gamma distribution with a mean of 1 and a standard deviation of 0.43; the mean is set to 1 so that the fecundability parameter represents the *relative* level of fecundability when compared to the average in fertile women. The standard deviation is based on the coefficient of variation in fertility levels, which was found to be relatively consistent at 0.56 in five different historical populations [73]. In the same study, it was estimated that the variation in fecundability could be decomposed into a ‘persistent’ component and a ‘temporary’ component, the former representing variation between couples that is fairly stable over time, and the latter representing ‘within couple’ variation in birth intervals. The persistent component accounted for 58% of the variation on average (across three historical populations) [73], and since it is the persistent component that we are interested in for the purpose of parameterizing our model, we set the standard deviation to 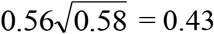.

Secondly, we describe the modelling of breastfeeding and its effect on fertility. Women are assumed not to be at risk of falling pregnant while breastfeeding (lactational amenorrhoea). Although menstruation may resume while a women is breastfeeding (particularly if she is no longer breastfeeding exclusively), we assume for the sake of convenience that there is negligible risk of pregnancy during this time. Assumptions about the duration of breastfeeding are the same as described previously [74, 75], viz. that 86.7% of women breastfeed and that in those women who do breastfeed, the duration of breastfeeding is Weibull-distributed with a median of 18 months and a shape factor of 2 [30]. If women are diagnosed HIV-positive, different breastfeeding parameters are assumed to apply: the probability of breastfeeding is set to 0.44 in the period prior to 2011, but is assumed to increase to 80% after 2011, following the phasing out of free formula milk for HIV-positive mothers [76]. For HIV-diagnosed mothers who choose to breastfeed, the duration of breastfeeding is simulated using a Weibull distribution with a median of 7 months and a shape parameter of 1 [77–79].

Thirdly, we describe the effect of sexual behaviour. A woman’s risk of failing pregnant is assumed to be proportional to her number of current partners (all other things being equal). In the case of sex workers who have no regular partners, we assume that the risk of falling pregnant is the same as that in women with one regular partner. The model also assumes that a woman is not at risk of falling pregnant if her current partner is temporarily absent due to migration or incarceration (in modelling circular migration, we assume that the pregnancy risk is concentrated during the periods in which the migrant partner returns home).

Fourthly, we describe the assumed effect of HIV on fertility. The adjustments to fertility rates in relation to the stage of HIV disease are similar to those assumed in the STI-HIV Interaction model [80]: for women in the acute stage of HIV infection, there is assumed to be no reduction in the fertility rate, but for other untreated HIV-positive women with CD4 counts of ≥350, 200-349 and <200, the fertility rate is assumed to be reduced by 8%, 20% and 27% respectively. In women on ART, the fertility rate is assumed to be 20% *greater* than that in HIV-negative women, based on findings suggesting a significant restoration of fertility after ART initiation [46].

In the interests of simplicity, we model only the incidence of pregnancies that lead to live births. This means that it is possible to work backwards from observed fertility rates to estimate the incidence of pregnancy. The ‘observed’ fertility rates are those estimated by the ASSA2008 model of the South African population, stratified by age, calendar year and race [22]. Suppose that *f_r_*(*x, t*) represents the fertility rate in year *t*, in women of race *r* who are aged *x* at the start of year *t*, as estimated by the ASSA2008 model. Further suppose that 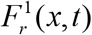 represents the rate of falling pregnant (where pregnancy leads to live birth) in year *t*, in women of race *r* who are aged *x* at the start of year *t*. Note that of women aged *x* at the start of year *t*, who fall pregnant in year *t*, approximately one quarter will deliver in year *t* and the remainder will deliver in year *t* +1 (assuming that conceptions occur uniformly over the period from time *t* to *t* + 1, and assuming the time to delivery is 9 months on average). Hence we can approximate 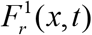 as

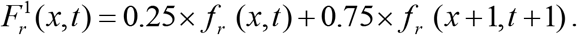

Now suppose that for the *i*^th^ women, who is aged *x_i_* at the start of year *t*, and is of race group *r_i_*, the fecundability parameter is *φ_i_*, the number of partners is *n_i_*, the level of condom use with her *j*^th^ partner is *c_ij_*, the reduction in the risk of conception due to other contraceptive methods is *θ_i_*, and *P_i_* is an indicator of whether the woman is currently pregnant or breastfeeding (1 if currently pregnant or breastfeeding, 0 otherwise). We define 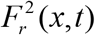 as the rate of falling pregnant (where pregnancy leads to live birth) in year *t*, in women of race *r* who are aged *x* at the start of year *t*, with average fecundability (relative fecundability parameter equal to 1) who have a single sexual partner and who are using no contraception. We relate 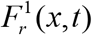 and 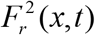 by noting that on average we would expect

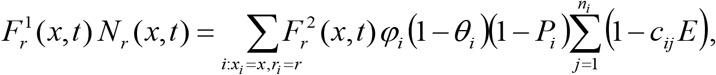

where *N_r_*(*x, t*) is the number of women of race *r* who are aged *x* at the start of year *t*, and *E* is the efficacy of condoms in preventing pregnancy. (For more detail on the assumed efficacy of condoms and other contraceptive methods, see section 3.3.6.) The summation across *i* is a summation across all sexually active women. This equation allows us to approximate 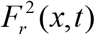:

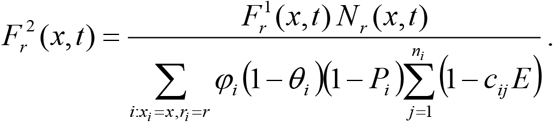

However, this relation will only hold *on average*. Because of stochastic variation, and because of the relatively small number of women in some age-race compartments, these rates may be quite volatile. To stabilize these rates, we define a smoothed set of pregnancy incidence rates, 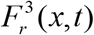. These are calculated as moving averages of the 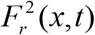 estimates:

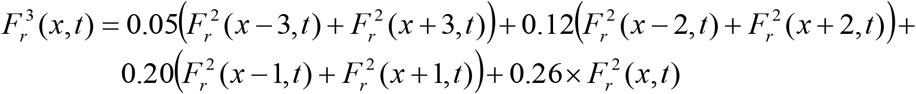

Thus the pregnancy incidence rate that applies to woman *i*, if she is sexually active, is

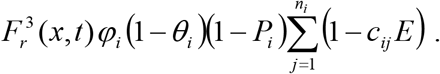

This pregnancy incidence rate is updated at monthly time steps as the woman’s contraceptive use, sexual behaviour and pregnancy/lactation status change. However, the calculations of 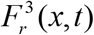 and *φ_i_* are only updated at the start of each year.

### 3.5 Educational attainment

In the mathematical model we assign to each individual two variables to represent their educational attainment. The first variable represents their highest level of completed education; this takes on a value from 0 (no education) to 13 (completed some form of tertiary education), with values of 1 to 12 corresponding to completion of grades 1 to 12. The second variable determines whether the individual is currently either enrolled in school/tertiary study or not enrolled. For the sake of simplicity, we assume that all children who start school do so before age 10, and that children who drop out of school do so only after grade 1 (studies show that the average age of starting schooling in recent cohorts is around 6 years [81], and that rates of dropout prior to grade 5 are negligible [82–84]). The model also ignores the possibility that individuals may temporarily interrupt their schooling; i.e. youth are assumed to remain continually enrolled in some form of educational institution until such time as their education is completed or they drop out permanently.

#### 3.5.1 Age of starting schooling

Current Department of Basic Education policy is that children should enrol in grade 1 at the start of the school year if they are aged 5 and turning 6 before 30 June, or older. However, entry into the schooling system is delayed for some children. The fraction of individuals who enter the formal schooling system has increased over time: data from the 2003-7 General Household Surveys show that the fraction of adults who have completed grade 1 has increased from 97.6% in the 1975-79 birth cohort to 98.9% in the 1985-9 birth cohort [23].

Table 3.5.1 shows the assumed annual probabilities of entering grade 1 at each age (conditional on not having previously started schooling) and the associated cumulative probabilities of having ever enrolled. ‘Age’ here is defined as age last birthday at the middle of the year prior to the start of the school year. Thus all children aged 5 (according to this definition) should enrol in the subsequent school year. These rates are based on the fraction of children at each age who reported being in school, averaged across the March Labour Force Surveys from 2000-2006 [23]. These data show that the fraction currently enrolled increases from age 7 to age 10, which is an indication of delays in school enrolment. For the purpose of these calculations, we assume that there is no dropout prior to age 10, which may lead to some under-estimation of enrolment rates (since some of the incomplete enrolment below age 10 is due to dropout but we are assuming it is all due to late registration). On the other hand, we are using data from the 2000-2006 Labour Force Surveys, which is likely to imply some over-estimation of the enrolment rates that existed in the 1980s and 1990s (since the model is not allowing for changes in the rate of entering grade 1 over time). The two sources of bias are likely to offset one another to some extent.

**Table 3.5.1:**
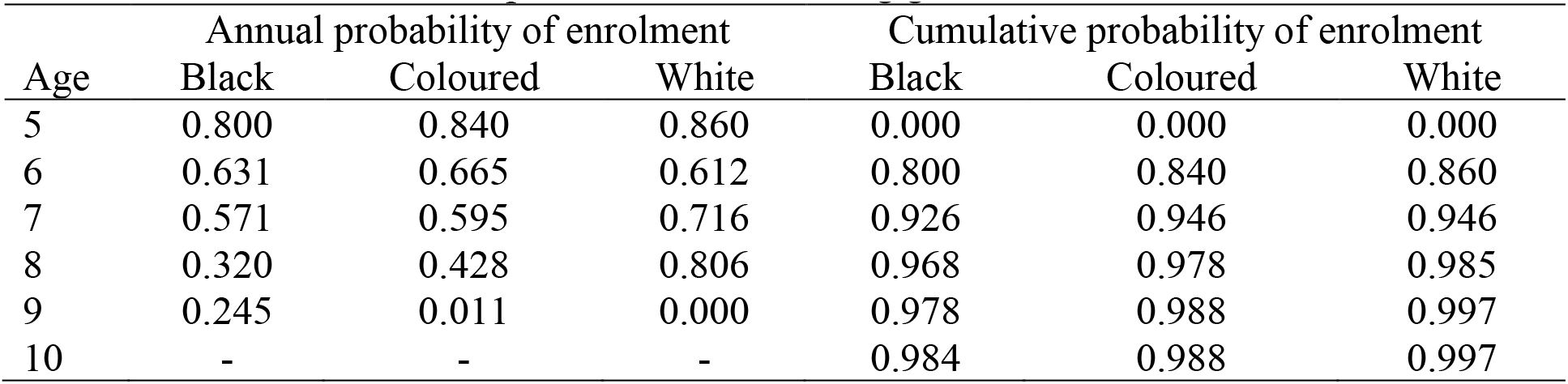
Assumed annual probabilities of entering grade 1 for the first time

| Age | Annual probability of enrolment |  |  | Cumulative probability of enrolment |  |  |
| --- | --- | --- | --- | --- | --- | --- |
|  | Black | Coloured | White | Black | Coloured | White |
| 5 | 0.800 | 0.840 | 0.860 | 0.000 | 0.000 | 0.000 |
| 6 | 0.631 | 0.665 | 0.612 | 0.800 | 0.840 | 0.860 |
| 7 | 0.571 | 0.595 | 0.716 | 0.926 | 0.946 | 0.946 |
| 8 | 0.320 | 0.428 | 0.806 | 0.968 | 0.978 | 0.985 |
| 9 | 0.245 | 0.011 | 0.000 | 0.978 | 0.988 | 0.997 |
| 10 | - | - | - | 0.984 | 0.988 | 0.997 |

#### 3.5.2 Rates of grade repetition

Studies show that rates of grade repetition differ substantially by grade, sex and race. Rates tend to be higher in boys than in girls, although at the more advanced grades, differences between boys and girls are relatively small, perhaps because of pregnancy-related grade repetition once girls become sexually active [81, 84]. Rates of grade repetition also tend to be highest in the grades 10 and 11 and lowest in the late primary phase [81, 84]. Grade repetition is very common in African children, lower in coloured children and lowest in white children [81].

The model assumptions about rates of grade repetition are based on data collected from round 1 of the National Income Dynamics Study (NIDS), conducted in 2008 [81]. In a nationally representative sample of individuals aged 30 and younger, individuals were asked about which grades they had repeated (and how many times). Branson and Lam [81] report the denominators for each grade, broken down by race and sex, as well as the fraction repeating each grade, broken down by race and sex. Because the sample sizes are relatively small in some categories, we fit a logistic regression model to the data. (For the purpose of fitting the model, we have excluded girls in grades 8-12, as much of the repetition in these grades is likely to be associated with pregnancy, which we are accounting for separately.) Table 3.5.2 summarizes the estimated probabilities of repetition that apply to African males (the baseline category). The crude rates from this logistic regression model are adjusted to take into account likely under-reporting of grade repetition. Although the extent of this under-reporting is difficult to quantify, Gustafsson [85] compared the implied levels of grade repetition in the NIDS data when calculated using two different sets of questions about grade repetition (the ‘bottom-up’ approach, based on questions about when schooling was started, is considered more reliable, although it is less complete than the ‘top-down’ approach). On average, the number of grade repetitions derived from the ‘bottom-up’ approach exceeds that derived from the ‘top-down’ approach by 0.5. Of all individuals interviewed, 42% reported at least one grade repetition; if it is conservatively assumed that these individuals reported an average of 2 grade repetitions, then the ratio of the true number of grade repetitions to the reported number of grade repetitions is 1.6 ((0.5 + 2 × 0.42)/(2 × 0.42)). We therefore multiply the crude grade repetition probabilities by factors of 1.6 in order to obtain the adjusted grade repetition probabilities.

**Table 3.5.2:** Probabilities of grade repetition in African males

| Grade | 1 | 2 | 3 | 4 | 5 | 6 | 7 | 8 | 9 | 10 | 11 | 12 |
| --- | --- | --- | --- | --- | --- | --- | --- | --- | --- | --- | --- | --- |
| Crude | 0.123 | 0.087 | 0.101 | 0.090 | 0.075 | 0.067 | 0.074 | 0.112 | 0.105 | 0.227 | 0.180 | 0.117 |
| Adjusted | 0.196 | 0.139 | 0.161 | 0.145 | 0.120 | 0.107 | 0.118 | 0.179 | 0.168 | 0.364 | 0.288 | 0.188 |

These rates are multiplied by factors of 0.48 and 0.14 for coloured and white children respectively, based on the odds ratios estimated by the logistic regression model. The rates are also multiplied by a factor of 0.54 in the case of girls, based on the same logistic regression model, although there is an additional probability of repetition that applies in the case of girls who fall pregnant while in school (see section 3.5.4). However, in the case of African girls in grades 8-12, we obtain the grade repetition rates by multiplying the male rates by an adjustment factor of 0.85 instead of 0.54, as we found that at these higher grades the differences in male and female repetition rates were too small to be explained entirely by teenage pregnancy.

The model does not make provision for changes over time in rates of grade repetition. Although Gustafsson [85] shows that the NIDS data suggest an increase in rates of grade repetition over time, he also notes that this appears to be contradicted by data from several other South African surveys, which suggest a trend towards completion of specific grades at earlier ages.

#### 3.5.3 Rates of school dropout

Assumed average rates of school drop-out, by grade, are estimated from a report commissioned by the Department of Education [23], based on average rates of grade attainment in the cohort of individuals born between 1975 and 1979, as reported in the General Household Surveys (GHSs) of 2003-2006 (very few individuals in this birth cohort would still be in the schooling system by the time of the 2003 survey, and the rates of grade attainment are therefore not affected by survivorship bias). The reported rates are summarized in Table 3.5.3 below. Because these reported rates represent the probabilities of dropping out before reaching the next grade, but our model requires annual probabilities, we divide the reported rates by (1 + the grade-specific probability of repetition) from the previous section in order to arrive at an adjusted average probability that can be applied on an annual basis.

**Table 3.5.3:** Probabilities of dropout by grade

| Grade | Probability of school dropout |  | Assumed annual probability of dropout |  |  |  |  |  |
| --- | --- | --- | --- | --- | --- | --- | --- | --- |
|  | Reported* | Annualized | Black |  | Coloured |  | White |  |
|  |  |  | M | F | M | F | M | F |
| 1 | - | - | 0.0000 | 0.0000 | 0.0000 | 0.0000 | 0.0000 | 0.0000 |
| 2 | 0.003 | 0.0027 | 0.0029 | 0.0025 | 0.0029 | 0.0025 | 0.0010 | 0.0008 |
| 3 | 0.005 | 0.0045 | 0.0049 | 0.0042 | 0.0049 | 0.0042 | 0.0016 | 0.0014 |
| 4 | 0.011 | 0.0100 | 0.0108 | 0.0092 | 0.0108 | 0.0092 | 0.0036 | 0.0031 |
| 5 | 0.016 | 0.0147 | 0.0159 | 0.0137 | 0.0159 | 0.0137 | 0.0053 | 0.0046 |
| 6 | 0.019 | 0.0177 | 0.0191 | 0.0164 | 0.0191 | 0.0164 | 0.0064 | 0.0055 |
| 7 | 0.031 | 0.0287 | 0.0310 | 0.0266 | 0.0386 | 0.0331 | 0.0103 | 0.0089 |
| 8 | 0.052 | 0.0460 | 0.0497 | 0.0426 | 0.0620 | 0.0531 | 0.0166 | 0.0142 |
| 9 | 0.071 | 0.0631 | 0.0681 | 0.0584 | 0.0849 | 0.0728 | 0.0227 | 0.0195 |
| 10 | 0.105 | 0.0821 | 0.0887 | 0.0760 | 0.1106 | 0.0948 | 0.0296 | 0.0253 |
| 11 | 0.155 | 0.1228 | 0.1327 | 0.1137 | 0.1654 | 0.1417 | 0.0442 | 0.0379 |
| 12 | 0.239 | 0.1975 | 0.2133 | 0.1828 | 0.2659 | 0.2279 | 0.0711 | 0.0609 |

\* Probabilities derived from GHS 2003-2007 data for 1975-79 birth cohort [23].

Because these rates derived from the GHSs are not stratified by sex or race, we apply additional adjustments to reflect the effect of these factors. A regression model fitted to data from the National Income Dynamics Study (NIDS) suggests that the baseline rate of school dropout (in African girls in grade 8) is increased by 16.7% in boys and by 24.7% in coloured students [83]. The latter is consistent with data from the Cape Area Panel Study, which show that coloured youth have substantially lower levels of retention in school than black youth [86], in spite of the lower rates of grade repetition noted previously. White youth, in comparison, have very low rates of school dropout [23, 86]. Based on these observations, we assume that

- The average rates of dropout in Table 3.5.3 are multiplied by factors of 1.080 for African males and 0.926 for African females.
- Rates of dropout in coloured children are the same as those in black children up to grade 6, after which coloured dropout rates are 1.25 times those in black children.
- Rates of dropout in white children are one third of the rates in black children [23].

The sex- and race-specific dropout rates are shown in Table 3.5.3. Because the GHS data do not permit an estimate of dropout rates in grade 1, we assume the dropout rate in grade 1 to be zero. Assumed rates of dropout are highest in grade 12, although other studies have estimated relatively low rates of dropout in grade 12, with the highest dropout being in grade 11 [82, 83]. This discrepancy could be due to reporting biases, as it has been noted previously that completion of grade 12 may be over-reported relative to other grades [85].

In setting these assumptions, we have not attempted to adjust for dropout related to pregnancy, although this is allowed for separately in the model (see section 3.5.4). However, pregnancy is not expected to have a major impact on average rates of dropout in girls, because most girls return to school after their pregnancy [82, 87, 88]. This is verified in the model calibration (section 3.5.8), which suggests that only about 15% of girls who drop out of school permanently do so because of pregnancy.

The report commissioned by the Department of Education suggests that over time there has been a decline in rates of school dropout, though this decline has only been noticeable in grades 1-9. This is illustrated in Table 3.5.4, which compares the dropout rates for the 1975-79 birth cohort (from Table 3.5.3) with the rates estimated for other birth cohorts from the same GHS data. The final column shows the average factor by which the grade-specific dropout rates decline, for each 5-year increase in birth cohort. The geometric average of this improvement factor, over grades 2-9, is 0.758 per 5-year increase, which is equivalent to an improvement factor of 0.946 per year (0.758^1/5^).

**Table 3.5.4:** Probabilities of dropout by grade and birth cohort

| Grade | Birth cohort |  |  |  | Average improvement factor* |
| --- | --- | --- | --- | --- | --- |
|  | 1970-74 | 1975-79 | 1980-84 | 1985-89 |  |
| 2 | 0.005 | 0.003 | 0.002 | 0.003 | 0.843 |
| 3 | 0.008 | 0.005 | 0.004 | 0.003 | 0.721 |
| 4 | 0.016 | 0.011 | 0.007 | 0.006 | 0.721 |
| 5 | 0.024 | 0.016 | 0.012 | 0.007 | 0.663 |
| 6 | 0.029 | 0.019 | 0.017 | 0.012 | 0.745 |
| 7 | 0.045 | 0.031 | 0.028 | 0.021 | 0.776 |
| 8 | 0.070 | 0.052 | 0.048 | 0.034 | 0.786 |
| 9 | 0.087 | 0.071 | 0.070 | 0.049 | 0.826 |
| 10 | 0.110 | 0.105 | 0.115 | - |  |
| 11 | 0.157 | 0.155 | 0.161 | - |  |
| 12 | 0.205 | 0.239 | 0.242 | - |  |

\* Calculated as $(r_4/r_1)^{1/3}$ , where $r_4$ is the rate for the 1985-89 birth cohort and $r_1$ is the rate for the 1970-74 birth cohort. Probabilities derived from GHS 2003-2007 data [23].

It is therefore assumed that in grades 2-9, dropout rates in black and coloured children decline by a factor of 0.946 per year. (No decline is assumed for white children, as dropout rates in white children were low to begin with and there is no strong evidence to suggest that they have declined over time.) Suppose that *D_r,g_*(*i*) is the probability of dropout for an individual of race *r* and sex *g*, in grade *i*, from the 1975-79 birth cohort (shown in Table 3.5.3). If it is assumed that these individuals were born in 1977 on average, and that they started school in 1984 on average, then the *D_r,g_*(2) probability applies in 1985. Similarly, if grade repetitions are ignored, the *D_r,g_*(9) probability applies in 1992. For an individual of race *r* and sex *g*, in grade *i* in year *t*, the probability of dropout is assumed to be

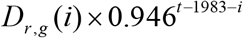

for *i* < 10. For white children and for grades 10-12, no adjustment is made for changes over time (i.e. the probability remains just *D_r,g_*(*i*)).

#### 3.5.4 Pregnancy-related dropout and grade repetition

Studies conducted in black South African girls show that a high proportion of those girls who become pregnant while in school either do not interrupt their schooling or ultimately return to school after temporarily interrupting their schooling. For example, in a study of young women in KwaZulu-Natal, Grant and Hallman [82] found that 27% of girls who fell pregnant while in school did not drop out of school during the year of pregnancy, and that of those who did drop out, 52% of those interviewed at ages 20-24 had returned to school after dropping out. Similarly, Ardington *et al* [87] found that among African girls in rural KwaZulu-Natal who reported falling pregnant while in school, only 34% reported having permanently dropped out of school as a result of this. Another study conducted in Cape Town reported that more than half of African girls who fell pregnant at age 16 or 17 were enrolled in school the following year [88].

In contrast, coloured girls who fall pregnant while in school appear to have a higher rate of dropout [82, 88]. Pregnancy is rare in white teenage girls [39], and there is thus a lack of information on whether the effect of pregnancy on educational attainment in white girls is different from that in other race groups.

In the model, the probability that a girl who has a baby while in school drops out of school permanently as a result of the pregnancy is assumed to be 0.35 if the girl is black or white, and 0.47 if the girl is coloured. The factor of 0.35 is calculated as 0.73 × (1 – 0.52), where 0.73 is the probability of dropout in the year of the pregnancy, and 0.52 is the probability of return to school after initial dropout, both estimated by Grant and Hallman [82]. It is also similar to the probability of 0.34 estimated by Ardington *et al* [87]. The factor of 0.47 is calculated as 0.35 × 1.33, where 1.33 is the relative rate of school dropout in coloured girls who fall pregnant, also estimated by Grant and Hallman [82]. Because the model does not allow for temporary dropout, we approximate the effect of pregnancy-related interruptions in education by assuming that all girls who do not permanently drop out of school repeat their grade in the year following their pregnancy. (In reality some might progress to the next grade despite their pregnancy, but this is to some extent offset by the effect of girls who take more than a year to re-enrol following their pregnancy [82].)

#### 3.5.5 Enrolment into tertiary education

In the model, individuals are classified as having tertiary education if they have completed an undergraduate degree, diploma or certificate. This does not include vocational diplomas and certificates obtained through FET (further education and training) colleges and AET (Adult Education and Training) centres, which are mostly completed by students who have not completed grade 12. FET colleges and AET centres are reported separately from higher education and training institutions (HEIs) in government statistics [89]. Of students who complete grade 12, a fraction is assumed progress to enrol in tertiary education (progression to tertiary education is assumed to be immediate, although in reality some students might only register for tertiary education a few years after completing grade 12). We use statistics from 2001 to set the assumptions about the fraction of students who progress to tertiary education by race, as shown in Table 3.5.5. These estimates are consistent with estimates from the 2002-2007 General Household Surveys, which show that coloured learners who complete grade 12 are slightly less likely to enrol in tertiary education than African learners, while white matriculants are substantially more likely to enrol in tertiary education than black matriculants [90].

**Table 3.5.5:** Progression to enrolment in tertiary education, in 2001

| Race | Number of 19-year olds in 2001<br>[91]* | Fraction completing grade 12<br>[23] | Number enrolled into tertiary education in 2001<br>[91] | % of matriculants who enrol in tertiary education |
| --- | --- | --- | --- | --- |
| African | 746899 | 0.36 | 77895 | 29.0% |
| Coloured | 76488 | 0.43 | 7455 | 22.7% |
| White | 70037 | 0.81 | 31627 | 55.8% |

Values in the final column are calculated as the number of new enrolments divided by the product of the number of 19-year olds and the fraction who complete grade 12. \* Age 19 is used as it is the median age of students entering tertiary education for the first time.

A limitation of the model is that it assumes a constant rate of advancement from completing grade 12 to enrolling in the first year of tertiary education. This is because we lack reliable data on trends in tertiary enrolment by race, although repeating the above calculations with 2007 data yields similar estimates. Another limitation is that the model does not distinguish male and female rates of enrolment in tertiary education, although Branson *et al* have found that among youth who complete grade 12, the fraction who go on to enrol in tertiary education is about 15% higher in females than in males [90].

#### 3.5.6 Dropout and completion of tertiary education

Individuals are classified as having completed grade 12 and as being enrolled at an educational institution for as long as they remain registered for their undergraduate degree, diploma or certificate. The model assumes that at the end of each year of enrolment, there is a probability of drop-out and a probability of graduation, both of which depend on the student’s race. The model does not allow for the probabilities to depend on time since first enrolment. Although it might be considered more realistic to specify a duration dependency, since most degrees are 3-or 4-year qualifications, many certificates and diplomas are for shorter durations, and thus some heterogeneity in completion times might be expected.

The model ceases to classify individuals as enrolled once they have dropped out or graduated. Although students who have graduated may well progress to postgraduate study, this is not modelled, and ‘13’ is thus the maximum level of educational attainment in the model. Students who drop out of tertiary education remain classified as level 12 educational attainment.

Suppose that *G_r_* is the probability that an individual of race *r*, who registers for a tertiary qualification, completes their qualification in a given year. Similarly, suppose that *D_r_* is the probability that an individual of race *r*, who registers for a tertiary qualification, drops out in a given year. Then the average number of times a student registers (or re-registers) is

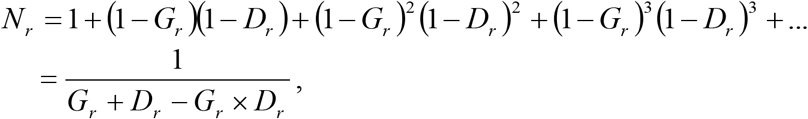

which follows because the first equation is a geometric series. From this it follows that

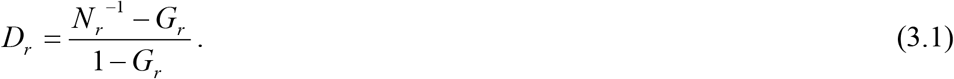

Our approach to setting the model parameters is first to estimate the *G_r_* parameters, then to estimate the *N_r_* parameters, and finally, to calculate the *D_r_* parameters using the above equation. Table 3.5.6 shows the data sources and the steps followed in deriving the parameters. (Although the model does not include Asians as a separate population group, we include them in the table because they are relevant in understanding how the parameters were calculated from the available data sources.) The resulting assumed annual probabilities of graduation are quite low, although substantially higher among white students (0.212) than students of other population groups. Despite this racial disparity, the assumed average number of times enrolled is similar among African, coloured and white students, at around 4 years. This consistency occurs because the effect of higher graduation rates in white students is offset by the effect of lower drop-out rates in white students. The average annual probability of dropout, across all races, is estimated to be 0.104, which is substantially lower than rates of 0.33 that have been reported in the first year of undergraduate study [92]. This is probably because the model assumes a constant annual probability of dropout, but the first year of undergraduate study is the one in which the probability of dropout is highest.

**Table 3.5.6:**
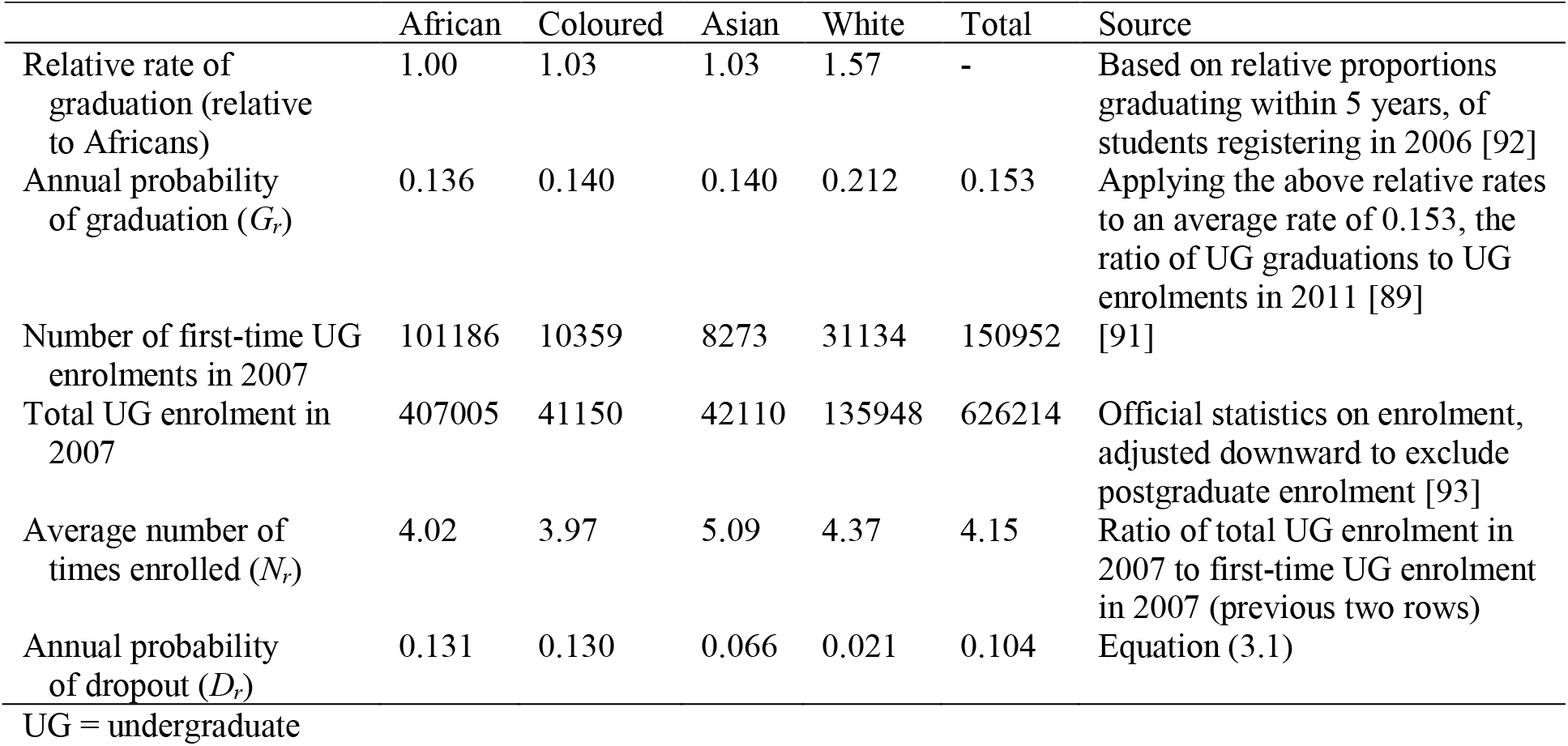
Assumed rates of graduation and dropout from tertiary education

|  | African | Coloured | Asian | White | Total | Source |
| --- | --- | --- | --- | --- | --- | --- |
| Relative rate of graduation (relative to Africans) | 1.00 | 1.03 | 1.03 | 1.57 | - | Based on relative proportions graduating within 5 years, of students registering in 2006 [92] |
| Annual probability of graduation ($G_r$) | 0.136 | 0.140 | 0.140 | 0.212 | 0.153 | Applying the above relative rates to an average rate of 0.153, the ratio of UG graduations to UG enrolments in 2011 [89] |
| Number of first-time UG enrolments in 2007 | 101186 | 10359 | 8273 | 31134 | 150952 | [91] |
| Total UG enrolment in 2007 | 407005 | 41150 | 42110 | 135948 | 626214 | Official statistics on enrolment, adjusted downward to exclude postgraduate enrolment [93] |
| Average number of times enrolled ($N_r$) | 4.02 | 3.97 | 5.09 | 4.37 | 4.15 | Ratio of total UG enrolment in 2007 to first-time UG enrolment in 2007 (previous two rows) |
| Annual probability of dropout ($D_r$) | 0.131 | 0.130 | 0.066 | 0.021 | 0.104 | Equation (3.1) |

UG = undergraduate

A limitation of this approach to parameterization is that it relies on data from different years (2007-2011). However, rates of enrolment, graduation and dropout are unlikely to have changed substantially over this relatively short period, and any bias due to the combination of estimates from different periods is therefore likely to be minimal. Another limitation is that the model assumes rates of graduation and dropout are the same for males and females, although there is some evidence to suggest that male students typically have lower rates of graduation and higher levels of academic exclusion [94].

#### 3.5.7 Baseline educational attainment (1985)

The baseline education profile of the population (mid-1985) is determined using the data from the 1985 census. For each age and race, the census data are used to calculate the fraction of individuals who have attained each level of educational attainment. Each individual in the population in 1985 is then assigned a level of educational attainment by sampling from the distribution of educational attainment levels for the relevant age and race. A limitation of the 1985 census data is that it excludes individuals living in the former ‘homeland’ states, and it might therefore not provide a representative picture of levels of educational attainment in black South Africans.

Another limitation of the 1985 census is that it reports on the fraction of individuals whose highest level of educational attainment is grades 1-3, without reporting the fraction in each individual grade. To overcome this problem, we use the previously-specified assumptions about entry into grade 1, grade repetition and drop-out to calculate the fraction of individuals who we would expect to have completed each grade, for a given age, and use these expected fractions to allocate the reported fraction with completed grades 1-3 to individual grades.

It is also necessary to assign to each individual an indicator of whether or not they are still enrolled in an educational institution. For all individuals over the age of 30, this indicator is set to zero (i.e. all individuals over the age of 30 are assumed to no longer be attending school or university [85]). For individuals aged 30 or younger, the indicator is sampled based on a modelled probability of enrolment that is dependent upon the individual’s age, highest educational attainment and race. The paragraphs below explain how these modelled probabilities are calculated.

Suppose that *D_r_*(*i*) is the probability that an individual of race *r*, who has enrolled in grade *i*, does not enrol in the following year (i.e. it is an annual probability of dropout). Similarly, suppose that *R_r_*(*i*) is the probability that an individual of race *r*, who has enrolled in grade i, repeats their current grade in the following year. These parameters are taken from tables 3.5.3 and 3.5.2 respectively (because we are ignoring sex differences here, we take the average of the male and female rates). Let *p_r_*(*i, x*) be the proportion of individuals of age *x* and race *r* who have grade *i* as their highest level of educational attainment, i.e.

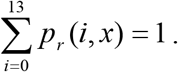

(The proportion *p_r_*(0, *x*) is the fraction of individuals who have never completed grade 1.) We define *π_r_*(*i, x*) to be the proportion of individuals (of age *x* and race *r*, with highest educational attainment of grade *i*) who are still enrolled in an educational institution. It is the *π_r_*(*i, x*) terms that we are principally interested in estimating. These terms can be estimated iteratively by noting that

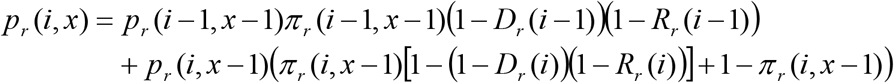

for *i* > 0 and *i* < 13. Following the same logic,

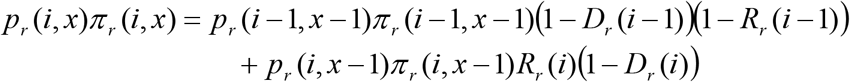

for *i* > 0 and *i* < 12. From this it follows that

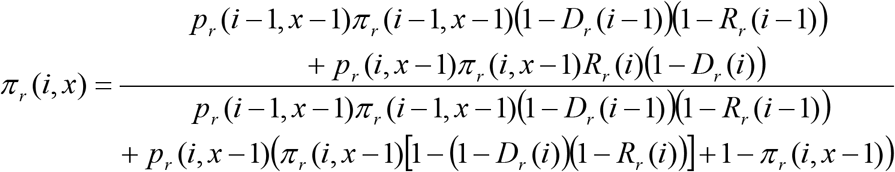

In the case where *i* = 0,

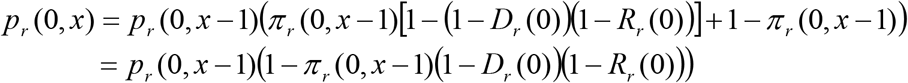

and

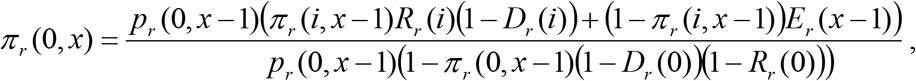

where *E_r_*(*x*) is the probability that a child who is aged *x* and not yet enrolled in school begins school before age *x* + 1. (These *E_r_*(*x*) probabilities have been specified previously in Table 3.5.1.) It is worth noting here the implicit assumption that individuals who pass a particular grade automatically enrol in the next grade the following academic year (in reality some may pass a particular grade but not enrol the following year). It is also implicitly assumed that dropout occurs at the end of each school year (but prior to determining whether the learner passes or fails) [85]. However, in the case of tertiary education, the model assumes that only a fraction of individuals who pass matric (*T_r_*) will enrol in tertiary education the next year (these *T_r_* fractions have been specified in section 3.5.5). Thus for *i* = 13

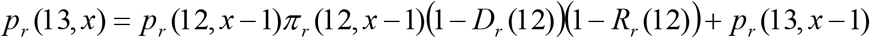

and for *i* = 12

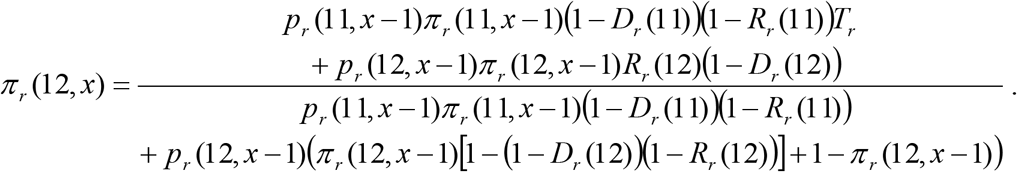

(Note that *π_r_*(13, *x*) = 0, since we cease to classify individuals as being enrolled at an educational institution once they have completed their undergraduate degree.)

A limitation of this approach to setting the baseline educational profile is that it does not take into account sex differences in educational attainment at baseline. Although girls have generally attained higher levels of educational attainment than boys in recent decades, data from older cohorts show the opposite, with men generally having higher educational attainment than women in the same age cohort [81].

#### 3.5.8 Model validation

This section aims to show that with the assumptions described in previous sections, the model yields results consistent with various other data sources. The model results presented in all comparisons are the median from 50 simulations. Figure 3.5.1 shows the fraction of youth (aged 15 to 24) who have completed matric and how this fraction has changed over time. In general the model results are quite consistent with the data from the Labour Force Surveys and the censuses. Among black and coloured youth, levels of grade 12 completion are relatively low, but they have been steadily increasing over the last two decades. Among white youth, levels of grade 12 completion are high and have remained relatively stable.

**Figure 3.5.1:**
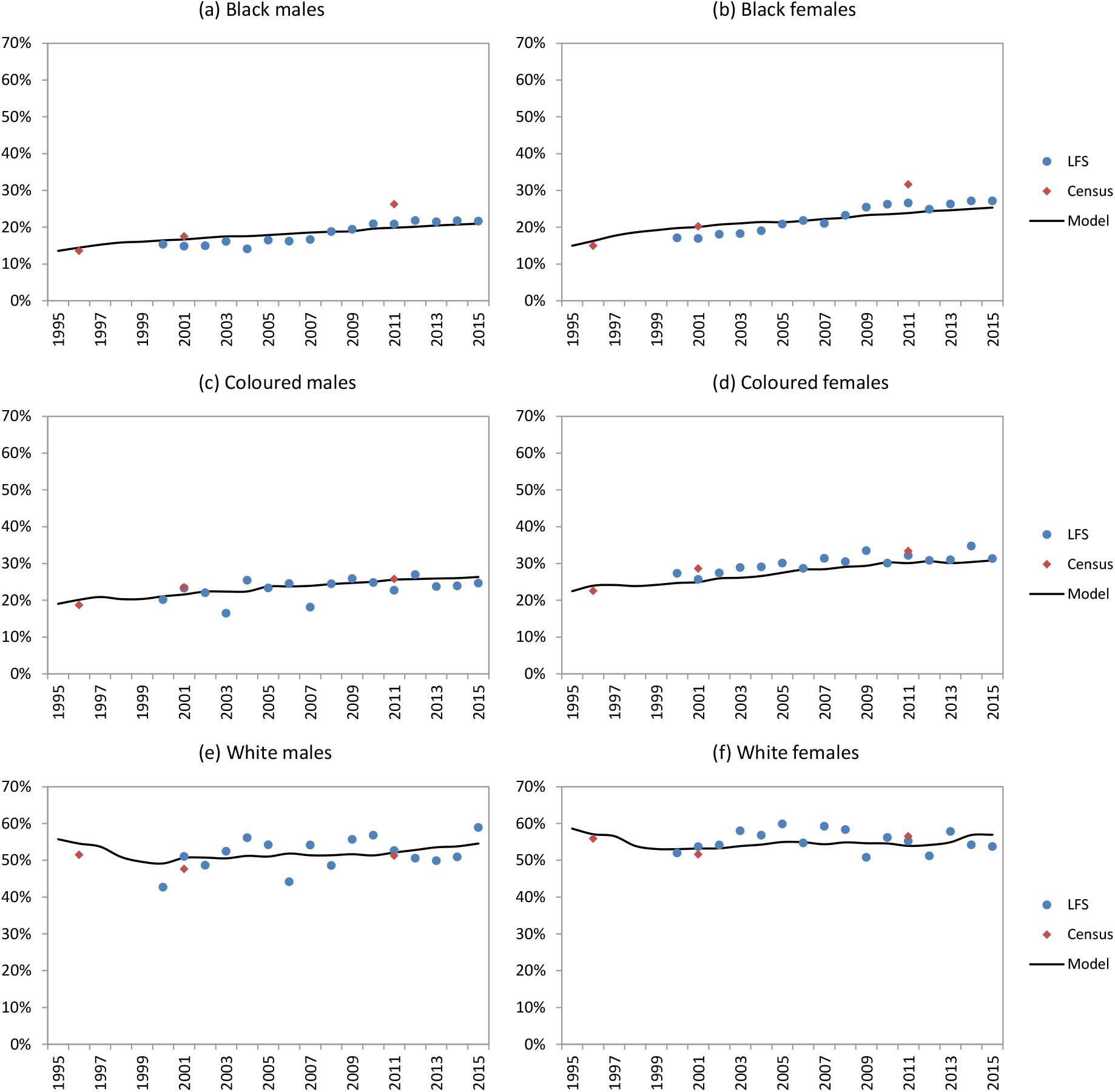
Fraction of 15-24 year olds who have completed grade 12. Labour Force Survey (LFS) results are shown only for the quarter 3 survey conducted in each year, since quarter 3 corresponds most closely to the mid-year time points at which the model results are calculated. Model results represent medians from 50 simulations.

Figure 3.5.2 compares the model estimates of the fraction of youth who are enrolled in school (excluding tertiary education) with 2001 data sources, and shows how this fraction varies in relation to age and race. In general, the model results are fairly consistent with the 2001 data, although it is clear that the model slightly under-estimates the proportion of African youth who were still in school in the 18-20 age range, and to a lesser extent the same is true among white youth. This could potentially be due to overstatement of current enrolment, which appears to be a particular problem in grade 12 (i.e. youth are reported as being in grade 12 when in fact they are out of school) [85]. Alternatively, the model assumptions about grade repetition might be too low. It is also possible that the model assumptions about rates of first enrolment in grade 1 may be too high (since the model slightly exaggerates enrolment at ages 7-9), although changing these assumptions would only partially resolve the problem of low enrolment at ages 18-20.

**Figure 3.5.2:**
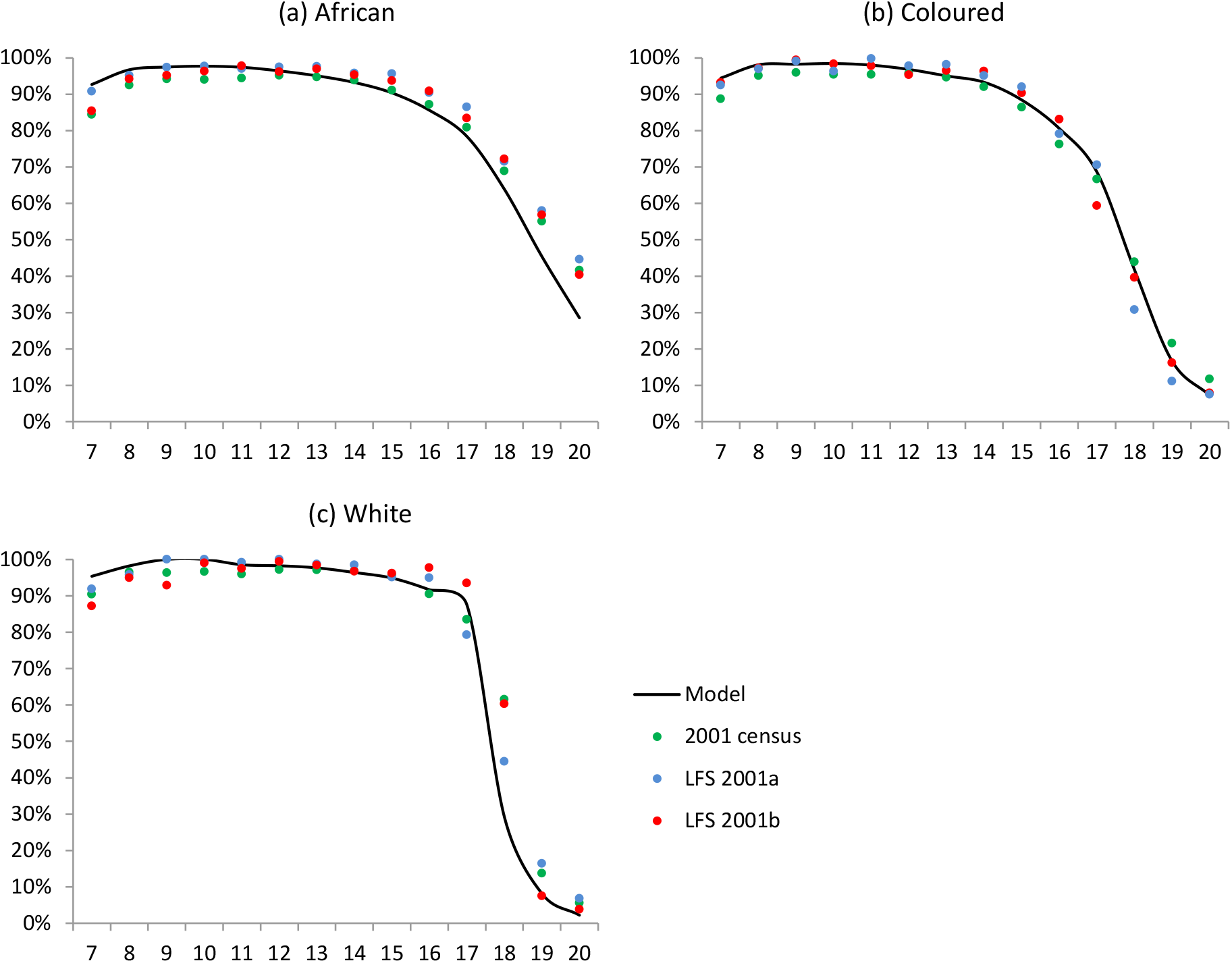
Fraction of youth enrolled in school, by age. Model results for 2001 are compared to data from the 2001 census, and the Labour Force Surveys conducted in March and September of 2001 (LFS 2001a and 2001b respectively) [23].

Figure 3.5.3 shows the fraction of the 1975-79 birth cohort with different levels of educational attainment (for the purpose of this analysis, individuals with tertiary education are included in the grade 12 category). Strictly speaking, this is not a validation, since we have used the same data in setting the model assumptions about rates of school dropout. Nevertheless, Figure 3.5.3 does confirm that the model produces results consistent with the General Household Surveys conducted between 2003 and 2006, for the 1975-79 birth cohort.

**Figure 3.5.3:**
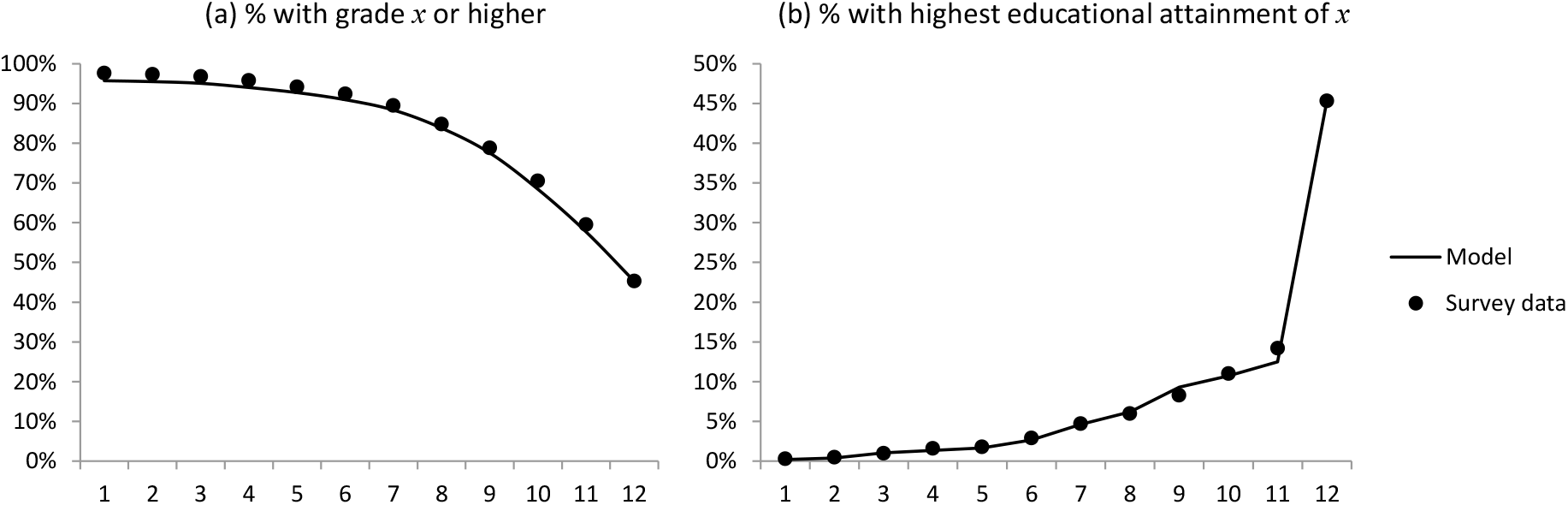
Levels of educational attainment in the 1975-79 birth cohort. Survey data are from the 2003-2006 General Household Surveys, as analysed previously [23].

Figure 3.5.4 compares the fractions of individuals who have completed grades 7 and 12 in different age cohorts, as recorded in the 2007 Community Survey and as estimated by the model in 2007. The model and the 2007 Community Survey are reasonably consistent when considering the fraction of the population who have completed grade 7, but there is less consistency when considering the fraction who have completed grade 12. At ages 45 and older, the model estimates a lower fraction who have completed grade 12 than the survey. This could possibly be because the model does not take into account the long-term survival advantage associated with higher education; individuals with lower levels of educational attainment are more likely to have died and are thus less likely to be represented in older cohorts. Alternatively, there may be problems with the assumptions about the baseline educational profile in 1985, which are derived from the 1985 census data. In the younger age cohorts (ages 25-24), the opposite problem occurs, with the model over-estimating the fraction who have completed grade 12 when compared with the data from the 2007 Community Survey. The explanations for this discrepancy are unclear, but it is worth noting that the 2007 Community Survey data at these young ages appear inconsistent with the 2003-2006 General Household Survey data for similarly-aged cohorts (Figure 3.5.3).

**Figure 3.5.4:**
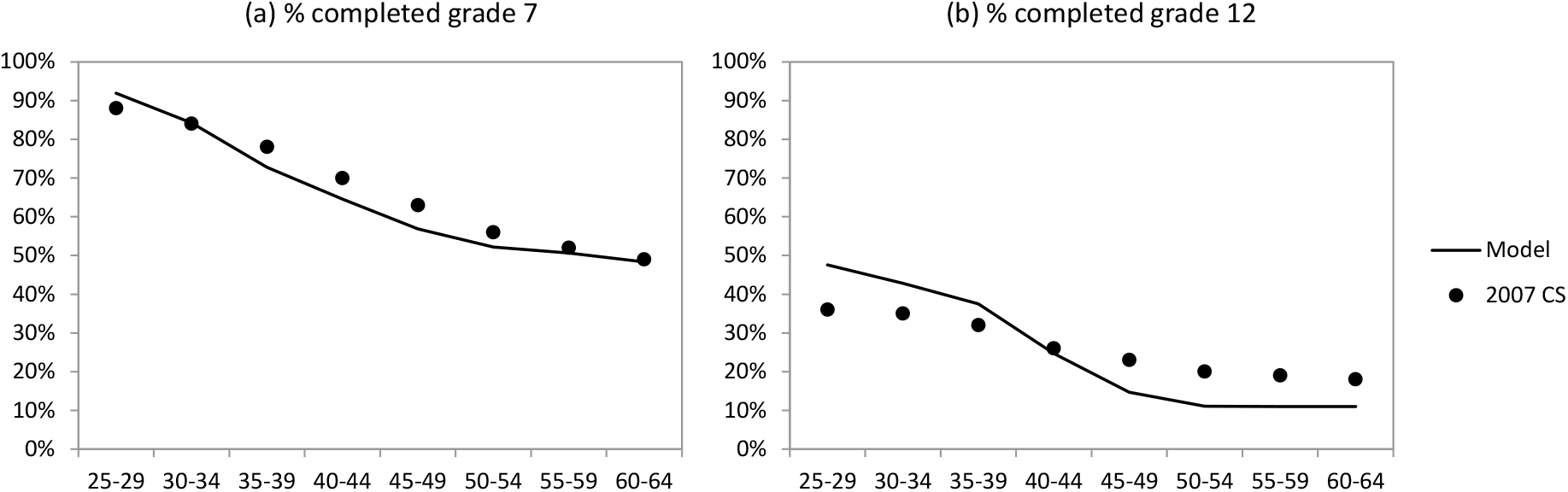
Levels of educational attainment in different birth cohorts, 2007. Data are from the 2007 Community Survey (CS), as analysed previously [81].

Finally, Figure 3.5.5 compares the gross enrolment ratio (GER) for tertiary education, defined as the ratio of the number of enrolments into tertiary education in a given year (including repeat enrolments) to the size of the population aged 20-24. Although the model considers only undergraduate enrolments, the model estimates have been adjusted upward based on estimates of the fraction of all enrolments that are postgraduate enrolments [93]. Despite this upward adjustment, the model estimates of the GER are somewhat below the recorded GERs, especially among coloured and white youth.

**Figure 3.5.5:**
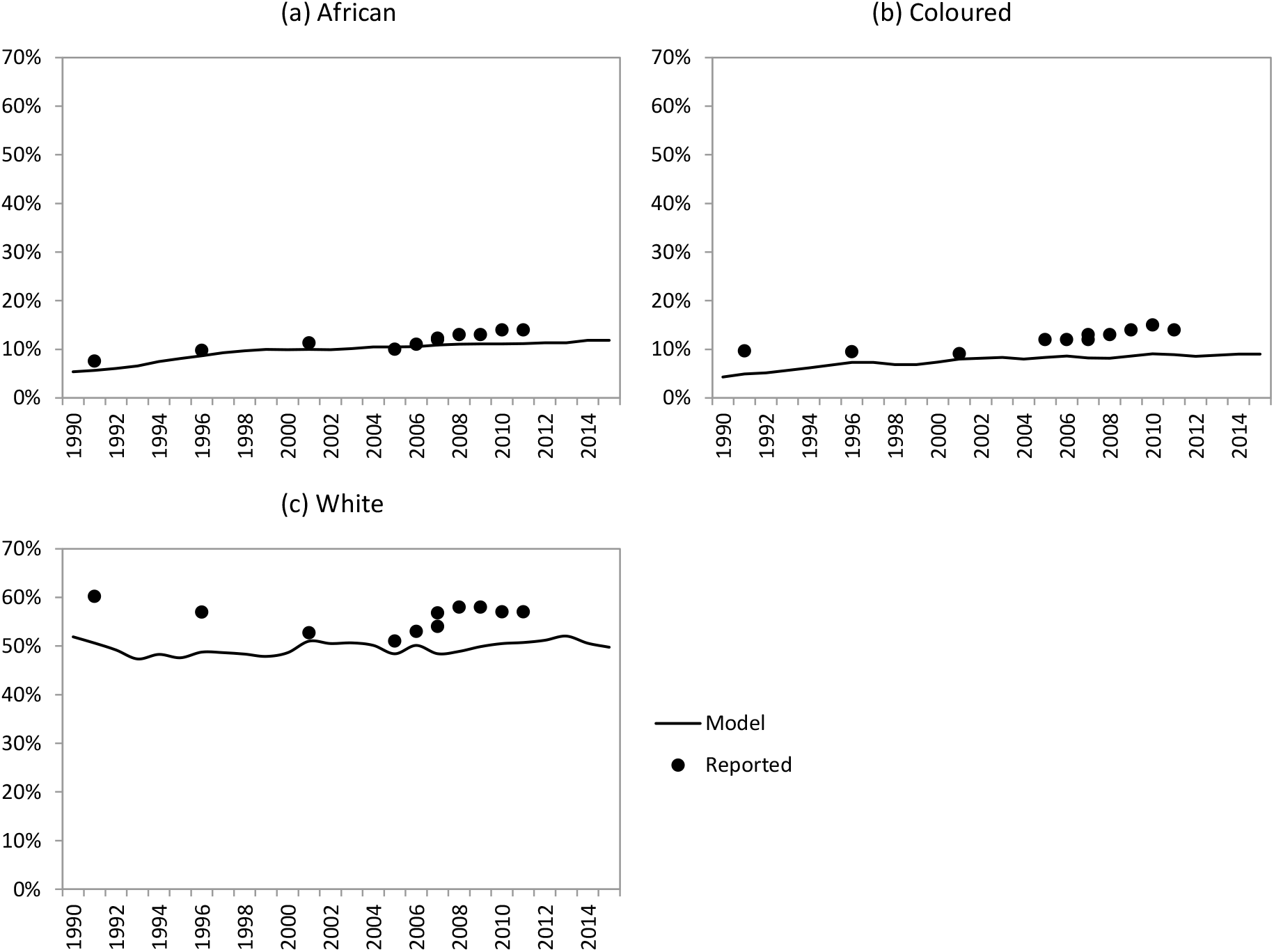
Tertiary education gross enrolment ratio. Reported ratios are derived from censuses conducted in 1991, 1996 and 2001 and the 2007 Community Survey [91], and from routine estimates from the Council for Higher Education over the 2005-2011 period [92].

### 3.6 Urbanization and urban-rural migration

The sections that follow describe the modelling of migration between urban and rural areas and the effects of urbanization and migration on sexual behaviour. It is worth noting that the current model simulates only internal migration, i.e. migration within South Africa. The model does not simulate migration out of South Africa or migration into South Africa. This means that although the model is representative of the South African population at the start of the simulation in 1985, it becomes less representative of the demographic profile of the South African population in recent years.

#### 3.6.1 Modelling the initial fraction of the population that is urbanized

The initial fraction of the population that is urbanized is calculated from the 1985 census data. The fraction is specified separately for each five-year age group, each sex and each population group. In the case of black South Africans, we also specify the proportion separately for different levels of educational attainment. For other population groups, the proportion urbanized is close to 100%, so the additional accuracy gained from calculating the rates separately for each education category is negligible.

Tables 3.6.1 and 3.6.2 shows the assumed urban proportions at the start of the simulation, in 1985, based on the 1985 census data. The urban fraction is highest among whites, lower among coloureds and lowest among black South Africans. However, the 1985 census excluded individuals living in the former homeland states, which were predominantly rural and occupied almost entirely by black Africans. This means that the urban fractions shown for black South Africans in Tables 3.6.1 and 3.6.2 are likely to be over-estimates, particularly among women, who had relatively low levels of employment in the former ‘white’ South Africa (pass laws did not apply to children under the age of 16, and the bias is therefore likely to be less significant at the younger ages). To overcome this problem, we multiply the urban fractions for black South African women (aged 10 or older in 1985) by a constant scaling factor of 0.85, to make allowance for the under-representation of rural black South African women in the census. This scaling factor was chosen to achieve consistency with the observed age and sex pattern of urbanization over time (see Figure 3.6.4). The observed patterns of urbanization by educational attainment suggest that urbanization is strongly associated with higher educational attainment. Among coloured and white South Africans, the fraction urbanized is slightly higher among women than in men, whereas among black South Africans, the urban fraction tends to be higher in men than in women.

**Table 3.6.1:** Fraction of the male population living in urban areas in 1985

| Age group | Highest grade passed (black Africans) |  |  |  |  | Tertiary | Coloured | White |
| --- | --- | --- | --- | --- | --- | --- | --- | --- |
|  | 0 | 1-5 | 6-7 | 8-11 | 12 |  |  |  |
| 0-4 | 0.2960 | - | - | - | - | - | 0.7604 | 0.8770 |
| 5-9 | 0.2215 | 0.3424 | - | - | - | - | 0.7602 | 0.8850 |
| 10-14 | 0.1189 | 0.2880 | 0.4503 | 0.5500 | - | - | 0.7770 | 0.8972 |
| 15-19 | 0.1571 | 0.2546 | 0.3742 | 0.5139 | 0.6346 | - | 0.7820 | 0.9160 |
| 20-24 | 0.2904 | 0.3952 | 0.5025 | 0.5911 | 0.6607 | 0.7686 | 0.7674 | 0.9094 |
| 24-29 | 0.3290 | 0.4425 | 0.5504 | 0.6712 | 0.7065 | 0.7865 | 0.7660 | 0.8924 |
| 30-34 | 0.3331 | 0.4686 | 0.5907 | 0.7158 | 0.7469 | 0.8004 | 0.7604 | 0.8932 |
| 35-39 | 0.3315 | 0.4777 | 0.6154 | 0.7467 | 0.7750 | 0.8042 | 0.7626 | 0.8920 |
| 40-44 | 0.3302 | 0.4957 | 0.6258 | 0.7555 | 0.7883 | 0.7947 | 0.7498 | 0.8866 |
| 45-49 | 0.3259 | 0.5068 | 0.6329 | 0.7644 | 0.7954 | 0.8134 | 0.7632 | 0.8758 |
| 50-54 | 0.3320 | 0.5389 | 0.6541 | 0.7708 | 0.7702 | 0.7890 | 0.7584 | 0.8710 |
| 55-59 | 0.3275 | 0.5566 | 0.6651 | 0.7707 | 0.7595 | 0.7895 | 0.7662 | 0.8624 |
| 60-64 | 0.3021 | 0.5417 | 0.6410 | 0.7502 | 0.7288 | 0.8588 | 0.7518 | 0.8632 |
| 65-69 | 0.2846 | 0.5062 | 0.6110 | 0.7124 | 0.7449 | 0.7444 | 0.7356 | 0.8646 |
| 70-74 | 0.2500 | 0.4867 | 0.5653 | 0.6707 | 0.6583 | 0.7059 | 0.7306 | 0.8688 |
| 75-79 | 0.2400 | 0.4618 | 0.5473 | 0.6342 | 0.6532 | 0.7895 | 0.7464 | 0.8776 |
| 80-84 | 0.2245 | 0.4443 | 0.4908 | 0.5851 | 0.6613 | 0.8000 | 0.7236 | 0.8954 |
| 85+ | 0.2090 | 0.3850 | 0.4718 | 0.5870 | 0.6129 | 1.0000 | 0.7244 | 0.8846 |

Source: 1985 census.

**Table 3.6.2:** Fraction of the female population living in urban areas in 1985

| Age group | Highest grade passed (black Africans)* |  |  |  |  |  | Coloured | White |
| --- | --- | --- | --- | --- | --- | --- | --- | --- |
|  | 0 | 1-5 | 6-7 | 8-11 | 12 | Tertiary |  |  |
| 0-4 | 0.2957 | - | - | - | - | - | 0.7586 | 0.8808 |
| 5-9 | 0.2214 | 0.3475 | - | - | - | - | 0.7638 | 0.8892 |
| 10-14 | 0.1122 | 0.2850 | 0.4398 | 0.5483 | - | - | 0.7800 | 0.9050 |
| 15-19 | 0.1037 | 0.1979 | 0.3427 | 0.5092 | 0.6358 | - | 0.7966 | 0.9364 |
| 20-24 | 0.1448 | 0.2452 | 0.4168 | 0.5926 | 0.6770 | 0.7424 | 0.7946 | 0.9288 |
| 24-29 | 0.1720 | 0.3127 | 0.5140 | 0.7005 | 0.7288 | 0.8441 | 0.8080 | 0.9048 |
| 30-34 | 0.1897 | 0.3565 | 0.5713 | 0.7556 | 0.7566 | 0.8750 | 0.8048 | 0.9020 |
| 35-39 | 0.1887 | 0.3848 | 0.6038 | 0.7767 | 0.7703 | 0.8824 | 0.8090 | 0.8984 |
| 40-44 | 0.2056 | 0.4223 | 0.6299 | 0.7932 | 0.7654 | 0.8587 | 0.8000 | 0.8906 |
| 45-49 | 0.2088 | 0.4556 | 0.6509 | 0.7980 | 0.7638 | 0.8452 | 0.8170 | 0.8844 |
| 50-54 | 0.2197 | 0.4960 | 0.6713 | 0.8046 | 0.7612 | 0.7624 | 0.8210 | 0.8818 |
| 55-59 | 0.2098 | 0.4995 | 0.6609 | 0.7881 | 0.7632 | 0.7833 | 0.8332 | 0.8870 |
| 60-64 | 0.2138 | 0.4736 | 0.6148 | 0.7446 | 0.7343 | 0.8125 | 0.8168 | 0.8938 |
| 65-69 | 0.2229 | 0.4932 | 0.6093 | 0.7238 | 0.6842 | 0.6250 | 0.8116 | 0.9022 |
| 70-74 | 0.2276 | 0.5079 | 0.5955 | 0.7045 | 0.7813 | 0.6667 | 0.8216 | 0.9124 |
| 75-79 | 0.2287 | 0.5027 | 0.5831 | 0.6854 | 0.6119 | 0.8000 | 0.8346 | 0.9210 |
| 80-84 | 0.2145 | 0.4695 | 0.5613 | 0.6811 | 0.6111 | 0.6667 | 0.8158 | 0.9322 |
| 85+ | 0.2149 | 0.4242 | 0.5501 | 0.6745 | 0.6522 | 1.0000 | 0.8308 | 0.9298 |

Source: 1985 census. \* Urban proportions for black South Africans are adjusted by a multiple of 0.85 to make allowance for exclusion of the homeland states from the census data.

#### 3.6.2 Rates of urban-to-rural and rural-to-urban migration

Rates of movement between urban and rural areas can be calculated using data from the 2011 census, which asked all individuals if they had moved to their current location from a different location over the last 10 years. Consistent with the approach adopted by Ginsburg *et al* [95] in their analysis of the same data, we limit the analysis to individuals who reported having moved in the last 5 years (any time from 2007 to 2011), to minimize recall bias that may arise when evaluating migration over longer time intervals. A challenge when using these data is that the 2011 census only established whether individuals were in an urban or rural location at the time of the census, and not five years previously. However, the census did collect information on the individual’s previous municipality if they reported having moved, and from this it is possible to approximate whether the individual was previously in an urban or rural area. For the purpose of this analysis, all municipalities in which the fraction of the population urbanized was more than 50% in 2011 were classified as urban, and the remainder were classified as rural. Individuals who migrated from an urban municipality were assumed to have migrated from an urban area, and individuals who migrated from a rural municipality were assumed to have migrated from a rural area. Although this approach is crude, the majority of municipalities are either more than 80% rural or more than 80% urban, so there is relatively little scope for misclassification. Applying this approach to the 2011 population, 90.4% of the population that is classified urban is truly urban, and 85.1% of the population that is classified rural is truly rural.

Figure 3.6.1 shows the resulting estimates of the probabilities of rural-to-urban and urban-to-rural migration, over the 5 years prior to the 2011 census, by age and sex. (Estimates are not shown for children under the age of 10 because the migration question was only asked of individuals aged 10 and older.) Individuals who were in their 20s at the time of the 2011 census reported the highest rates of rural-to-urban migration, and in adults, migration rates were consistently higher in men than in women (panel (a) of Figure 3.6.1). Rates of urban-to-rural migration were substantially lower than rates of rural-to-urban migration, but were less sharply peaked at the young ages (panel (b) of Figure 3.6.1).

**Figure 3.6.1:**
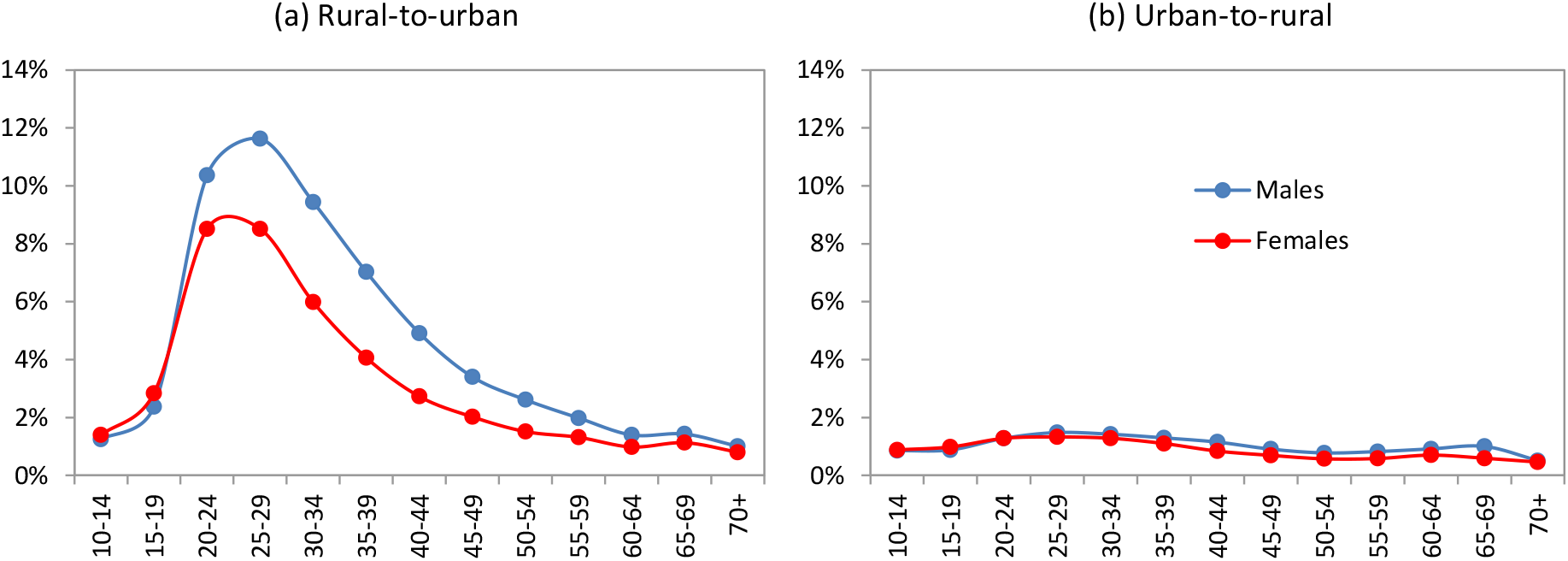
Probability of migration in the 5 years before the 2011 census, by sex and age at the time of the 2011 census

Table 3.6.3 shows that the fraction of adults who report having migrated over the last 5 years has remained relatively stable over time, at around 12% (though higher among white South Africans). Although the measure is not defined consistently across censuses, this suggests that rates of migration have been relatively stable over time. The fraction of the population that is urbanized has changed linearly over the 1985-2001 period [24], which also suggests relatively stable rural-to-urban and urban-to-rural flows.

**Table 3.6.3:** Fraction of individuals who report having migrated over the last 5 years

| Population group | Migration period |  |  |  |
| --- | --- | --- | --- | --- |
|  | 1975-1980 | 1992-1996 | 1996-2001 | 2007-2011 |
| Black African | 9% | 9% | 11% | 12% |
| Coloured | 10% | 9% | 13% | 11% |
| White | 25% | 21% | 26% | 26% |
| Total | 13% | 11% | 12% | 13% |

Source: Estimates for the first 3 periods are from Kok and Collinson [24]. Estimates for 2007-2011 are the author's own calculations, based on the 2011 census data. In the first 2 periods, the unit of analysis when defining migration is the magisterial district, while in the 2001 census, the unit of analysis was the 'main place', and in the 2011 census the unit of analysis was the local municipality.

Logistic regression models have been fitted to the 2011 census data to assess the effects of educational attainment and race on rates of migration (Table 3.6.4). In the case of rural-to-urban migration, the logistic regression models differ somewhat from the results in Table 3.6.3, suggesting higher relative rates of urban-to-rural migration in white and coloured South Africans. However, in the case of urban-to-rural migration, white rates are similar to black rates, while rates are significantly lower among coloured South Africans. Rural-to-urban migration tends to be more frequent in more educated individuals. However, urban-to-rural migration is only weakly affected by educational attainment; urban dwellers with tertiary education are 1.28 times as likely to make an urban-to-rural migration as urban dwellers with no education.

**Table 3.6.4:** Logistic regression estimates of the effects of race and educational attainment on probabilities of rural-to-urban and urban-to-rural migration, in the last 5 years

|  | Rural-to-urban migration<br>Adjusted OR (95% CI) | Urban-to-rural migration<br>Adjusted OR (95% CI) |
| --- | --- | --- |
| Highest grade passed |  |  |
| None | 1 | 1 |
| 1-5 | 0.86 (0.80-0.92) | 1.11 (1.01-1.21) |
| 6-7 | 1.10 (1.03-1.18) | 1.03 (0.94-1.12) |
| 8-11 | 1.51 (1.41-1.62) | 0.86 (0.78-0.94) |
| 12 | 1.71 (1.59-1.84) | 0.89 (0.81-0.97) |
| Tertiary education | 2.60 (2.42-2.79) | 1.28 (1.17-1.40) |
| Race |  |  |
| Black African | 1 | 1 |
| Coloured | 2.53 (2.45-2.62) | 0.15 (0.15-0.16) |
| White | 5.06 (4.99-5.14) | 1.06 (1.04-1.08) |

Both regression models control for age and sex (coefficients not shown). Models are applied only to individuals who were aged 30 or older at the time of the 2011 census, since most of these individuals would have completed their education at the time of their migration (if we included individuals at younger ages, we may introduce confounding due to individuals migrating for the purposes of education).

An alternative approach to calculating rates or rural-to-urban migration is to compare the proportions urbanized in two successive censuses, using a ‘synthetic cohort’ approach. For the purpose of this analysis, we focus on the 1996 and 2001 censuses. Suppose that the data from the first census are used to calculate 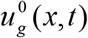, the fraction of individuals of sex *g*, aged *x* in year *t*, who are urbanized, and the data from the second census are used to calculate 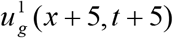, the fraction of individuals of sex *g*, aged *x* + 5 in year *t* + 5, who are urbanized. Then the ‘net probability’ that an individual living in a rural area at the time of the first census moves to an urban area by the time of the second census is

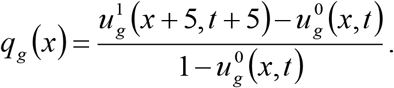

This is not a probability in the strict sense, because the numerator is net of urban-to-rural migration, and can thus be negative. The 5-year migration probabilities calculated using this formula (Figure 3.6.2) appear very different from those in Figure 3.6.1. The net probabilities of rural-to-urban migration in young adults are roughly two times higher than those suggested by the Figure 3.6.1 data, and the net probabilities of urban-to-rural migration around the retirement ages are also much higher than the Figure 3.6.1 data suggest (especially in men). The reason for the discrepancies is not clear. Although it is possible that the differences in rates might be attributable to changes in urban-rural migration flows over time (Figure 3.6.1 being based on 2011 census data, and Figure 3.6.2 being based on 1996 and 2001 census data), this is not supported by the data in Table 3.6.3, which suggest relatively stable migration patterns over time. A more likely explanation is that the lower migration rates in Figure 3.6.1 are the result of under-reporting of past moves in the 2011 census, possibly due to recall bias (the synthetic cohort approach, in contrast, does not rely on self-reported data). For example, of individuals who reported having moved from rural municipalities to urban municipalities in the last 5 years, at the time of the 2011 census, 14% reported having moved in 2007 and 33% reported having moved in 2011, which suggests relative under-reporting of moves of longer duration. It is also possible that the synthetic cohort approach may be biased because of the implicit assumption of a closed population; if there is net international in-migration and in-migrants are settling mainly in urban areas, the closed population assumption may lead to over-estimation of rural-to-urban flows. However, we consider the synthetic cohort approach a more reliable approach, as it leads to model estimates more consistent with census data (as demonstrated below).

**Figure 3.6.2:**
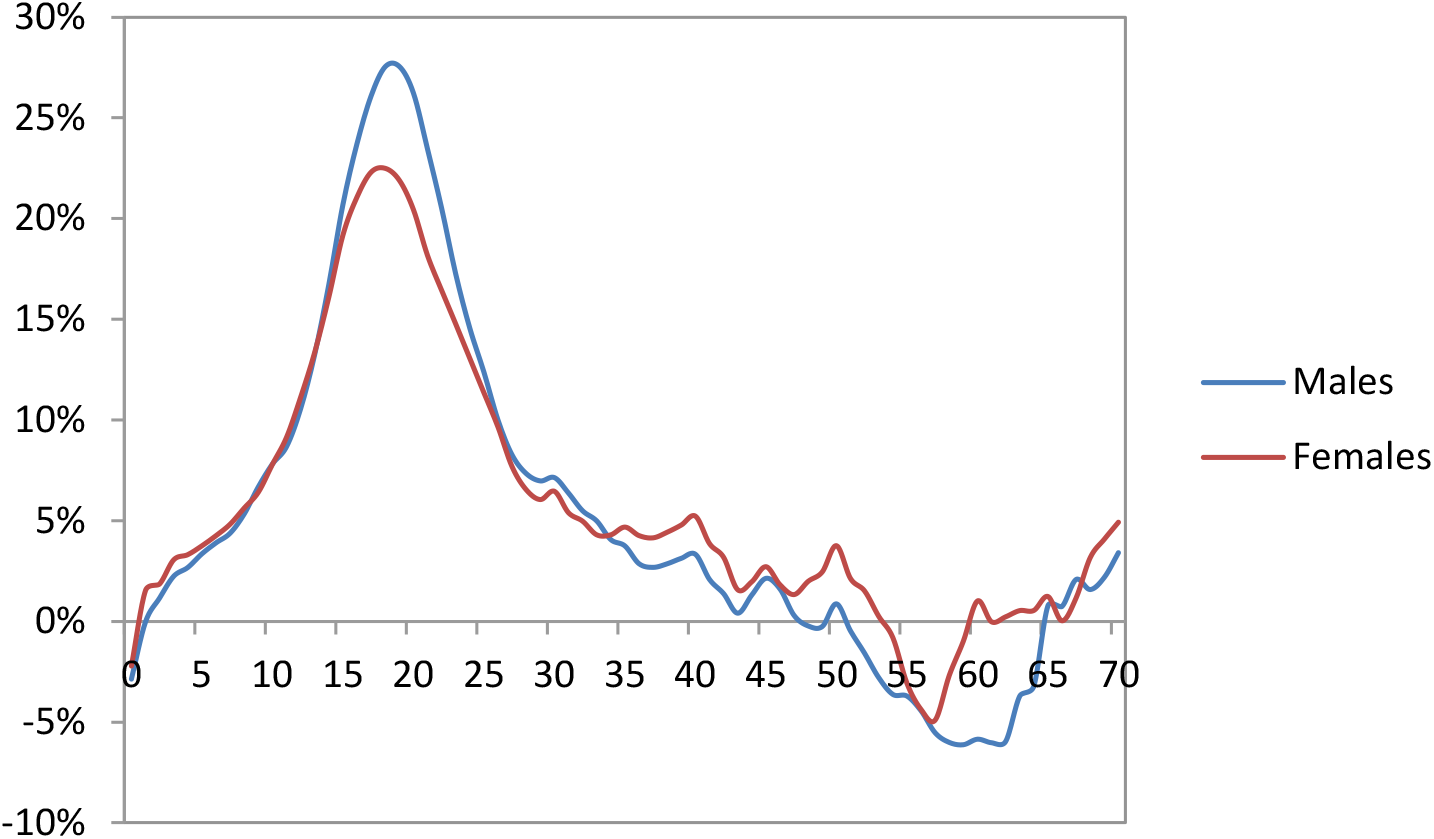
Net probabilities of urban-to-rural migration over the 1996-2001 period, by sex and age at the time of the 1996 census

Suppose we define *μ*_1_(*x, g*) to be the average annual rural-to-urban migration rate for individuals of sex *g*, aged *x* to *x* + 4, and we define *μ*_2_(*x, g*) to be the corresponding urban-to-rural migration rate. Then

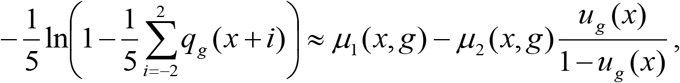

where *u_g_*(*x*) is the proportion of *x* to *x* + 4 year olds who were living in urban areas at the start of the inter-census interval. Because there are two unknowns in this equation (*μ*_1_(*x, g*) and *μ*_2_(*x, g*)), we approximate *μ*_1_(*x, g*) by fixing the *μ*_2_(*x, g*) at the values estimated from the 2011 census data (Figure 3.6.1), but subject to the constraint that *μ*_1_(*x, g*) cannot be less than 0.0016 (the minimum annual rural-to-urban migration rate estimated from the 2011 census data). In other words,

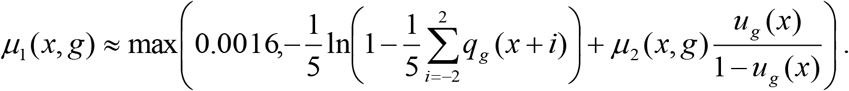

The *μ*_2_(*x, g*) values are then updated to take into account the constraint, using the estimated values of *μ*_1_(*x, g*). The resulting estimates of the rural-to-urban and urban-to-rural migration rates are shown in Table 3.6.5. The only age groups in which the urban-to-rural rates are increased relative to those estimated from the 2011 census data are the 0-4 age group (probably a reflection of children born in urban areas being sent to be looked after by grandparents or other relatives in rural areas) and the 55+ age group. Based on the stability of migration rates over time (Table 3.6.3), these urban-rural migration rates are assumed to remain constant over time.

**Table 3.6.5:**
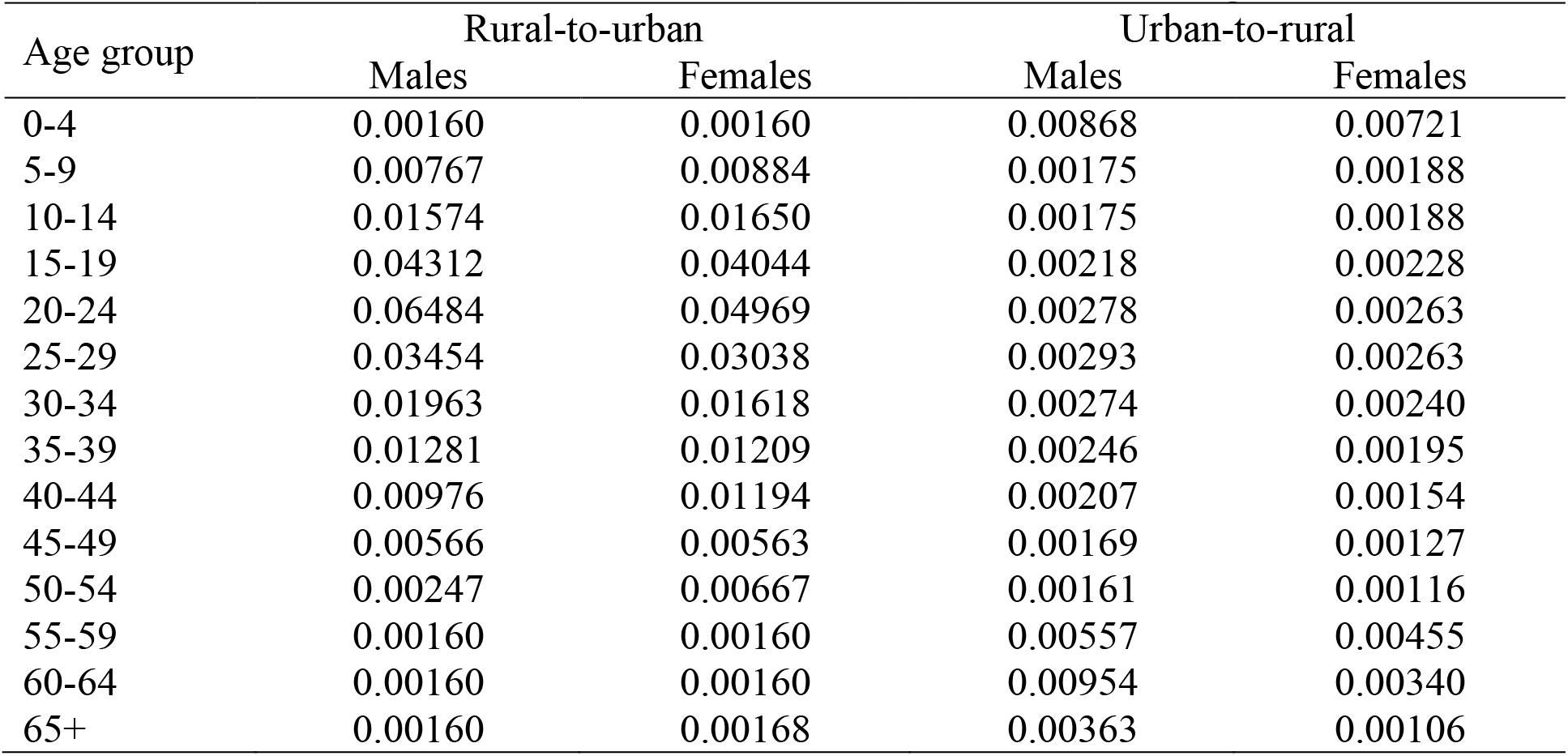
Assumed annual rates of rural-to-urban and urban-to-rural migration

| Age group | Rural-to-urban |  | Urban-to-rural |  |
| --- | --- | --- | --- | --- |
|  | Males | Females | Males | Females |
| 0-4 | 0.00160 | 0.00160 | 0.00868 | 0.00721 |
| 5-9 | 0.00767 | 0.00884 | 0.00175 | 0.00188 |
| 10-14 | 0.01574 | 0.01650 | 0.00175 | 0.00188 |
| 15-19 | 0.04312 | 0.04044 | 0.00218 | 0.00228 |
| 20-24 | 0.06484 | 0.04969 | 0.00278 | 0.00263 |
| 25-29 | 0.03454 | 0.03038 | 0.00293 | 0.00263 |
| 30-34 | 0.01963 | 0.01618 | 0.00274 | 0.00240 |
| 35-39 | 0.01281 | 0.01209 | 0.00246 | 0.00195 |
| 40-44 | 0.00976 | 0.01194 | 0.00207 | 0.00154 |
| 45-49 | 0.00566 | 0.00563 | 0.00169 | 0.00127 |
| 50-54 | 0.00247 | 0.00667 | 0.00161 | 0.00116 |
| 55-59 | 0.00160 | 0.00160 | 0.00557 | 0.00455 |
| 60-64 | 0.00160 | 0.00160 | 0.00954 | 0.00340 |
| 65+ | 0.00160 | 0.00168 | 0.00363 | 0.00106 |

The ‘base’ rates shown in Table 3.6.5 are adjusted to reflect the effects of educational attainment and race on migration. Based on the results in Table 3.6.4, it is assumed that the base rates of rural-to-urban migration are multiplied by factors of 0.66 in individuals with no education, 0.57 in individuals whose highest grade passed is 1-5, 0.73 in individuals whose highest grade passed is 6 or 7, 1.00 in individuals whose highest grade passed is 8-11, 1.13 in individuals whose highest grade passed is 12, and 1.72 in individuals with tertiary education. Since education appears to have relatively little effect on urban-to-rural migration, base rates of rural-to-urban migration are adjusted by a factor of 1.30 only if the individual has tertiary education.

Since the data from the 2011 census (Table 3.6.4) and the data in Table 3.6.3 suggest somewhat different conclusions about the relative levels of migration in different race groups, we have compromised by setting the racial adjustments to the rural-to-urban rates as the average of the multiples estimated from the two sources. This means setting the adjustments to the rural-to-urban rates to 1 for black South Africans, 1.8 for coloured South Africans and 3.8 for white South Africans. However, the assumed urban-to-rural adjustment factors are set based only on the data in Table 3.6.4 (i.e. 1 for black South Africans, 0.15 for coloured South Africans and 1.06 for white South Africans), since urban-to-rural migration is a relatively small fraction of total migration.

The base rates in Table 3.6.5 are assumed to apply to unmarried individuals. It is assumed that women are 50% less likely to migrate if they are currently living with their partner than if they are not (no adjustment is made in the case of men). In the case of individuals who are married but not currently living together with their partner (because they are geographically separated), it is assumed that the rate of migration is doubled (i.e. they are more likely to migrate so that they can be with their partner). Although we lack data to support these assumptions, Kok and Aliber have found that South Africans who were married expressed lower intention to migrate than those who were unmarried, even after controlling for age and other factors [96].

It is also assumed that of births to black women in urban areas, 10% are relocated to rural areas soon after birth. It was necessary to make this assumption in order to match the low fractions of children urbanized in the censuses, relative to the fraction urbanized in women of reproductive age (see Figure 3.6.4).

#### 3.6.3 Calibration to urbanization data

Figure 3.6.3 shows the model estimates of the change in the fraction of the population that is urbanized, over time. The model estimates are reasonably consistent with the census data, confirming the steady upward trend in the fraction of the population that is urbanized. However, there is less consistency when comparing the model estimates with the estimates of the General Household Surveys, which do not appear internally consistent (for example, the jump in the fraction urbanized between 2004 and 2005 does not appear plausible).

**Figure 3.6.3:**
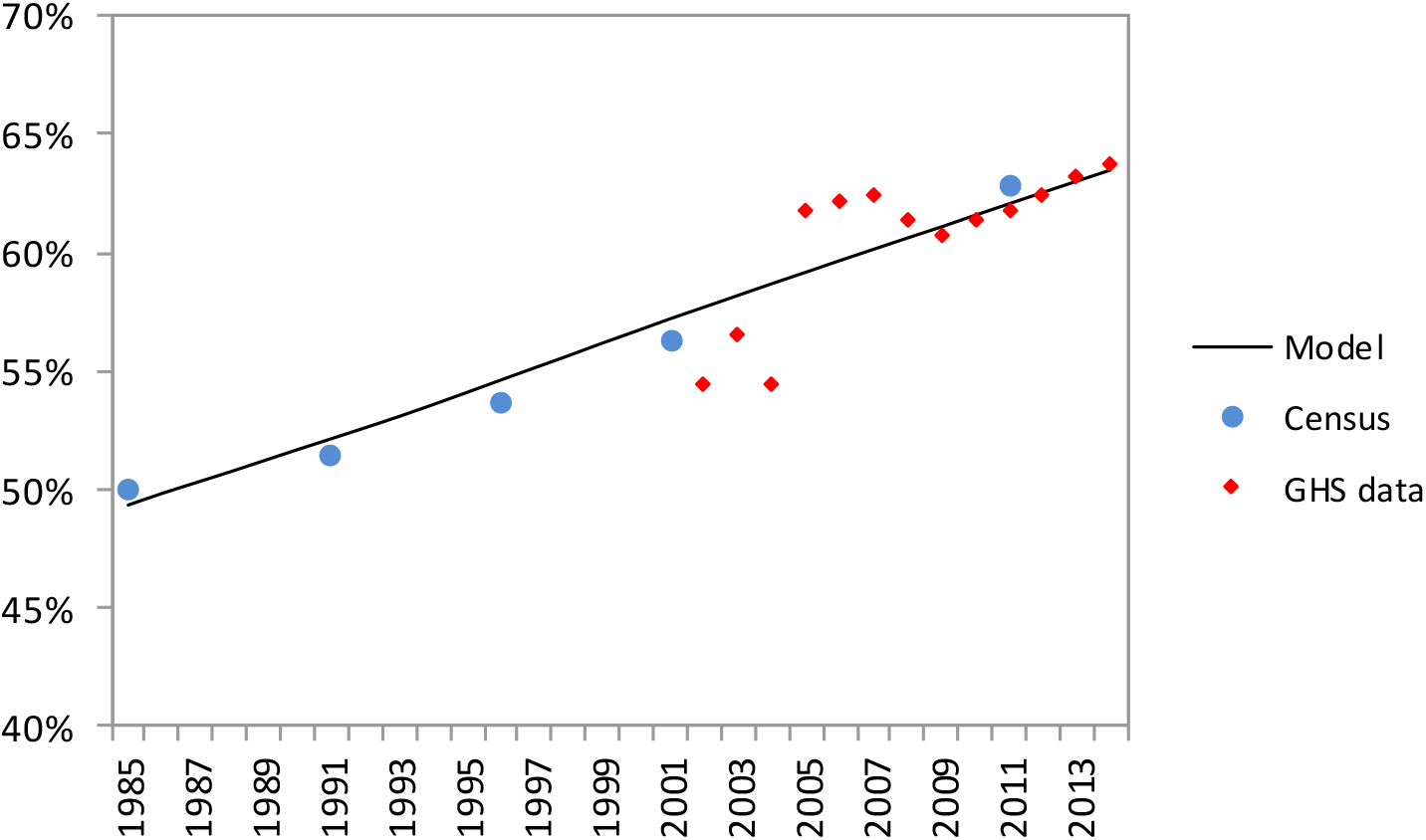
Trend in the fraction of the population urbanized. Census estimates for 1985 and 1991 are from Kok and Collinson [24], who report having adjusted their estimates to take into account the exclusion of the homeland states in these two censuses. Other census estimates and estimates from the General Household Surveys (GHSs) have been calculated directly from the data files on the Statistics South Africa website. Model estimates are the average from 100 simulations.

Figure 3.6.4 shows the patterns of urbanization by age and sex, in 1996 and 2011. The model is roughly consistent with the census in both years, showing a steep rise in the fraction urbanized in the early twenties, and a gradual decline in the fraction urbanized at older ages.

The model is not as consistent with the 1996 census data at the older ages, which could possibly be a reflection of age misreporting in the census.

**Figure 3.6.4:**
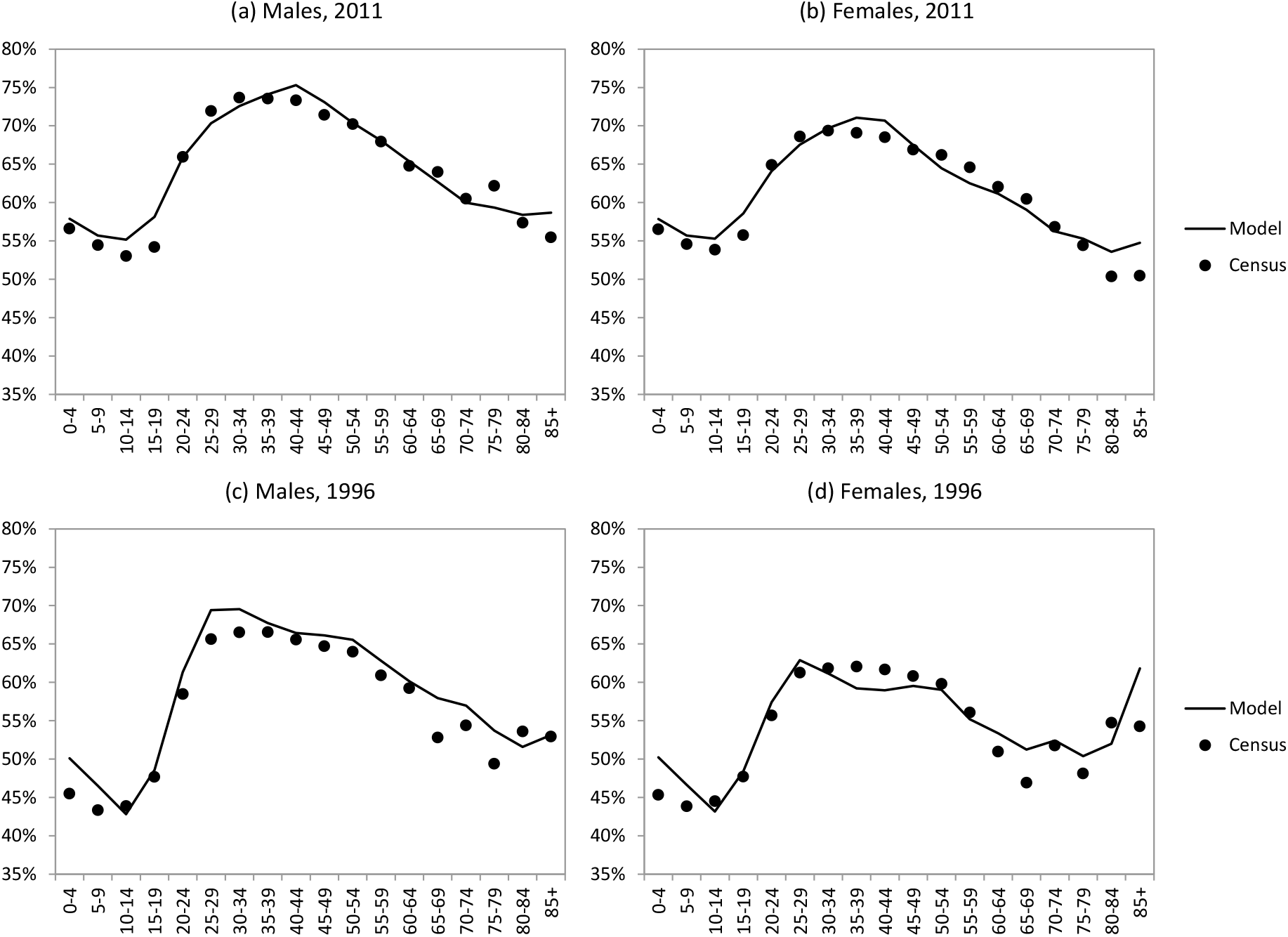
Fraction of the population urbanized, by age and sex. Census estimates have been calculated directly from the data files on the Statistics South Africa website. Model estimates are the average from 100 simulations.

Figure 3.6.5 shows the model estimates of the fraction of the population urbanized, by race. Again, there is a fair degree of consistency between the model and the census, with the fraction urbanized being substantially lower among black South Africans than among coloured and white South Africans. This suggests that the assumed race-specific adjustments to the base rates of rural-to-urban and urban-to-rural migration are reasonable.

**Figure 3.6.5:**
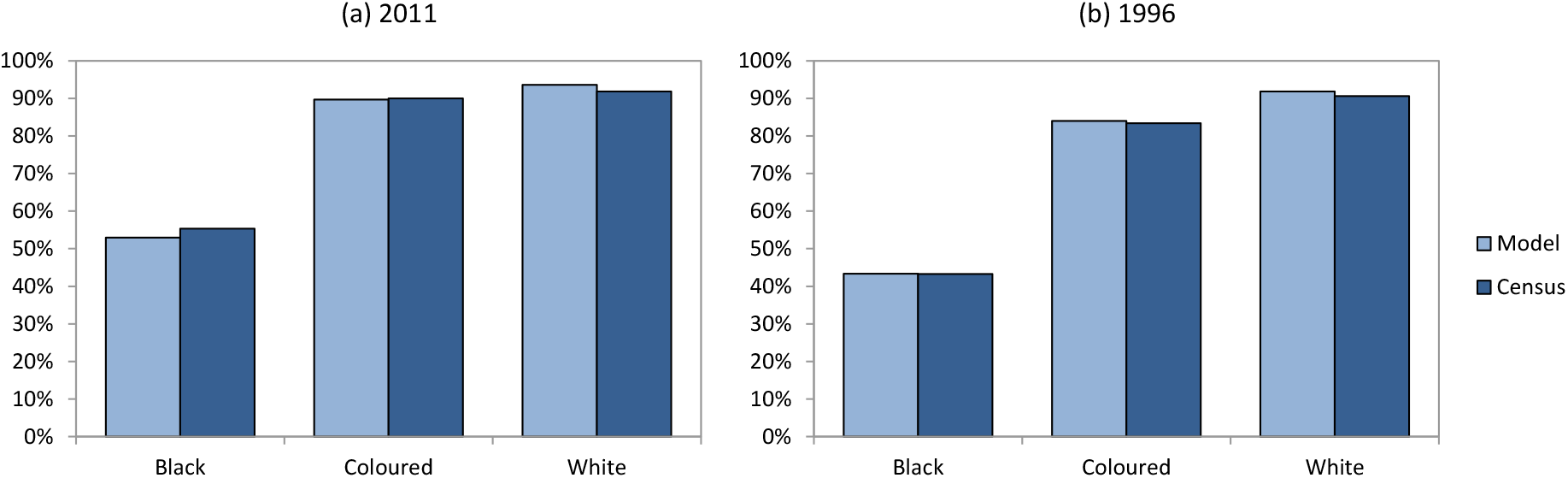
Fraction of the population urbanized, by race. Census estimates have been calculated directly from the data files on the Statistics South Africa website. Model estimates are the average from 100 simulations.

Finally, Figure 3.6.6 compares the model estimates and 2011 census estimates of the fraction of the population urbanized, by highest educational attainment. Although the model is in good agreement with the data for grades 6 and higher, the model over-estimates the fraction urbanized among individuals with little or no education. This could potentially be because the model assumes rates of school enrolment, grade advancement and school dropout are the same in urban and rural areas; if in fact school enrolment and grade progression is lower among rural children than among urban children, the model may be over-estimating the fraction urbanized among children with poor educational attainment. Although data from the National Income Dynamics Study do not suggest an independent effect of urban location on rates of dropout [83], dropout was significantly associated with many other socioeconomic factors, which are likely to be differentially distributed between urban and rural areas.

**Figure 3.6.6:**
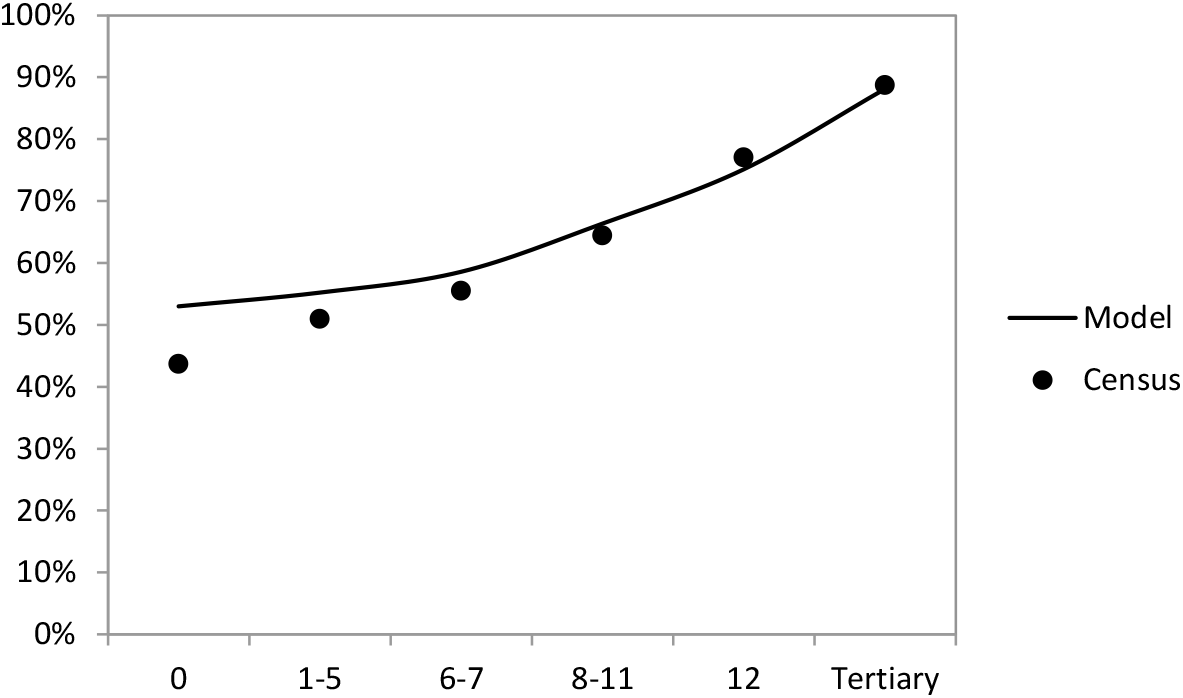
Fraction of the population urbanized in 2011, by highest educational attainment. Census estimates have been calculated directly from the data files on the Statistics South Africa website. Model estimates are the average from 100 simulations.

#### 3.6.4 The effect of migration on sexual behaviour

Several South African studies have documented a positive association between migration and either sexual risk behaviour or HIV infection. Welz *et al* [97] found that in rural KwaZulu-Natal, the risk of HIV was significantly higher in adults who were away from home for 10 or more nights in the last 4 months (aOR 1.3 for women and 1.4 for men). Comparing co-resident couples in the same community with couples in which the male partner worked in an urban centre, Lurie *et al* [98] found that male HIV prevalence was significantly higher in the migrant couples (aOR 2.65, 95% CI: 1.12-6.26). This study also found that the migrant men were significantly more likely to report having casual partners than the non-migrant men (20% vs 6%), but other sexual behaviour parameters were similar in the two groups (or the difference could be explained by age confounding). A more recent study in the same community found that after controlling for age and time, non-resident household members were significantly more likely to report multiple partners in the last year than resident household members (aOR 3.10 (95% CI: 1.88-5.12) in women and 1.66 (95% CI: 1.32-2.09) in men), and similar differences were seen in the reporting of concurrent partnerships, but HIV prevalence did not differ significantly between the two groups [99].

Although the previously-cited studies all relate to the same rural community in KwaZulu-Natal, similar differentials between migrants and non-migrants are seen in other settings. In Carletonville, an urban community close to Johannesburg, Williams *et al* [100] found that the HIV prevalence in men was substantially higher among migrants than in non-migrants (29% versus 19%), and HIV prevalence in women was also substantially higher in migrants than in non-migrants (53% versus 39%). Cluver *et al* [101] found that among adolescents girls in Mpumalanga and Western Cape, the number of moves between homes was significantly associated with the odds of reporting unprotected sex (aOR 1.22, 95% CI: 1.00-1.49), although the same was not true for adolescent boys or for any other measure of sexual risk behaviour. However, in a study of men in rural Limpopo, Collinson *et al* [102] found that men who were employed outside of the study area (temporary migrants) were *less* likely to report more than 1 partner in the last year when compared to men who were employed inside the study area. The study nevertheless found that among the men who were employed outside of the study area, those who returned home more frequently were less likely to report having more than one partner in the last year, which suggests that men who are able to return to their partners in rural areas on a regular basis may feel less need to form additional partnerships in the areas where they work.

The interpretation of the evidence is unclear: is the de-stabilization associated with migration the cause of increased risk behaviour, or is separation from one’s regular partner the principal cause of the increased risk behaviour? It is also possible that there may be a selection effect; if individuals who are ‘risk seekers’ are more likely to migrate, we might expect higher levels of sexual risk behaviour in such individuals. For example, Kok and Aliber [96] found that when South Africans were asked about whether they intended to migrate in the next five years, individuals who were risk averse were less likely to report intention to migrate, even after controlling for age and other confounding factors. It has also been noted that the association between HIV and migration may be due to an effect of HIV on migration, rather than the other way round [103]. However, strong associations between migration and HIV have been observed even in the early stages of the South African epidemic [104], when most HIV-positive individuals were asymptomatic, and hence the association between HIV and migration cannot be attributed only to health seeking or ‘going home to die’ [105].

In the model we assume that separation from one’s primary partner is the primary cause of increased risk behaviour. Individuals who migrate from one location to another, and who are in the high risk group, are assumed to have an altered rate of secondary partner acquisition. Suppose that *c* is the rate at which a high risk individual acquires their primary partner, and *cθ* is the rate at which a high risk individual acquires their secondary partner if their primary partner is in the same location (in general it is assumed that *θ* < 1). In the event that the primary partner is not in the same location, the rate of secondary partner acquisition changes by raising *θ* to the power of *λ*^*a*/365^, where *λ* is a scaling parameter that determines the extent of the behaviour change and *a* is the average number of days per annum that the individual spends away from their primary partner. The adjustment for the number of days away ensures that individuals who are hardly ever away from their primary partner have adjustment factors close to *θ*, while those who are almost always away from their primary partner have adjustment factors close to *θ^λ^*. The *λ* parameter has to be between 0 and 1 (these limits corresponding to the maximum and minimum behaviour changes respectively). In initial attempts to calibrate the model, we attempted to set the *λ* parameter in such a way that the model matched the observed relative differences in HIV prevalence between migrants and non-migrants. However, the model results were found to be relatively insensitive to the *λ* parameter, and the parameter was therefore set arbitrarily to 0.5.

#### 3.6.5 Urban/rural differences in sexual behaviour

Although several South African studies have reported on differences in HIV prevalence between urban and rural areas, there is a lack of South African data on urban-rural differences in sexual behaviour. Cluver *et al* [101] found relatively few differences in sexual behaviour between urban and rural youth: the only differences that were statistically significant were a lower odds of age-disparate sex and a higher odds of multiple partners among girls in urban areas, and a higher probability of unprotected sex reported by boys in urban areas. This is somewhat inconsistent with the pattern observed in other African countries, where age-disparate sex tends to be more common in young women in urban areas than in rural areas, and where condom use tends to be more common in urban areas than in rural areas [106]. However, the finding of a higher reporting of multiple partnerships in young women in urban areas when compared to rural areas is consistent with data from other African countries [106]. Maughan-Brown *et al* [107] found that in a national survey in 2012, men in rural areas were significantly less likely to report concurrency than men in urban areas, but levels of condom use were similar in urban and rural areas, both for men and women.

The 2012 HSRC household survey found that rates of early sexual debut (before age 15) were similar in urban and rural youth (10.2% and 11.0% respectively), and adult reporting of condom use at last sex was also similar (35.9% in urban and 33.4% in rural), but reporting of multiple partnerships was slightly more common in urban areas than in rural areas (12.9% and 10.7% respectively) [25]. The 2005 HSRC household survey found that among 15-24 year olds the proportion who reported past sexual experience was similar in urban and rural areas, but the fraction of adults who reported more than one partner in the last year was slightly higher in urban areas than in rural areas, and the fraction of adults who reported condom use at last sex was also slightly higher in urban areas than in rural areas [108]. Similar results were obtained in the 2002 HSRC household survey [109]. However, none of these analyses controlled for confounding factors such as age and sex, and these comparisons therefore need to be treated with caution.

There is some evidence to suggest that commercial sex activity may be more frequent in urban areas than in rural areas. In a review of the international literature, Vandepitte *et al* [110] found that in African countries, the fraction of women who reported having sex in exchange for money or goods was consistently higher in urban areas than in rural areas. A South African sex worker size estimation study sampled a number of towns and cities across South Africa [111]. Across the 8 cities with populations of 200 000 or more, the average fraction of women engaged in sex work was estimated to be 0.60%. Although relatively few smaller centres were sampled, the sex worker fraction was estimated to be 0.27% in Thohoyandou (other smaller centres were sampled, but because they were all either border towns or towns on major trucking routes, they were not considered representative of rural areas). A study in Zambia found that among married men, urban residence was associated with a significantly higher probability of visiting a sex worker in the last year (aOR 3.14, 95% CI: 1.68-5.85), although there was no significant difference between urban and rural areas when comparing unmarried men [112].

In view of the lack of strong evidence of differences in sexual behaviour between urban and rural areas, and in view of the similarity of HIV prevalence levels in urban and rural areas in recent years (discussed below), we assume sexual behaviour parameters are the same in urban and rural locations. However, men in rural areas are assumed to visit sex workers at a rate that is 0.5 times the national average, while men in urban areas are assumed to visit sex workers at a rate that is 1.5 times the national average (i.e. a 3-times higher rate of male contact with sex workers in urban areas than in rural areas).

#### 3.6.6 Urban/rural differences in initial HIV prevalence

Many studies in developing countries have found higher levels of HIV prevalence in urban areas compared to rural areas [113]. In South Africa, early surveys found HIV prevalence levels in urban areas 4-5 times those in rural areas; however, in more recent surveys differences in HIV prevalence between urban and rural areas have become less significant and the ratio of urban to rural prevalence is close to 1 (Table 3.6.6). This suggests that the epidemic is likely to have started in urban areas and gradually spread to rural areas. To capture this dynamic, we assume in our model that all initial HIV infections in 1990 occur in urban centres.

**Table 3.6.6:** Ratio of HIV prevalence in urban areas to HIV prevalence in rural areas

| Reference | Population | Year | Ratio |
| --- | --- | --- | --- |
| Cronje *et al* [114] | Women in households in Free State | - | 3.8 |
| Department of National Health and Population Development [115] | SA antenatal clinics | 1991 | 6.5 |
|  |  | 1992 | 4.2 |
|  |  | 1993 | 2.7 |
|  |  | 1994 | 4.5 |
| Coleman & Wilkinson [116] | Antenatal clinics in KwaZulu-Natal | 1995 | 1.4 |
| HSRC [109] | SA households | 2002 | 1.6 |
| Reproductive Health Research Unit [117] | SA youth (aged 15-24) in households | 2003 | 1.2 |
| Shisana *et al* [108] | SA households | 2005 | 1.0 |
| Shisana *et al* [25] | SA households, ages 15-49 | 2012 | 0.9 |

#### 3.6.7 Modelling of trips away from home

Individuals who are geographically separated from their primary or secondary partner are assumed to make trips away from their main place of residence on a regular basis, in order to visit these partners. Based on data collected from male migrants who had partners in the Hlabisa district, Coffee *et al* [14] estimated that the number of return trips per annum varied between 4 per year for men who were working in Carletonville (an urban area in Gauteng) and 12 per year for men who were working in Richards Bay (an urban location close to Hlabisa). The average duration of these return trips was estimated at 5 days and 4 days respectively. Based on these data, we assume that an individual’s frequency of visit to a partner in a different location is sampled from a gamma distribution with a mean of 8 visits per year and a standard deviation of 6. If the individual’s frequency of visit is *f* per annum, their assumed average length of stay is 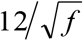, so that individuals who visit their partner more frequently will tend to stay for shorter average durations. For the average value of *f* = 8, this is equivalent to an average duration of 4.2 days, close to the values estimated by Coffee *et al* [14]. The frequency of visits is assumed not to depend on age or sex, as data from the 2001 census suggest that temporary cohabitation among married couples (an indicator of the frequency of visits among geographically separated couples) is similar by age and sex.

It is likely that the frequency of sex during these return visits is higher than that in relationships in which partners are normally living in the same location, and some mathematical models make provision for this [13]. Suppose that *K* is the total annual number of sex acts that would be expected if the couple were not geographically separated. Then the coital frequency (per day) that is expected during the periods when the couple is together is

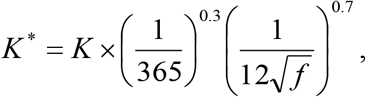

which is a weighted geometric average of the minimum coital frequency (*K*/365, the coital frequency that would be expected if couples had the same coital frequency during periods together as couples who are always together) and the maximum coital frequency 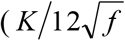, the coital frequency that would be expected if geographically-separated couples had the same total amount of sex in a year as couples who are not geographically separated). The weights of 0.3 and 0.7 were chosen so that a geographically-separated couple with the average frequency of visits (8 per year) has roughly half as much sex in a year as a couple who are not geographically separated.

In the model, an individual’s ‘partner visit’ status is updated 48 times per annum (i.e. roughly at weekly time steps). If an individual is randomly assigned to visit their partner in the current time step, and their average length of stay is less than 365/48 days, it is assumed that they will automatically return to their normal place of residence in the next time step. During this time step, their expected number of sex acts with their geographically-separated partner will be 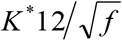 and their expected number of sex acts with their partner in their normal place of residence (if they have one) will be

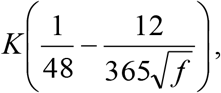

assuming the same coital frequency (*K*) applies in their other relationship. If an individual is assigned to visit their partner in the current time step, and their average length of stay is greater than 365/48 days, it is assumed that they will return to their normal place of residence in the next time step with probability

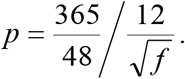

(The same formula applies in the following time step; this formula ensures that the expected total duration of a single visit remains equal to 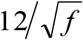.) During this time step, their expected number of sex acts with their geographically-separated partner will be *K*\* 365/48, and they will have no sex with any regular partners in their usual place of residence.

In the above equations, it is implicitly assumed that only one partner initiates the visits, but in reality, either partner could initiate the visit. To account for this, suppose *f*_1_ and *f*_2_ are the desired visit frequencies for the male and female partner respectively, in a heterosexual couple who are geographically separated. In a given time step, the probability that the male partner initiates a visit is 0.5 *f*_1_/48 and the probability that the female partner initiates a visit is 0.5 *f*_1_/48, so that if both partners have the same desired visit frequency, the probability of either partner visiting the other is approximately *f*_1_/48. The average visit duration and expected number of sex acts during the visit are determined based on the desired visit frequency of the partner who initiated the visit.

#### 3.6.8 Calibration to HIV data

Figure 3.6.7 compares the model estimates of the ratio of HIV prevalence in urban areas to that in rural areas and the survey estimates of the urban-to-rural prevalence ratio summarized in Table 3.6.6. The model and the surveys both show a sharp decline in the urban-to-rural HIV prevalence ratio in the early stages of the epidemic, though there is wide variation in the ratios early on in the epidemic (both simulated and observed).

**Figure 3.6.7:**
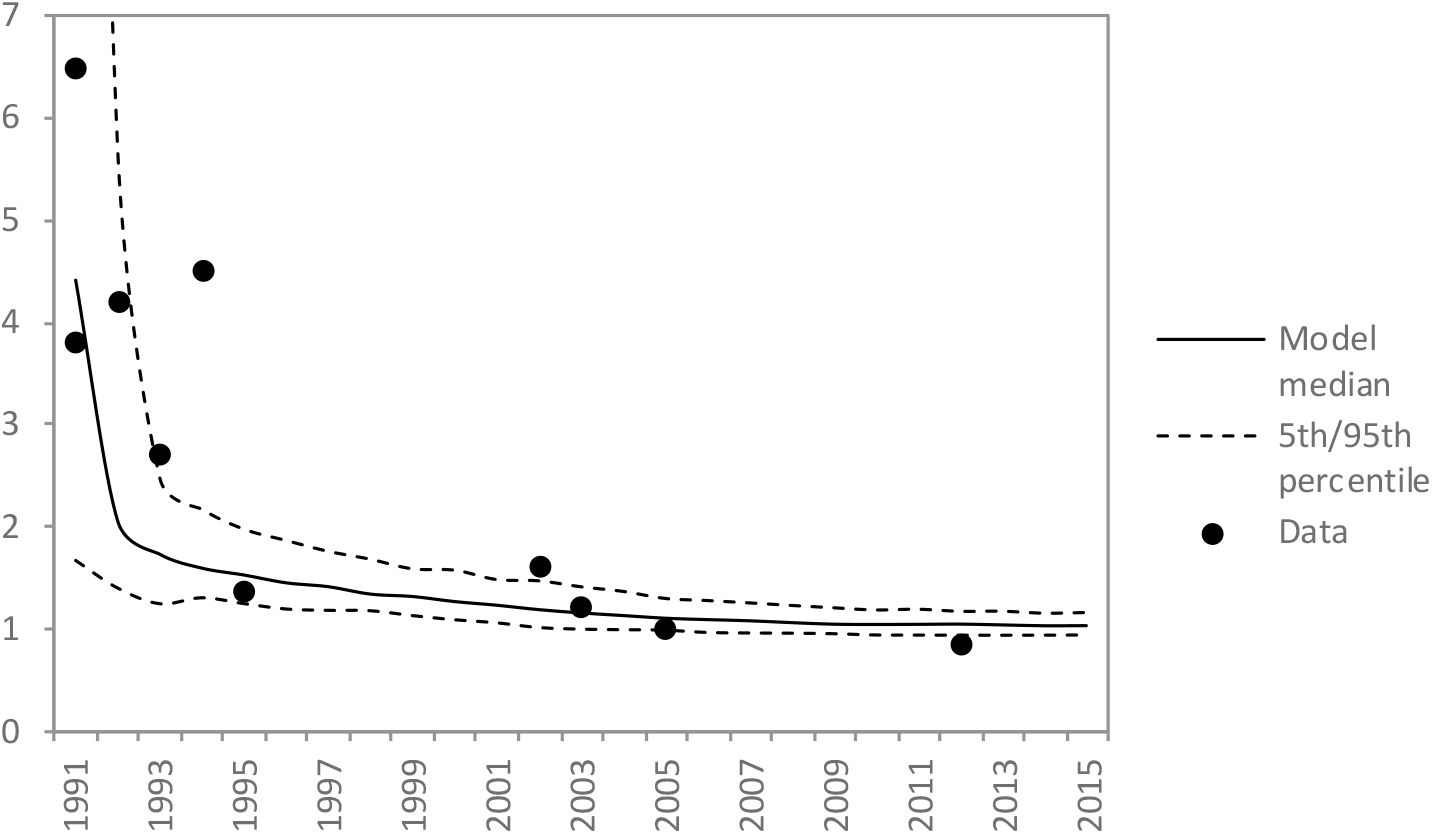
Ratio of HIV prevalence in urban areas to rural areas

Figure 3.6.8 compares model estimates of the association between migration and HIV with empirical estimates. The model estimates have been calculated by running a logistic regression on the individual-level model outputs for each year (migration is defined as having moved from an urban to a rural area or vice versa in the last 2 years, and the analysis is limited to individuals who are either living in rural areas or recent migrants, to be consistent with the approach adopted in most of the studies to which the model is compared). The data to which the model results are compared are mostly from rural South Africa [97–99, 118]. Consistent with the data, the model estimates of the migration-HIV association are consistently positive, and decline over time. This is likely to be because the migration-HIV association is mediated by the higher HIV prevalence in urban areas when compared to rural areas (i.e. migrants are more likely to have had sexual contact with individuals in urban areas and are thus at a higher HIV risk). Because the urban-rural HIV prevalence differentials diminish over time, the association between migration and HIV also becomes more attenuated in the advanced stages of the HIV epidemic.

**Figure 3.6.8:**
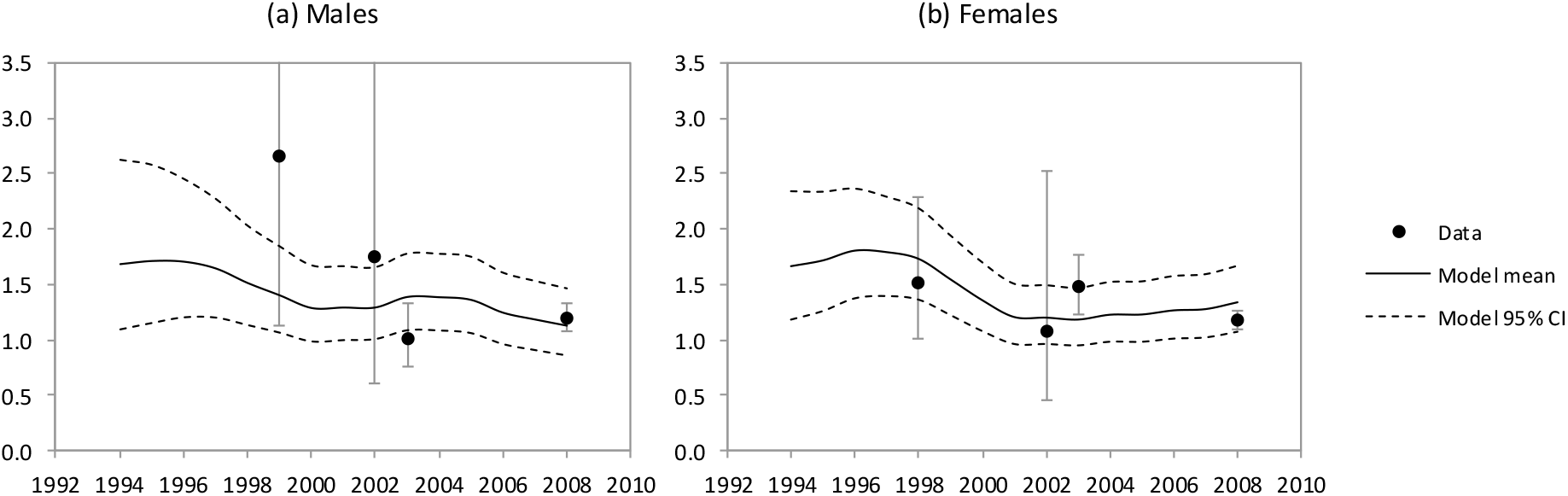
Odds ratios for the association between HIV and recent migration. Model results are calculated by running a multivariate logistic regression on the model outputs (controlling for age, age squared, race, marital status and recent migration), separately for each year (the adjusted odds ratios have been smoothed using a moving average). The data summarized are also all from studies that used multivariate logistic regression to control for major confounders between HIV and recent migration [97–99, 118].

### 3.7 Incarceration

South Africa has among the highest rates of incarceration in the world [119, 120]. We model incarceration using a relatively simple model in which prisoners are classified as either unsentenced (awaiting trial) or sentenced. As 98% of prisoners are male [121], the model simulates only male prisoners.

#### 3.7.1 Rates of incarceration and length of incarceration

Rates at which men are incarcerated are assumed to depend on a number of factors: age, educational attainment, race, risk group and prior incarceration. Table 3.7.1 shows the annual rates at which men in the ‘baseline’ category (incomplete schooling, African, with no prior incarceration) are assumed to enter the prison system. These have been roughly estimated from the age distribution of unsentenced prisoners in 2004 [122] and estimates of the number of men in the population in the corresponding age groups in 2004 [123]. These base rates are adjusted by a factor of 0.5 in men who have completed high school (i.e. a 50% lower rate of imprisonment), 1.50 in coloured men, 0.25 in white men and 10.0 in men who have previously been incarcerated. These adjustment factors were chosen such that the modelled prison population matched the demographics of prisoners in various studies (see Calibration section below).

**Table 3.7.1:** Annual rates at which men are incarcerated

| 15-19 | 20-24 | 25-29 | 30-34 | 35-39 | 40-44 | 45-49 | 50-54 | 55-59 | 60-64 | 65+ |
| --- | --- | --- | --- | --- | --- | --- | --- | --- | --- | --- |
| 0.0080 | 0.0135 | 0.0075 | 0.0047 | 0.0030 | 0.0020 | 0.0015 | 0.0011 | 0.0008 | 0.0005 | 0.0000 |

Statistics from the Department of Correctional Services indicate that 48% of unsentenced prisoners have been in prison for more than 3 months [124]. Based on this, we assume that the average time spent in the ‘unsentenced’ stage is 0.34 years (−0.25/ln(0.48)). Upon leaving the ‘unsentenced’ state, a proportion ***θ*** of prisoners are assumed to be released (i.e. the cases against them are dismissed or they are proven innocent) and the remainder (1 – *θ*) are sentenced. The ***θ*** parameter has been estimated based on the observation that the fraction of prisoners who are sentenced is around 71% [121, 122, 125]. If the average sentence length is 1.17 years (see below), then the fraction of prisoners who are sentenced is

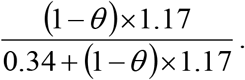

Setting this equal to 0.71 and solving for ***θ*** gives an estimate of 0.29, and the assumed fraction of prisoners who are released without being sentenced is therefore set to 0.29.

Among those who are sentenced, the time in prison after sentencing is randomly sampled from a mixture of two exponential distributions: one representing the standard distribution of sentence lengths (at the time of sentencing), and the other representing the distribution of prison times (after sentencing) in prisoners who are granted parole. The former distribution is assigned a mean of 1.7 years, this parameter being chosen in order to match the distribution of sentence lengths among newly-sentenced prisoners in 1997 [126]. It is worth noting that the distribution of sentence lengths among *newly-sentenced* prisoners is not the same as the distribution of sentence lengths among *current* prisoners (which is more commonly reported). The second distribution is assigned a mean that is half of the corresponding mean for the first distribution, since South African law has specified (since 2004) that prisoners should be considered for parole when they have served half of their sentence. Parole is not granted automatically, and it is therefore necessary to specify a parameter *ψ*, which represents the fraction of newly-sentenced prisoners who are ultimately granted parole. Limited data are available on the number of parolees as a fraction of the total number of individuals serving sentences, but data from the 2013-2015 period indicate that parolees comprise 31% of the combined parolee and sentenced prisoner populations [121]. From this it follows that 0.31 = 0.5*ψ* (assuming that the granting of parole is independent of the sentence length and that parole is granted halfway through the sentence length), i.e. *ψ* = 0.62. This value therefore determines the mixture weight assigned to the second exponential distribution, while a weight of 0.38 is assigned to the first exponential distribution. From this it follows that the overall mixture mean (i.e. the average length of time spent in prison after sentencing) is 0.38 × 1.7 + 0.62 × 1.7 × 0.5 = 1.17 years.

A limitation of the model is that it does not allow for the effect of age or previous conviction on sentence length. South African law requires that judges consider individual circumstances (e.g. previous convictions) in determining the sentence length, and offenders are typically given shorter sentences if it is their first offence. Another limitation is that the model parameters have been derived on the assumption that rates of incarceration, sentencing practices and parole practices have remained stable over time, though in fact there have been substantial changes in practices since the late 1990s [127].

#### 3.7.2 Association between incarceration and sexual behaviour

Little is known about sexual behaviour in South African prisons. Although there is qualitative evidence to suggest that rape and consensual sex between prisoners are both common [128, 129], data quantifying the extent of sex in prisons are limited. In a survey of prisoners in the KwaZulu-Natal and Mpumalanga provinces, Sifunda *et al* [130] found that only 6% reported having engaged in anal intercourse (either voluntarily or involuntarily) while in prison. In a survey of MSM in Cape Town and Johannesburg, it was found that of those who had ever been in prison, only 12.0% reported consensual unprotected anal intercourse (UAI) and 7.6% reported forced UAI [131]. In a review of the international literature, Dolan *et al* [132] found that very few studies quantified the extent of HIV incidence in prisons, and the few that had found very low rates of HIV transmission between prisoners. In South Africa, HIV-positive prisoners were segregated from HIV-negative prisoners until 1996 [133], making HIV transmission almost impossible. A more recent survey found that HIV prevalence in recently-sentenced prisoners was similar to that in prisoners who had been incarcerated for longer durations, despite the latter being older on average [134] – this is the opposite of what would be expected if significant levels of HIV transmission were occurring in prison. Lane *et al* [135] also found that among MSM in Soweto, there was no difference in HIV prevalence between those who reported prior incarceration and those who did not. Similarly, Jewkes *et al* [136] found that among young men in the Eastern Cape, prior incarceration was not significantly associated with HIV prevalence in multivariate analysis. Given that relatively few men report engaging in UAI while in prison, and given the lack of clear evidence of significant HIV incidence in prisons, we have not modelled transmission of HIV between prisoners.

It is possible that men in the general population who engage in high risk behaviours may be more likely to engage in criminal activity that would lead to incarceration. Stephens *et al* [137] found that among prisoners in KwaZulu-Natal and Mpumalanga, 55% reported having ever had an STI – this suggests a relatively high level of sexual risk behaviour. However, a case-control study, which included former prisoners (cases) and individuals who had never been incarcerated (controls) in four countries that included South Africa, found that prior incarceration was not significantly associated with HIV risk behaviour in multivariate analysis [138]. In initial analyses, we found that it was not possible to match our model estimates of HIV prevalence in prisoners to those observed in surveys unless we assumed a greater rate of incarceration in sexually active high-risk men than in other men. Rates of incarceration in high-risk sexually experienced men are therefore assumed to be double those in other men. (The rates of incarceration specified in section 3.7.1 are multiplied by factors of 1.5 and 0.75 in high-risk and low-risk men respectively.)

It is assumed that while men are in prison, they do not have any sex with heterosexual partners and do not form any new heterosexual relationships. Consistent with international literature [15, 139], we assume that women whose partners are currently in prison are more likely to acquire additional partners while their partner is in prison. The procedure is the same as that used to model the effect of partners being geographically separated, i.e. assuming that women in the high risk group are more likely to acquire additional partners (see section 3.6.4).

#### 3.7.3 HIV prevalence in prisoners

Very few studies have evaluated HIV prevalence in South African prisoners. The four studies that have been identified are summarized in Table 3.7.2. HIV prevalence estimates vary between 7.3% and 22.3%. However, the reliability of the highest prevalence estimate is questionable, as it is based on routine HIV testing data from prisons (rather than a survey) and only a third of prisoners were tested [124]. The two smaller studies in Johannesburg and Cape Town are also not nationally representative.

**Table 3.7.2:** Estimates of HIV prevalence in South African prisons

| Study | Year | Location | n | HIV prevalence |
| --- | --- | --- | --- | --- |
| Lim'uvune Consulting [140] | 2006 | National | 5288 | 19.8% |
| Department of Correctional Services [124] | 2009-10 | National | 47011 | 23.2% |
| Telisinghe *et al* [134] | 2009-10 | Johannesburg | 957 | 17.2%* |
| Skiti *et al* [141] | 2013 | Cape Town | 1176 | 7.3% |

\* Adjusted to reflect the reported sensitivity and specificity of the assay used.

#### 3.7.4 Calibration to incarceration data

Table 3.7.3 summarizes the model estimates of the demographic and HIV profile of the prison population, and compares them with the data sources used in the model calibration. The model is consistent with the calibration data sources, suggesting a relatively young age profile (43% of prisoners aged 25 or younger) and a relatively high proportion of prisoners who are repeat offenders (44%). The model estimates that the fraction of the adult male prison population has remained fairly stable over time, at around 0.83%. Although this is consistent with data for recent periods, it is worth noting that there have been substantial changes in the size of the prison population over time [127], and thus the model might not adequately reflect the demographics of the prison population in earlier periods.

**Table 3.7.3:** Incarceration calibration

| Indicator | Model estimate* | Calibration target: mean, range | Calibration sources |
| --- | --- | --- | --- |
| % of men (aged 15+) who are in prison, 2012-14 | 0.83% | 0.83% (0.82-0.84%) | [121] |
| % of prisoners aged ≤ 25, 2000-10 | 43% | 41% (35-46%) | [122, 124, 142] |
| % of prisoners who have completed grade 12, 2000-2010 | 18% | 21% (11-29%) | [137, 142] |
| HIV prevalence in prisoners, 2005-2015 | 15.5% | 17% (7-23%) | See Table 3.7.2 |
| % of prisoners previously incarcerated, 2000-10 | 44% | 41% (39-42%) | [128, 134, 142] |
| % of prisoners who are unsentenced, 2000-10 | 30% | 30% (29-32%) | [121, 122, 125] |
| % of prisoners by race group in 2007 |  |  |  |
| Black | 80.0% | 80.4% | [143] |
| Coloured | 18.1% | 17.5% | [143] |
| White or Asian | 1.9% | 2.2% | [143] |

\* The model estimate presented is the average for the period stated, averaged across 100 simulations.

## 4. Sexual behaviour

The model of sexual behaviour is an adaptation of that described in a previous agent-based modelling study [18]. Figure 4.1 summarizes the structure of the sexual behaviour model for heterosexual adults. Individuals are divided into two risk groups: high risk (having a propensity for concurrent partnerships and commercial sex) and low risk (serially monogamous and never engaging in commercial sex). Two types of regular relationships are considered: short-term (non-cohabiting) and long-term (marital and/or cohabiting). In addition, men in the high risk group are assumed to have once-off contacts with sex workers. Individuals in the high risk group can have up to two regular partners at any time, while individuals in the low risk group are (by definition) limited to one partner at a time. As polygamy is rare in South Africa [144], it is assumed that no individual has more than one long-term partner at a time. All long-term relationships are assumed to start as short-term partnerships. Women engaging in sex work are assumed not to form short-term or long-term relationships during the periods in which they are active as sex workers.

**Figure 4.1:**
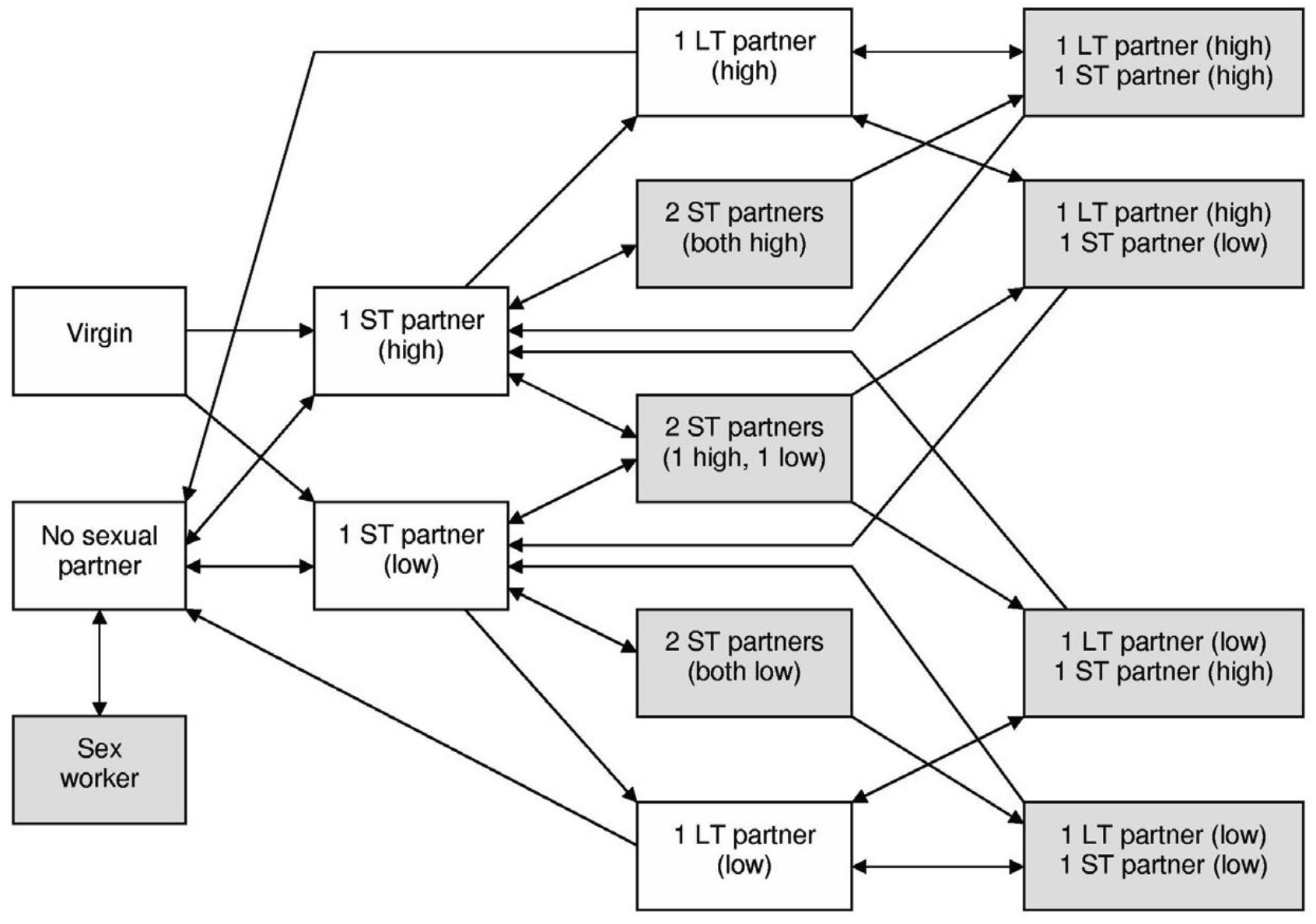
Model of sexual behaviour of ‘high risk’ females. LT = long-term (spousal). ST = short-term (non-spousal). ‘High’ and ‘low’ refer to the risk group of the sexual partner. The model for low risk females is the same as that shown here, except that the shaded states are omitted. The model for high risk heterosexual men is also the same as that shown here, except that the ‘sex worker’ state is omitted.

In modelling the sexual behaviour of MSM, we adopt a similar model structure, but with a few modifications. Firstly, commercial sex is not modelled. Although male sex work does occur in South Africa, male sex workers comprise only about 5% of all sex workers [145, 146], and given the relatively small size of the MSM and sex worker sub-populations, it is not practical to model such a small sub-group within these sub-populations. Secondly, we model casual sex relationships (defined as lasting less than a week) between MSM by assuming that MSM intermittently go through phases of engaging in casual sex. Low risk men are allowed to engage in casual sex even when in a regular relationship, as evidence from high-income settings suggests that a high proportion of MSM engage in casual sex even when in regular relationships [147–149]. This means that the definition of low risk in heterosexuals (‘serially monogamous’) is not strictly applicable to MSM; low risk MSM should rather be defined as men who never have more than one *regular* partner.

In heterosexual adults, the high risk proportion has been set to 35% and 25% in men and women respectively [150–152]. Although the definition of ‘high risk’ is not the same in MSM and heterosexual men, the proportion of men in the high risk group is assumed to be the same for MSM and heterosexual men. Few South African MSM studies report on the prevalence of concurrency. One study in Soweto found that only 39% of sexual relationships between MSM did not involve overlap with other partners [153]. However, this is likely to be an under-estimate of the fraction of MSM who are low risk, because (a) low risk individuals will tend to have fewer sexual partners than high risk individuals and (b) our definition of low risk in MSM does not exclude concurrent casual partnerships.

It is worth noting that this model does not consider same-sex relationships between women. Although same-sex attraction is more prevalent in women than in men [154–156], the primary focus of this modelling analysis is on HIV transmission, which is negligible in the context of sexual relationships between women. It has therefore been assumed for the sake of simplicity that women only enter into relationships with men.

### 4.1 Sexual preference

Although sexual preference is often considered a fixed attribute, studies in high-income countries suggest that sexual preference and same-sex activity can change substantially over the life course [155–157]. Rather than assign men a fixed sexual preference, we assign to each man a ‘male preference’ variable, *Y_i_*, which can vary over the course of his life. For each individual, the pattern of male preference over the life course is defined by two parameters: an initial male preference (*a_i_* for individual *i*), which is assumed to apply up to age 20, and an annual rate of change in male preference (*b_i_* for individual *i*), which applies after age 20. The model considers three categories of men:

1. Men who are exclusively heterosexual throughout their life course (*a_i_* = 0 and *b_i_* = 0)
2. Men who are exclusively homosexual throughout their life course (*a_i_* = 1 and *b_i_* = 0)
3. Men who can engage in both heterosexual and homosexual relationships, either throughout their life course or at different stages of their life (0 < *a_i_* < 1)

For the sake of convenience, we will refer to these three groups of men as ‘heterosexual’, ‘gay’ and ‘bisexual’ in all subsequent sections. For heterosexual and gay men, *Y_i_* will always be 0 and 1 respectively, while for a bisexual man aged *x*, the *Y_i_* value will be

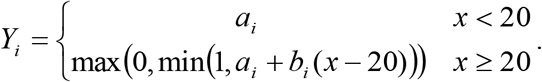

The fraction of men who are in either the gay or bisexual category is assumed to be 5%, based on two surveys. The first of these studies, by Dunkle *et al* [158], used computer-assisted self-interviews, which have generally been found to be associated with lower social desirability bias, and found that 5.4% of men reported having ever engaged in consensual sex with another man. The second survey, conducted in four provinces, found that 5.5% of sexually experienced young men reported having ever had sex with another man [159].

Of the men who are gay or bisexual, 77% are assumed to be in the bisexual category. This estimate is the average of the 100 best-fitting parameters obtained when the model was previously fitted to South African sexual behaviour data from MSM [160]. The data are from respondent-driven sampling (RDS) studies, in which the proportion of MSM who reported having ever had sex with women varied between 36% and 87% (average value of 58%) [131, 135, 161–163]. These studies are likely to under-estimate the proportion of gay/bisexual men who are bisexual because the RDS samples exclude men who have not recently been sexually active with other men. In a more representative household survey, it was found that of men who reported having ever had sex with men, the fraction who reported having ever had sex with a woman was 99% [158], although the sample size was relatively small.

Among bisexual men, there is likely to be heterogeneity in both the extent of their same-sex preference and the degree to which their sexual preference changes over their life course. To reflect this heterogeneity, we sample the *a_i_* values from a beta distribution with mean 0.23 and standard deviation 0.09 and sample the *b_i_* values from a normal distribution with mean - 0.015 and standard deviation 0.16. As before, each parameter estimate is calculated as the average of the 100 best-fitting parameters obtained when the model was previously fitted to South African sexual behaviour data from MSM [160]. The model thus suggests that men in the bisexual category tend to have more female partners than male partners, and that there is a slight tendency towards relatively more female partners as men get older (although with substantial variation between individuals).

### 4.2 Sexual debut

Sexual debut is assumed to occur between the ages of 10 and 30. Sexual debut is assumed to occur upon entry into a short-term relationship, i.e. the rate of sexual debut is specified as a desired rate of entry into short-term sexual relationships (described in more detail in section 4.3).

#### 4.2.1 Effect of educational attainment on sexual debut

Although several African studies have observed an association between low educational attainment and early sexual debut [164], it is not clear if this represents an effect of education on sexual debut or an effect of sexual debut on educational attainment, or possibly confounding due to other factors. For example, in a cross-sectional study conducted in rural Limpopo, Hargreaves *et al* [165] found a strong association between sexual debut and school enrolment in boys; in girls there was also an association, although it ceased to be statistically significant after controlling for women’s fertility history (suggesting that the association in women might be due only to women dropping out of school if they fall pregnant).

Considering first the evidence of an effect of education on sexual debut, the most reliable South African data come from a longitudinal study in rural KwaZulu-Natal [166]. In this study, boys who had dropped out of school prior to completing secondary education started sexual activity at a rate 1.44 times (95% CI: 1.17-1.77) that in boys of the same age who were still in school. A larger effect was seen in girls, although the extent of the increase was strongly age-dependent, and girls who had completed secondary school were significantly more likely to become sexually active than girls of the same age who were still in school (an effect not seen in boys). In contrast, a longitudinal study conducted in Cape Town found that the rate of sexual debut increased as the number of years of completed education increased (after controlling for age) – the increase was statistically significant in girls but not in boys [167]. This study did not control for whether youth were *currently* in school at the time of sexual debut, and the observed positive association in girls could possibly be because girls who have higher levels of completed education are more likely to be out of school. In another study based on the same dataset, school enrolment in 2002 was found to reduce the rate of sexual debut between 2002 and 2005, both in males and females, although neither association reached statistical significance [88]. Another study that used the same dataset found that school enrolment in 2002 was associated with a reduced rate of later sexual debut, although this was only statistically significant in boys [168]. A recent study in Mpumalanga province found that girls who had dropped out of school had 1.85 times as many sexual partners as girls of the same age who did not drop out of school, although this study did not specifically assess sexual debut or control for possible confounding due to pregnancy-related dropout [169]. The evidence therefore suggests that being in school reduces the chance of starting sexual activity, but it is less clear whether educational attainment affects sexual debut, after controlling for age and current school enrolment. Some evidence suggests that youth who are enrolled in school but with relatively low grade advancement have lower rates of sexual debut [88], while other evidence suggests the opposite [82], and some studies suggest no significant effect of grade repetition on sexual behaviour [166, 169].

We next consider whether there is evidence of an effect of sexual debut on educational attainment. In girls it is obvious that pregnancy will often lead to school dropout, and the modelling of this dynamic is described elsewhere (see section 3.5.4). However, there has been no research on whether sexual behaviour predicts school dropout and grade repetition independent of pregnancy.

Lastly, it is possible that some of the observed association between early sexual debut and low educational attainment could be due to unmeasured confounders, and might not be due to any effect of sexual debut on educational attainment or vice versa. For example, it is possible that there is greater social desirability bias affecting the reporting of sexual experience among youth in school than there is among youth who are not in school. Mensch *et al* [170] found that among youth in Kenya, the reporting of sexual debut was much more sensitive to the interview format in girls who were enrolled in school than in girls who were not in school, which suggests greater social desirability bias in schools.

Given the lack of clear evidence of a causal relationship between schooling and sexual debut (independently of that mediated by teenage pregnancy, which is already allowed for in the model), we have made the conservative assumption that neither being in school nor current educational attainment has any effect on rates of sexual debut. The age of sexual debut is also assumed to have no effect on schooling outcomes, except insofar as teenage pregnancy can lead to girls dropping out of school.

#### 4.2.2 Effect of race on sexual debut

Several South African studies have noted a strong association between race and age of sexual debut. This cannot be explained by the effect of school enrolment noted previously, as rates of school enrolment tend to be similarly high across race groups [23], and in the one analysis that did simultaneously control for school enrolment and race when assessing predictors of sexual debut, race remained highly significant [168]. Table 4.2.1 summarizes the studies that have examined the association between race and sexual debut. On average, the rate of sexual debut among coloured youth is 0.75 times that in black youth, while the rate among white youth is 0.47 times that in black youth.

**Table 4.2.1:**
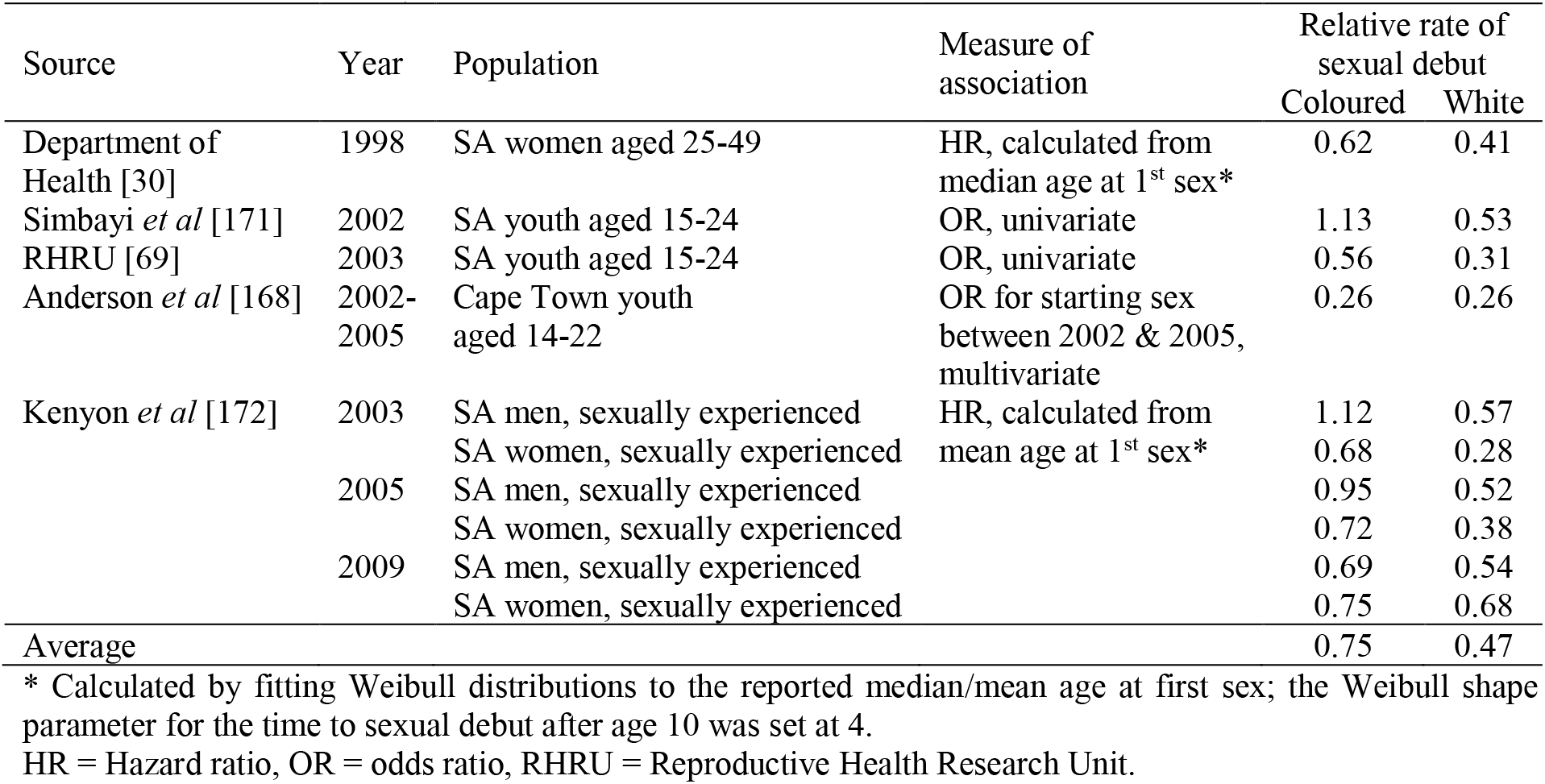
Rates of sexual debut in coloured and white South Africans (as a fraction of rates in black South Africans)

Although the odds ratio (OR) in a cross-sectional study is not the same as the hazard ratio (HR) in a longitudinal study, simulations show that the two are likely to be quantitatively similar (Table 4.2.2). In these simulations, the time to sexual debut after age 10 in black South Africans is assumed to follow a log-logistic distribution, with a median age at sexual debut of 18 years [30]. Regardless of the assumed log-logistic shape parameter, and regardless of the assumed HR relating sexual debut in other race groups to that in black South Africans, the OR for sexual experience in the 15-24 age group is similar to the assumed HR. This indicates that it is reasonable to use the OR estimates in Table 4.2.1 as approximations to the HR.

**Table 4.2.2:**
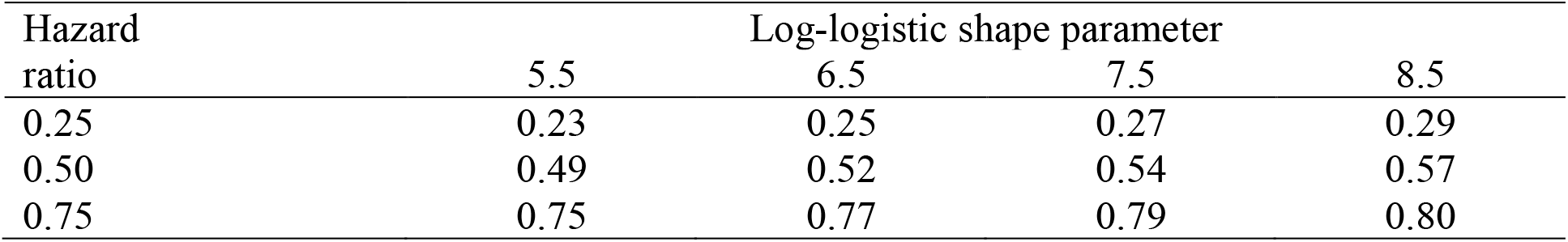
Simulated odds ratios for given hazard ratios, when time to sexual debut is assumed to be log-logistically distributed

| Hazard ratio | Log-logistic shape parameter |  |  |  |
| --- | --- | --- | --- | --- |
|  | 5.5 | 6.5 | 7.5 | 8.5 |
| 0.25 | 0.23 | 0.25 | 0.27 | 0.29 |
| 0.50 | 0.49 | 0.52 | 0.54 | 0.57 |
| 0.75 | 0.75 | 0.77 | 0.79 | 0.80 |

In the model, it is assumed that the relative rates of sexual debut in coloured and white South Africans (expressed as multiples of the rates in black South Africans) are 0.75 and 0.47 respectively, the same as the average values summarized in Table 4.2.1.

#### 4.2.3 Effect of risk group on sexual debut

Several South African studies suggest that individuals who start having sex at early ages are more likely to engage in ‘risky’ sexual practices than those who become sexually active at older ages. For example, Dunkle *et al* [173] found that after controlling for other factors, women who had become sexually active after the age of 20 were both less likely to report having ever engaged in transactional sex (OR 0.26, 95% CI: 0.13-0.51) and less likely to report having ever had more than one concurrent partner (OR 0.67, 95% CI: 0.46-0.99). Negative correlation between age at sexual debut and subsequent risk behaviour is also suggested by Lurie *et al* [174], who found that the risk of HIV infection in a couple was significantly increased if the couple’s average age at sexual debut was 16 or less (OR 2.45, 95% CI: 1.08-5.55), in a multivariate analysis. Harrison *et al* [175] also found that male sexual debut before age 15 significantly increased the odds of reporting more than three partners in the last three years (OR 10.26, p = 0.008), and Mpofu *et al* [176] found that sexual debut before age 16 was significantly associated with multiple partnerships in the year before the survey. Fraser-Hurt *et al* [177] found that in both South African women and men, younger age at sexual debut was strongly associated with higher rates of partnership formation. Similar associations have been observed in other African countries [178, 179].

For the purpose of mathematical models of sexual behaviour, it is necessary to estimate the relative rates of sexual debut in those who have a propensity for ‘high risk behaviour’ (however that may be defined) and in those with no such propensity. Suppose that *h*_1_(*x*) is the rate at which ‘high risk’ individuals begin sexual activity at age *x*, and *h*_2_(*x*) is the corresponding rate at which ‘low risk’ individuals begin sexual activity. Further suppose that the hazard ratio *a* represents the ratio of *h*_2_(*x*) to *h*_1_(*x*). Although *h*_2_(*x*) and *h*_1_(*x*) are usually not measured directly, *a* can nevertheless be estimated by noting that if *p*_1_(*y*) and *p*_2_(*y*) are the probabilities of beginning sexual activity at age *y* or older, in the high risk and low risk groups respectively, then

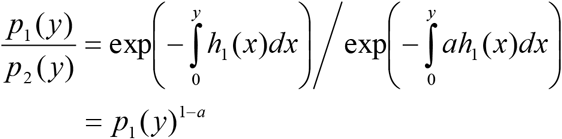

so that *a* can be estimated by the formula

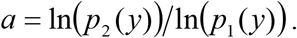

Table 4.2.3 summarizes the estimates of *a* obtained using this equation, from various studies of sexually experienced individuals. Estimates lie between 0.46 and 0.71, with an average of 0.58.

**Table 4.2.3:**
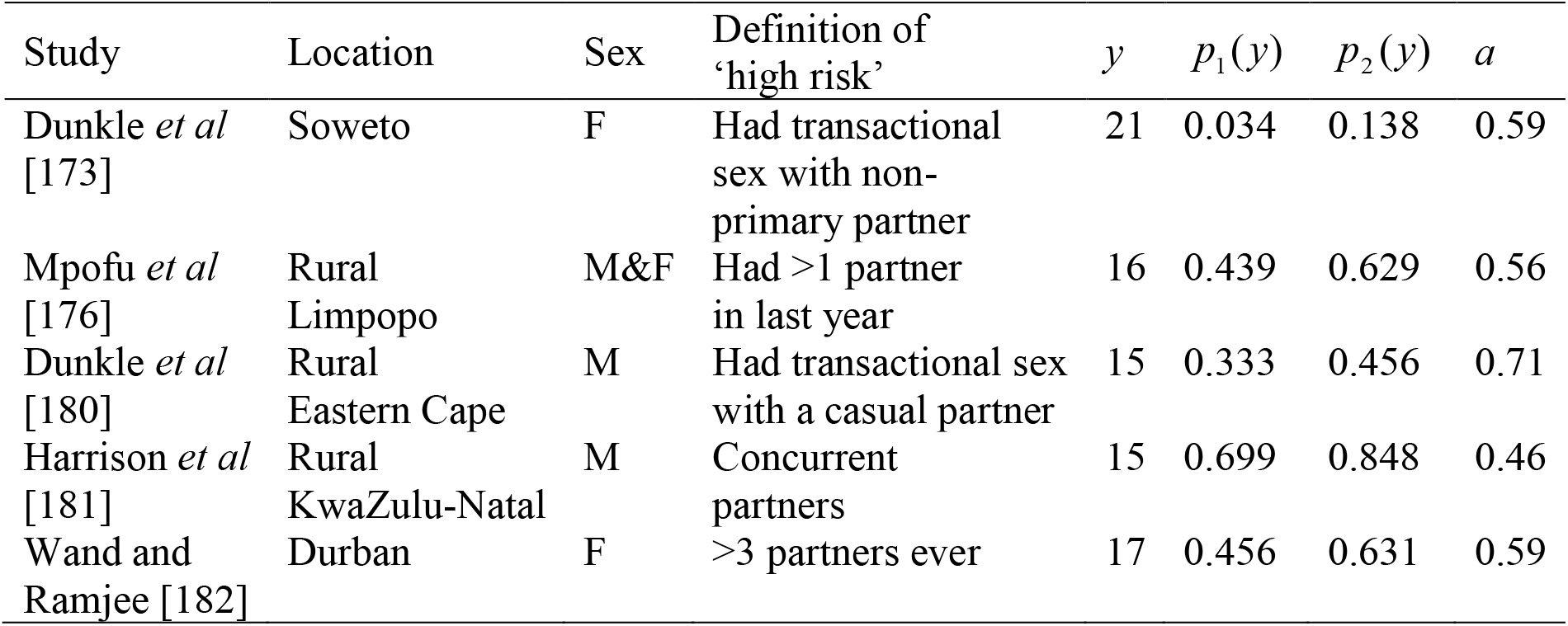
Relative rates of sexual debut in low risk and high risk individuals

In the model, it is assumed that the rate of sexual debut in low risk individuals is 0.58 times that in high risk individuals of the same age, sex and race, after controlling also for current school enrolment.

#### 4.2.4 Assumed rates of sexual debut in baseline category

The hazard function for the rate of sexual debut in the baseline category (black youth in the high risk group) is assumed to be log-logistic in form, with an offset of 10 to prevent sexual debut at ages below 10. The baseline hazard is set separately for males and females. Mathematically, the probability that an individual in the baseline category, aged *x* and of sex *g*, is sexually experienced is

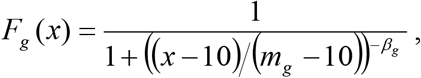

for *x* > 10. In this equation, *m_g_* is the median age at sexual debut, and *β_g_* is the shape parameter that determines the extent to which the rate of sexual debut changes in relation to age. The rate of sexual debut at exact age *x* is then

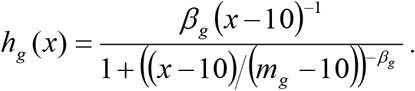

For individuals in the low risk group, this hazard is multiplied by the factor of 0.58 mentioned previously. Similarly, for coloured and white youth the hazard is multiplied by the factors of 0.75 and 0.47 respectively.

The median age at sexual debut in the baseline category (*m_g_*) has been set to 17 for males and 16.5 for females, while the shape parameter (*β_g_*) has been set to 7.5 for males and 8.5 for females. These parameters have been chosen in such a way that the modelled levels of sexual experience at each age are consistent with the results of various national surveys [108, 183, 184], as shown in Figure 4.2.1. For the purpose of this comparison, we have calculated the average age-specific prevalence of sexual experience across the three surveys; however, for women we have adjusted the reported rates by an odds ratio of 2, to reflect likely under-reporting of sexual experience in young women [170, 185]. The model results show the prevalence of sexual experience in 2005, averaged across 10 simulations.

**Figure 4.2.1:**
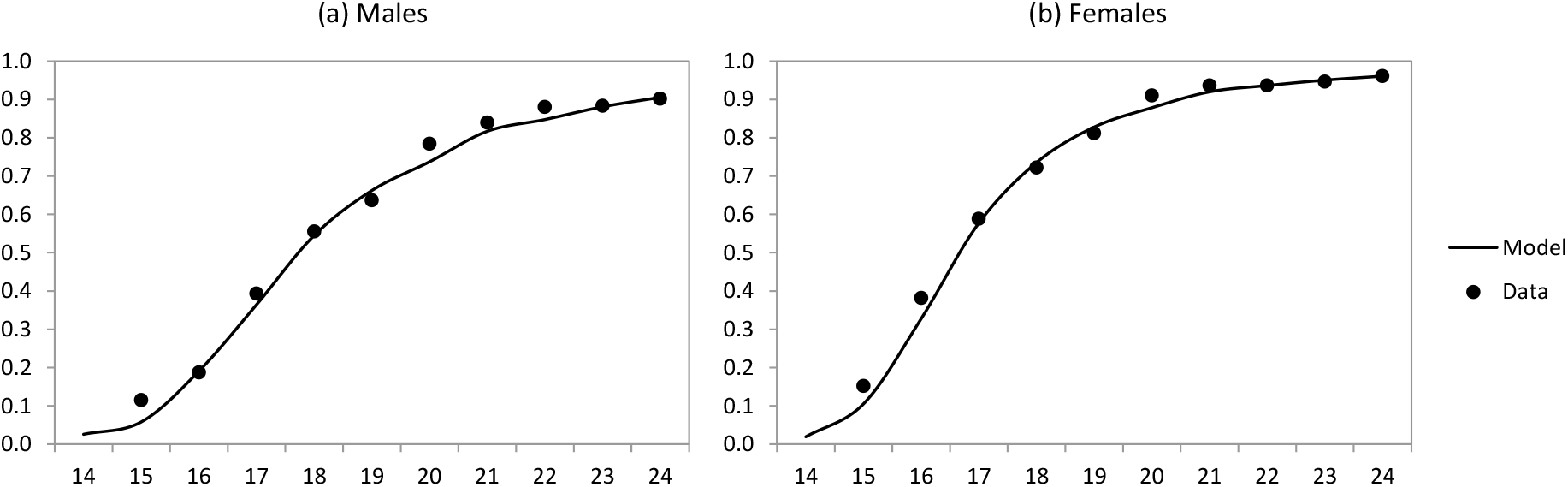
Fraction of youth who are sexually experienced, by age

There is little evidence to suggest a significant difference in the age of sexual debut between heterosexual men and MSM. Two South African surveys have both found that among sexually experienced youth, the average age at first sex did not differ significantly between individuals who self-identified as gay/bisexual and those who self-identified as heterosexual [154, 159]. The rates of sexual debut specified previously for men are therefore assumed to apply to both MSM and heterosexual men.

### 4.3 Rates of short-term partnership formation

Sexually experienced individuals are assumed to acquire new short-term partners at a rate that depends on their age, risk group, race and current relationship status. The parameter 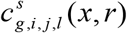 is defined as the annual rate at which a sexually-experienced individual of sex *g* wishes to form new short-term partnerships if they are in risk group *i* (1 for high risk, 2 for low risk), aged *x*, in race group *r*, in HIV disease state *s*, with *j* current partners and marital status *l* (0 for unmarried, 1 for married). A gamma probability density function is used to represent age differences in rates of partnership formation; for the purpose of calculating a constant rate over a five-year age interval, *x* is taken as the mid-point of the age interval (e.g. 17.5 in the 15-19 year age group). The rate at which individuals wish to form new partnerships is calculated as

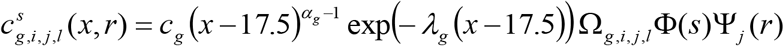

where *C_g_* is the desired rate in the baseline group (single, HIV-negative Africans in the high risk group who are aged 15-19), *λ_g_* and *α_g_* are the parameters of the gamma probability density function, Ω*_g,i,j,l_* is an adjustment factor taking into account the individual’s risk group and current relationship status, Φ(*s*) is an adjustment factor that takes into account the individual’s HIV status, and Ψ_*j*_(*r*) represents an effect of race on acquisition of secondary partners. The values assumed in the model are summarized in Table 4.3.1. Some of these parameter values were previously estimated by fitting a similarly-structured deterministic model to data on numbers of current sexual partners, by age and sex, in a nationally-representative 2005 survey [108]. (The calibration to sexual behaviour data made allowance for misreporting of partner numbers, as evidenced by inconsistencies in the numbers of current partners reported by men and women.) The sexual behaviour parameters were also partially determined based on the age and sex patterns of HIV prevalence in nationally representative household surveys and antenatal surveys [108, 186]. A full description of the model calibration procedure is provided elsewhere [187].

**Table 4.3.1:**
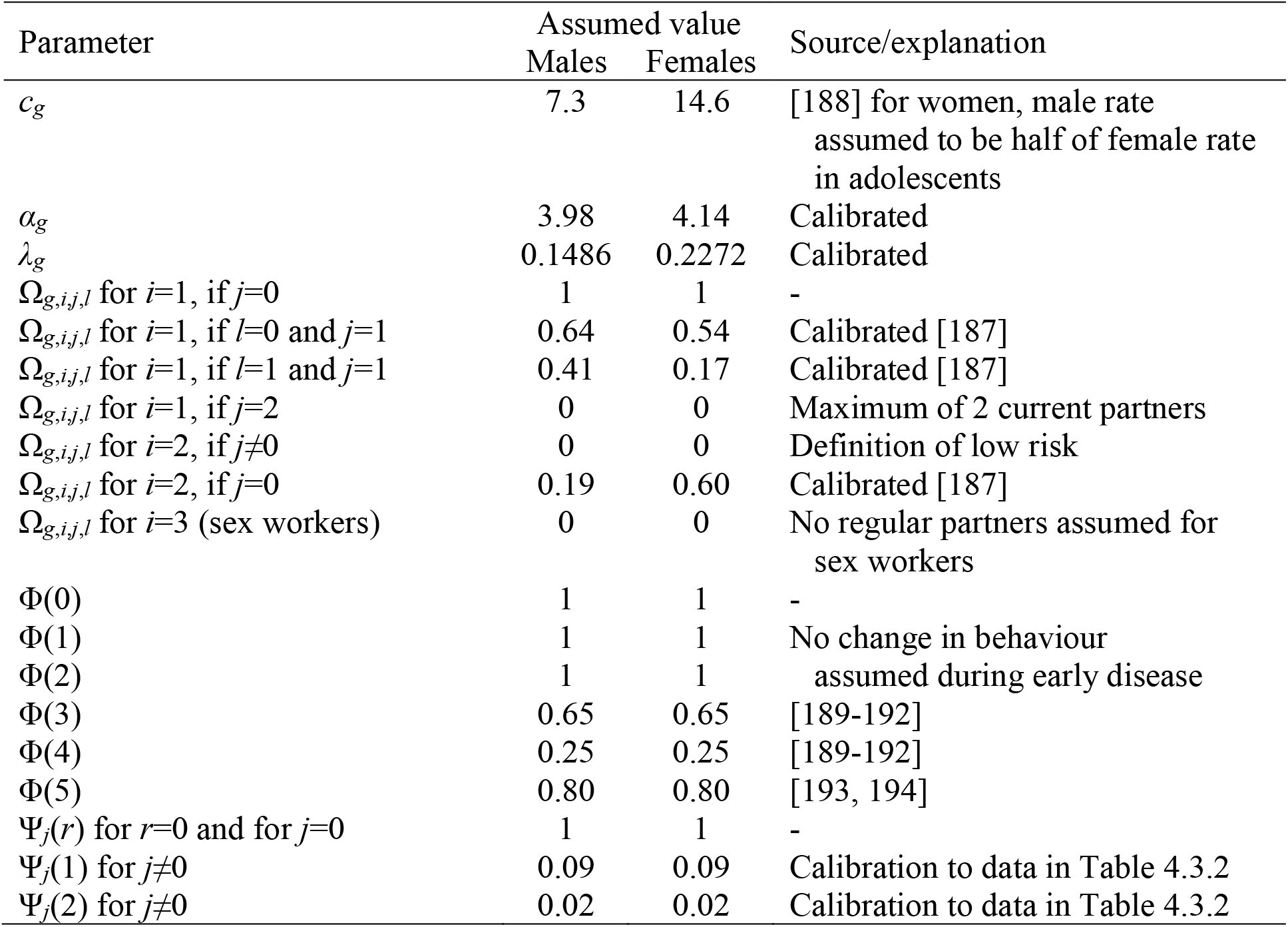
Parameters determining rates of short-term partnership formation, in sexually-experienced adults

Because male demand for new short-term partnerships with women may differ from female demand for new short-term partnerships with men, the model averages the demand of the two sexes when calculating the actual numbers of new short-term heterosexual partnerships formed in each period.

The previously-cited deterministic model [187] did not stratify the population by race, and it is therefore necessary to consider racial differences in partnership formation separately. Although overall numbers of partners appear roughly similar across risk groups, there is strong evidence of racial differences in the prevalence of concurrent partnerships, summarized in Table 4.3.2. Some estimates have been excluded from Table 4.3.2 because they relate only to women, and it would appear that women substantially under-report concurrent partnerships. The results show that the prevalence of concurrency among white South Africans is typically only one sixth of that in black South Africans, while that in coloured South Africans is about half of that in black South Africans. These racial differences may be due to cultural differences in the acceptance of concurrent partnerships [150], or may be due to differences in socioeconomic factors (for example, circular migration among black South Africans is a legacy of apartheid that may lead to increased concurrency).

**Table 4.3.2:**
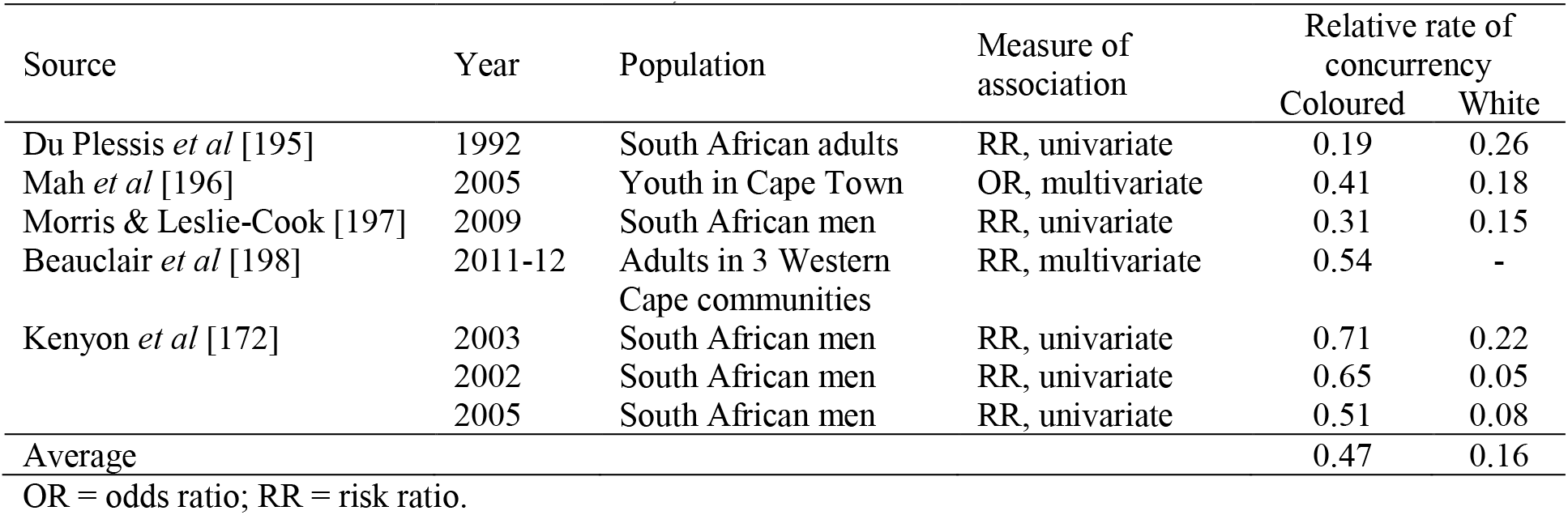
Rates of concurrent partnerships in coloured and white South Africans (as a fraction of rates in black South Africans)

The model assumes rates of primary partnership formation are the same across race groups, but allows for racial differences in the rate at which secondary partnerships are formed, among high risk individuals who are currently in relationships. The rates at which secondary partnerships are formed are multiplied by factors of 0.09 and 0.02 in coloured and white South Africans respectively, in order to produce model estimates of the relative prevalence of concurrency by race consistent with the average of the ratios estimated from empirical investigations (Table 4.3.2). These rate adjustments are expressed as hazard ratios, which is why they differ so substantially from the odds ratios and risk ratios shown in Table 4.3.2.

The model makes similar assumptions about the rates at which men in partnerships visit sex workers, since this can also be considered a form of concurrency. This means that although the rates at which single men visit sex workers are the same across race groups, the rates of sex worker contact among white men in relationships are assumed to be 0.02 times the corresponding rates in black men with the same relationship status, age and risk group. Similarly, the rates of sex worker contact among coloured men in relationships are 0.09 times the corresponding rates in black men with the same characteristics.

Although it is possible that rates of entry into short-term partnerships may differ for heterosexual men and MSM (even after controlling for differences in age and relationship status), we lack data to inform MSM-specific assumptions. South African studies suggest that individuals who identify as gay or bisexual have more sexual partners than individuals who identify as heterosexual [154, 159], but this could be because of differences in casual sex, which are accounted for separately (see section 4.7). It is therefore assumed that the desired rate of entry into short-term relationships is the same for MSM and heterosexual men. For bisexual men, the desired rate of entry into short-term relationships is split between men and women in proportion to their male preference parameter (parameter *Y_i_* in section 4.1).

### 4.4 Assortativeness of sexual mixing

Sexual mixing patterns are important in defining the patterns of disease spread in a population. If sexual mixing is random (i.e. individuals have no preferences regarding the characteristics of their partners), STIs will tend to spread relatively uniformly through the population. However, if sexual mixing is highly assortative (meaning that individuals tend to select partners with characteristics similar to their own), then STIs will tend to cluster in specific sub-populations.

#### 4.4.1 Mixing between education categories

International literature suggests that individuals tend to choose partners with levels of educational attainment similar to their own [199, 200]. In our model, we specify a relative probability of selecting an individual as a sexual partner if their educational attainment is different from the individual’s educational attainment (expressed as a fraction of the probability that would apply if the two individuals had the same educational attainment). Suppose that two individuals have educational attainment levels *i* and *j*, with values ranging from 0 (no education) to 13 (having a tertiary qualification). The relative probability of them forming a partnership is calculated as

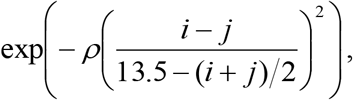

where the *ρ* parameter determines the extent of the educational assortativeness (zero implying that there is no assortativeness, i.e. individuals do not choose partners on the basis of their educational attainment). It is therefore assumed that the greater the difference between *i* and *j*, the lower is the likelihood of the individuals forming a partnership. However, the adjustment in the denominator of the exponent means that absolute differences in educational attainment are more important at high levels of educational attainment than at low levels of educational attainment.

In calibrating the model to South African data, we have set the *ρ* parameter to 1.5. Figure 4.4.1 compares the estimated fraction of married partners in each educational attainment category, for men at each education level. The model results are calculated for men who are married in 2011, and who are aged 40 or younger (the analysis is restricted to this age range to avoid any dependence on the baseline assumptions set in 1985, i.e. assuming that all of those men who were married in 2011 and aged 40 or younger got married after 1985). The model results are compared with the results of the 2011 General Household Survey (limiting the analysis to cases where there was a male household head with a spouse/cohabiting partner in the same house). The results of the model are roughly consistent with the survey, except in the case of men with tertiary qualifications, where the fraction of partners with tertiary education is under-estimated by some 20%. This might be because the model slightly under-estimates the fraction of the population with tertiary education (see Figure 3.5.5), or because the model ignores the fact that women have a higher rate of entry into and completion of tertiary education than men (see section 3.5.5).

**Figure 4.3.1:**
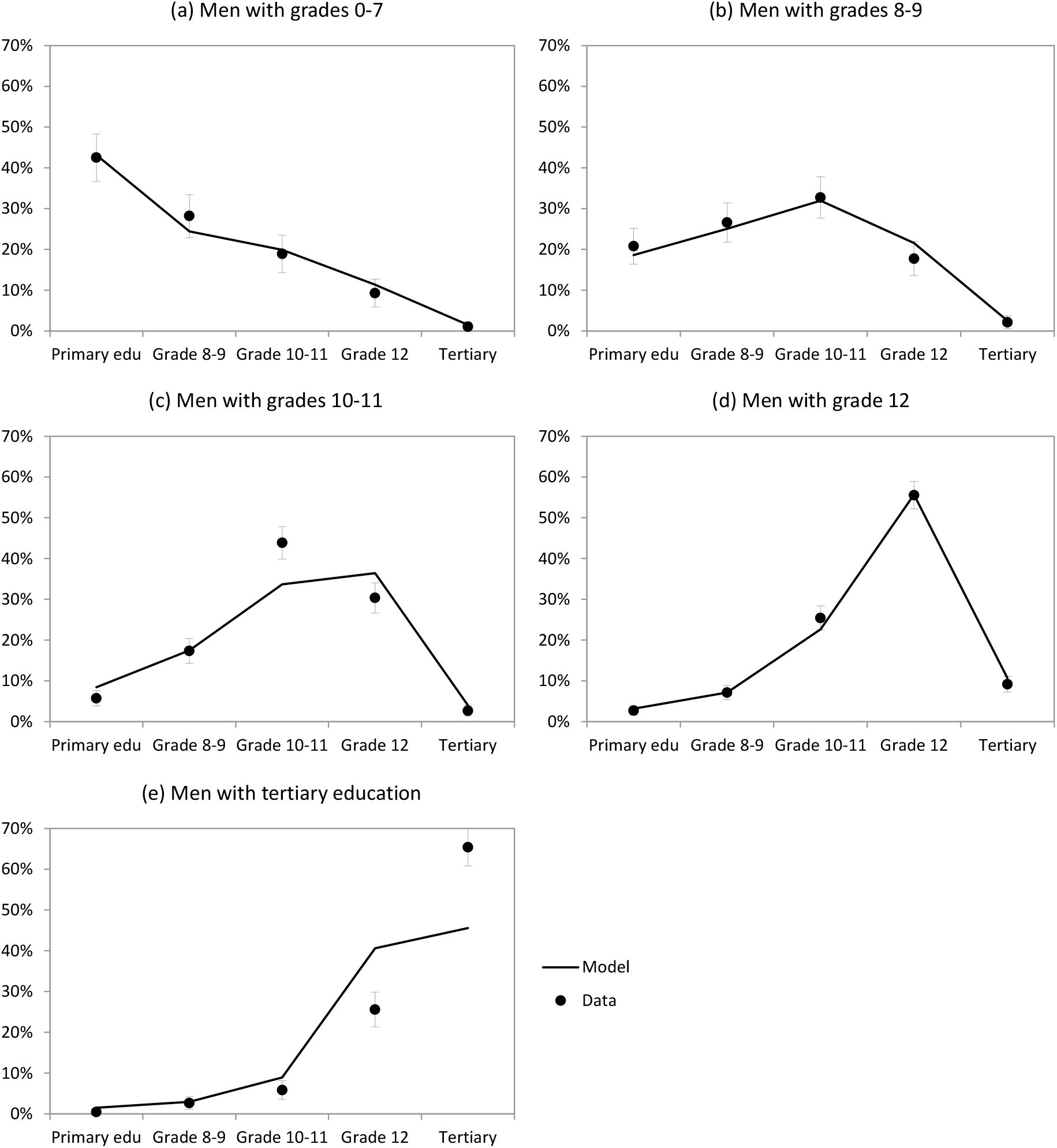
Fraction of partners with different education levels. The denominator in each panel is men who are married/cohabiting and aged 40 or younger in 2011; the numerator is the number of partners in each education category. Data are from the 2011 General Household Survey; error bars represent 95% confidence intervals around the survey estimates.

Although the above formula is applied when modelling all new partnerships that form after 1985, there is no adjustment to reflect assortativeness by education when simulating the initial population profile in 1985. This is unlikely to be a major problem for the purpose of modelling HIV transmission, as the short-term partnerships that exist in 1985 will mostly have broken up by 1990 (when HIV transmission is introduced into the simulated population), and the long-term relationships that exist in 1985 are unlikely to play an important role in HIV transmission after 1990 [187].

Educational attainment is assumed to have no effect on how clients select sex workers or on how MSM select casual partners.

#### 4.4.2 Mixing between race groups

South African studies show that sexual mixing by race is highly assortative, i.e. individuals almost always select partners from their own race group [201–203]. Up until 1985, the Immorality Act in South Africa prohibited all sexual interaction between whites and other race groups. The model therefore assumes that at the start of the simulation (in 1985) all partnerships are with individuals of the same race. Following 1985 (when the Immorality Act was repealed), the model allows for some degree of racial mixing in the selection of sexual partners. Similar to the approach taken in modelling assortativeness by education, we specify a relative probability of selecting an individual as a sexual partner if their race is different from the individual’s race (expressed as a fraction of the probability that would apply if the two individuals were of the same race). This relative adjustment has been set to 0.015, 0.003 and 0.0015 in black, coloured and white South Africans respectively. These parameters have been chosen so that the fractions of partners in different race groups are consistent with those observed in the 2011 General Household Survey (Figure 4.4.2). As before, the model results are calculated for men who are married in 2011, and who are aged 40 or younger.

**Figure 4.4.2:**
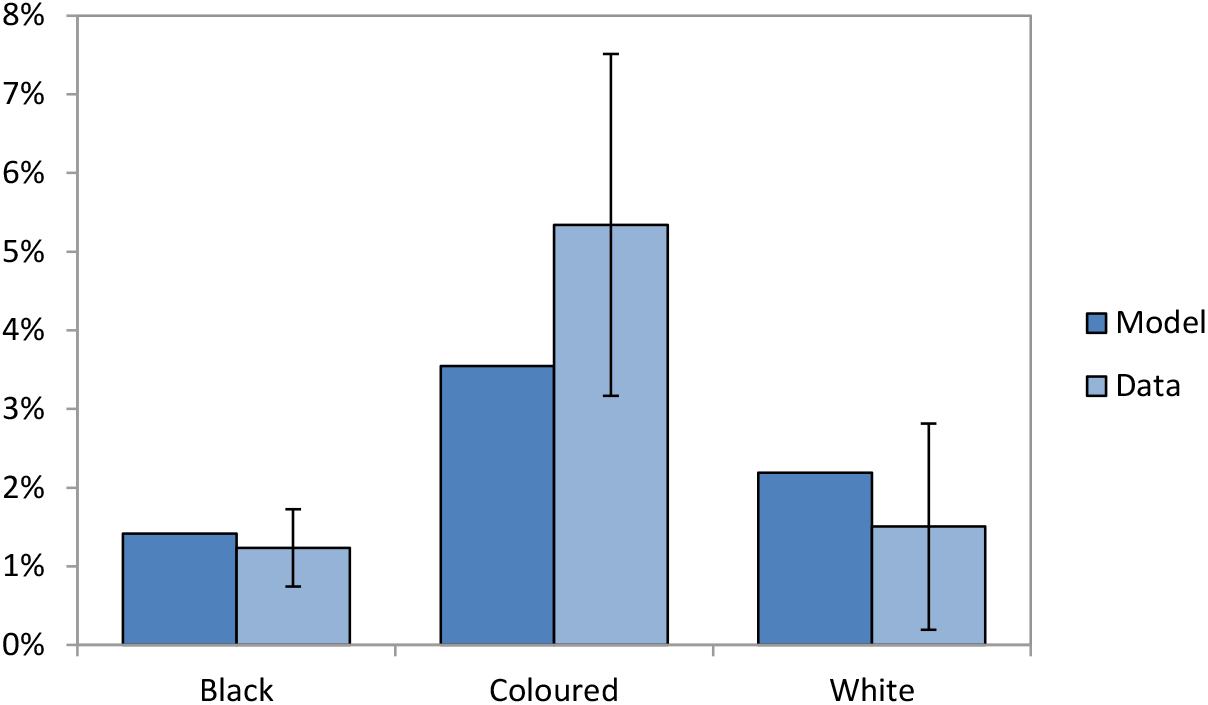
Fraction of men in relationships with partners of other race groups. The denominator in each bar is the number of men who are married/cohabiting and aged 40 or younger in 2011; the numerator is the number of primary partners who are not of the same race. Data are from the 2011 General Household Survey; error bars represent 95% confidence intervals around the survey estimates.

A weakness of this approach is that it assumes the assortativeness of mixing is the same in short-term and long-term partnerships, which might not be the case. The model is calibrated to data on the assortativeness of mixing in long-term relationships, which might not be typical of mixing patterns in short-term relationships.

For the sake of simplicity, we assume that sexual contacts between sex workers and their clients are completely assortative with respect to race. This may also be unrealistic, as there is anecdotal evidence of men engaging sex workers of other race groups [204]. In the context of MSM who engage in casual sex, we assume that sexual mixing is random with respect to race. This is because the relatively small size of the MSM casual sex pool means that it would be difficult to allocate casual partners to MSM of racial minorities (whites and coloureds) if we assumed their partner selection was highly assortative.

#### 4.4.3 Mixing between age groups

For heterosexuals, an age mixing matrix is specified for each sex. This mixing matrix determines the fraction of partners in each five-year age group, for individuals in each age group. Tables 4.4.1 and 4.4.2 show the age mixing matrices for women and heterosexual men respectively, which are the same as assumed previously [18, 187]. The female age mixing matrix is estimated based on the ages of spousal partners reported by women in the 1998 Demographic and Health Survey [30] and the age differences reported by women in non-spousal partnerships in smaller studies [100, 108, 205, 206]. The heterosexual male age mixing matrix has been calculated to be consistent with the female age mixing matrix.

**Table 4.4.1:** Percentage of women’s primary partners in each age group

| Female age | Age of male partner |  |  |  |  |  |  |  |  |  |  |  |  |  |  |  |
| --- | --- | --- | --- | --- | --- | --- | --- | --- | --- | --- | --- | --- | --- | --- | --- | --- |
|  | 10-14 | 15-19 | 20-24 | 25-29 | 30-34 | 35-39 | 40-44 | 45-49 | 50-54 | 55-59 | 60-64 | 65-69 | 70-74 | 75-79 | 80-84 | 85+ |
| 10-14 | 46.8 | 43.1 | 8.1 | 1.5 | 0.3 | 0.1 | 0.0 | 0.0 | 0.0 | 0.0 | 0.0 | 0.0 | 0.0 | 0.0 | 0.0 | 0.0 |
| 15-19 | 0.0 | 47.5 | 40.9 | 8.8 | 2.1 | 0.5 | 0.0 | 0.1 | 0.0 | 0.0 | 0.0 | 0.0 | 0.0 | 0.0 | 0.0 | 0.0 |
| 20-24 | 0.0 | 0.0 | 48.9 | 35.2 | 11.1 | 3.6 | 0.8 | 0.3 | 0.0 | 0.1 | 0.0 | 0.0 | 0.0 | 0.0 | 0.0 | 0.0 |
| 25-29 | 0.0 | 0.0 | 1.3 | 43.8 | 35.7 | 12.7 | 4.4 | 1.0 | 0.3 | 0.3 | 0.1 | 0.2 | 0.1 | 0.0 | 0.0 | 0.0 |
| 30-34 | 0.0 | 0.0 | 0.2 | 3.3 | 37.5 | 38.4 | 13.3 | 4.1 | 1.4 | 0.9 | 0.2 | 0.4 | 0.1 | 0.0 | 0.0 | 0.0 |
| 35-39 | 0.0 | 0.0 | 0.0 | 0.3 | 3.9 | 37.2 | 36.3 | 14.0 | 4.4 | 1.4 | 0.9 | 1.2 | 0.3 | 0.0 | 0.0 | 0.0 |
| 40-44 | 0.0 | 0.0 | 0.0 | 0.1 | 1.4 | 5.3 | 35.4 | 35.3 | 15.6 | 3.7 | 1.2 | 1.6 | 0.4 | 0.1 | 0.0 | 0.0 |
| 45-49 | 0.0 | 0.0 | 0.0 | 0.0 | 0.0 | 1.2 | 9.2 | 35.3 | 31.9 | 14.2 | 3.8 | 3.4 | 0.9 | 0.1 | 0.0 | 0.0 |
| 50-54 | 0.0 | 0.0 | 0.0 | 0.0 | 0.0 | 0.0 | 1.6 | 14.4 | 32.1 | 29.8 | 15.3 | 5.2 | 1.3 | 0.2 | 0.0 | 0.0 |
| 55-59 | 0.0 | 0.0 | 0.0 | 0.0 | 0.0 | 0.0 | 0.0 | 2.0 | 15.8 | 33.6 | 29.7 | 13.9 | 4.0 | 0.8 | 0.1 | 0.0 |
| 60-64 | 0.0 | 0.0 | 0.0 | 0.0 | 0.0 | 0.0 | 0.0 | 0.0 | 1.8 | 15.9 | 36.5 | 30.9 | 11.9 | 2.5 | 0.3 | 0.0 |
| 65-69 | 0.0 | 0.0 | 0.0 | 0.0 | 0.0 | 0.0 | 0.0 | 0.0 | 0.0 | 1.4 | 16.2 | 41.2 | 31.4 | 8.7 | 1.0 | 0.1 |
| 70-74 | 0.0 | 0.0 | 0.0 | 0.0 | 0.0 | 0.0 | 0.0 | 0.0 | 0.0 | 0.0 | 1.0 | 17.4 | 47.6 | 29.1 | 4.6 | 0.2 |
| 75-79 | 0.0 | 0.0 | 0.0 | 0.0 | 0.0 | 0.0 | 0.0 | 0.0 | 0.0 | 0.0 | 0.0 | 0.6 | 20.3 | 56.5 | 21.3 | 1.2 |
| 80-84 | 0.0 | 0.0 | 0.0 | 0.0 | 0.0 | 0.0 | 0.0 | 0.0 | 0.0 | 0.0 | 0.0 | 0.0 | 0.7 | 26.7 | 57.9 | 14.8 |
| 85+ | 0.0 | 0.0 | 0.0 | 0.0 | 0.0 | 0.0 | 0.0 | 0.0 | 0.0 | 0.0 | 0.0 | 0.0 | 0.0 | 1.4 | 35.0 | 63.6 |

**Table 4.4.2:** Percentage of men’s primary female partners in each age group

| Male age | Age of female partner |  |  |  |  |  |  |  |  |  |  |  |  |  |  |  |
| --- | --- | --- | --- | --- | --- | --- | --- | --- | --- | --- | --- | --- | --- | --- | --- | --- |
|  | 10-14 | 15-19 | 20-24 | 25-29 | 30-34 | 35-39 | 40-44 | 45-49 | 50-54 | 55-59 | 60-64 | 65-69 | 70-74 | 75-79 | 80-84 | 85+ |
| 10-14 | 100.0 | 0.0 | 0.0 | 0.0 | 0.0 | 0.0 | 0.0 | 0.0 | 0.0 | 0.0 | 0.0 | 0.0 | 0.0 | 0.0 | 0.0 | 0.0 |
| 15-19 | 10.8 | 89.2 | 0.1 | 0.0 | 0.0 | 0.0 | 0.0 | 0.0 | 0.0 | 0.0 | 0.0 | 0.0 | 0.0 | 0.0 | 0.0 | 0.0 |
| 20-24 | 0.8 | 30.7 | 66.5 | 1.8 | 0.2 | 0.0 | 0.0 | 0.0 | 0.0 | 0.0 | 0.0 | 0.0 | 0.0 | 0.0 | 0.0 | 0.0 |
| 25-29 | 0.1 | 5.6 | 40.9 | 49.8 | 3.3 | 0.2 | 0.1 | 0.0 | 0.0 | 0.0 | 0.0 | 0.0 | 0.0 | 0.0 | 0.0 | 0.0 |
| 30-34 | 0.0 | 1.4 | 13.5 | 42.5 | 38.7 | 3.0 | 0.9 | 0.0 | 0.0 | 0.0 | 0.0 | 0.0 | 0.0 | 0.0 | 0.0 | 0.0 |
| 35-39 | 0.0 | 0.4 | 4.7 | 16.4 | 43.0 | 31.2 | 3.6 | 0.7 | 0.0 | 0.0 | 0.0 | 0.0 | 0.0 | 0.0 | 0.0 | 0.0 |
| 40-44 | 0.0 | 0.0 | 1.2 | 6.9 | 18.0 | 36.6 | 29.6 | 6.7 | 1.0 | 0.0 | 0.0 | 0.0 | 0.0 | 0.0 | 0.0 | 0.0 |
| 45-49 | 0.0 | 0.1 | 0.6 | 1.8 | 6.4 | 16.3 | 34.0 | 29.6 | 10.0 | 1.2 | 0.0 | 0.0 | 0.0 | 0.0 | 0.0 | 0.0 |
| 50-54 | 0.0 | 0.0 | 0.1 | 0.7 | 2.6 | 6.3 | 18.4 | 32.9 | 27.3 | 11.0 | 0.7 | 0.0 | 0.0 | 0.0 | 0.0 | 0.0 |
| 55-59 | 0.0 | 0.0 | 0.2 | 0.9 | 2.2 | 2.5 | 5.4 | 18.4 | 32.0 | 29.7 | 7.9 | 0.5 | 0.0 | 0.0 | 0.0 | 0.0 |
| 60-64 | 0.0 | 0.0 | 0.0 | 0.4 | 0.8 | 2.2 | 2.4 | 6.4 | 21.4 | 34.2 | 23.8 | 8.2 | 0.3 | 0.0 | 0.0 | 0.0 |
| 65-69 | 0.0 | 0.0 | 0.0 | 0.9 | 1.6 | 3.4 | 3.7 | 7.0 | 8.7 | 19.2 | 24.3 | 25.3 | 5.8 | 0.0 | 0.0 | 0.0 |
| 70-74 | 0.0 | 0.0 | 0.0 | 0.4 | 0.7 | 1.5 | 1.7 | 3.1 | 3.7 | 9.6 | 16.1 | 33.1 | 27.3 | 2.7 | 0.0 | 0.0 |
| 75-79 | 0.0 | 0.0 | 0.0 | 0.2 | 0.3 | 0.6 | 0.6 | 1.1 | 1.8 | 4.7 | 8.3 | 22.5 | 40.9 | 18.3 | 0.8 | 0.0 |
| 80-84 | 0.0 | 0.0 | 0.0 | 0.0 | 0.0 | 0.0 | 0.0 | 0.0 | 1.4 | 3.6 | 5.4 | 13.6 | 32.4 | 34.7 | 8.7 | 0.2 |
| 85+ | 0.0 | 0.0 | 0.0 | 0.0 | 0.0 | 0.0 | 0.0 | 0.0 | 2.7 | 5.9 | 6.2 | 10.6 | 19.6 | 24.5 | 26.8 | 3.6 |

Few studies report on age mixing patterns in MSM relationships in the South African setting. Arnold *et al* [153] found that in 758 male-male sexual relationships in Soweto, the average partner age difference was small (0.25 years) but there was high variation in partner age differences (standard deviation of 5.8 years). Based on what is known about the age distribution of sexually active MSM in South Africa, it is possible to use this information to determine how patterns of age mixing vary in relation to age. If *S*(*x*) is the age distribution of sexually active MSM and *f*(*y*|*x*) represents the proportion of male partners aged *y* for an MSM aged *x*, then for a random sample of MSM, the expected proportion of their partners who are aged *y* is

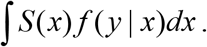

If the sample of sexually active MSM is truly representative, then we would expect that this proportion should be the same as *S*(*y*). We would also expect that

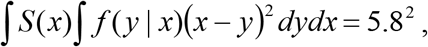

if the estimated standard deviation of 5.8 years [153] is correct. These constraints allow us to determine the likely patterns of sexual mixing. It is assumed that *S*(*x*) is a gamma distribution, with a mean of 25 years and a standard deviation of 7 years [153, 163], with an age offset of 14 years to prevent implausible levels of sexual activity in very young boys. The *f*(*y* | *x*) distribution is also assumed to be of gamma form, with mean of *μ*(*x*) = max(*x* –10, *x* + *A*(25 – *x*)) and variance of *B*^2^ (again, with an offset of 14 years to prevent sexual activity at young ages). The two free parameters, *A* and *B*, have been set to 0.45 and 5.0 years respectively, to yield a variance of partner age differences equal to 5.8^2^, as well as a distribution of 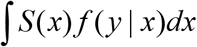 values roughly consistent with the distribution of *S*(*y*) values (Figure 4.4.3).

**Figure 4.4.3:**
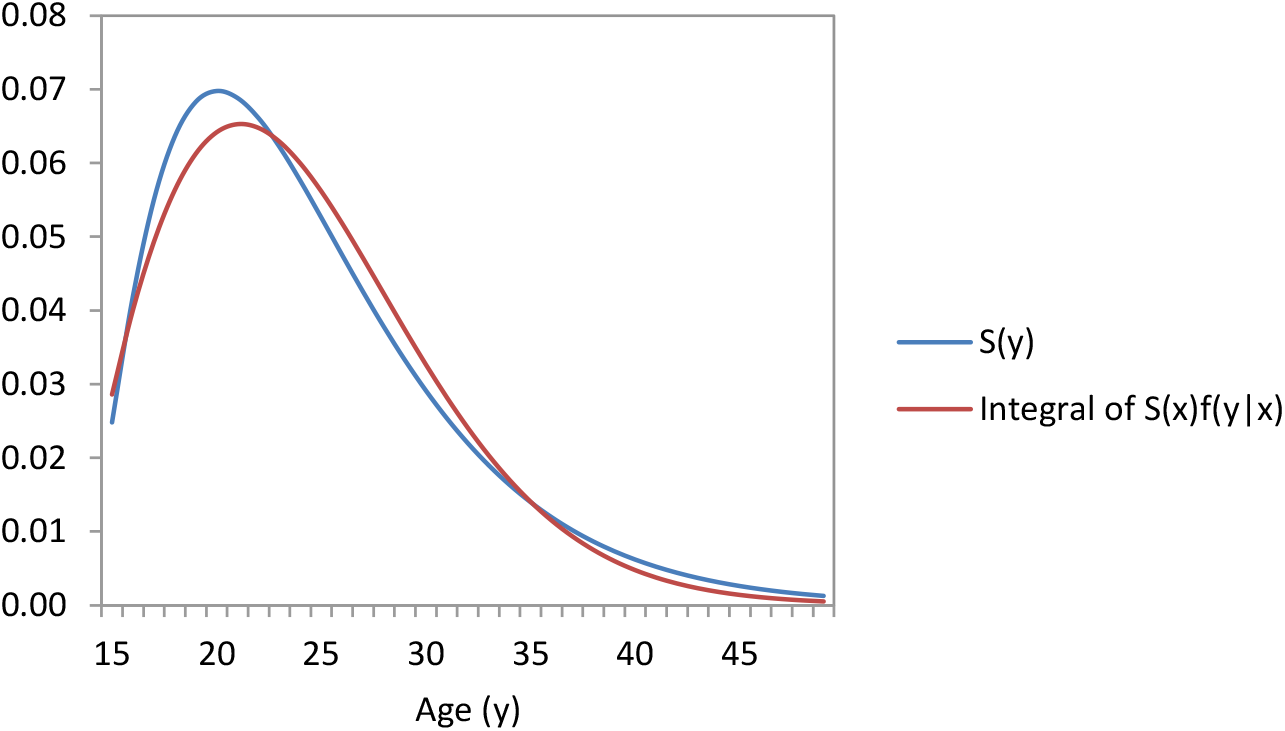
Age distribution of sexual activity in South African MSM

Using the above assumptions about the form of *f*(*y | x*), we calculate the proportion of sexual partners in each 5-year age group, for men in each age category (Table 4.4.3).

**Table 4.4.3:** Percentage of men’s male partners in each age group

| Male age | Age of male partner |  |  |  |  |  |  |  |  |  |  |  |  |  |  |  |
| --- | --- | --- | --- | --- | --- | --- | --- | --- | --- | --- | --- | --- | --- | --- | --- | --- |
|  | 10-14 | 15-19 | 20-24 | 25-29 | 30-34 | 35-39 | 40-44 | 45-49 | 50-54 | 55-59 | 60-64 | 65-69 | 70-74 | 75-79 | 80-84 | 85+ |
| 10-14 | 18.7 | 51.6 | 18.8 | 6.9 | 2.5 | 0.9 | 0.3 | 0.1 | 0.0 | 0.0 | 0.0 | 0.0 | 0.0 | 0.0 | 0.0 | 0.0 |
| 15-19 | 5.0 | 47.8 | 29.7 | 11.8 | 4.0 | 1.3 | 0.4 | 0.1 | 0.0 | 0.0 | 0.0 | 0.0 | 0.0 | 0.0 | 0.0 | 0.0 |
| 20-24 | 0.4 | 28.3 | 40.1 | 21.0 | 7.4 | 2.1 | 0.5 | 0.1 | 0.0 | 0.0 | 0.0 | 0.0 | 0.0 | 0.0 | 0.0 | 0.0 |
| 25-29 | 0.0 | 9.6 | 38.0 | 32.8 | 14.2 | 4.2 | 1.0 | 0.2 | 0.0 | 0.0 | 0.0 | 0.0 | 0.0 | 0.0 | 0.0 | 0.0 |
| 30-34 | 0.0 | 1.7 | 22.9 | 39.4 | 24.7 | 8.7 | 2.1 | 0.4 | 0.1 | 0.0 | 0.0 | 0.0 | 0.0 | 0.0 | 0.0 | 0.0 |
| 35-39 | 0.0 | 0.1 | 8.5 | 33.6 | 35.3 | 16.7 | 4.7 | 0.9 | 0.1 | 0.0 | 0.0 | 0.0 | 0.0 | 0.0 | 0.0 | 0.0 |
| 40-44 | 0.0 | 0.0 | 1.8 | 19.3 | 38.5 | 27.7 | 10.1 | 2.3 | 0.4 | 0.0 | 0.0 | 0.0 | 0.0 | 0.0 | 0.0 | 0.0 |
| 45-49 | 0.0 | 0.0 | 0.2 | 6.8 | 29.4 | 36.9 | 19.8 | 5.7 | 1.0 | 0.1 | 0.0 | 0.0 | 0.0 | 0.0 | 0.0 | 0.0 |
| 50-54 | 0.0 | 0.0 | 0.0 | 0.5 | 8.7 | 30.3 | 35.6 | 18.6 | 5.2 | 0.9 | 0.1 | 0.0 | 0.0 | 0.0 | 0.0 | 0.0 |
| 55-59 | 0.0 | 0.0 | 0.0 | 0.0 | 0.6 | 8.8 | 29.9 | 35.7 | 18.9 | 5.1 | 0.8 | 0.1 | 0.0 | 0.0 | 0.0 | 0.0 |
| 60-64 | 0.0 | 0.0 | 0.0 | 0.0 | 0.0 | 0.7 | 8.9 | 29.6 | 35.8 | 19.0 | 5.1 | 0.8 | 0.1 | 0.0 | 0.0 | 0.0 |
| 65-69 | 0.0 | 0.0 | 0.0 | 0.0 | 0.0 | 0.0 | 0.8 | 8.9 | 29.4 | 35.9 | 19.2 | 5.1 | 0.7 | 0.1 | 0.0 | 0.0 |
| 70-74 | 0.0 | 0.0 | 0.0 | 0.0 | 0.0 | 0.0 | 0.0 | 0.8 | 8.9 | 29.2 | 36.0 | 19.3 | 5.0 | 0.7 | 0.1 | 0.0 |
| 75-79 | 0.0 | 0.0 | 0.0 | 0.0 | 0.0 | 0.0 | 0.0 | 0.0 | 0.8 | 8.9 | 29.1 | 36.0 | 19.4 | 5.0 | 0.7 | 0.1 |
| 80-84 | 0.0 | 0.0 | 0.0 | 0.0 | 0.0 | 0.0 | 0.0 | 0.0 | 0.0 | 0.9 | 8.9 | 29.0 | 36.0 | 19.4 | 5.0 | 0.7 |
| 85+ | 0.0 | 0.0 | 0.0 | 0.0 | 0.0 | 0.0 | 0.0 | 0.0 | 0.0 | 0.0 | 0.6 | 6.2 | 21.7 | 32.0 | 24.3 | 15.2 |

The previously specified age preference matrices are assumed to apply when individuals select their primary partner. However, South African surveys suggest that different age preferences may apply in the context of concurrent partners (i.e. secondary partners). In a national youth survey, Steffenson *et al* [207] found that young men who reported recent concurrency were significantly more likely to report that their most recent partner was the same age or younger; conversely, young women who reported recent concurrency were more likely to report that their most recent partner was 5 or more years older (OR 1.6, 95% CI: 1.1-2.5). In a survey of young women in Carletonville in 1999, reporting concurrency was also significantly associated with reporting a partner 5 or more years older (aOR 2.5, 95% CI: 1.3-4.6) [208]. In the 2012 National Communication Survey, men who reported being in age-disparate relationships with women aged 16-24 were more likely to report concurrent partnerships, although this difference was not quite significant in multivariate analysis (aOR 1.39, 95% CI: 0.94-2.07) [107]. Similar results were obtained in the 2009 National Communication Survey, in which the age difference between men and their primary partners was substantially less than the age difference between men and their secondary partners [209]. In men aged 31-49, those who reported being in concurrent partnerships were significantly more likely to report an age gap of 10 or more years in respect of their secondary partner than in respect of their primary partner (OR 3.6, p < 0.01). Studies in Zimbabwe have also found associations between concurrency and age-disparate relationships [210].

To model the effect of concurrency on partner age differences, we assume that individuals who acquire secondary partners sample from a different partner age distribution to that from which primary partner ages are selected. Suppose that *F_g_*(*x|y*) represents the probability that an individual of sex *g* and age *y* selects a partner of age *x* or younger when choosing their primary partner. We sample a partner age *x* by randomly drawing a value *u* from the range (0, 1) and setting 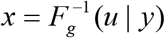. In selecting a secondary partner age, we follow a similar process, but replace *u* with *u^β^* in men and with *u*^1/*β*^ in women, where *β* is a parameter ≥1 that determines the extent of the change in age preference when selecting a secondary partner. The value of *β* has been set to 4.0, this value having being chosen such that the modelled odds ratio for the association between concurrency and age-disparate sex in partnered women aged 15-24 in 2000 was 1.57 (95% CI: 1.52-1.61), roughly consistent with the odds ratio reported by Steffenson *et al* [207] (1.6).

A limitation of this approach to defining age mixing patterns is that we have used data on age differences in *prevalent* relationships to determine age differences in *incident primary* relationships. It is possible that the two distributions may in fact be different, and it is therefore important to validate the model by comparing the modelled age distributions to actual data on age distributions. Figure 4.4.4 compares the modelled fraction of sexually active individuals whose partners are 5 or more years older or younger with corresponding data from the 2005 HSRC household survey [108]. The 2005 survey data have been used for this purpose as this is the survey with the most detailed reporting of partner age differences. The model is generally in good agreement with the data. However, the model over-estimates the fraction of young women (15-19) who report having a partner 5 or more years older (18.5% in the survey). This data point is almost certainly an outlier, as five other national surveys have found the proportion to be between 28% and 39% (average 33%) [25, 68, 184, 211].

**Figure 4.4.4:**
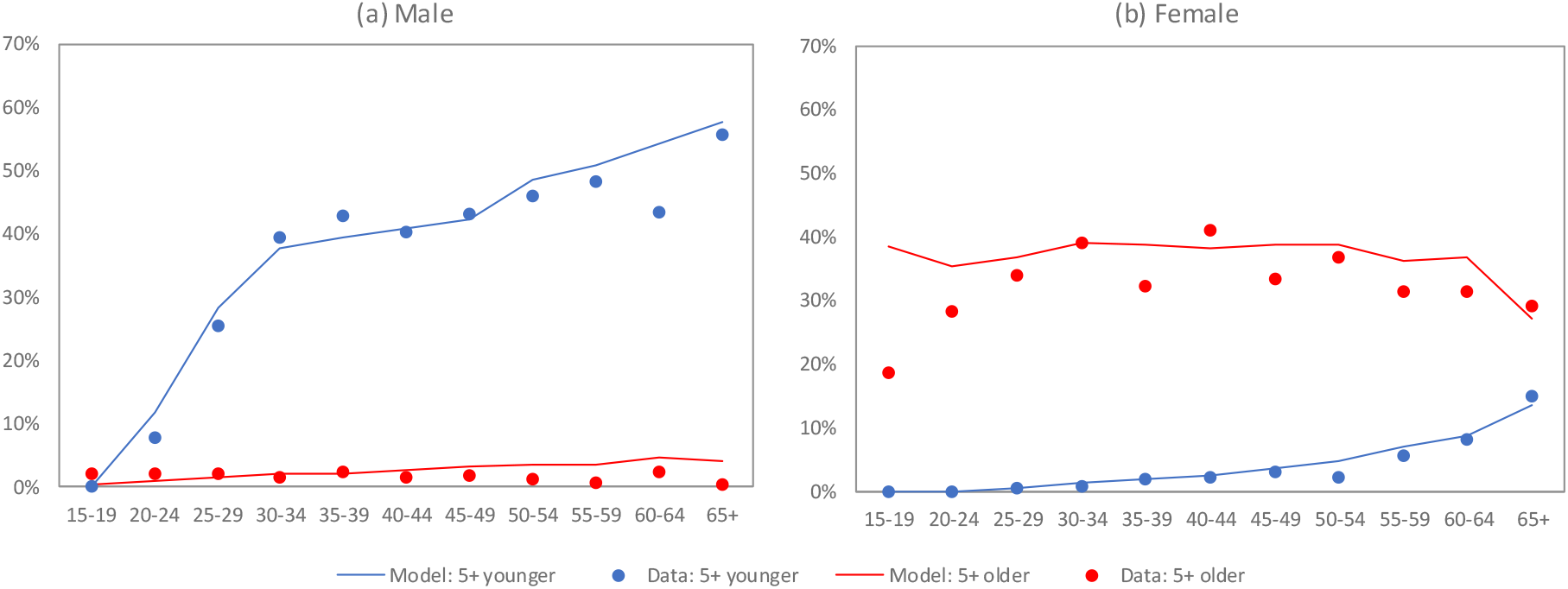
Proportions of sexually active individuals who are in age-disparate relationships. Data are from the 2005 HSRC household survey [108]. Model estimates are calculated in 2005; results presented are averages across 10 simulations.

Although there is some evidence to suggest that there may be differences in age mixing patterns by race [172, 198], the evidence is not consistent, and after averaging the results from different studies, racial differences in age preference appear to be relatively modest. We therefore do not allow for racial differences in age mixing in the current model.

#### 4.4.4 Mixing between urban and rural areas

The model allows for individuals to select partners from both urban and rural areas, regardless of their current location. However, the model assumes that an individual will be 94% less likely to choose an individual as their partner if they are not living in the same location (urban/rural) than they would be if they were living in the same location. This parameter has been chosen in such a way that the model matches the 1996 census data. In the census, 5.2% of married individuals were reported to be migrant workers; this is consistent with the model estimate of the fraction of married individuals who lived in urban areas while their partners resided in rural areas in 1996 (5.4%). Although the model definition does not exactly match the census definition of migrant worker (since some migrant workers might be working in rural areas, or urban migrant workers might be married to individuals in different urban centres), it is expected that the vast majority of married migrant workers would be working in urban areas while their partners resided in rural areas.

Note that although we apply this 94% penalty to reduce the likelihood of regular partnerships forming between individuals in urban and rural locations, we do not apply the same penalty to once-off sex acts between sex workers and clients, or between MSM engaging in casual sex. This is partly because the numbers of sex workers and MSM engaging in casual sex are relatively small, and splitting the respective risk pools between urban and rural areas may lead to small/empty pools from which individuals can choose casual/commercial sex partners, creating implausible scenarios. However, the assumption of no geographical penalty is also to some extent justified by the highly mobile nature of sex work in South Africa and the willingness of individuals to travel for once-off sexual encounters.

#### 4.4.5 Mixing between risk groups and by sexual orientation

The parameter *ρ*_1,*i,j*_(*t*) is defined as the desired proportion of new short-term partners who are in risk group *j*, for a man in risk group *i* at time *t*, who chooses a female partner. Mathematically, it is calculated according to the following formula:

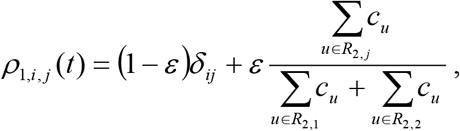

where *δ_ij_* = 1 if *i* = *j* and 0 otherwise, *ε* is the degree of sexual mixing, *R*_2,*j*_ is the set of women in risk group *j* and *c_u_* is the desired rate of short-term partnership formation in individual *u* (calculated as defined in section 4.3). The degree of sexual mixing can be any value from 0 to 1, with lower values of the parameter indicating greater tendency to form partnerships with individuals in the same sexual activity class. Similarly, the parameter *ρ*_2,*i,j*_(*t*) is defined as the desired proportion of new short-term partners who are in risk group *j*, for a woman in risk group *i* at time *t*:

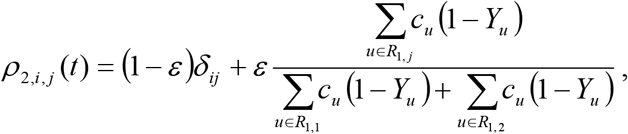

where *R*_1,*j*_ is the set of men in risk group *j*, and *Y_u_* is the male preference parameter for individual *u* (defined as the proportion of partners who are men). In the case of MSM, we define *ρ*_3,*i,j*_(*t*) as the desired proportion of new short-term partners who are in risk group *j*, for a man in risk group *i* at time *t*, who chooses a male partner:

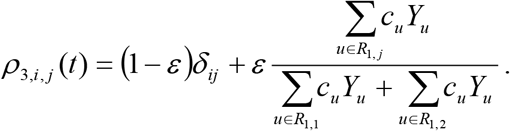

Because we lack data on patterns of mixing between high and low risk groups in South Africa, we assign the same value of *ε* to women, heterosexual men and MSM. The value of the *ε* parameter is difficult to estimate. Data from high-income settings suggest values of between 0.65 and 0.92 [212–215], and data from Botswana suggest a value of 0.53 [216]. However, it is difficult to estimate *ε* reliably from empirical data, and Ghani *et al* [217] demonstrate that sampling bias is likely to lead to significant overestimation of *ε*. Although our previous models of HIV in South Africa fit the HIV prevalence data best when values of *ε* were assumed to be around 0.5 [80, 187], these models were not stratified by race and educational attainment, and thus ignored two of the most important sources of assortativeness in sexual mixing. To represent the substantial uncertainty around the *ε* parameter, we therefore assign a prior distribution that is uniform on the interval (0, 1).

For MSM, mixing between risk groups is assumed to be random in the context of casual sex, i.e. MSM seeking casual partners are assumed not to select on the basis of the potential partner’s risk group.

### 4.5 Rates of marriage

Previous studies have shown that there are substantial differences in marriage rates between South African race groups [218, 219]. In the model, the rates at which individuals in short-term (non-cohabiting) partnerships marry their short-term partners (or equivalently, start cohabiting) is assumed to depend on a number of factors, including age, sex, number of partners, risk group and race. This section will start with an explanation of how the incidence of marriage is estimated (by age, sex and race), and will then be followed by an explanation of how the incidence rates are converted into probabilities of marriage per short-term partnership.

#### 4.5.1 Estimation of annual marriage incidence rates by age, sex and race

Suppose that _5_*m*_*g,r*_(*x*) is the probability that an unmarried individual of sex *g* and race *r*, who is aged between *x* and *x* + 4, gets married over the next 5 years. Further suppose that *P_g,r_*(*x, t*) is the proportion of individuals of sex *g* and race *r*, aged between *x* and *x* + 4, who are married at time *t*, and that _5_*d_g,r_*(*x*) is the probability that their marriage ends (either due to widowhood or divorce) over the next 5 years. It follows that

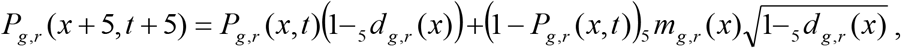

if we ignore the possibility of remarriage in the period immediately after a marriage ends, and if we assume that newly-married individuals get divorced/widowed at the same rate as individuals of the same age and sex who have been married for longer durations. Hence it is possible to estimate the probability of marriage in a given age cohort by comparing the proportion of the cohort that is married in two successive censuses, 5 years apart, if we know the rate of divorce/widowhood over the inter-census period:

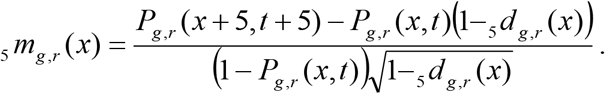

We have followed this approach in estimating the marriage rates from 1996 and 2001 census data. Estimates of divorce/widowhood rates are derived from previously-assumed age-specific divorce rates [187], previously-assumed age mixing patterns [187] and estimates of age-specific mortality probabilities in 1998-99 from the ASSA2008 AIDS and Demographic model [22] (the age mixing patterns and mortality probabilities together determine the probability of widowhood). Table 4.5.1 summarizes the resulting estimates of marriage probabilities (results are limited to the 15-59 age range, as estimates of marriage incidence at older ages are sensitive to inaccuracies in the estimated probabilities of widowhood). Female 5-year marriage probabilities are highest in the 25-29 age group, while male rates peak in the 35-39 age group. Rates of marriage are highest in white South Africans and lowest among black South Africans.

**Table 4.5.1:** Estimated probabilities of marriage in the 5 years between 1996 and 2001

| Age group<br>in 1996 | African |  | Coloured |  | White |  |
| --- | --- | --- | --- | --- | --- | --- |
|  | Male | Female | Male | Female | Male | Female |
| 15-19 | 0.0603 | 0.1795 | 0.1119 | 0.1920 | 0.1349 | 0.3008 |
| 20-24 | 0.2185 | 0.2964 | 0.3546 | 0.3987 | 0.4800 | 0.5998 |
| 25-29 | 0.3706 | 0.3416 | 0.4634 | 0.4107 | 0.6417 | 0.6221 |
| 30-34 | 0.4402 | 0.3177 | 0.4563 | 0.3895 | 0.6599 | 0.5343 |
| 35-39 | 0.4449 | 0.2444 | 0.4794 | 0.3390 | 0.6660 | 0.4413 |
| 40-44 | 0.4411 | 0.1935 | 0.4514 | 0.3032 | 0.5588 | 0.3861 |
| 45-49 | 0.4221 | 0.1063 | 0.3597 | 0.1823 | 0.5538 | 0.3492 |
| 50-54 | 0.3815 | 0.0435 | 0.3662 | 0.1456 | 0.5371 | 0.3044 |
| 55-59 | 0.3329 | 0.0000* | 0.1700 | 0.0509 | 0.4340 | 0.1910 |

\* Negative result has been reset to zero.

The microsimulation model assumes the effect of age on marriage incidence is the same for all races, and the average age effects are calculated from the race-specific estimates in Table 4.5.1. We do this by fitting logistic regression models to these estimates, separately for men and women. For the purpose of this regression, the five-year probabilities have been converted into annual probabilities, and zero values have been excluded from the regression. The results are summarized in Table 4.5.2. In converting the logistic regression model estimates into rates that apply at individual ages, we use the constant term from the logistic regression model and note that marriage rates in individuals aged 15-19 in 1996, over the 1996-2001 period, apply at age 20 on average (thus male rates peak at age 40 and female rates peak at age 30). Marriage rates are assumed to be zero at age 15 and are assumed to decline linearly after age 60.

**Table 4.5.2:** Age differences in marriage rates, and assumed baseline marriage rates

| Age<br>group | Logistic regression results (OR, 95% CI) |  | Age | Model assumptions |  |
| --- | --- | --- | --- | --- | --- |
|  | Males | Females |  | Males | Females |
| 15-19 | 1 | 1 | 15 | 0.0000 | 0.0000 |
| 20-24 | 4.18 (2.89-6.05) | 2.29 (1.43-3.66) | 20 | 0.0303 | 0.0873 |
| 25-29 | 6.92 (4.79-10.01) | 2.52 (1.58-4.03) | 25 | 0.1210 | 0.1897 |
| 30-34 | 7.57 (5.24-10.95) | 2.19 (1.37-3.49) | 30 | 0.1931 | 0.2070 |
| 35-39 | 7.85 (5.43-11.34) | 1.68 (1.05-2.68) | 35 | 0.2095 | 0.1817 |
| 40-44 | 6.81 (4.71-9.85) | 1.37 (0.86-2.19) | 40 | 0.2163 | 0.1493 |
| 45-49 | 5.98 (4.14-8.65) | 0.86 (0.54-1.38) | 45 | 0.1904 | 0.1180 |
| 50-54 | 5.66 (3.91-8.18) | 0.55 (0.34-0.88) | 50 | 0.1690 | 0.0757 |
| 55-59 | 3.52 (2.43-5.09) | 0.33 (0.19-0.55) | 55 | 0.1313 | 0.0488 |
|  |  |  | 60 | 0.1029 | 0.0293 |
|  |  |  | 65 | 0.0875 | 0.0251 |
|  |  |  | 70 | 0.0724 | 0.0209 |
|  |  |  | 75 | 0.0575 | 0.0167 |
|  |  |  | 80 | 0.0428 | 0.0125 |
|  |  |  | 85 | 0.0283 | 0.0083 |
|  |  |  | 90 | 0.0141 | 0.0041 |

The baseline marriage rates in Table 4.5.2 are multiplied by race-specific adjustment factors, which have been chosen in such a way that the modelled fractions of 15-64 year olds who are married are consistent with the corresponding estimates from the 2001 and 2011 censuses (Figure 4.5.1). These adjustment factors are 0.40 in Africans, 0.65 in coloureds and 1.30 in whites.

**Figure 4.5.1:**
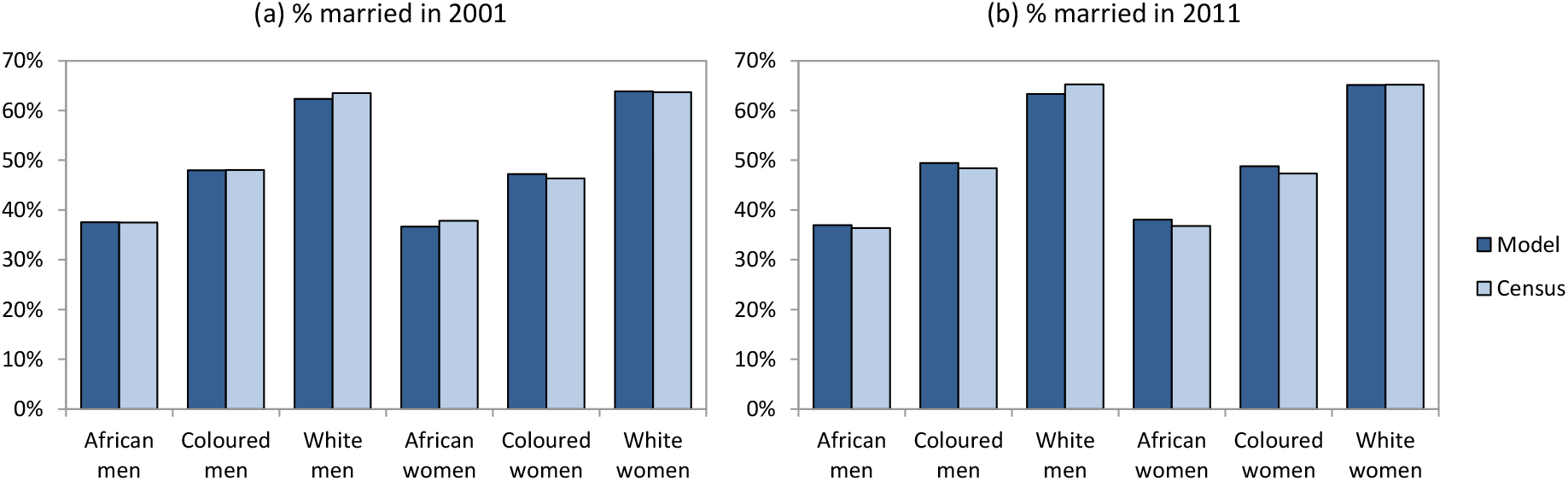
Fraction of 15-64 year olds who are in marital/cohabiting relationships. Model estimates are the median from 100 simulations.

A limitation of this approach is that it assumes the same age pattern for all race groups, though differences between race groups may be more pronounced in some age groups than in others. Figure 4.5.2 shows the calibration to the 2001 census data in more detail. Although the modelled age patterns are consistent with the census for Africans, there is less consistency in white and coloured men; the modelled fraction married is less than that estimated in the census in the 25-34 age range, but is more than that estimated in the census in the 45-64 age group. Similar patterns are seen in the 2011 census data (results not shown). This suggests that differences in male marriage rates by race might be more pronounced at younger ages than at older ages.

**Figure 4.5.2:**
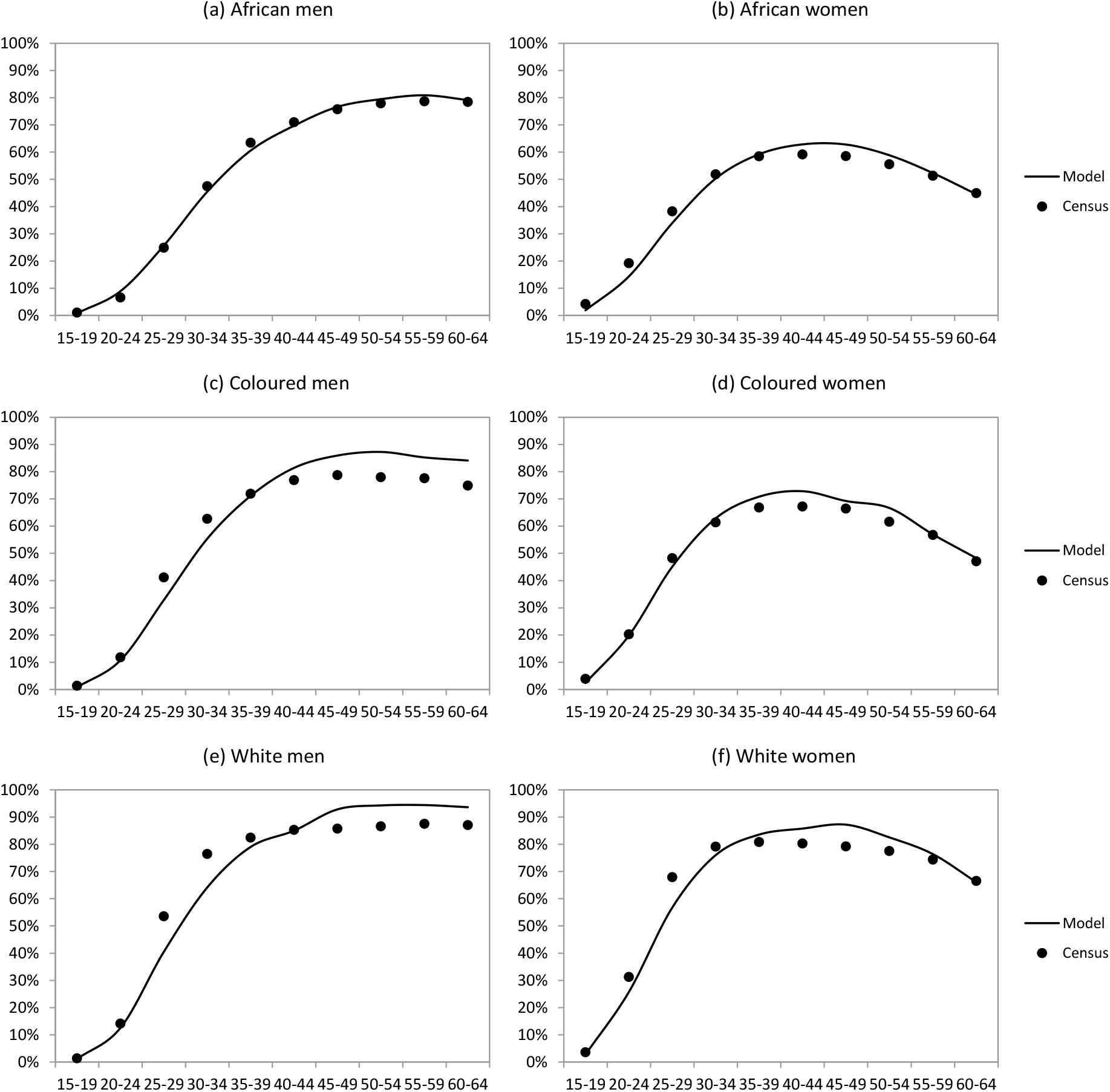
Fraction of individuals who are in marital/cohabiting relationships in 2001. Model estimates are the median from 100 simulations.

#### 4.5.2 Effect of education

The model assumes that after controlling for age, race and sex, educational attainment has no effect on rates of marriage. This appears to be plausible, as the model estimates of the fraction married, stratified by highest educational attainment, are consistent with the census estimates (Figure 4.5.3), even though the model makes no allowance for an independent effect of education on marriage. To the extent that there are differences in the fraction married by level of educational attainment, these are probably explained by age and race differences in educational attainment.

**Figure 4.5.3:**
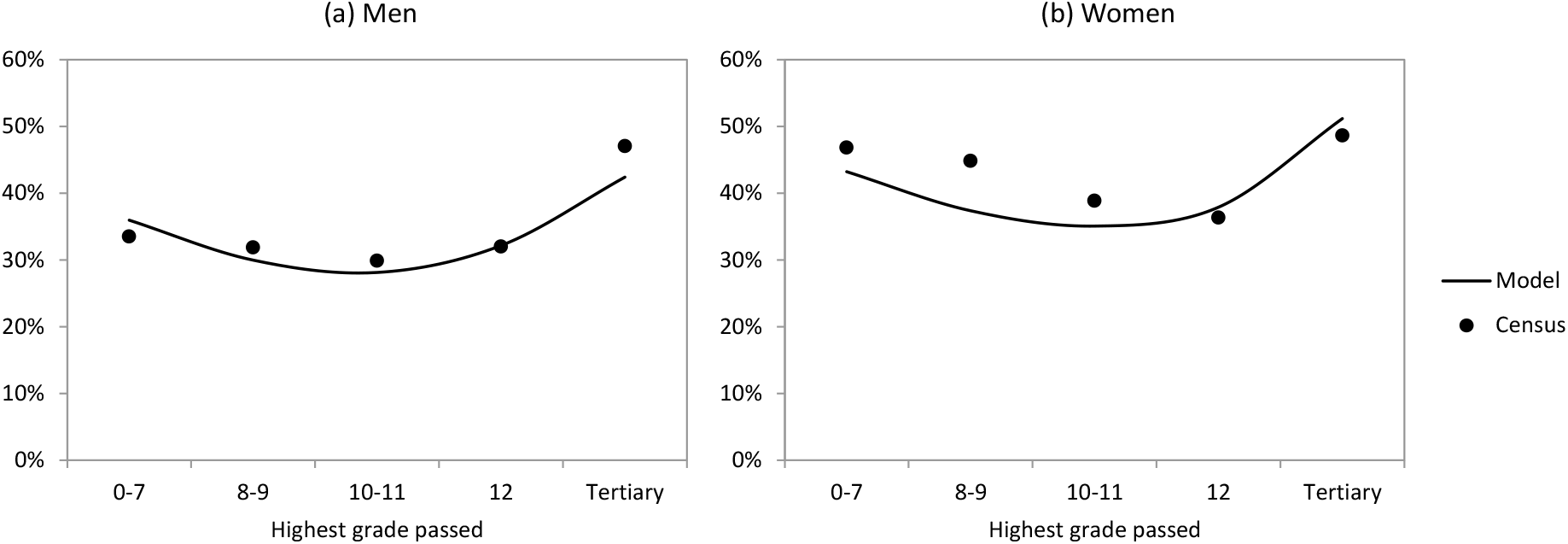
Fraction of 20-39 year olds who are married in 2011

#### 4.5.3 Effect of risk group on marriage incidence rates

An obvious approach to converting the previously-specified marriage incidence rates into rates that apply per short-term partnership is to divide the expected numbers of new marriages (in each age, race and sex category) by the corresponding number of unmarried individuals in short-term relationships. However, this would mean that high risk individuals have a greater chance of getting married, because they tend to have more short-term partnerships. There is no evidence to suggest that this is the case (if anything, evidence suggests the opposite [112, 150], although the interpretation of the evidence is unclear because some of the association between marriage and low risk behaviour may be due to an effect of marriage on high risk behaviour, not the other way around). Therefore we modify the calculations to ensure that the overall incidence of marriage is approximately the same in the high and low risk groups.

#### 4.5.4 Marriage and same-sex relationships

The previously-specified marriage assumptions relate to heterosexual couples. For same-sex couples, it is likely that rates at which short-term relationships transition to cohabiting/marital relationships are lower than in heterosexual couples, as progressing to a cohabiting or marital relationship implies a degree of openness about one’s relationship status. Although South Africa legislation allows for same-sex marriage, such openness may not be possible in communities in which there are high levels of homophobia [220]. This was confirmed in a recent South African study, which found significantly lower rates of marriage in MSM when compared to heterosexual men [221]. We therefore reduce the heterosexual marriage rates by a constant factor when modelling the rate at which short-term relationships between MSM become marital. This factor has been set to 0.48, the average of the 100 best-fitting parameters obtained when the model was fitted to behavioural data from South African surveys of MSM [160].

### 4.6 Divorce and partnership dissolution

Regular partnerships can be terminated through death of either partner, through ‘divorce’ (in the case of marital/long-term relationships) or through ‘break up’ (in the case of short-term relationships). For short-term heterosexual relationships, the annual rate of dissolution has been set at 2, which implies an average duration of short-term relationships equal to 6 months, roughly consistent with average durations of 3-12 months observed in African studies of non-spousal relationships [188, 222, 223]. For long-term heterosexual relationships, the rates of relationship dissolution have been estimated by multiplying estimated rates of divorce in 2004 (by age and sex) by a factor of 2 [224]. (This upward adjustment makes allowance for the fact that rates of dissolution are higher in cohabiting non-marital relationships than in marital relationships, and many married individuals are separated although not formally divorced.) These assumptions about relationship break-up are the same as in our previous models of sexual behaviour in South Africa [18, 187].

There is limited information on rates of relationship dissolution in South African MSM. Arnold *et al* [153] found that among MSM in Soweto, the mean duration of reported relationships (over a 6-month period) was 2.5 months. This is almost certainly an under-estimate of the duration that would be expected in short-term relationships, (a) because men could not report durations of more than 6 months, and (b) because reporting included casual sex encounters (which would in most cases have been of very short duration). The average duration of short-term relationships in MSM has been set to 0.48 years (5.8 months), the average of the 100 best-fitting parameters obtained when the model was fitted to behavioural data from surveys of South African MSM [160].

Although same-sex marriage has been legal in South Africa since 2005, published marriage and divorce statistics do not disaggregate heterosexual and same-sex relationships [225]. It is therefore assumed that rates of divorce/separation in long-term same-sex relationships are the same as in long-term heterosexual relationships.

### 4.7 Casual sex

It is assumed that MSM intermittently go through phases of engaging in casual sex (defined as once-off sex acts). This is modelled by assigning MSM a casual sex ‘indicator’ (1 if the individual is regularly engaging in casual sex, and 0 otherwise). The rate at which MSM enter this casual sex phase is assumed to depend on their age, risk group, relationship status and the extent of their same-sex preference. Mathematically, the annual rate of entry into the casual sex ‘phase’ is calculated as

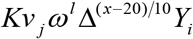

where *K* is the ‘base’ rate (applying to single high-risk gay men aged 20), *v_j_* is the effect of being in risk group *j* (1 for the high risk group), *ω* is the effect of being in a regular relationship (*l* = 1), Δ is the factor by which the rate of entry into casual sex is reduced per 10-year increase in age (*x*), and *Y_i_*, as before, is the male preference value for individual *i*. The annual rate at which men leave the casual sex phase is calculated as

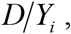

where *D* is the rate of exit for gay men.

Little local information is available on the frequency of casual sex in MSM. International literature shows that engagement in casual sex is significantly reduced in MSM who have regular partners [149, 226] but is only weakly associated with age [149, 226, 227]. International literature also suggests that men do not remain in the casual sex ‘phase’ for long periods of time [227]. In other accounts, gay men describe casual sex as a phase that precedes more regular relationships [228]. We have fixed the *D* parameter at 1 (i.e. assuming that gay men remain in the casual sex phase for a year on average).

The values assigned to the remaining parameters are summarized in Table 4.7.1. All of these parameters have been calculated as the average of the 100 best-fitting parameters obtained when the model was fitted to behavioural and HIV prevalence data from South African surveys of MSM [160]. As might be expected, low-risk MSM are estimated to have a lower rate of entry into casual sex than high-risk MSM. Consistent with the international literature [149, 226], the model assumes that men in regular relationships are less likely to engage in casual sex than men who are single. There is substantial uncertainty regarding the effect of age on entry into casual sex. Although international studies suggest limited association between age and casual sex [149, 226, 227, 229], Sandfort *et al* [202] found that in South African MSM, the age distribution of men whose last partner was a ‘casual neighbourhood’ partner was significantly older than that of men in other relationship categories, and the model assumption of a positive relationship between age and casual sex is consistent with this.

**Table 4.7.1:** Assumed rates of entry into casual sex

| Parameter description | Symbol | Value |
| --- | --- | --- |
| Annual rate of entry into casual sex in single high risk gay men aged 20 | $K$ | 5.20 |
| Relative rate of entry into casual sex in low risk men | $v_2$ | 0.59 |
| Relative rate of entry into casual sex in men who have a regular partner | $\omega$ | 0.62 |
| Relative rate of entry into casual sex per 10-year increase in age | $\Delta$ | 1.07 |

### 4.8 Coital frequencies and types of sex act

In short-term heterosexual relationships, sex is assumed to occur at a rate of 3 times per month on average, based on South African studies reporting on coital frequencies among youth [206, 230, 231]. In long-term heterosexual relationships, coital frequencies are assumed to depend on the age and sex of the individual. In married women, coital frequencies are assumed to reduce exponentially from a rate of 5 per month in the 20-24 age group, declining by a factor of 50% for each 20-year increase in age (i.e. reducing to 2.5 times per month in 40-44 year olds, to 1.25 times per month in 60-64 year olds, etc.). Although the parameters are specified for married women, coital frequencies in married men are calculated to be consistent with the assumed female frequencies, taking into account the age mixing matrices, and it is the age of the male partner that is assumed to determine the frequency of sex in any given married relationship. These assumed frequencies of sex in spousal and non-spousal partnerships result in numbers of sex acts that are roughly consistent with the aggregate reported coital frequencies in the 15-24 and 25-49 age bands in a 2005 national household survey [108]. In heterosexual relationships, all sex acts are assumed to be penile-vaginal sex acts.

In modelling sex between MSM, we distinguish between ‘sexual episodes’ and ‘sex acts’, since a single sexual episode may consist of multiple sex acts (e.g. insertive and receptive anal intercourse, each of which carries a different transmission risk). South African data suggest that many sexual episodes do not involve anal intercourse [153, 158, 232]. A possible explanation for the low rates of engagement in anal intercourse is HIV risk mitigation, since oral and other forms of sex carry negligible HIV transmission risk. Another possible explanation for the low rates of anal intercourse may be incompatibility in role preferences. South African studies show that MSM have strong role preferences, with relatively low rates of role versatility [153, 163]. Role preference appears to be strongly related to the way in which MSM self-identify; for example Lane *et al* [232] found a strong association between bisexual identity and exclusive practice of insertive anal intercourse (*r* = 0.9). Partners with incompatible role preferences may prefer to engage in non-anal intercourse.

To model the heterogeneity in role preference, we assign to each MSM a role preference indicator. In bisexual men, the probabilities of being assigned to the exclusively insertive, versatile and exclusively receptive categories are 0.5, 0.3 and 0.2 respectively, while the corresponding proportions in gay men are 0.2, 0.3 and 0.5 respectively. It is assumed that MSM do not choose their partners on the basis of their role preference, but that role preferences determine the type of sex and frequency of sex. If both partners are versatile, a single sexual episode is assumed to involve acts of both insertive and receptive anal intercourse with 50% probability (i.e. there is an average of 1.5 sex acts per sexual episode). Although we lack South African data to support this assumption, it is roughly consistent with data from North America, South America and Australia [233–235]. If both partners are exclusively insertive, or both are exclusively receptive (i.e. incompatible role preferences), no anal intercourse occurs. For all other permutations of role preference, a single sexual episode is assumed to involve a single anal sex act (the nature of which will be determined by the partner with the exclusive preference).

There is also a lack of data on the frequency of sexual episodes among MSM in the South African context. In a study of MSM in Soweto, which was limited to partnerships that involved anal sex, the average number of anal sex episodes was 10.28 per partnership (over a 6-month period) in 758 partnerships [153]. Only 284 of the relationships were reported to be regular relationships; if it is assumed that the remaining 474 partnerships were all casual ‘once-off’ partnerships, then the total number of anal sex episodes with regular partners must have been 7318 (758 × 10.28 – 474), which is equivalent to 25.8 anal sex episodes per regular partnership over 6 months (or 4.3 anal sex episodes per month, assuming regular partnerships were sustained over the full 6 months). This is similar to the mean coital frequency assumed in regular heterosexual relationships. This is congruent with data from high-income countries, which show similar coital frequencies in heterosexual and gay relationships [229]. We therefore assume that the frequencies of sexual episodes in short- and long-term MSM relationships are the same as the corresponding frequencies of vaginal sex in heterosexual relationships.

There is very little data on the frequency of casual sex for men who are engaging in casual sex. Data from high income settings suggest that the frequency of sexual episodes in the context of casual partnerships is roughly half of that in the context of regular partnerships [229]. The assumed monthly frequency of anal sex episodes in MSM who are engaging in casual sex is set to 1.56, the average of the 100 best-fitting parameters obtained when the model was fitted to HIV and behavioural data from South African studies of MSM [160].

No assumptions are made about the frequency of oral sex, as this is assumed to carry negligible HIV transmission risk [10].

### 4.9 Condom usage

Rates of condom usage are assumed to depend on a number of factors, including age, sex, education, race, type of relationship, knowledge of HIV status and calendar period. The sections that follow describe the approach to modelling each of these effects. In many cases, these assumptions are based on our previous deterministic modelling of condom usage in South Africa [80, 236]. However, a key change to the previous deterministic modelling approach is that we allow for heterogeneity between relationships in consistency of condom use. In our deterministic model, we assumed that all individuals of the same age, sex and relationship status have the same probability of condom use in any given sex act. In reality, the probability that a condom is used in any given sex act is likely to be strongly determined by whether a condom was previously used at the last sex act with the same partner. South African studies show that condoms tend to be used in the early stage of a relationship before being discontinued once the relationship is well-established [153, 237–239], and the assumption that condom use at any sex act is independent of that at preceding sex acts is likely to be unrealistic.

In the individual-based model, each individual is assigned three variables:

- a ‘condom preference’ variable, which represents the extent to which the individual regards condom use as acceptable, and is able and willing to use condoms (a value of 1 implies that the individual has the same condom acceptance as the national average for individuals of their age and sex);
- a ‘condom primary’ variable, which represents the fraction of sex acts with their current primary partner that are protected (this variable will be 0 or 1, implying non-use or consistent use respectively, depending on whether the relationship is in its early or more stable phase and depending on other factors such as exposure to HIV communication programmes);
- a ‘condom secondary’ variable, similarly defined for the individual’s relationship with their secondary partner (if they have a secondary partner).

The condom preference parameter is assumed to depend on the individual’s educational attainment, race and HIV diagnosis (as described below). This parameter is fixed at the individual level, although it can change when the individual’s educational attainment increases or if the individual learns that they are HIV-positive. For individual *i*, the condom preference parameter is calculated as Ω*_i_R*(*r_i_*)*E*(*g_i_*)*D_i_*, where Ω_*i*_ is the individual’s innate condom preference, *R*(*r_i_*) is an adjustment factor to represent the effect of the individual’s race (*r_i_*) on their odds of condom use, *E*(*g_i_*) is similarly defined as an adjustment factor to represent the effect of the individual’s educational attainment (*g_i_*) on their odds of condom use, and *D_i_* is an adjustment to represent the effect of HIV diagnosis. Heterogeneity in the innate condom preference is modelled by sampling Ω_*i*_ from a gamma distribution with a mean of 1. The standard deviation of the gamma distribution has been set at 1.4, which has been chosen so that the simulated correlation in condom use across multiple partnerships is roughly consistent with that observed in two South African studies [238, 239].

#### 4.9.1 Condom use in short-term relationships

At the time a new partnership is formed, we calculate the probability that a condom is used at the first sex act. This probability is calculated as the average of the probabilities determined for each partner.

The parameter *γ*_2_(*x, t*) represents the probability that a woman aged *x* uses a condom in her first act of sex with a new partner at time *t*, i.e. the population average before taking into account the individual-specific condom preference mentioned previously. This parameter is calculated in relation to a ‘baseline’ rate of condom usage, *γ*\*, which is the probability of condom use for a woman aged 15-19 in a short-term relationship in 1998 (1998 has been chosen as the baseline because it is the year for which the most condom usage data are available, and because there is little reliable data on condom usage prior to 1998). The following formula is used to calculate *γ*(*x, t*):

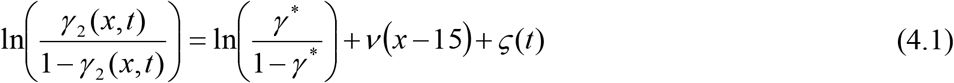

where exp(*ν*) is the factor by which the odds of condom use reduces, per year of age, and exp(*ς*(*t*)) is the odds of using a condom in year *t*, relative to that in 1998. The *ς*(*t*) function is a linear combination of a constant term and two cumulative Weibull distribution functions. The constant term represents the initial rate of condom usage, prior to the start of the HIV epidemic in South Africa, the first Weibull distribution corresponds to the increase in condom usage following the introduction of HIV communication programmes in the mid-1990s, and the second Weibull distribution represents the reversal in condom usage rates in recent years. In mathematical terms,

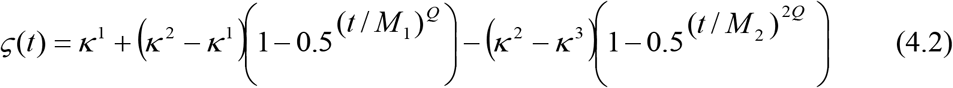

where *t* is time in years since 1985, and the other variables are defined as follows:

*κ*^1^ represents the initial rate of condom use in 1985 (relative to the baseline in 1998);
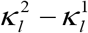 represents the increase in condom use, following initial HIV communication programmes;
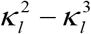 represents the reduction in condom use, following reductions in condom promotion/risk compensation;
*M* = the median for the first Weibull distribution;
*M*_2_ = the median for the second Weibull distribution;
*Q* = the Weibull shape parameter controlling the speed of behaviour change.

The logistic transformation prevents rates of condom use greater than 100%, and facilitates a ‘logistic regression’ interpretation of the condom parameters. Based on logistic regression models fitted to data on condom usage in the 1998 and 2003 South African DHSs [30, 36], it is assumed that the parameter *ν* is −0.025 (i.e. an odds ratio of 0.975 for the effect of each year of increase in age on the odds of condom use). The proportion of African women reporting condom usage for contraceptive purposes was found to be 0.13% in the 1987-89 DHS [240], compared to 1.8% in the 1998 DHS, and on the basis of this information, the ratio of the initial odds of condom use to that in 1998 (exp(*κ*^1^)) is assumed to be 0.07.

The parameters *γ*\*, *κ*^2^ and *Q* have been set separately for two scenarios: a scenario in which women are assumed to report accurately on their levels of condom use, and a scenario in which women are assumed to overstate their levels of condom use substantially. A condom reporting bias parameter, *θ*, is used to interpolate linearly between the parameter values in these two scenarios, with *θ* = 0 corresponding to the scenario in which there is no bias and *θ* = 1 corresponding to the scenario in which there is substantial over-reporting of condom use. The assumed parameter values for the two scenarios are summarized in Table 4.9.1. Parameters in the ‘no bias’ scenario were chosen so that the modelled proportions of young women using condoms were reasonably consistent with data on the proportion of young women reporting having used a condom the last time they had sex [25, 30, 68, 69, 108, 109, 184, 211]. Parameters in the ‘high bias’ scenario were chosen so that the modelled proportions of young women using condoms were consistent with proportions of sexually active women who reported using condoms for contraceptive purposes in the Demographic and Health Surveys (on the assumption that these would be less affected by social desirability bias and would represent a minimum on the true rate of condom use). Although the assumed relative level of condom usage in the early stages of the epidemic is the same in all scenarios, this parameter has little influence on HIV incidence, and potential bias in the estimation of these parameters is therefore of little consequence. The parameter *M*_2_ has been set to 26 (i.e. assuming the median time of the reversal in condom use is 2011, 26 years after the start of the simulation), and the *κ*^3^ value has been set to 75% of *κ*^2^. These values have been chosen to ensure approximate model consistency with the observed trends in condom usage (as shown in Figure 4.9.1).

**Table 4.9.1:** Differences in condom usage parameters between scenarios

| Parameter | Symbol | No bias scenario<br>($\theta = 0$) | High bias scenario<br>($\theta = 1$) |
| --- | --- | --- | --- |
| Probability of condom use in women aged 15-19, in short-term relationships, in 1998 | $\gamma^*$ | 0.35 | 0.08 |
| Ultimate odds of condom use, relative to the odds of condom use in 1998 | $\exp(\kappa^2)$ | 20 | 3 |
| Shape parameter controlling the speed of behaviour change | $Q$ | 4.0 | 2.5 |

**Figure 4.9.1:**
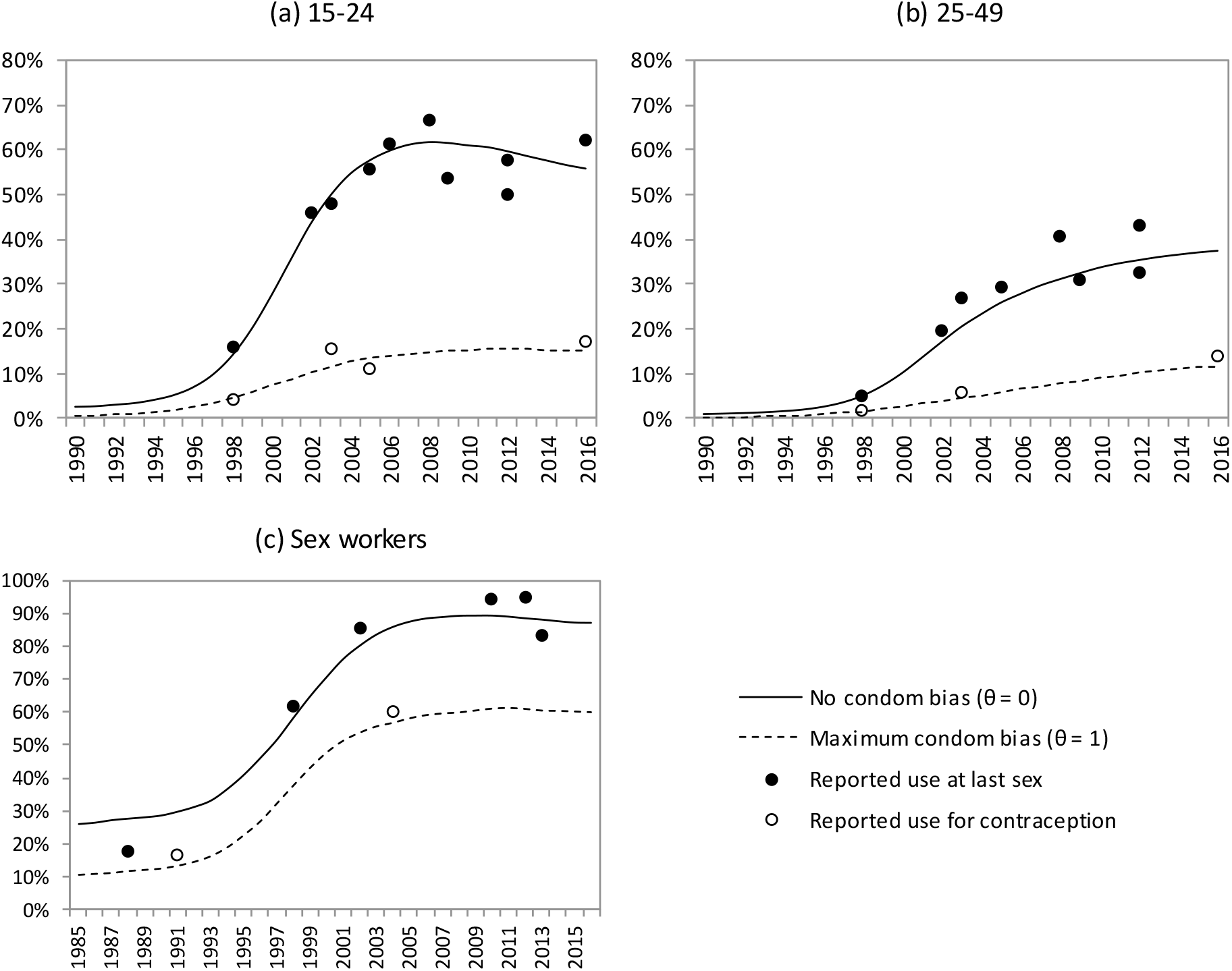
Calibration of model of condom use to reported condom use data. Model results are the average from 100 simulations. Modelled estimates of condom usage are calculated as the fraction of women who are consistently using condoms with their primary partners.

For the purpose of this analysis, the *θ* parameter has been set to 0.80, consistent with results of a previous analysis that defined the *θ* parameter similarly [80]. The median parameter *M*_1_ is calculated by noting that the ‘baseline’ parameters relate to 1998, and hence when *t* = 13 (i.e. in 1998)

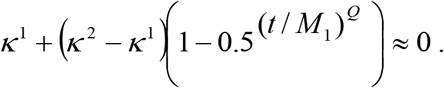

(The approximation is valid because the value of the third term in the *ς*(*t*) equation is close tc zero when *t* = 13.) The parameter *M*_1_ is therefore calculated as a function of *κ*^1^, *κ*^2^ and *Q*.

To ensure that male and female assumptions are consistent, the average probability that a heterosexual man uses a condom in a short-term relationship is calculated as

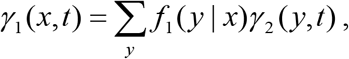

where *f*_1_(*y | x*) is the probability that a female partner is aged *y*, if the male partner is aged *x*.

To illustrate how these variables are applied in the model, consider the example of a new partnership formed at time *t*, between a woman aged *x* and a man aged *y*, with condom preference parameters of 0.87 and 1.33 respectively. The probability that they use a condom at their first sexual encounter will be

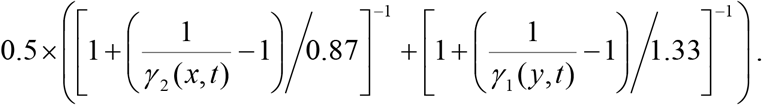

A random number between 0 and 1 is compared to this probability; if the random number is less than the probability, they are assumed to use a condom. If they use a condom at their first sexual encounter, it is assumed that they will continue to use condoms consistently, at least to the time when they start cohabiting. However, if they do not initiate condom use at first sex, it is assumed that they will never initiate condom use, unless one of the partners discloses that they are HIV-positive (discussed in section 5.2.2). This probability of adopting condoms following disclosure is calculated by comparing the rate of condom use if neither partner is diagnosed and the rate that would be expected if the HIV-positive partner had disclosed their HIV status to a partner whose HIV status was either negative or unknown. Extending the previous example, suppose that the male partner discovered he was HIV-positive and disclosed his HIV-status to the female partner (whose HIV status was unknown). If the 1.33 represents his condom preference *before* having learned his HIV status, then the probability of condom use prior to disclosure is calculated in the same way as before:

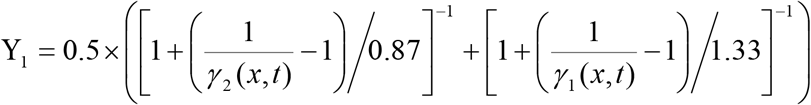

The probability of condom use if HIV disclosure had occurred at the start of the relationship is

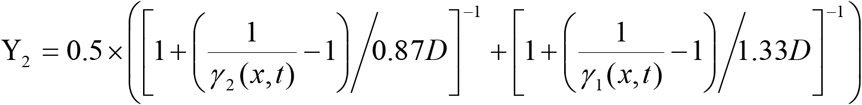

where *D* is the increase in the odds of condom use associated with HIV diagnosis and disclosure (set to 3, as explained in section 5.2.2). Given that condom use was not initiated at the start of the relationship, the probability that condom use is adopted following disclosure of HIV status is

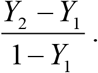

However, this formula is only applied in the event that the partner’s HIV status is negative or unknown (undiagnosed HIV). If disclosure occurs but the partner already knows they are HIV-positive, there is assumed to be no change in condom use.

#### 4.9.2 Condom use in long-term relationships

If the couple uses condoms consistently, in the initial phase of their relationship, it is assumed that there is a probability that they discontinue condom use at the point that their relationship becomes cohabiting, provided that neither partner has disclosed that they are HIV-positive. This probability, 0.46, is calculated based on logistic regression models fitted to data on condom usage in the 1998 and 2003 South African DHSs [30, 36], which compare condom usage in cohabiting and non-cohabiting women after controlling for age. If a randomly-generated uniform variate is less than 0.46, or if either partner has disclosed that they are HIV-positive, it is assumed that condoms continue to be used consistently after the relationship becomes cohabiting, otherwise condom use is discontinued. As before, condom use is only resumed if one partner is diagnosed positive and they disclose their HIV status to their partner.

#### 4.9.3 Condom use by sex workers and their clients

A similar approach to that described above is adopted in modelling condom usage by sex workers and their clients. The key difference is that the rates of condom use are assumed to be independent of age (i.e. the v term in equation (4.1) is omitted). The parameters in the high bias and no bias scenarios are summarized in Table 4.9.2. These assumptions have been set in order to ensure model consistency with levels of condom use reported by sex workers in surveys (Figure 4.9.1, panel c). The base probabilities of condom use among sex workers in 1998 (0.84 and 0.57 in the no bias and high bias scenarios respectively) are higher than those seen in the figure in 1998 because the effect of the allowance for heterogeneity in condom preference between sex workers and their clients is to lower the average level of condom use.

**Table 4.9.2:** Differences in condom usage parameters between scenarios

| Parameter | Symbol | No bias scenario<br>($\theta = 0$) | High bias scenario<br>($\theta = 1$) |
| --- | --- | --- | --- |
| Probability of condom use by sex workers in 1998 | $\gamma^*$ | 0.84 | 0.57 |
| Ultimate odds of condom use,<br>relative to the odds of condom use in 1998 | $\exp(\kappa^2)$ | 24 | 3.0 |
| Shape parameter controlling the speed of behaviour change | $Q$ | 4.0 | 4.0 |

#### 4.9.4 Condom use by MSM

Levels of condom use in same-sex relationships generally appear similar to those in heterosexual relationships [154, 159, 221]. It is therefore assumed that the rates of condom use for men in regular partnerships (short-or long-term) with other men are the same as the corresponding rates in heterosexual relationships.

There is evidence to suggest that MSM engaging in casual sex use condoms more consistently than MSM in regular partnerships [241]. Arnold *et al* [153] found, in a study of MSM in Soweto, that the number of acts of unprotected anal intercourse increased by a factor of 1.81 (95% CI: 1.43-2.28) if the partner was a regular partner (after controlling for number of anal sex episodes and other factors). The fraction of anal sex episodes that were unprotected was 0.24 in the average relationship, and 284 out of 758 male-male relationships were reported to be regular. If 73 is the probability of condom use in casual relationships, then

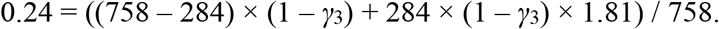

Solving for *γ*_3_ gives 0.816, and an associated probability of condom use of 0.667 in regular relationships. This implies an odds ratio of 2.21 for the increase in the odds of condom use in casual sex relative to regular relationships. We therefore assume that the odds of condom use in a casual MSM relationship is 2.2 times that in a short-term MSM relationships (on the assumption that almost all of the ‘regular’ relationships included in the Arnold study were short-term partnerships).

#### 4.9.5 Effect of educational attainment on condom usage

Evidence from sub-Saharan Africa consistently shows a strong association between educational attainment and condom use [164]. In South Africa, several studies show an association between education and condom use. For example, in a trial involving youth in rural South Africa, Hargreaves *et al* [238] found that the odds of condom use at last sex increased by factors of 1.26 (95% CI: 0.86-1.86) and 1.97 (95% CI: 1.28-3.03) in youth with some secondary and completed secondary education respectively, when compared to youth with no secondary education. Maharaj and Cleland [242] found that among married/cohabiting couples in KwaZulu-Natal, condom use was strongly associated with both partners’ educational attainments, and similar results were found in a study of youth in KwaZulu-Natal [50]. In an analysis of the South African 1998 DHS data, Katz [243] found that condom use at last sex was significantly associated with having completed secondary education. However, some South African studies have shown that the strong association between education and condom use observed in univariate analyses ceases to be significant after controlling for factors such as age (which is strongly associated with education) and knowledge about HIV and condoms (which is likely to lie on the causal pathway between education and condom use) [244].

In our model, we have relied on levels of condom use reported by women in non-cohabiting relationships in the 1998 and 2003 DHSs (summarized in Table 4.9.3) [30, 36]. Average odds ratios have been calculated, relative to the grade 8-11 category (summarized in the last row). Taking baseline as grade 10, we set the odds ratios for condom use in the model to 0.22 for no education, 0.36 for grade 3, 1.42 for grade 12 and 2.38 for tertiary education. The odds ratios for all other grades are linearly interpolated between these values (with a value of 0.59 applying at the mid-point between grades 6 and 7). However, when modelling uptake of condom use using these parameters, it was found that women who had completed high school had levels of condom use that were too low, relative to the levels reported in the 1998 and 2003 DHSs. The odds ratio for the effect of grade 12 was therefore increased from 1.42 to 1.60 and the odds ratio for the effect of tertiary education was therefore increased from 2.38 to 5.0, in order to achieve greater consistency between the model estimates and survey estimates (see Figure 4.9.2).

**Table 4.9.3:** Fraction of women reporting condom use at last sex (in non-cohabiting relationships), by highest educational attainment

|  | Highest educational attainment |  |  |  |  |  |
| --- | --- | --- | --- | --- | --- | --- |
|  | No education | Grades 1-5 | Grades 6-7 | Grades 8-11 | Grade 12 | Tertiary education |
| 1998 data | 3.6% | 5.3% | 11.9% | 17.3% | 20.2% | 35.1% |
| 2003 data | 16.4% | 26.1% | 29.5% | 43.6% | 55.8% | 62.8% |
| Odds ratio |  |  |  |  |  |  |
| 1998 | 0.18 | 0.27 | 0.65 | 1.00 | 1.21 | 2.59 |
| 2003 | 0.25 | 0.46 | 0.54 | 1.00 | 1.63 | 2.18 |
| Average | 0.22 | 0.36 | 0.59 | 1.00 | 1.42 | 2.38 |

**Figure 4.9.2:**
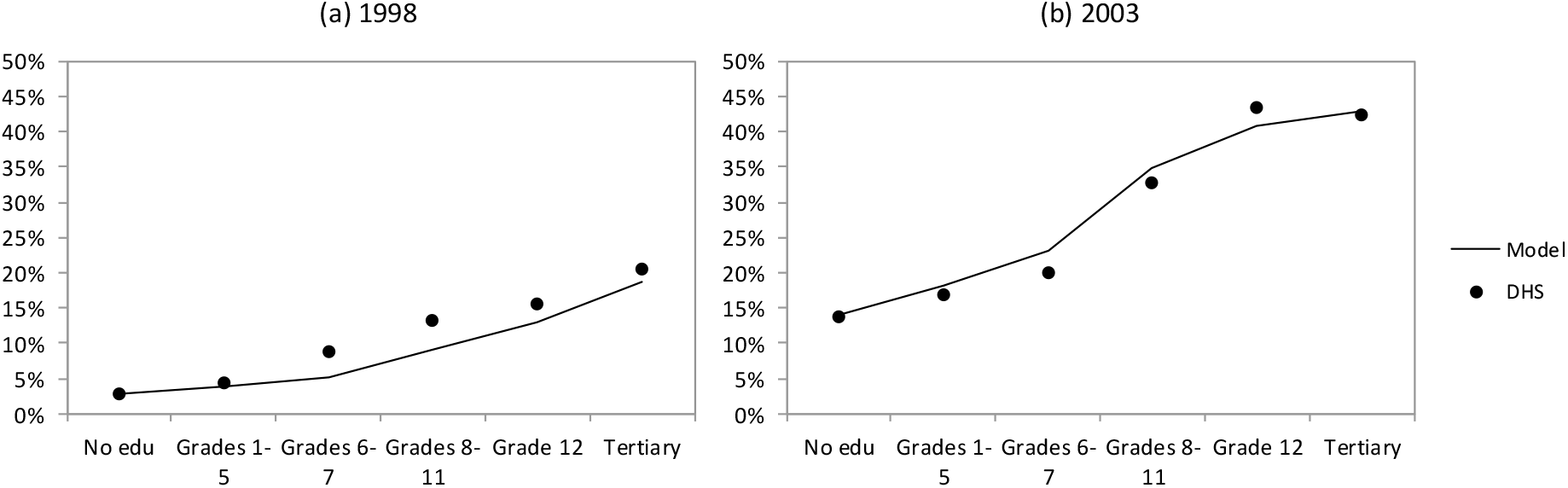
Female condom use at last sex, by educational attainment. Model results are the average from 100 simulations. These are the results obtained when there is assumed to be no bias in the reporting of condom use (i.e. ***θ*** = 0). Modelled estimates of condom usage are calculated as the fraction of women who are consistently using condoms with their primary partners.

#### 4.9.6 Effect of race on condom usage

Several studies have shown that levels of condom usage differ between South African race groups. Table 4.9.4 summarizes the evidence from various studies. In general, levels of condom usage appear to be lower among white South Africans than among black South Africans (average OR of 0.58), and lowest among coloured South Africans (average OR of 0.42). This could potentially reflect the greater burden of HIV in the black South African population and a greater behaviour change among black South Africans in response to the HIV epidemic. Early studies conducted in the late 1980s and early 1990s, prior to the introduction of HIV communication programmes, found relatively low rates of condom use among black South Africans when compared with other race groups [33, 195], and thus it seems plausible that the relatively high levels of condom usage among black South Africans in recent surveys are the result of greater response to AIDS awareness programmes. However, it is important to bear in mind that most of these odds ratios are univariate and thus do not control for many of the confounding factors noted previously; this means that there could be other explanations for the high levels of condom usage observed in black South Africans (e.g. a lower proportion married or a younger age distribution).

**Table 4.9.4:**
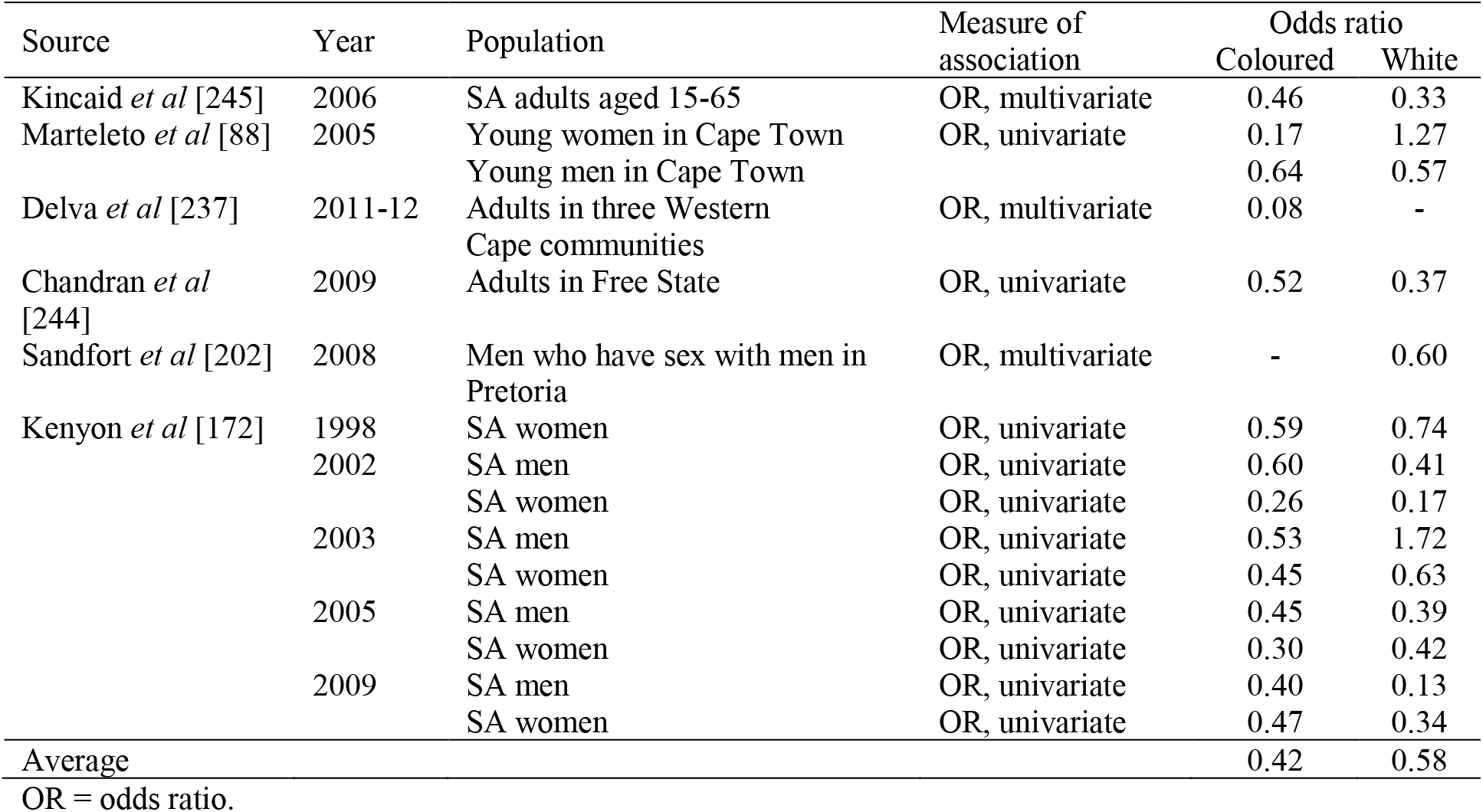
Odds of condom usage in coloured and white South Africans (relative to black South Africans)

In the model, we set the *R*(*r_i_*) variables in such a way that the modelled odds of condom use in coloured and white South Africans (relative to those in black South Africans) are consistent with the average odds ratios in Table 4.9.4. This means setting *R*(*r_i_*) to 0.35 for coloured South Africans and 1.50 for white South Africans. The ratio of 1.50 is very different from the univariate association in Table 4.9.4 (0.58) because the latter is confounded by differences in marriage rates (rates of marriage being substantially higher among white South Africans than among black South Africans).

#### 4.9.7 Model calibration

Figure 4.9.2 shows the simulated fractions of women who report having used a condom the last time they had sex compared with the actual levels measured in the 1998 and 2003 DHSs. The model results match the observed association between condom use and educational attainment quite closely, although the model estimates are slightly too low for women with incomplete secondary education in 1998.

### 4.10 Commercial sex

For a sexually experienced man with ID *u*, the annual rate of contact with female sex workers, *w*(*u*), is assumed to depend on his age (*x_u_*), race (*r_u_*), urban/rural location (*b_u_*), same-sex preference (*Y_u_*), risk group (*i_u_*), marital status (*l_u_*) and number of current partners (*j_u_*). As in section 4.3, a gamma probability density function is used to represent the age differences in rates of male contact with sex workers; for the purpose of calculating a constant rate over each five-year age band, *x_u_* is taken to be the midpoint of the age group (e.g. 17.5 for men aged 15 to 19). The following formula is used to calculate *w*(*u*):

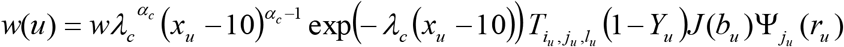

where *λ_c_* and *α_c_* are the parameters of the gamma probability density function, *T_i,j,l_* is an adjustment factor to represent the man’s risk group and relationship status, *J*(*b_u_*) is an adjustment factor to represent the effect of urban/rural location, and Ψ_*j*_(*r_u_*) is the same adjustment factor described in section 4.3 (i.e. allowing for racial differences in the incidence of concurrency). The assumed values of the parameters are summarized in Table 4.10.1. Men are assumed not to have contact with sex workers if they are currently in prison or are virgins. The base rate of sex worker contact (*w*, which applies to high risk men who currently have no partner and are aged 20-24) has been set in such a way that the demand for commercial sex is sufficient to match the estimated size of the South African sex worker population in a recent study [246], when it is assumed that sex workers have 750 clients per annum on average [204, 247–253]. Using this sex worker population size estimate and the assumed number of 750 clients per annum implies roughly 5 sex worker contacts per annum for men age 15-49. Hence the *w* parameter has been calculated so that the average number of sex worker contacts per year, averaged across all men aged 15 to 49 at the start of the simulation, is 5, i.e.

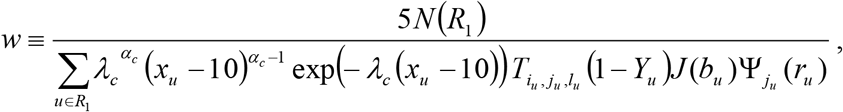

where *R*_1_ is the set of all men aged 15-49 who are alive, and *N*(*R*_1_) represents the number of men in this set.

**Table 4.10.1:**
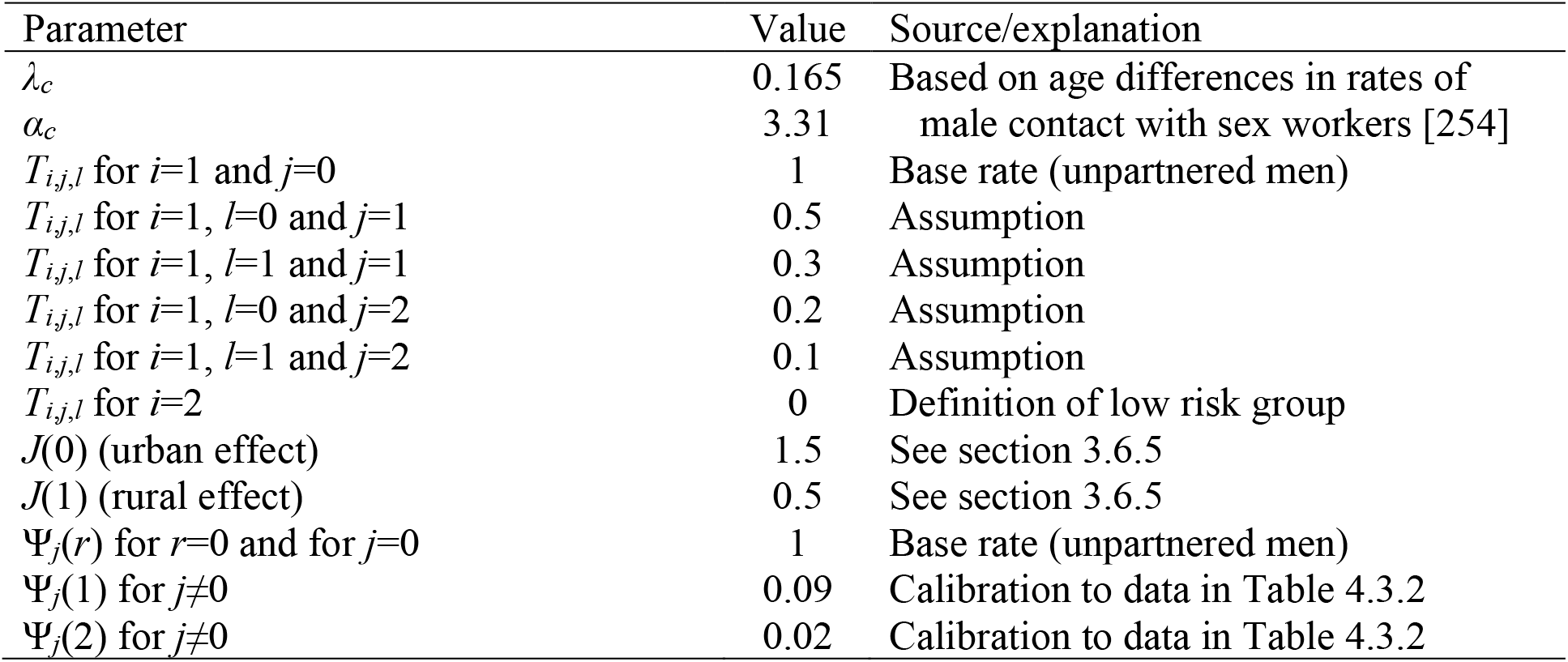
Assumed male rates of sex worker contact

Since only men in the high risk group are assumed to have contact with sex workers, and since all sex workers are assumed to be recruited from the high risk group, no assumptions about mixing between risk groups are required for the purpose of modelling commercial sex. It is assumed in the interests of simplicity that clients have no preference regarding the age of their commercial sex contacts, although age preferences are accounted for implicitly by assuming a sex worker age distribution (based on data from Johannesburg [255]) and setting the age-specific rates of entry into sex work in such a way that the simulated age distribution remains roughly stable over time and matches that assumed. Women are assumed to remain active as sex workers for two years on average, before returning to the unpartnered state.

The number of new sex workers required over the period [*t,t* + *d*) in order to satisfy male demand, Δ_*c*_(*t,t* + *d*), is calculated as

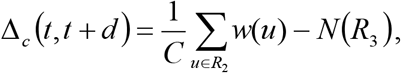

where *C* is the assumed annual number of clients per sex worker (750), *R*_2_ is the set of all men who are sexually experienced at time *t*, and *R*_3_ is the set of women engaged in sex work at time *t*. The probability that a woman in the high risk group, who has no partners and is aged *x* and in HIV disease state *s* at time *t*, becomes a sex worker over the period [*t,t + d*) is calculated as

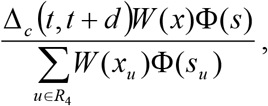

where *W*(*x*) is the factor by which the rate of recruitment into the ‘sex worker’ group is multiplied when the woman is of age *x* in order to match the target sex worker age profile, Φ(*s*) is the factor by which the rate of recruitment into commercial sex is adjusted in HIV disease stage *s* (these are the same as the factors assumed in modelling formation of short-term partnerships – see Table 4.3.1) and *R*_4_ is the set of unpartnered women in the high risk group at time *t*.

As noted previously, the model does not consider male sex workers.

## 5. HIV interventions

This chapter describes the HIV interventions that have been introduced in South Africa to date. The condom promotion and HIV communication programmes that were introduced in the mid-1990s, and their effect on condom usage, have already been discussed in section 4.7 and are not repeated here. Prevention of mother-to-child transmission programmes are also allowed for in the model, but are not described here, as the focus of this report is limited to sexual transmission of HIV.

### 5.1 Male circumcision

The model assumes that rates of male circumcision prior to 2008 depended only on age and race. Following 2008, when campaigns were introduced to promote medical male circumcision (MMC) as an HIV prevention strategy, the model assumes that rates of male circumcision changed over time, with the extent of the increase in male circumcision uptake again being age-dependent. The sections below describe the modelling of male circumcision uptake over each of these two periods.

#### 5.1.1 Male circumcision prior to 2008

The approach to modelling of male circumcision prior to 2008 is similar to that in the Thembisa model [256], with the main difference being that the approach has been extended to allow for differences in rates of male circumcision between population groups. We assume that the prevalence of male circumcision at age *x*, in population group *r*, is determined by the function

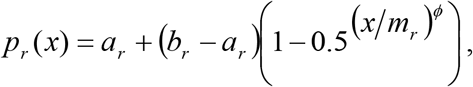

where *a_r_* is the proportion of males of race *r* who are circumcised soon after birth, *b_r_* is the level at which circumcision prevalence ‘stabilizes’ in older men, *m_r_* is the median age at circumcision in men who get circumcised after birth, and *ϕ* is the ‘shape’ parameter that determines the concentration of the distribution of circumcision ages (post-birth) around the median. The assumed values of these parameters are shown in Table 5.1.1; these have been set in such a way that the model matches the fraction of men who report being circumcised and the distribution of reported ages at circumcision, in the 2002 HSRC household survey [257]. The shape parameter *ϕ* has been set to 4.5, the same values as assumed previously when the Thembisa model was fitted to age-specific circumcision prevalence data [256]. The parameter estimation takes into account likely misreporting of male circumcision status, as described previously [256].

**Table 5.1.1:** Male circumcision parameters

|  | Black | Coloured | White |
| --- | --- | --- | --- |
| Fraction circumcised at birth ($a_r$) | 0.068 | 0.143 | 0.223 |
| Fraction ultimately circumcised ($b_r$) | 0.450 | 0.280 | 0.330 |
| Median age at circumcision, after birth ($m_r$) | 21.2 | 20.0 | 18.0 |

The calculated values of *p_r_*(*x*) are used to randomly assign a circumcision status to each male at the start of the simulation (in 1985). Similarly, whenever a male child is added to the simulated population, he is randomly assigned a circumcision status at birth (with the probability of circumcision at birth being just *a_r_*). For every uncircumcised male in the population, the male circumcision status is updated at annual intervals, with the probability of becoming circumcised – for a man currently aged *x* and of race *r* – being calculated as

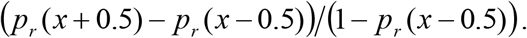

#### 5.1.2 Male circumcision in 2008 and subsequent years

In 2008 and subsequent years, the above formula is modified to take into account additional demand for MMC, which is assumed to be dependent on the individual’s age. For an individual of age *x*, who is HIV-negative, the probability of becoming circumcised changes to

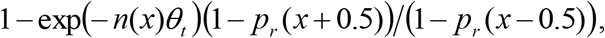

where *θ_t_* is the annual rate of MMC as part of MMC campaigns in year *t*, for a male aged 10-14, and *n*(*x*) is the relative rate of MMC uptake at age *x* (compared to the 10-14 age category). The *θ_t_* factor is estimated from *R*(*t*), the ratio of total MMC operations in year *t* to the total number of South African men aged 15-49 at the middle of year *t* (shown in Table 5.1.2). The *n*(*x*) factors, shown in Table 5.1.3, are estimated based on the age distribution of men seeking MMC in South Africa in recent years [258], and are similar to the age adjustment factors assumed in the Thembisa model [259]. If *N*(*x, t*) is the number of uncircumcised, HIV-negative men at age *x* in year *t*, and *M*(*t*) is the number of men aged 15-49 in year *t*, then the *θ_t_* parameter is calculated as

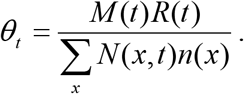

**Table 5.1.2:** Ratio of MMC operations to male population aged 15-49

|  | 2008 | 2009 | 2010 | 2011 | 2012 | 2013 | 2014 |
| --- | --- | --- | --- | --- | --- | --- | --- |
| MMC operations* | 5190 | 9168 | 131117 | 347973 | 422262 | 331668 | 508404 |
| Men aged 15-49 (millions)† | 13.06 | 13.19 | 13.34 | 13.49 | 13.67 | 13.84 | 14.00 |
| Ratio ($R(t)$) | 0.0004 | 0.0007 | 0.0098 | 0.0258 | 0.0309 | 0.024 | 0.0363 |

\* Source: [260-263]. † From the ASSA2008 AIDS and Demographic model [22].

**Table 5.1.3:** Relative rates of MMC operation by age

| 10-14 | 15-19 | 20-24 | 25-29 | 30-34 | 35-39 | 40-44 | 45-49 | 50-54 | 55-59 | 60-64 | 65-69 | 70+ |
| --- | --- | --- | --- | --- | --- | --- | --- | --- | --- | --- | --- | --- |
| 1.00 | 0.60 | 0.35 | 0.27 | 0.21 | 0.16 | 0.12 | 0.08 | 0.05 | 0.03 | 0.02 | 0.01 | 0.00 |

It is assumed that men who are HIV-positive do not get circumcised through MMC campaigns, as these campaigns encourage HIV testing prior to male circumcision, and there would be little motivation to get circumcised if the individual was already HIV-positive. (HIV-positive men may nevertheless want to get circumcised for cultural reasons or reasons unrelated to the HIV prevention benefit, so the rate at which they become circumcised is assumed to be the same as the rates of male circumcision in the general population prior to 2008.)

For the years that follow 2014, we assume that the same annual ratio of MMC operations to men aged 15-49 (0.0363) applies when calculating the number of men who get circumcised through MMC campaigns.

#### 5.1.3 Effect of male circumcision on HIV transmission probabilities

Men who are circumcised are assumed to have a 60% lower rate of HIV acquisition from HIV-positive female partners, per act of sex [264]. However, male circumcision is assumed to have no effect on male-to-female transmission of HIV [265] or male-to-male transmission of HIV (as there is only limited evidence of a protective effect in MSM [233]).

### 5.2 HIV counselling and testing

#### 5.2.1 Modelling HIV diagnosis

The modelling of the uptake of HIV counselling and testing (HCT) is an extension of that described in the Thembisa model [266]. Individuals are classified according to their HIV testing history: never tested, previously tested negative, or diagnosed positive. Eight modes of HIV testing are simulated, each described in more detail below.

##### 5.2.1.1 General HIV testing

General HIV testing refers mainly to self-initiated HIV testing and all other HIV testing not included in the categories outlined below. The approach to modelling general HIV testing is similar to that described in the Thembisa model [266], which is to say that the rate of general HIV testing is assumed to depend on the calendar year and the individual’s age, sex, sexual experience, educational attainment and HIV testing history. Base HIV testing rates are specified for each sex and each year (Table 5.2.1). These base HIV testing rates are set in such a way that the model matches the total recorded numbers of HIV tests performed in South Africa in each year (see Figure 5.2.1(a)).

**Figure 5.2.1:**
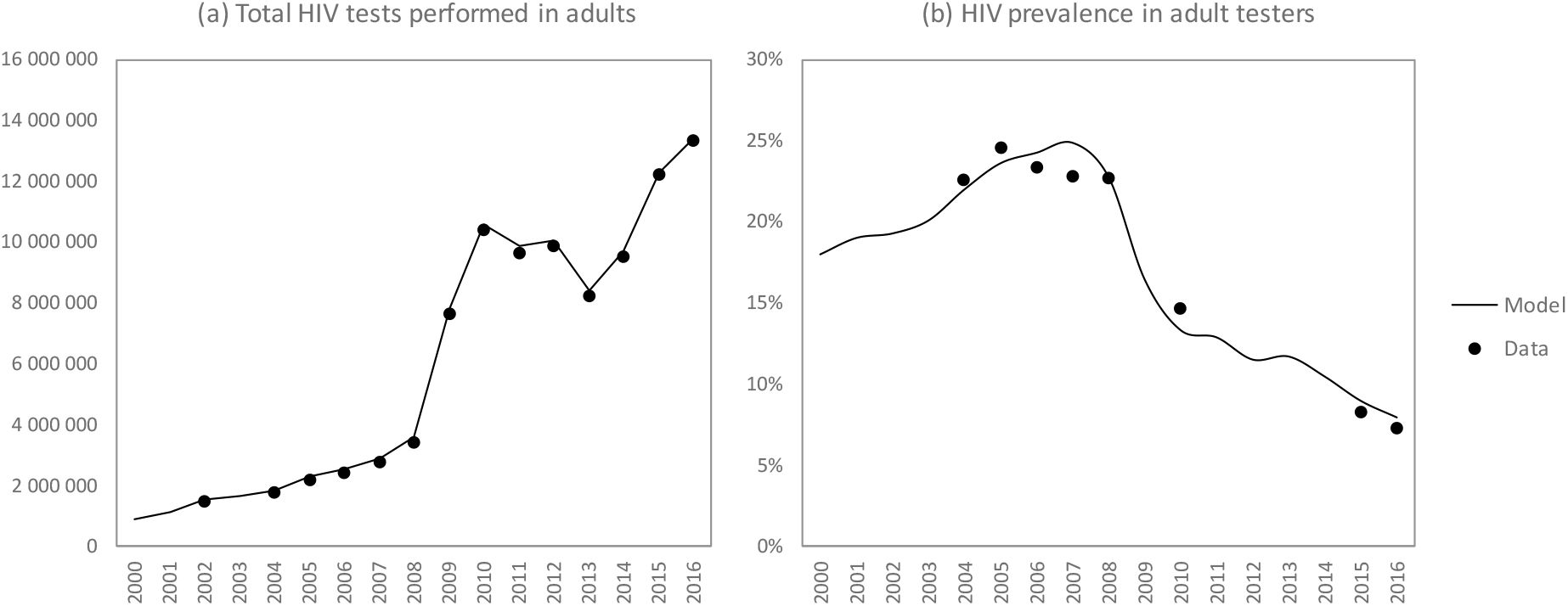
Total HIV tests performed in South African adults and HIV prevalence in adults tested for HIV

**Table 5.2.1:** Assumed HIV testing rates in OI patients and pregnant women (undiagnosed)

| Year | Base general testing rate |  | Testing of OI patients |  | Testing of pregnant women |  | Retesting of HIV-negative pregnant women |  |
| --- | --- | --- | --- | --- | --- | --- | --- | --- |
|  | Male | Female | Rate | Sources | Rate | Sources | Rate | Sources |
| Pre-1999 | - | - | 5% |  | 0.0% |  | 0% |  |
| 1999-00 | 0.0347 | 0.0558 | 5% |  | 0.9% |  | 0% |  |
| 2000-01 | 0.0365 | 0.0586 | 5% |  | 2.9% |  | 0% |  |
| 2001-02 | 0.0374 | 0.0602 | 5% |  | 7.5% | [281] | 0% |  |
| 2002-03 | 0.039 | 0.0626 | 5% |  | 15.6% | [282, 283] | 0% |  |
| 2003-04 | 0.0308 | 0.0506 | 5% |  | 31.3% |  | 0% |  |
| 2004-05 | 0.0228 | 0.0384 | 8% | [284] | 42.0% | [285] | 0% |  |
| 2005-06 | 0.0263 | 0.0453 | 15%* | [286] | 54.5% | [287] | 0% |  |
| 2006-07 | 0.0265 | 0.0467 | 21%* | [286] | 72.2% | [288] | 0% |  |
| 2007-08 | 0.0241 | 0.0435 | 29%* | [286, 289] | 84.0% | [290] | 5% | [291] |
| 2008-09 | 0.0382 | 0.0707 | 34%* | [286, 289] | 89.0% | [292] | 15% |  |
| 2009-10 | 0.1477 | 0.2803 | 43%* | [286, 293] | 93.0% |  | 25% | [294] |
| 2010-11 | 0.1982 | 0.3861 | 55%* | [293, 295] | 97.0% | [296] | 35% | [294] |
| 2011-12 | 0.1550 | 0.3020 | 78%* | [293] | 98.0% | [297] | 45% | [294] |
| 2012-13 | 0.1472 | 0.2867 | 83%* | [293] | 98.0% |  | 55% |  |
| 2013-14 | 0.1043 | 0.2031 | 87%* | [293] | 98.0% |  | 65% |  |
| 2014-15 | 0.1199 | 0.2336 | 92%* | [293] | 98.0% |  | 75% |  |
| 2015-16 | 0.1677 | 0.3268 | 93%* | [293] | 98.0% |  | 85% |  |
| 2016-17 | 0.1851<sup>‡</sup> | 0.3605<sup>‡</sup> | 93%*<sup>†</sup> |  | 98.0%<sup>†</sup> |  | 95%<sup>†</sup> |  |

\* Assumed rates are slightly lower than those quoted in the cited sources because the sources present the fraction of TB patients with a known HIV status, which is higher than the fraction of TB patients who receive an HIV test. † Rates are assumed to remain constant at the 2016/17 level after 2017. ‡ The annual testing rate is set to 0.1448 and 0.2821 in men and women respectively (the average over the 2012-17 period), for all years after 2017.

The base HIV testing rates are assumed to apply at age 25. Multiplicative age adjustment factors are applied to these base rates in calculating the rates that apply at other ages. A gamma probability density function is used to represent the relative rate of HIV testing at different ages; in males, the mean and standard deviation of the gamma distribution are set at 37.2 and 28.0 respectively, while in women the mean and standard deviation are set at 22.3 and 19.4 respectively, the values estimated when the Thembisa model was fitted to age- and sex-specific HIV testing data in South Africa [266].

The base HIV testing rates are assumed to apply in individuals whose highest educational attainment is grade 11. For each additional grade completed, the rate at which individuals seek general HIV testing is assumed to increase by a factor of 1.12. (For example, the rate of HIV testing in someone who has completed grade 8 would be 0.71 (1.12^8–11^) times the base rate that applies in an individual who has completed grade 11.) The factor of 1.12 is the estimated effect of educational attainment on the uptake of HIV testing by black South Africans between 2010 and 2012, after controlling for age, sex and other factors, which was estimated using data from nationally-representative surveys [267].

General HIV testing is assumed not to occur in youth who are not yet sexually experienced, but apart from this, no effect of sexual risk behaviour on the uptake of general HIV testing is assumed. These assumptions are consistent with South African studies. For example, in a recent study of youth in Western Cape and Mpumalanga, those who reported being sexually experienced were 4.7 times (95% CI: 3.1-7.1) as likely to report having tested for HIV as those who were virgins, after controlling for age and other factors (Franziska Meinck, personal communication). Similarly, in a household survey conducted by Venkatesh *et al* [268], virgins were significantly less likely to report previous HIV testing than youth who were sexually experienced, after controlling for age and other factors. However, among youth who were sexually experienced, the lifetime number of sexual partners was not a significant determinant of the individual’s previous testing, either in men or women. Most other South African studies have also not found significant associations between HIV testing history and numbers of partners [269–271].

The base rates apply to individuals who have never tested for HIV before. Individuals who have previously been tested for HIV (and tested negative) are assumed to have a higher rate of HIV testing than individuals who have never previously sought testing, with a multiple of 1.97 being applied to the base rates. This multiple is the same as that estimated when the Thembisa model was fitted to South African HIV testing data [266], and is consistent with various South African studies that show higher rates of testing uptake in previously-tested individuals compared to previously-untested individuals [272–274]. It is assumed that individuals who have previously been diagnosed positive might also get retested, either because they seek confirmation of the positive test result or because they need to test again in order to link to HIV care. This rate of retesting is assumed to be 0.50 times the rate of testing that applies in individuals who have never previously tested for HIV, if the individual is not on ART. If the individual is on ART, their rate of testing is assumed to be 0.15 times the rate that applies in individuals who have never been tested. These parameter were chosen such that the model matched the HIV prevalence trends in individuals tested for HIV in South Africa (see Figure 5.2.1(b)).

##### 5.2.1.2 Testing of patients with HIV-related opportunistic infections

The CD4 counts of HIV-positive adults are dynamically updated (see section 6.2), and the incidence of HIV-related opportunistic infections (OIs) is assumed to be a function of the individual’s CD4 count and receipt of ART. The annual incidence of OIs in untreated HIV-positive adults is assumed to be 0.05 per annum during acute HIV infection, 0.08 per annum after acute infection in individuals with CD4 counts of 350 cells/μl or higher, 0.27 per annum in individuals with CD4 counts of 200-349 cells/μl and 0.90 per annum in individuals with CD4 counts <200 cells/μl. These rates are based on studies of the incidence of OIs in HIV-positive adults in sub-Saharan African settings [275, 276], and are the same as the values assumed in the Thembisa model [266]. The incidence of HIV-related OIs in HIV-negative adults has been set to 0.019 per annum; this assumption is derived by dividing the annual incidence of TB in the HIV-negative South African population (0.004 per annum [277]) by the fraction of OI patients that have TB at high CD4 counts (20.8% in a Cape Town study [275]). The incidence of OIs in ART patients is assumed to be 0.10 per annum, which is 0.11 times the rate in untreated patients with CD4 <200 and 0.37 times the rate in untreated patients with CD4 200-349. These ratios are roughly consistent with studies that have shown the incidence of HIV-related morbidity after ART initiation to be between 0.16 and 0.39 times that in untreated patients, after controlling for differences in baseline CD4 count [278–280].

Few data exist on the proportion of OI patients who are tested for HIV, except in the case of TB. It is therefore assumed that the fraction of OI patients tested for HIV is the same as the rate of HIV testing in patients with TB symptoms. Table 5.2.1 summarizes the model assumptions about the fraction of OI patients tested for HIV and the associated data sources. In individuals who have previously been diagnosed HIV-positive, it is assumed that the probability of being HIV-tested again if they experience an OI is 50% of the probability that applies to patients who have not previously been HIV-diagnosed (or 15% of that probability if the individual is already on ART). These assumptions are the same as apply to general HIV testing.

##### 5.2.1.3 Testing of pregnant women

The modelling of fertility is described in more detail in section 3.4. Table 5.2.1 shows the assumed proportions of pregnant women tested for HIV in each year. These proportions apply to all women who have not previously been diagnosed positive. In women who have previously been diagnosed positive, the probability of retesting if they fall pregnant is assumed to be 50% of the probability that would apply if they had not previously been diagnosed (or 15% of that probability if they are already on ART). This is the same adjustment that applies in the case of general HIV testing.

In recent years it has become common for women who test negative on their first antenatal HIV test to be offered retesting in late pregnancy. Although these retests are usually not included in the total numbers of HIV tests reported by the Department of Health, and are not included in the denominator when reporting the HIV prevalence among women tested for HIV antenatally, they do nevertheless affect the cost-effectiveness of antenatal HIV screening and are therefore important to model. Table 5.2.1 shows the assumed fraction of women testing HIV-negative antenatally who receive retesting in late pregnancy. The proportion is assumed to increase to 95% in 2016 and all subsequent years, based on recent unpublished DHIS data, which suggest that almost all HIV-negative pregnant women receive retesting in late pregnancy.

##### 5.2.1.4 Testing of partners of diagnosed individuals

Currently in South Africa, there is no policy of active partner notification when an HIV diagnosis is made. Although newly-diagnosed individuals may be encouraged by health workers to disclose their HIV status to their sexual partner(s), the health worker makes no attempt to contact these sexual partners on behalf of the patient. This ‘passive’ approach to partner notification is generally not shown to be very effective in getting partners of newly-diagnosed individuals tested for HIV. As discussed in section 5.2.2.1, HIV-diagnosed individuals often do not disclose their HIV status to their sexual partners, with disclosure being particularly uncommon in the case of non-spousal relationships [298–300]; it has also been found that rates of disclosure are relatively low in women and in individuals who have not yet initiated ART [44, 299, 301, 302]. Even when individuals disclose their HIV status, there is no guarantee that this will result in their sexual partner seeking HIV testing.

Table 5.2.2 summarizes the results of African studies that have assessed the proportion of sexual partners who come for HIV testing following HIV diagnosis of the index patient. With the exception of the studies of Brown *et al* [303] and Kiene *et al* [304], all of these studies have been done in the context of antenatal screening, and there is thus limited African data on the success of partner notification when the index patient is male. The weighted average fraction of partners receiving HIV testing was 30% (after weighting by the number of index cases in each study). It is also important to note that all of these studies have been conducted in settings in which a relatively high proportion of index patients are married or cohabiting with their sexual partner. In two of the studies, there was a strong association between male partner testing and marital status (OR 4.50 (95% CI: 2.20-9.19) in Tanzania [305] and OR 2.11 (95% CI: 1.14-3.92) in Kenya [306]).

**Table 5.2.2:** Proportion of partners coming for HIV testing following diagnosis of index HIV case (passive partner notification approaches)

| Study | Location | % married | # individuals diagnosed | % of partners seeking testing |
| --- | --- | --- | --- | --- |
| Msuya *et al* [305] | Moshi, Tanzania | 91% | 184 | 18.5% |
| Aluisio *et al* [306] | Nairobi, Kenya | 84%* | 456 | 30.7% |
| Brown *et al* [303] | Lilongwe, Malawi | 73% | 82 | 24.4% |
| Rosenberg *et al* [307] | Lilongwe, Malawi | 100% | 100 | 52.0% |
| Kiene *et al* [304] | Mpigi, Uganda | - | 38<sup>†</sup> | 35.1% |

\* Monogamous marriages only. † The number of partners of newly-diagnosed individuals.

In our model, the probability of HIV disclosure at the time of HIV diagnosis is assumed to depend on the relationship type and the individual’s sex (see section 5.2.2). A disclosure probability of 50% is assumed for women who are in non-marital relationships, and this disclosure probability is adjusted by odds ratios of 1.25 in men and 2.5 in married couples (i.e. the probability of disclosure from a married woman to her husband is 71%, and the probability of disclosure from an unmarried man to his partner is 55%). If 30% of the husbands of newly-diagnosed married women receive HIV testing following their wives’ HIV diagnosis, this implies that the probability of men seeking HIV testing, given that their wives have disclosed their HIV-positive status, is 0.42 (i.e. 0.30/0.71). In the absence of more detailed data, the same 0.42 probability of seeking testing (conditional on disclosure) is assumed to apply regardless of whether the partner is male or female, and regardless of whether the relationship is marital or non-marital. For example, if an unmarried woman is diagnosed positive, the probability that she discloses her HIV status to her male partner and he gets tested as a result of this disclosure is 0.50 × 0.42 = 0.21.

##### 5.2.1.5 Testing of STI patients

Early syndromic management protocols in South Africa did not explicitly mention the need for HIV testing as part of STI patient consultations [308, 309], and the importance of HIV testing has only been mentioned in guidelines issued since 2009 [310]. Few data exist on the extent to which HIV testing is actually offered. The only nationally representative data are from surveys conducted in 2002 and 2014, in which standardized patient actors were used to estimate that 8% and 67% of providers respectively offered HIV testing to STI patients [282, 311]. A longitudinal study in North West province in 2013 found that 54% of standardized patient actors were offered HIV testing at baseline [312]. Earlier studies in Cape Town and KwaZulu-Natal found intermediate proportions: Leon *et al* conducted a trial and found that in non-intervention STI clinics, the rate of HIV testing increased from around 30% prior to 2007 to 43% in 2007 [313], and the latter was similar to the rate found in KwaZulu-Natal in 2006 [314]. Table 5.2.3 shows the assumed proportions of STI patients receiving HIV testing in each year, based on these studies. These rates of HIV testing are assumed to apply only in the public health sector, as the only available data are from the public sector. The limited data available from the South African private sector suggest that relatively few private practitioners follow syndromic management protocols [315], and thus few would be expected to offer HIV testing as part of the STI patient consultation.

**Table 5.2.3:** Assumed HIV testing rates in public-sector STI patients and prisoners (undiagnosed)

| Year | Testing of STI patients |  | Testing of prisoners |  |
| --- | --- | --- | --- | --- |
|  | Proportion | Sources | Annual rate | Sources |
| Pre-2000 | 0% |  | 0.0 |  |
| 2001-02 | 3% |  | 0.0 |  |
| 2002-03 | 8% | [282] | 0.0 |  |
| 2003-04 | 13% |  | 0.0 |  |
| 2004-05 | 18% |  | 0.0 |  |
| 2005-06 | 24% |  | 0.05 |  |
| 2006-07 | 30% | [313] | 0.10 |  |
| 2007-08 | 43% | [313] | 0.19 | [125] |
| 2008-09 | 47% |  | 0.29 |  |
| 2009-10 | 51% |  | 0.39 | [124] |
| 2010-11 | 55% |  | 0.49 |  |
| 2011-12 | 58% |  | 0.58 | [316] |
| 2012-13 | 61% |  | 0.68 | [316] |
| 2013-14 | 64% | [312] | 0.97 | [317] |
| 2014-15 | 67% | [311] | 1.13 | [121] |
| 2015-16 | 69%* |  | 1.13* |  |

\* Rates are assumed to remain constant at the 2015/16 level after 2016.

Our model simulates the incidence of five STIs in addition to HIV: syphilis, genital herpes, gonorrhoea, chlamydia and trichomoniasis. In addition, the model simulates the incidence of vaginal candidiasis and bacterial vaginosis in women. For each of these infections, an assumed proportion of cases becomes symptomatic, with individuals experiencing symptoms of genital ulceration or discharge. Symptomatic individuals are assumed to seek treatment at a weekly rate, with a proportion of these cases seeking treatment in public health facilities. Assumptions regarding STI treatment seeking are described in more detail elsewhere [318], and the model calibration to South African STI prevalence data is also described elsewhere [18]. The model simulates HIV testing in STI patients by applying the testing probabilities in Table 5.2.3 to the weekly rates of seeking treatment in public health facilities, in individuals who have any symptoms of genital ulceration or discharge. A limitation of this modelling approach is that it does not consider other conditions that could prompt patients to seek treatment at STI clinics (for example, human papillomavirus infections, which cause genital warts). The model assumes that individuals who have previously been diagnosed positive are only 50% as likely to be tested for HIV on developing STI symptoms as other individuals with STI symptoms (or 15% as likely if they are on ART).

##### 5.2.1.6 Testing of men who seek medical male circumcision (MMC)

It is standard practice for men who get medically circumcised to receive HIV testing prior to circumcision. WHO guidelines on MMC recommend HIV testing prior to circumcision [319], and most donor-supported programmes also recommend this. For example, Lesotho’s MMC programme, supported by USAID, has provided HIV testing to 97% of all men presenting for MMC [320], and in a South African study, almost all men seeking MMC received HIV testing [321].

Assumptions about the annual rate of MMC uptake in uncircumcised men are described in section 5.2.1. It is assumed that only men who have not been diagnosed positive would seek MMC, and that all men who seek MMC are tested for HIV. In the event that men are diagnosed positive for the first time on seeking MMC, it is assumed that they choose not to get circumcised. (Although protocols do not restrict MMC to men who are HIV-negative, it is assumed that HIV risk reduction is the primary reason for seeking MMC, and thus a man who is diagnosed HIV-positive would have less incentive to get medically circumcised.)

##### 5.2.1.7 Testing of men in prison

HIV testing is commonly offered to prisoners at the time of admission to prison. In addition, prisoners are often able to request HIV testing at other times. The Department of Correctional Services only publishes the total number of HIV tests performed in prisons each year (i.e. with no split between new prisoners and those who have been incarcerated for longer durations), and we therefore model HIV testing in prisons as occurring at an annual rate, which is independent of the time since entry into prison. The model assumption about the rate of testing in a given year is obtained by dividing the reported total number of HIV tests in each year by the estimated number of prisoners in that year. The resulting assumed rates of testing are shown in Table 5.2.3. Previously-diagnosed prisoners are assumed to be 50% as likely to test for HIV as prisoners who have not previously tested for HIV (or 15% as likely if they are already receiving ART).

##### 5.2.1.8 Testing of individuals receiving pre-exposure prophylaxis

This is described in more detail in section 5.5.5.

#### 5.2.2 Modelling effect of HIV diagnosis on existing partnerships

It is assumed that individuals who are already in a sexual relationship at the time they are diagnosed with HIV do not modify their condom usage or coital frequency with their partner until they have disclosed their HIV status to their partner. Disclosure is modelled as occurring with a once-off probability, at the time of HIV diagnosis, or – if disclosure does not occur at that time – at a continuous rate thereafter (taking into account that some individuals may need time before they are able to disclose their status to their partner). It is assumed that following HIV disclosure, there is both heightened risk of the relationship ending and increased likelihood of condoms being used.

##### 5.2.2.1 Probabilities of disclosure

Table 5.2.4 summarizes South African studies that have measured rates of disclosure to sexual partners. The proportion of HIV-positive patients who have disclosed their HIV status to their sexual partners is highly variable, ranging between 17% and 95%. However, several factors account for this variation. A recent study in Uganda found that rates of disclosure tended to be higher at longer durations since HIV diagnosis, higher in the ART era, higher in relationships of longer duration, and higher in men when compared to women [301]. A Tanzanian study also found that disclosure was significantly associated with marital status, time since diagnosis and being on ART, although this study did not focus on disclosure to sexual partners specifically [302]. It was also found that the rate of HIV disclosure increased significantly if the HIV-diagnosed individual had experienced AIDS symptoms (aHR 1.56), was in pre-ART care (aHR 2.09) or was on ART (aHR 3.26). In a South African study conducted in two communities (Soweto and Vulindelela), Wong *et al* [322] found that disclosure to sexual partners was significantly associated with longer time since HIV diagnosis, older age, and higher socio-economic status. In another South African study in Pretoria, HIV-diagnosed women were more likely to disclose their HIV status to sexual partners if they were married (aOR 2.3, 95% CI: 1.2-4.5) or if their partner had tertiary education (aOR 2.8, 95% CI: 1.3-5.9) [298]. In a study conducted in Cape Town, disclosure was significantly higher in patients who reported living with their sexual partner and in patients who were on ART (aOR 1.6, 95% CI: 1.1-2.3) [299]. Thus the low rates of disclosure observed in the early studies (52% or less) may be explained by the fact that these studies were mostly done in the early 2000s, when access to ART was still very limited; in addition, most of these studies sampled only women who had been recently diagnosed [57, 298, 323, 324]. Women receiving PMTCT in the early 2000s would mostly have been receiving only a single dose of nevirapine at the time of going into labour, whereas women receiving PMTCT in 2008 and later years would have received daily antiretroviral prophylaxis that was more difficult to conceal from sexual partners; this may explain the higher rates of disclosure in some of the more recent PMTCT studies [325, 326].

**Table 5.2.4:** South African studies of HIV disclosure to sexual partners

| Reference | Year | Population | Time since diagnosis | % who had disclosed |
| --- | --- | --- | --- | --- |
| Kuhn *et al* [323] | 1997 | HIV-positive mothers | - | 34% |
| Hilderbrand *et al* [324] | - | Women participating in PMTCT programme | Around 6 months<sup>a</sup> | 45% |
| Sethosa & Peltzer [327] | - | VCT clients in a hospital setting | 5 months | 17% |
| Wong *et al* [322] | 2003 | HIV+ individuals identified in a household survey | 13 months | 64% |
| Myer *et al* [44] | 2005 | Women on ART | >11 months<sup>b</sup> | 85% |
|  |  | Men on ART | >8 months<sup>b</sup> | 95% |
| Peltzer *et al* [57] | 2003-4 | Women participating in PMTCT programme | 12 months<sup>c</sup> | 51.7% |
| Eisele *et al* [328] | 2006 | Patients who started ART <3 months ago | - | 67.6% |
|  | 2007 | Patients who started ART >12 months ago | >12 months<sup>b</sup> | 72.7% |
| Le Roux *et al* [326] | 2009-2010 | Pregnant women | - | 78.2% |
| Mnyani & McIntyre [325] | 2009 | Pregnant women | >3 months | 79.9% |
| McGrath *et al* [329] | 2009-2011 | Pre-ART patients, CD4 <200 | >1 month<sup>d</sup> | 72% |
|  |  | Pre-ART patients, CD4 >500 | >9 months<sup>d</sup> | 69% |
| Makin *et al* [298] | 2003-2004 | Pregnant women | 1 week | 47.7% |
| Varga *et al* [330] | 2002-2003 | Pregnant women | Around 7 months<sup>e</sup> | 80% |
| Mkwanazi *et al* [300] | - | Women diagnosed during pregnancy | 6-8 years | 65% |
| Skogmar *et al* [331] | 2003 | Patients at HIV clinics (35% on ART) | >2 years | 79.1% |
| Simbayi *et al* [332] | - | Patients at HIV clinics | 2.7 years | 58% |
| Nachega *et al* [333] | 2002 | Patients at HIV clinics | 3.5 years | 62% |

<sup>a</sup> Not stated, but most women would have been diagnosed around the time of their first antenatal visit (usually 3 months before delivery) and the average age of their infants at the time of the interview was 3 months. <sup>b</sup> Only the duration of ART was reported; the time since diagnosis would be longer. <sup>c</sup> Not stated, but most women would have been diagnosed around the time of their first antenatal visit (usually 3 months before delivery) and the median age of their infants at the time of the interview was around 9 months. <sup>d</sup> Only the time since entry into HIV care was reported; the time since diagnosis would be longer. <sup>e</sup> Women were enrolled within 3 months after receiving the results of their infant's HIV test; since the first antenatal test would usually occur around 3 months prior to birth and collection of infant PCR results would usually occur around 2-3 months after birth, it is assumed that the average time since diagnosis was 7 months at enrolment.

For women who are diagnosed with HIV, not yet on ART, and not in a marital relationship, we set the assumed probability of disclosure at the time of diagnosis to 0.50, based on the average of the 5 studies that were conducted in pregnant women prior to the widespread availability of ART (most pregnant women are not in cohabiting/marital relationships) [57, 298, 323, 324, 330]. This baseline rate is adjusted by an odds ratio of 1.25 for men [44, 301, 302] and by an odds ratio of 2.5 if the partner is a marital/cohabiting partner [298–302]. Evidence regarding the effect of socio-economic factors on disclosure is conflicting [298, 299, 322, 334], and hence no adjustment is made in relation to educational attainment.

If individuals do not disclose their HIV status to their sexual partner at the time of diagnosis, it is assumed that the annual rate of disclosure thereafter is 0.06 [301, 328, 331]. As in the previous paragraph, this baseline rate applies to unmarried women who are not on ART; the baseline rate is adjusted by the same multiplicative factors as before to account for the effects of sex and marital status. In addition, the baseline rate is adjusted by an odds ratio of 3 if the individual is on ART [44, 299, 301, 302, 328].

##### 5.2.2.2 Effect of disclosure on condom use

Most South African studies have found that disclosure of HIV-positive status to sexual partners is associated with increased condom use. Wong *et al* [322] found that in HIV-diagnosed individuals, 70% of those who had disclosed their HIV status to their sexual partner reported always using condoms, compared to 44% in those who had not disclosed. Similarly, Simbayi *et al* [332] found that among HIV patients, the reported rate of condom use was 75% in those who had disclosed to all their recent partners, compared to 46% among those who had not disclosed their HIV status. Eisele *et al* [335] also found that HIV-positive individuals in Cape Town were more than twice as likely to report using condoms at last sex if they had disclosed their HIV status to their partner (adjusted odds ratios were 2.57 for men and 2.84 for women). However, Toska *et al* [336] found that among HIV-positive adolescents in the Eastern Cape (most of whom acquired HIV through vertical transmission), condom use was not significantly associated with disclosure. The authors note that many adolescents reported postponing the disclosure until the relationship was judged to be stable, and thus any effect of disclosure on condom use may have been offset by the effect of relationship duration on condom use (as noted in section 4.9, condom use is less frequent in more established relationships).

Levels of condom use after HIV disclosure may depend also on whether the other partner already knows they are HIV-positive. Although HIV-positive individuals are usually counselled to use condoms even if they know their partner is HIV-positive (in order to avoid super-infection), studies in South Africa suggest that condom use in HIV-discordant relationships is typically more frequent than in concordant-positive relationships [332, 337, 338]. However, one study in rural KwaZulu-Natal found that condom use was less frequent in discordant relationships (or relationships in which the partner’s HIV status was unknown) than in concordant positive relationships [329]. This might be because individuals who know their partner is HIV-positive are likely to have disclosed their HIV status to one another, whereas partnerships in which the partner’s HIV status is unknown are likely to be partnerships in which the HIV-positive individual has not disclosed their HIV status. Thus the effect of HIV-positive concordance may be confounded with the effect of HIV disclosure. Since none of the South African studies had sufficiently large numbers of individuals in known serodiscordant relationships (or did not report the number separately from the number of individuals who were unsure of their partner’s HIV status), it is difficult to quantify the effect of the partners’ HIV concordance on the rate of condom use independently of the effect of disclosure. Since a substantial fraction of individuals in concordant-positive relationships (35-75%) report consistent condom use, and since evidence from other African countries suggests that concordant-positive couples actually have higher condom use than concordant-negative couples [339], it is unlikely that HIV-concordant couples substantially reduce condom use after disclosure. It is therefore assumed that if an HIV-positive individual discloses their HIV status to a partner who already knows they are HIV-positive, there is no change in condom use, but that if they disclose their HIV status and their partner’s status is either negative or unknown, there is a 3-fold increase in the odds of condom use (relative to what it would have been if the couple had not been aware of their HIV status). The multiple of 3 is based on the studies cited in the previous paragraph. If a couple starts to use condoms consistently, and one or both partners have been diagnosed positive, it is assumed that there will be no discontinuation of condom use when the relationship becomes cohabiting, due to concern about potential HIV transmission.

It is worth noting that there is evidence from other African countries of over-reporting of condom use by individuals who have been diagnosed HIV-positive [340], similar to the evidence of over-reporting of condom use in the general population of South Africa [80]. For this reason we specify the effect of disclosure on condom use as an odds ratio rather than as an absolute level of condom use, since the modelled baseline levels of condom use are already adjusted for likely over-reporting in sexual behaviour surveys.

##### 5.2.2.3 Effect of disclosure on relationship dissolution

No South African studies have assessed the effect of HIV status on rates of relationship dissolution. However, the fraction of relationships that terminate soon after disclosure of HIV-positive status appears to vary substantially in studies of HIV-positive South African women: from 2.8% in a study conducted in Pretoria [341] to 16% in a Johannesburg study [330] and 27% (8/30) in a study conducted in Cape Town [342]. There is evidence from other African countries to suggest that partnerships in which one or both partners are diagnosed HIV-positive have a higher rate of dissolution than partnerships in which both partners are HIV-negative (Table 5.2.5). This is especially true when the female partner is HIV-positive and the male partner is HIV-negative. Anglewicz and Reniers [343] also found that in Malawi, married couples were more likely to get divorced if they were HIV-positive, although this effect was only statistically significant in women. There appears to be little difference in rates of dissolution when comparing partnerships in which both partners are HIV-negative and partnerships in which the male partner is HIV-positive. Based on these data, it is assumed that the rate of relationship dissolution is increased by a factor of 2.5 if the female partner discloses her HIV-positive status and her partner is HIV-negative (or has undiagnosed HIV), but that there is no difference in rates of union dissolution if the male partner discloses his HIV status.

**Table 5.2.5:** Effect of HIV status on rates of relationship dissolution

| Study | Location | Definition of dissolution | Dissolution by couple serostatus |  |  |  |
| --- | --- | --- | --- | --- | --- | --- |
|  |  |  | M-F- | M+F- | M-F+ | M+F+ |
| Mackleprang *et al* [344] | Kenya | Dissolution over 2 years of follow-up | - | 22% | 29% | - |
| Floyd *et al* [345] | Malawi | Annual rate of dissolution<sup>a</sup> | 0.0261 | 0.0277<sup>b</sup> | 0.0578<sup>b</sup> | - |
| Grinstead *et al* [346] | Kenya, Trinidad and Tanzania | Marital dissolution | 3% | 0% | 20% | 7% |
|  |  | Relationship dissolution | 22% | 38% | 45% | 23% |
| Porter *et al* [347] | Uganda | Odds ratio for divorce/separation | 1.00 | 1.04 | 3.05 | 1.57 |

<sup>a</sup> Calculated from reported numbers of surviving relationships, separations and deaths. <sup>b</sup> HIV status of partners was not reported, so it is assumed partners were HIV-negative.

#### 5.2.3 Modelling effect of HIV diagnosis on new partnerships

It is assumed that due to the effect of HIV-related illness, HIV-positive individuals with low CD4 counts are less likely to form new partnerships than HIV-negative individuals or HIV-positive individuals with high CD4 counts (see section 4.3). However, HIV diagnosis is assumed to have no effect on the rate at which new partners are acquired. For the sake of simplicity, the effect of HIV diagnosis is assumed to be limited to relationship dissolution and condom use (as discussed previously) and the probability of choosing an HIV-positive partner.

Although many studies have shown evidence of gay men in high income countries choosing sexual partners of the same HIV status (‘serosorting’), there has been almost no research on the extent of serosorting in generalized heterosexual epidemics [348]. However, there is anecdotal evidence from various African countries of ART patients meeting partners through HIV support groups, and of match-making agencies matching individuals on the basis of HIV status [348–352]. A recent study from rural KwaZulu-Natal also found that HIV-positive individuals married other HIV-positive individuals at a higher rate than that at which HIV-negative individuals married HIV-positive partners (aHR 2.52, 95% CI: 1.43-4.45) [353]. To model this dynamic, we assume that ART patients are 50% less likely to select an individual as a new partner if they are HIV-negative or untreated, than if the individual was on ART. Although there may also be a degree of selection when individuals are HIV-positive but not on ART, the literature reviewed in section 5.2.2 suggests that untreated HIV-positive individuals are generally less open about their HIV status. Since they do not attend HIV services as frequently as ART patients, they would probably also find it more difficult to identify HIV-positive partners.

It is assumed that if a new partnership forms between two individuals who are both on ART, both partners disclose their HIV status to each other at the start of the new relationship, and the level of condom use in that relationship is the same as would be expected in an established relationship in which disclosure takes place between two HIV-positive partners. The assumption of immediate disclosure is made because it is assumed that many ART patients would meet partners through HIV support groups, or if they met outside of the context of a support group, would be unlikely to conceal their HIV status for long.

In all other cases, the annual rate of disclosure in a new relationship is assumed to be the same as that in established relationships, i.e. disclosure occurs at a rate of 0.06 per annum in the case of unmarried women who are not on ART, with this rate increasing by a factor of 3 in ART patients, by a factor of 2.5 in married individuals and by a factor of 1.25 in men. Prior to disclosure, the probability of condom adoption is multiplied by a factor of 2, and after disclosure, the probability of condom adoption is multiplied by a factor of 3 if the other partner is HIV-negative or is HIV-positive but undiagnosed (in both cases, relative to the probability that would be expected if neither partner knew their HIV status). The adjustment to condom use in serodiscordant relationships is thus the same as that assumed previously in established relationships (where diagnosis occurs after the relationship was established). However, we assume that HIV-diagnosed individuals would be more likely to attempt to initiate condom use in a new relationship, even if they have not disclosed their HIV status, and the adjustment factor of 2 is therefore the average of the adjustment factor that would apply if only the HIV-diagnosed partner could determine the condom use (3) and the adjustment factor that would apply if only the HIV-negative/undiagnosed partner could determine the condom use (1). It follows that if both partners are HIV-diagnosed but they have not disclosed their HIV status to each other, the condom adjustment factor is 3 (the average of 3 and 3).

A mathematical explanation of the approach to modelling the effect of disclosure on condom use is provided in section 4.9.1.

#### 5.2.4 Model validation

The model assumptions about rates of HIV diagnosis and changes in behaviour after HIV diagnosis can be validated in a number of ways. Firstly, Figure 5.2.2 compares the fraction of HIV-positive adults who have been diagnosed in the Thembisa and MicroCOSM models. Since MicroCOSM assumptions about rates of HIV testing are similar to those in Thembisa, we would expect the diagnosed proportions to be similar, as Figure 5.2.2 shows.

**Figure 5.2.2:**
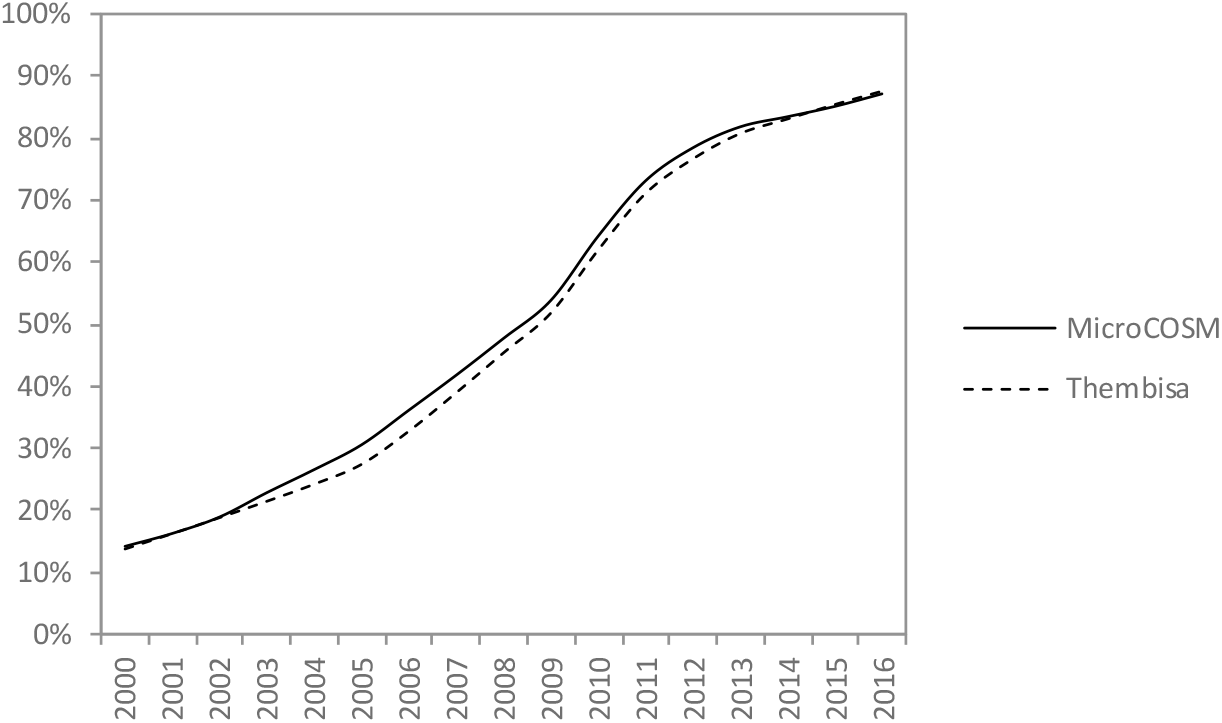
Fraction of HIV-positive adults (ages 15 and older) who have been diagnosed HIV-positive. The diagnosed fraction in the Thembisa model is calculated using Thembisa version 2.5 [354]. The diagnosed fraction in MicroCOSM is the average from 100 simulations.

A second approach to validation is to compare the model estimates of the relative frequency of unprotected sex in HIV-diagnosed adults with (a) empirical estimates and (b) estimates from the Thembisa model. Table 5.2.6 summarizes empirical estimates of the relative frequency of unprotected sex in HIV-diagnosed adults and adults who are either HIV-negative or HIV-positive but undiagnosed. Studies were only included in this table if they were conducted in developing countries and if they used multivariate analysis to control for confounding factors. Applying a random effects meta-analysis to the empirical data yields a pooled odds ratio of 0.39 (95% CI: 0.28-0.56) for the effect of HIV diagnosis on unprotected sex, with significant heterogeneity between studies (p < 0.001). The pooled odds ratio differed when considering only the studies in which the controls were HIV-negative (OR 0.60, 95% CI: 0.53-0.68) and the studies in which the controls were HIV-positive but undiagnosed (OR 0.22, 95% CI: 0.08-0.61), with the heterogeneity between studies being significant for the latter group of studies (p < 0.001) but not the former (p = 0.48). The pooled odds ratio was also higher when considering only the four studies conducted in South Africa (OR 0.55, 95% CI: 0.45-0.68), with little evidence of heterogeneity (p = 0.11).

**Table 5.2.6:**
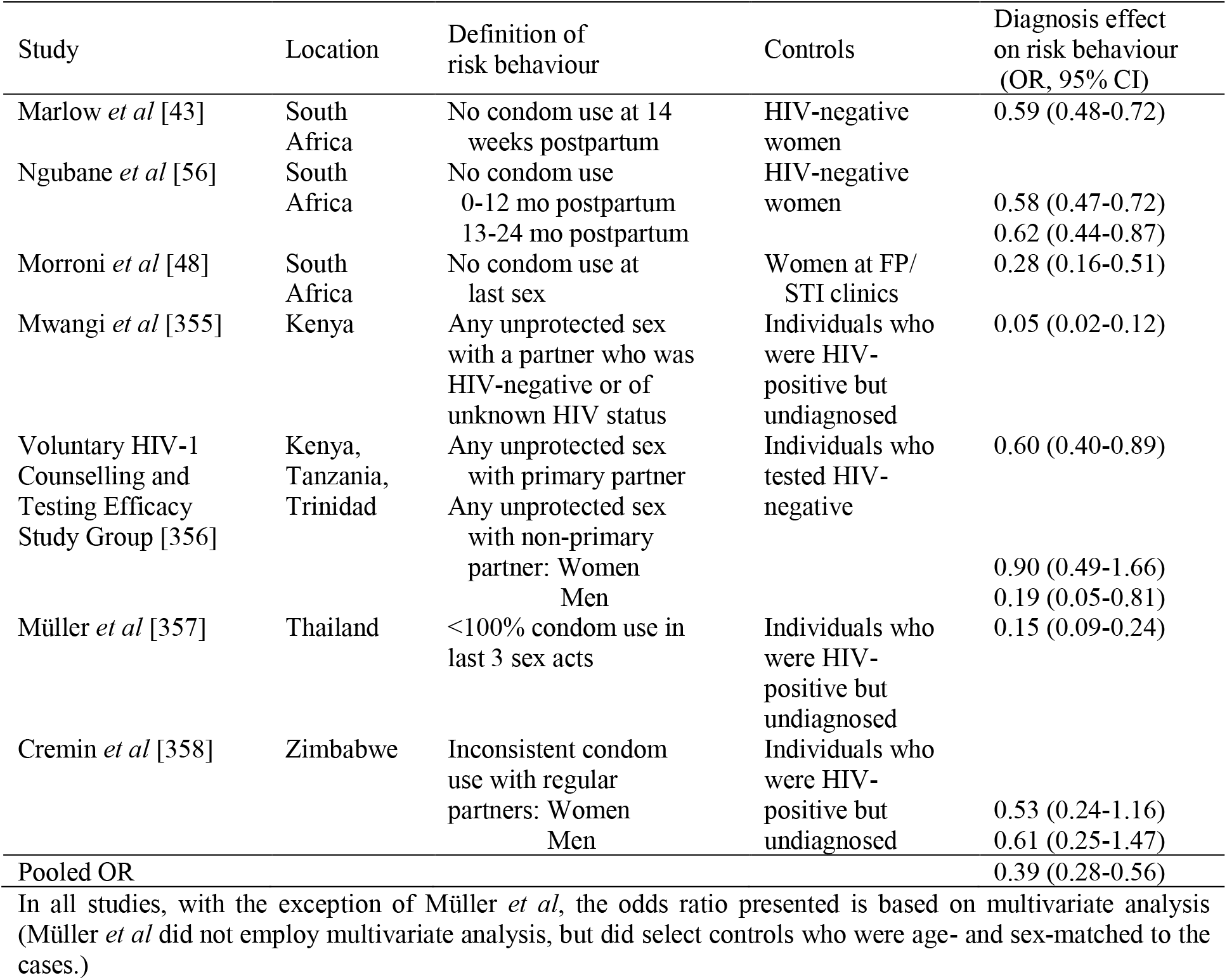
Studies evaluating the effect of HIV diagnosis on unprotected sex in developing countries

The Thembisa model adopts a Bayesian approach in estimating the proportionate reduction in unprotected sex following HIV diagnosis. A prior distribution is specified based on the same empirical evidence as summarized in Table 5.2.6, and a likelihood function is specified to represent the goodness of model fit to the age-specific HIV prevalence trends, for a given set of model assumptions. The posterior estimates of the reduction in unprotected sex after HIV diagnosis are then calculated by integrating the prior and likelihood function. The posterior estimate of the relative rate of unprotected sex following HIV diagnosis was estimated as 0.63 (95% CI: 0.57-0.69) in Thembisa version 1.0 [256] and 0.54 (95% CI: 0.42-0.67) in Thembisa version 2.4 [236]. These rates are thus consistently higher than the pooled odds ratio in Table 5.2.6, which indicates that the HIV prevalence data in South Africa do not suggest reductions in HIV risk behaviour as dramatic as those estimated from self-reported data. The Thembisa model makes adjustments to take into account the effects of factors such as age, sex and marital status on the level of unprotected sex; in addition, further reductions in unprotected sex are assumed to occur after individuals start ART.

To meaningfully compare these estimates with the estimates from the microsimulation model, we store the microsimulation results for all adults who were sexually active in 2005, across 3 simulations (*n* = 43 111) and perform a multivariate logistic regression on the results to evaluate the effects of HIV diagnosis on unprotected sex. The analysis is based on 2005 results because inclusion of later results could be biased by a substantial fraction on ART (since the Thembisa estimates represent the effect of HIV diagnosis on unprotected sex prior to ART initiation, and since most of the HIV-diagnosed individuals in the empirical studies in Table 5.2.6 were also not on ART). After controlling for age, sex, marital status and number of partners, sexually active HIV-diagnosed adults were found to have a significantly reduced odds of engaging in unprotected sex when compared to HIV-negative adults (aOR 0.68, 95% CI: 0.60-0.77), and the reduction was more substantial in a separate analysis in which the comparison group was HIV-positive adults who were undiagnosed (aOR 0.57, 95% CI: 0.50-0.66). This difference in odds ratios occurs because the individuals who acquire HIV are usually those with a lower condom preference, and thus HIV-negative individuals tend to have higher levels of condom use than HIV-positive undiagnosed individuals.

Figure 5.2.3 compares the estimates of the relative frequency of unprotected sex after HIV diagnosis from the different sources. The microsimulation model estimates appear to be roughly consistent with the Thembisa model estimates, although strictly speaking, the metrics being compared are not defined consistently (the Thembisa metric is a relative frequency of unprotected sex, while the microsimulation metric is an odds ratio for any unprotected sex). The model estimates are higher than the empirical estimates, as represented by the meta-analysis. However, the extreme heterogeneity in the empirical estimates suggests that the pooled odds ratio should be treated with caution; it is likely that the true extent of the reduction in risk behaviour after diagnosis differs between settings. When the meta-analysis is limited to the South African studies, there is greater consistency between the pooled odds ratio (OR 0.55, 95% CI: 0.45-0.68) and the modelled ratios.

**Figure 5.2.3:**
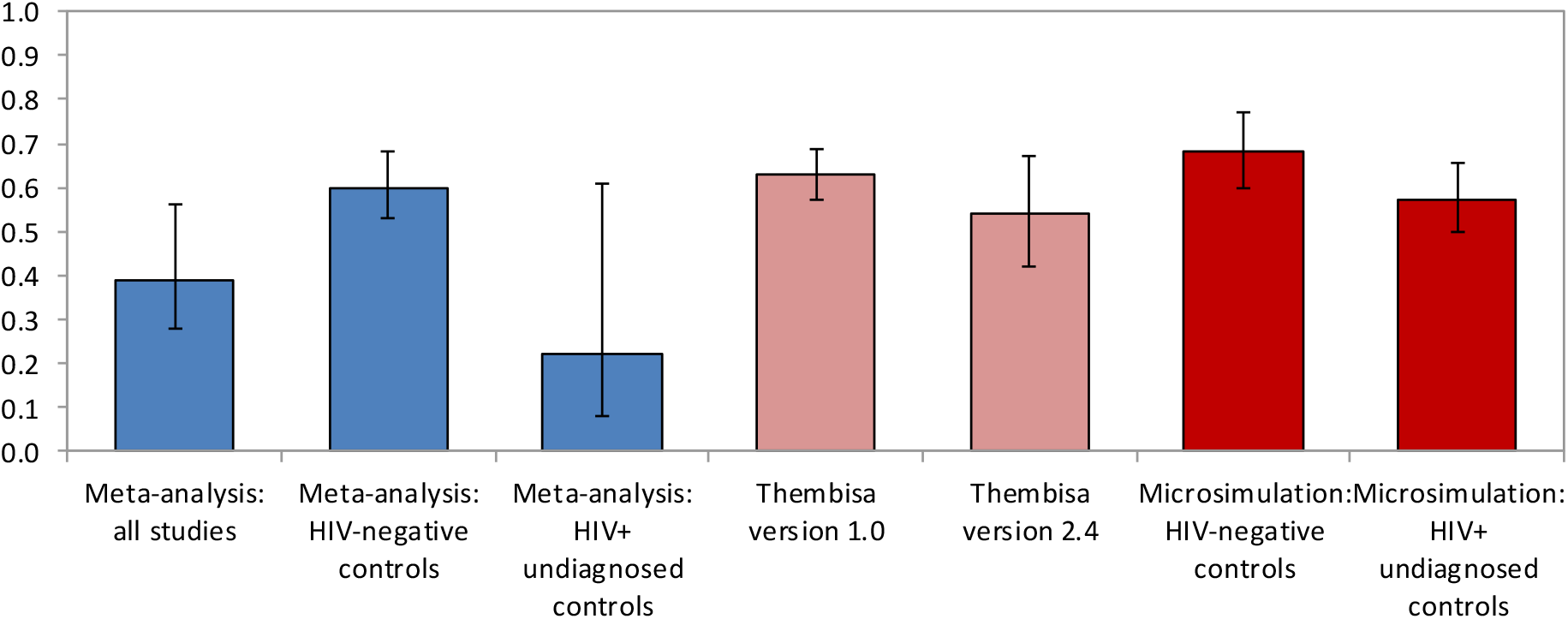
Ratio of unprotected sex in HIV-diagnosed individuals to that in HIV-negative or undiagnosed individuals. The meta-analysis and microsimulation estimates are expressed as odds ratios, while the Thembisa estimates are expressed as relative frequencies of unprotected sex.

### 5.3 Antiretroviral treatment (ART)

The approach adopted in modelling adult uptake of ART is similar to that in the Thembisa model [236]. Individuals are assumed to start ART either at the time of HIV diagnosis (if they are eligible) or at a later time, based on an assumed rate of ART initiation in previously-diagnosed individuals. ART eligibility assumptions are the same as in the Thembisa model: eligibility is assumed to have changed over time, and is defined with reference to the individual’s CD4 count, pregnancy status and experience of HIV-related symptoms. After starting ART, HIV-related mortality is assumed to depend on both time since ART initiation and baseline CD4 count (see section 6.3).

#### 5.3.1 ART initiation soon after HIV diagnosis

Table 5.3.1 summarizes the assumed proportions of HIV-positive adults who are eligible to start ART in different CD4 categories, in different periods (these assumptions apply only to adults who are asymptomatic and not pregnant). These assumptions have been set with reference to ART guidelines in South Africa [359–363], but have also been modified to take into account partial access to ART through private providers and NGO programmes that were using more liberal ART eligibility criteria. The table also shows the assumed proportions of eligible individuals who start ART immediately after a diagnosis (again assuming that individuals are asymptomatic and not pregnant). These proportions are assumed to have remained stable at 40% in recent years, in line with a review of linkage studies [364]. The South African linkage studies included in this review estimated that on average only 40% of patients who were newly diagnosed linked to ART soon after diagnosis. The 40% is scaled down in earlier years in order to reflect the limited availability of ART in the early stages of the ART rollout.

**Table 5.3.1:** Adult ART eligibility and uptake after HIV diagnosis (asymptomatic, not pregnant)

| Year | % eligible at CD4 count |  |  |  | Starting ART soon after HIV diagnosis | Annual ART initiation rate if previously diagnosed, with CD4 <200 |  |
| --- | --- | --- | --- | --- | --- | --- | --- |
|  | <200 | 200-349 | 350-499 | 500+ |  | Male | Female |
| Pre-2000 | 0% | 0% | 0% | 0% | 0.0% | 0.000 | 0.000 |
| 2000-01 | 100%* | 10%* | 0% | 0% | 1.0% | 0.069 | 0.055 |
| 2001-02 | 100%* | 10%* | 0% | 0% | 1.2% | 0.059 | 0.045 |
| 2002-03 | 100%* | 10%* | 0% | 0% | 1.2% | 0.053 | 0.039 |
| 2003-04 | 100%* | 10%* | 0% | 0% | 1.8% | 0.102 | 0.080 |
| 2004-05 | 100% | 10%* | 0% | 0% | 6.3% | 0.162 | 0.183 |
| 2005-06 | 100% | 10%* | 0% | 0% | 11.3% | 0.161 | 0.208 |
| 2006-07 | 100% | 10%* | 0% | 0% | 14.5% | 0.221 | 0.302 |
| 2007-08 | 100% | 10%* | 0% | 0% | 17.8% | 0.225 | 0.341 |
| 2008-09 | 100% | 10%* | 0% | 0% | 22.3% | 0.281 | 0.432 |
| 2009-10 | 100% | 10%* | 0% | 0% | 27.5% | 0.373 | 0.539 |
| 2010-11 | 100% | 20% | 0% | 0% | 32.1% | 0.471 | 0.774 |
| 2011-12 | 100% | 80% | 0% | 0% | 37.7% | 0.301 | 0.524 |
| 2012-13 | 100% | 100% | 0% | 0% | 40.0% | 0.252 | 0.498 |
| 2013-14 | 100% | 100% | 0% | 0% | 40.0% | 0.212 | 0.458 |
| 2014-15 | 100% | 100% | 50% | 0% | 40.0% | 0.152 | 0.224 |
| 2015-16 | 100% | 100% | 100% | 0% | 40.0% | 0.187 | 0.311 |
| 2016-17 | 100% | 100% | 100% | 80% | 40.0% | 0.223 | 0.400 |
| 2017-18 | 100%† | 100%† | 100%† | 100%† | 40.0%† | 0.261 | 0.490 |

\* Reflecting availability in the private and NGO sectors. † The same rates are assumed to apply in all years after 2018.

Different eligibility criteria and rates of ART initiation are assumed to apply in the context of pregnant women and patients with HIV-related opportunistic infections (OIs), as shown in Table 5.3.2. As in Table 5.3.1, all individuals with CD4 counts <200 cells/μl are assumed to be eligible in the period after 2000 (assumptions not shown). The proportions of OI patients eligible in each CD4 category is calculated based on the relative incidence of WHO stage III and IV conditions in each CD4 stage [275, 276] and the guidelines on which clinical conditions defined eligibility for ART in different CD4 stages. The fraction of women diagnosed with HIV during pregnancy who start ART soon after diagnosis is set to 93% in 2015-16 [293], but lower fractions are assumed to apply in earlier periods, in part because of the limited access to ART in the early years of the ART programme, and in part because ART services were not integrated into antenatal care in the early years of the programme [365–367]. The fraction of OI patients who start ART soon after HIV diagnosis has been found to be lower than that in pregnant women [368], and the assumed fraction of OI patients starting ART after diagnosis has been set to 78% in 2015-16, slightly lower than the fraction of TB patients with HIV who were on ART in this year (85% [293]) because many of these TB patients were already on ART before diagnosis. Uptake in OI patients is assumed to have changed over time, in proportion to the annual changes in uptake among pregnant women.

**Table 5.3.2:** Adult ART eligibility and uptake after HIV diagnosis (symptomatic patients and pregnant women)

| Year | % pregnant women eligible at CD4 |  | % of OI patients eligible at CD4 |  |  | Starting ART soon after HIV diagnosis |  |
| --- | --- | --- | --- | --- | --- | --- | --- |
|  | 200-349 | 350+ | 200-349 | 350-499 | 500+ | Pregnant | OI patient |
| Pre-2000 | 0% | 0% | 0% | 0% | 0% | 0% | 0% |
| 2000-01 | 10%* | 0% | 24% | 17% | 20% | 1.9% | 1.6% |
| 2001-02 | 10%* | 0% | 24% | 17% | 20% | 2.5% | 2.1% |
| 2002-03 | 10%* | 0% | 24% | 17% | 20% | 2.5% | 2.1% |
| 2003-04 | 10%* | 0% | 24% | 17% | 20% | 3.6% | 3.0% |
| 2004-05 | 10%* | 0% | 24% | 17% | 20% | 12.6% | 10.6% |
| 2005-06 | 10%* | 0% | 24% | 17% | 20% | 22.6% | 19.0% |
| 2006-07 | 10%* | 0% | 24% | 17% | 20% | 29.1% | 24.4% |
| 2007-08 | 10%* | 0% | 24% | 17% | 20% | 35.5% | 29.8% |
| 2008-09 | 10%* | 0% | 24% | 17% | 20% | 44.5% | 37.3% |
| 2009-10 | 50% | 0% | 30% | 17% | 20% | 55.0% | 46.1% |
| 2010-11 | 100% | 0% | 37% | 17% | 20% | 64.1% | 53.8% |
| 2011-12 | 100% | 0% | 37% | 17% | 20% | 75.4% | 63.2% |
| 2012-13 | 100% | 0% | 100% | 100% | 100% | 75.9% | 63.7% |
| 2013-14 | 100% | 0% | 100% | 100% | 100% | 76.3% | 64.0% |
| 2014-15 | 100% | 50% | 100% | 100% | 100% | 91.2% | 76.5% |
| 2015-16 | 100% | 100% | 100% | 100% | 100% | 93.0% | 78.0% |
| 2016-17 | 100% | 100% | 100% | 100% | 100% | 93.0% | 78.0% |
| 2017-18† | 100%† | 100%† | 100%† | 100%† | 100%† | 93.0%† | 78.0%† |

\* Reflecting availability in the private and NGO sectors. † The same rates are assumed to apply in all years after 2018.

The model also allows for the possibility that individuals who have already been diagnosed may be ‘re-diagnosed’ and a once-off probability of ART initiation is assumed to apply at the time of re-diagnosis, in the same way as for first diagnoses. The probability of starting ART immediately after the retest is assumed to be the same as it would have been if they were being diagnosed for the first time, as a South African study found that previously-diagnosed individuals (who had not previously linked to care) had the same odds of linking to care after a retest as individuals who were newly diagnosed [369].

#### 5.3.2 ART initiation in previously-diagnosed individuals

Individuals who do not start ART soon after diagnosis can initiate ART at a later time. The model assumes that the annual rate of ART initiation in previously-diagnosed individuals (i.e. not recently diagnosed or re-diagnosed) changes over time, and depends on sex and CD4 count. The assumed annual rates that apply for men and women with CD4 counts <200 cells/μl are shown in the last two columns of Table 5.3.1. These are estimated from the Thembisa model (version 2.5), which has been calibrated to recorded total numbers of ART patients in South Africa up to 2015 [354]. Rates rose to their highest levels in 2010-11, then steadily declined. Although it is unclear what rates might be expected in the post-2015 period, we have optimistically assumed that the average rate will tend towards 0.333 and 0.667 per annum in men and women respectively.

Studies suggest that HIV-diagnosed individuals who were previously diagnosed have a lower rate of ART initiation and retention in pre-ART care the higher their CD4 count [370–372]. Based on this evidence, it is assumed that the rates of ART initiation for previously-diagnosed individuals with CD4 counts <200 cells/μl (as shown in Table 5.3.1) are scaled down by factors of 0.70 and 0.45 at CD4 counts of 200-349 and ≥350 cells/μl respectively. These rates apply only if the individual is eligible to start ART (as shown in the first columns of Table 5.3.1).

#### 5.3.3 ART interruption and ART re-initiation

The model allows for the possibility that individuals may interrupt ART, and also allows for the possibility that ART may be re-initiated after an interruption. Based on recent South African studies [373–375], we assume that the annual rate of interrupting ART is 0.25 and the annual rate of resuming ART after an interruption is 0.90. This implies that modelled ART interruptions are frequent but usually of relatively short duration. Figure 5.3.1 shows that with these assumptions about ART initiation, interruption and re-initiation, the model matches quite closely the results of the Thembisa model, which had been calibrated to South African ART data [354].

**Figure 5.3.1:**
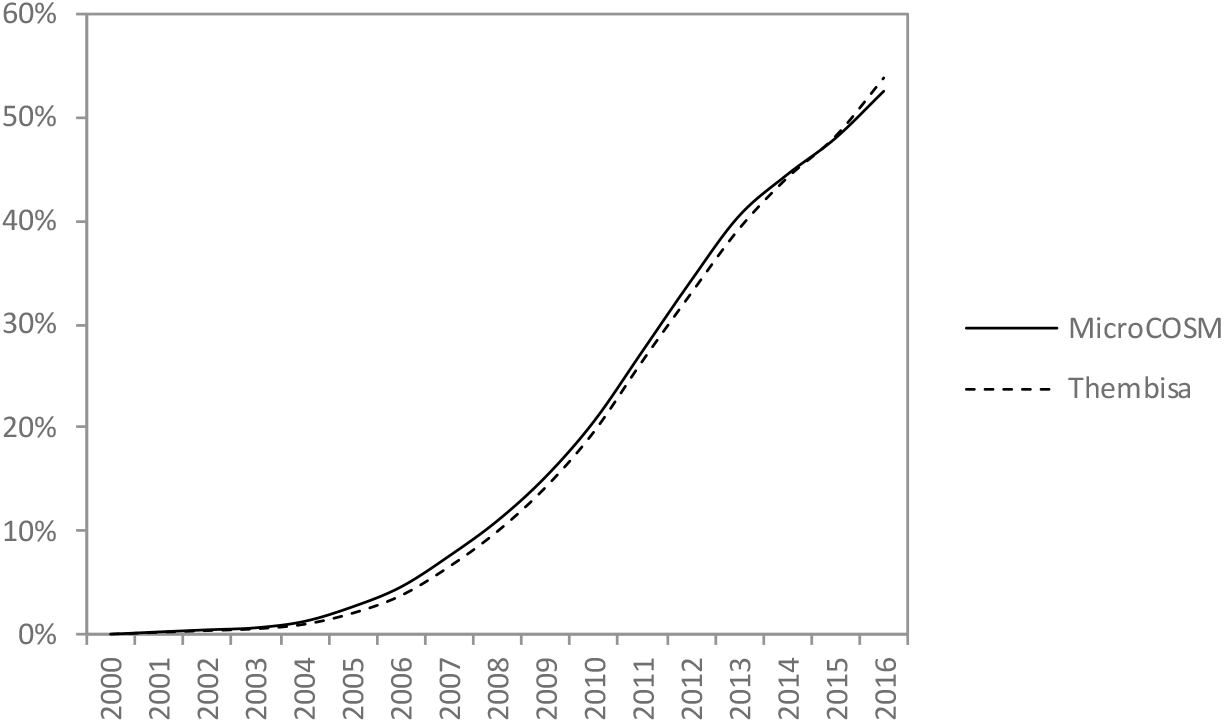
ART coverage among HIV-positive adults. The ART coverage in the Thembisa model is calculated using Thembisa version 2.5 [354]. The diagnosed fraction in the MicroCOSM model is the average from 100 simulations.

### 5.4 Sexually transmitted infection (STI) treatment

Assumptions about STI treatment are the same as in the STI-HIV Interaction model, as described previously [318]. Briefly, individuals who are experiencing STI symptoms are assumed to seek STI treatment at a rate that depends on their age and sex (the model does not consider treatment for asymptomatic infections, as current syndromic management protocols in South Africa do not address asymptomatic STIs). Individuals are assumed to seek STI treatment either from public clinics, formal private providers or traditional healers. STI patients are managed according to syndromic management protocols with a given probability, this probability depending on the type of provider (highest in the public sector) and the calendar year (syndromic management protocols having been phased in after 1994). The probability that the STI is cured is assumed to depend on whether syndromic management protocols are followed, with treatment practices prior to syndromic management generally being considered inferior [315]. The probability of cure is also assumed to depend on the individual STI, and is assumed to change over time as a result of drug stockouts (a particular challenge in the public sector) and drug resistance (a particular problem in the case of gonorrhoea).

### 5.5 Pre-exposure prophylaxis (PrEP)

#### 5.5.1 PrEP efficacy and adherence

It is well-established that the efficacy of PrEP is highly dependent on PrEP adherence [376, 377], with efficacy being close to 90% in populations in which there is high adherence [378, 379]. Unfortunately, levels of PrEP adherence have been relatively low in the PrEP trials that have been conducted in South Africa, and local PrEP trials have therefore produced disappointing results [380, 381]. In a recent meta-analysis, it was estimated that PrEP reduced the rate of heterosexual HIV transmission by an average of 46%, as compared to an average reduction of 66% in MSM [377]. Recent evidence suggests that a higher dosing frequency is required to achieve protection in the female genital tract than in rectal tissue [382], which may explain this difference in efficacy.

It is worth noting that the data included in the meta-analysis all came from randomized controlled trials (RCTs), which were conducted prior to the efficacy of PrEP being established. Trial participants might have been less motivated to adhere to PrEP because of uncertainty regarding its efficacy as well as uncertainty about whether they had been randomized to the PrEP or placebo arm. In subsequent studies, in which patients knew that they were taking PrEP and that it was effective, higher levels of efficacy have been established, for example 86% in two recent studies of MSM in high-income countries [378, 379]. Recent PrEP studies in South African women have found relatively high rates of adherence [383, 384]. This suggests that the ‘real world’ efficacy of PrEP may in fact be greater than that measured in the blinded RCTs.

In this analysis, PrEP efficacy is assumed to be 85% in same-sex relationships and 65% in heterosexual relationships, based on the emerging evidence of higher adherence in field settings compared to randomized trial settings. Although it may be more realistic to model a difference in PrEP efficacy based on the type of sex act (i.e. higher in the context of receptive anal intercourse), there is a lack of data on the relative efficacy of PrEP in the context of insertive vaginal or anal intercourse. The limited data from randomized trials in heterosexuals suggest that PrEP has similar efficacy in heterosexual males and females [385, 386], but confidence intervals around these estimates are too wide to permit firm conclusions. In the absence of better data, we therefore assume that PrEP efficacy depends on the type of relationship (same-sex or heterosexual) but not the type of sex act.

#### 5.5.2 PrEP uptake

Few studies have investigated the acceptability of PrEP among high risk groups. In a study of high risk groups in four countries (Kenya, India, Peru and Ukraine), Eisingerich *et al* [387] found that more than 90% of individuals reported that they would probably or definitely use PrEP if it was available. In another Kenyan study, 80% of sex workers and MSM reported that they would use PrEP if it was found to be effective [388]. However, stated acceptability may differ from actual uptake. In a South African PrEP demonstration project, only 219 out of 351 HIV-negative sex workers started PrEP, and only 117 returned for their first visit 1 month after baseline [383], suggesting an effective adoption rate of 33% (117/351). By the end of December 2017, approximately 3000 sex workers had started PrEP in South Africa, 1.5 years after the South African sex worker PrEP programme was introduced (Hasine Subedar, personal communication). This is approximately 8% of the Thembisa estimate of the number of HIV-negative sex workers in South Africa. We assume that from mid-2016 the mean annual rate at which sex workers adopt PrEP is 0.07.

There is limited local data on PrEP uptake among MSM, as this only became policy in April of 2017. To be consistent with the assumptions made about PrEP uptake in sex workers, we set the assumed annual rate of PrEP uptake in MSM to be 0.07 per annum from July of 2017.

#### 5.5.3 PrEP discontinuation

Rates at which individuals discontinue PrEP are highly variable between studies, ranging from rates of 0.23 per annum in American MSM [389] to rates of 0.45 and 0.80 per annum in studies that have followed individuals after the completion of randomized controlled trials of PrEP [376, 390]. Many of these discontinuations are due to side effects, changes in risk perception, difficulty in adhering to the medication and relocation [376, 389]. In a South African PrEP demonstration project in sex workers, it was found that although there was high dropout in the first month after starting PrEP (only 53% of women returned for the month 1 visit), the fraction of those retained at 12 months (of those who attended the month 1 visit) was 42% [383]. Our model assumes that individuals would stop PrEP if they cease to meet the PrEP eligibility criteria (for example, if they cease to engage in sexual risk behaviour), but also allows for PrEP discontinuation for other reasons, such as side effects and relocation. Individuals who start PrEP are assumed to discontinue PrEP for these other reasons at a rate of 0.33 per annum, which is towards the lower end of the range of discontinuation rates noted previously because discontinuations due to changes in risk behaviour are accounted for separately.

#### 5.5.4 Risk compensation

Although data from randomized trials generally do not show evidence of risk compensation in PrEP recipients [381, 385, 386], it is difficult to extrapolate from the data collected in these randomized trials, as trial participants would have been counselled on the uncertainty regarding the efficacy of PrEP, and even if they believed PrEP to be effective, would not have known whether they were receiving the study drug or the placebo. In a recent analysis of changes in behaviour after the unblinding of the Partners PrEP trial data, a statistically significant 10% increase was noted in unprotected extramarital sex, amongst individuals who were receiving open-label PrEP [390]. Another microbicide acceptability study found that women were resistant to the idea of using both condoms and microbicides simultaneously [391]. This suggests that some reduction in condom use could occur. However, in a study of MSM and transgender women who were offered PrEP following news of its efficacy, unprotected anal intercourse declined similarly over the course of the study in those who chose to receive PrEP and those who did not take PrEP [376]. Given the uncertainty regarding the extent of risk compensation, we have assumed zero condom migration due to PrEP in this analysis.

#### 5.5.5 HIV testing and PrEP

Because of the risk of HIV drug resistance developing if HIV-positive individuals take PrEP, it is important that individuals who wish to use PrEP be screened for HIV prior to starting PrEP and at regular intervals after starting PrEP. In our model, we therefore assume that any individual who wants to take PrEP first gets tested for HIV. If they test positive, we assume the same behaviour changes associated with HIV diagnosis as described in section 5.2.2. Similarly, if an individual acquires HIV while on PrEP, they are assumed to be diagnosed positive at their next clinic visit, with associated changes in behaviour. HIV testing in PrEP recipients is assumed to occur on average three times per annum, although local guidelines recommend four tests per annum after the baseline assessment [392, 393]. Part of the benefit of PrEP thus lies in the more rapid diagnosis of people who acquire HIV, and the associated reductions in sexual risk behaviour.

## 6. HIV disease progression

When individuals become infected with HIV, they are assumed to progress through an acute phase of HIV infection, which lasts for an average of three months. During this time, individuals are assumed to be highly infectious, but their infection is assumed not to be detected on rapid antibody tests. After progression to the chronic phase of infection, the model then simulates individual-level changes in viral load and CD4 count over the course of HIV infection. This chapter focuses mainly on the modelling of untreated HIV infection in adults. Although mortality rates after ART initiation are modelled (as described in section 6.3), the effect of ART on CD4 and viral load is not yet modelled, i.e. the model records only baseline CD4 count and viral load at ART initiation. CD4 and viral load changes in HIV-positive children are also not modelled.

### 6.1 Model of changes in viral load

At the point of HIV acquisition, all adults are assigned a set point viral load (SPVL), which represents the initial viral load at the time when they progress from acute HIV infection to the chronic phase of HIV infection. The changes in viral load that occur during the acute stage of HIV infection are not modelled, as these are difficult to characterize. The SPVL is measured on a log10 scale, and is randomly sampled from a normal distribution. The mean of this distribution is 4.17 in women (based on a review of viral load distributions in early disease in East Africa and Southern Africa [394]) and 4.36 in men (studies suggest that levels of viral load in early infection tend to be higher in men than in women [395], and the difference of 0.19 is the lower confidence limit of the difference estimated in an analysis of viral load data from across Europe [396]). The standard deviation for the normal distribution is set at the same value (0.9) for males and females, based on the reported inter-quartile range in a study of Kenyan sex workers [397], after correcting for variation due to measurement error (assumed to be 0.25^2^ on the log10 scale [398]).

The viral load is updated deterministically, at weekly time steps, according to the following formula

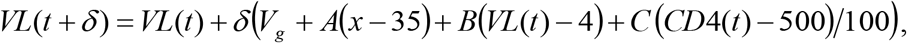

where *VL*(*t*) is the viral load at time *t, V_g_* is the average annual increase in viral load for individuals of sex *g, A* is the effect of age (*x*) on the average annual change in viral load, *B* is the effect of viral load on the change in viral load, *C* is the effect of CD4 count on the change in viral load, and *CD*4(*t*) is the CD4 count at time *t* (discussed below). This model is based loosely on the model of viral load changes fitted by Nakagawa *et al* [399] to data from Europe, with some adaptations to improve the fit to South Africa data sources. Parameters taken directly from the Nakagawa study include the effect of age (*A* = 0.0022), the effect of viral load (*B* = −0.026) and the effect of CD4 count (*C* = −0.004). The baseline rate of change in men has been set at a slightly lower rate than that estimated by Nakagawa *et al* (*V*_0_ = 0.061), but the baseline rate of change in women has been set at a higher rate (*V*_1_ = 0.122), as the sex difference in the Nakagawa study (0.004) is much smaller than the sex difference observed in a US study (0.118) [400], and cross-sectional data from another US study also suggest a more rapid increase in viral load in women than in men [401]. We have therefore taken the average of the 0.004 and 0.118 differences, and added this to the 0.061 parameter for men, in order to get the *V*_1_ parameter for women. The change in the viral load over each time step (the second term on the right-hand side of the above equation) is assumed to be subject to a lower limit of zero, to prevent instances in which the viral load declines in the absence of ART.

Although changes in viral load after ART initiation are not modelled, it is assumed that if individuals interrupt ART, their viral load returns immediately to the level it was prior to ART initiation. The change in viral load at each subsequent time step is then modelled in the same way as the change in viral load in ART-naïve individuals.

### 6.2 Model of changes in CD4 count

At the time of progression from acute HIV infection to chronic HIV infection, each HIV-positive adult is randomly assigned a CD4 count. The CD4 count is modelled on a natural log scale, and the natural log of the initial CD4 count is drawn from a normal distribution. South African data sources are inconsistent regarding the initial values of the CD4 count. Data from urban Gauteng suggest a median CD4 count in HIV-negative adults of 1128 cells/μl [402], but data from rural KwaZulu-Natal suggest much lower levels (683 in men and 833 in women) [403]. There is assumed to be a 22.4% reduction in CD4 count from prior to HIV acquisition to the time of entry into chronic HIV infection [404]. Using the data from the KwaZulu-Natal study, we might estimate normal means of 6.27 and 6.47 for males and females respectively (e.g. 6.27 = ln(683×(1-0.224)) in men). However, if we were instead to use the data from the Gauteng study, we might estimate means of 6.61 and 6.81 in men and women respectively, assuming the same gender difference as observed in the KwaZulu-Natal study. We have used the average of the values from the two studies to obtain a mean of 6.44 for men. For women we have assumed a mean of 6.54, which is 0.1 less than the average of the two studies – this slight reduction in the gender differential is incorporated as the gender difference of 0.2 estimated from the KwaZulu-Natal data is greater than that estimated in other studies in HIV-negative adults [405–407]. The standard deviation of the normal distribution from which the initial CD4 counts are sampled is set at 0.219 in men and 0.224 in women, based on the inter-quartile ranges of the CD4 distributions observed in HIV-negative adults in KwaZulu-Natal (after correcting for variation due to measurement error, assumed to be 0.25^2^ on the natural log scale [398]).

The CD4 count is updated deterministically, at weekly time steps, according to the following formula

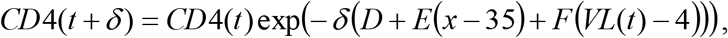

where *D* is the average reduction in CD4 count per annum (on the natural log scale), *E* is the effect of age on the change in CD4 count, and *F* is the effect of viral load (on the log_10_ scale) on the change in CD4 count. This model is adapted from the model fitted by Nakagawa *et al* to longitudinal CD4 data collected from ART-naïve patients in Europe [399]. The model of Nakagawa *et al* was applied only to patients with CD4 counts above 200 cells/μl, and the rates of change in CD4 count were measured on a normal scale (not a natural log scale), which made it inappropriate for modelling CD4 changes below the 200 cells/μl threshold. We have instead expressed our CD4 change model on a natural log scale, which means that a coefficient of *β* estimated in the Nakagawa study would need to be transformed as −ln(1 + *β*/500) before it could be entered in our model (the CD4 count of 500 is the baseline defined in the Nakagawa model). Using this transformation, we obtain an estimate of the *F* parameter of 0.078.

Using the same approach to estimate the *D* parameter yields an estimate of 0.148; this estimate changes to 0.124 if the baseline changes to a CD4 count of 300. We have set the value of the *D* parameter at 0.139.

Using the same approach to estimate the *E* parameter yields an estimate of 0.00034. However, using data from the same European collaboration, Lodi *et al* [408] estimated a more substantial effect of age on the rate of CD4 decline, with each year of increase in age accounting for a roughly 0.01 greater annual decline in CD4 count, on the square root scale (on the natural log scale, this is roughly equivalent to an age effect of 0.0013). Although the Lodi model is not standardized on viral load, in the way that the Nakagawa model is, it yields estimates of the age effect more consistent with the age differences observed in practice (discussed below), and we have therefore set *E* to be 0.0013 rather than 0.00034.

The rate of change in CD4 count is assumed not to depend on sex, as evidence is conflicting. Some studies suggest a significantly faster CD4 decline in women than in men [401, 409], but others suggest the opposite [371], and most of these analyses do not control for confounding factors that could explain the reported sex differential. In the Nakagawa model, sex was found to have no significant effect on the rate of CD4 decline.

The factor by which the CD4 count changes over each time step (the exp(…) term in the above equation) is subject to an upper limit of 1, to prevent instances in which the CD4 count might increase in the absence of ART.

Although changes in CD4 count after ART initiation are not modelled, it is assumed that if individuals interrupt ART, their CD4 count returns immediately to the level it was prior to ART initiation. The change in CD4 count at each subsequent time step is then modelled in the same way as the change in CD4 count in ART-naïve individuals. This is roughly consistent with data from structured treatment interruption trials, which show that the CD4 count drops rapidly during the first 3 months after an ART interruption, then declines at a more steady rate thereafter [410–412].

### 6.3 Model of mortality as a function of CD4 count

Unlike the models of change in CD4 count and viral load, which are deterministic, the model of HIV-related mortality is stochastic. The annual mortality rate in untreated adults is assumed to be a function of the individual’s current CD4 count. Mathematically, the annual HIV-related mortality rate at time *t* is

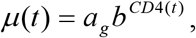

where *a_g_* is the theoretical maximum HIV mortality rate that an individual of sex *g* would experience if they had a CD4 count of zero, and *b* is the factor by which the mortality risk reduces per unit increase in the CD4 count. This model has been proposed previously [256], and in this analysis we use the same parameters for *b* (0.9886) and for a0 in males (0.645) as estimated previously. However, there is some evidence to suggest that women experience a higher mortality rate than men when CD4 count is controlled for [413], and we therefore use a value of *a*_1_ that is 10% higher than that in males (i.e. 0.710).

In individuals who are ART-experienced, HIV-related mortality is assumed to be a function of the individual’s baseline CD4 count and the time since ART initiation, with this function following a Makeham form [414]. If *x* is the individual’s CD4 count at the time they started ART and *d* is the length of time (in years) since ART initiation, the annual HIV-related mortality rate is assumed to be

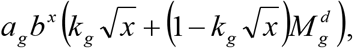

where *k_g_* and *M_g_* are factors controlling the extent of the reduction in HIV-related mortality per year after ART initiation, in individuals of sex *g*. When *d* = 0, the HIV-related mortality is the same as in untreated individuals. For *M_g_* < 1, mortality decreases as the duration of ART increases. However, the inclusion of the *k_g_* term ensures that the decline does not follow a pattern of exponential decay, since the decline in mortality appears to be more rapid at early ART durations than at longer durations. The inclusion of the 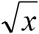 term further ensures that the decline is more modest (in relative terms) for patients starting ART at higher baseline CD4 values. The parameters *k_g_* and *M_g_* have been set to 0.0105 and 0.117 respectively in men, and to 0.0079 and 0.049 respectively in women. These values were chosen to yield the best fit to HIV-specific mortality estimates at different ART durations in a recent analysis of data from eight South African ART cohorts [415]. The correspondence between the empirically-derived mortality estimates and the model are presented in Figure 6.3.1.

**Figure 6.3.1:**
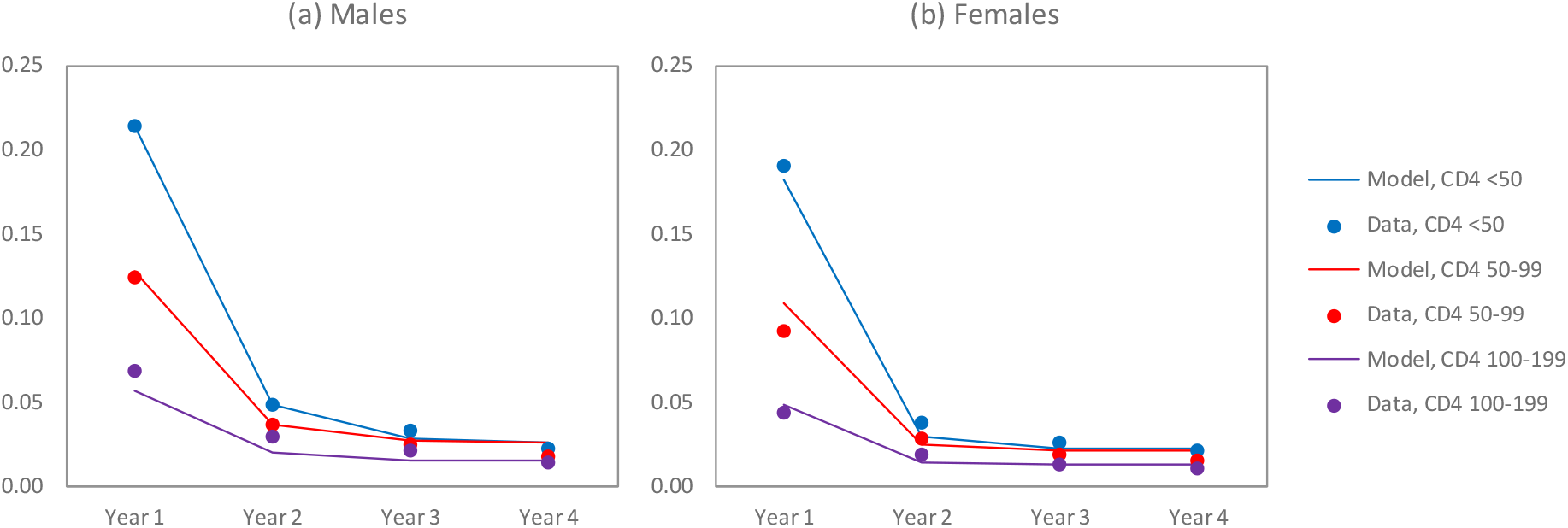
Annual HIV-related mortality rates, by time since ART initiation. For the purpose of comparison, empirically-derived estimates over the <50, 50-99 and 100-199 CD4 ranges are compared to model estimates for baseline CD4 values of 25, 75 and 150 respectively. Model parameters were estimated by minimizing the sum of squared differences between model estimates and empirically-derived estimates.

### 6.4 Baseline values at age 15

Although we are not modelling viral load and CD4 changes that occur during childhood, it is necessary to assign a baseline value that applies if children survive to age 15 in the absence of ART. As there are relatively few such children, we do not consider the complexities of allowing for variation between individuals, but assign to all children the same baseline value at age 15: a log viral load of 4.8 and a CD4 count of 300. The former is the average baseline viral load prior to ART initiation in a cohort of South African adolescents [416], and the latter is based on CD4 distributions in HIV-positive Zimbabwean adolescents (median age 14) who were believed to have acquired HIV through mother-to-child transmission [417]. After reaching age 15, changes in CD4 and viral load are modelled relative to this baseline level, in the same way as in adults.

### 6.5 Model validation

Figure 6.5.1 compares the model estimates of the fraction of the HIV-positive adult population in four different CD4 categories with cross-sectional data from three surveys conducted in the period up to 2005 [402, 418, 419]. (More recent surveys are not included as the CD4 distributions in more recent surveys will have been influenced by ART rollout, and the objective of this exercise is not to validate the assumptions about ART rollout or CD4 recovery after ART initiation.) Although none of the surveys are nationally representative, the model estimates are roughly consistent with the results from these surveys. This suggests that the overall patterns of CD4 decline in the model are realistic.

**Figure 6.5.1:**
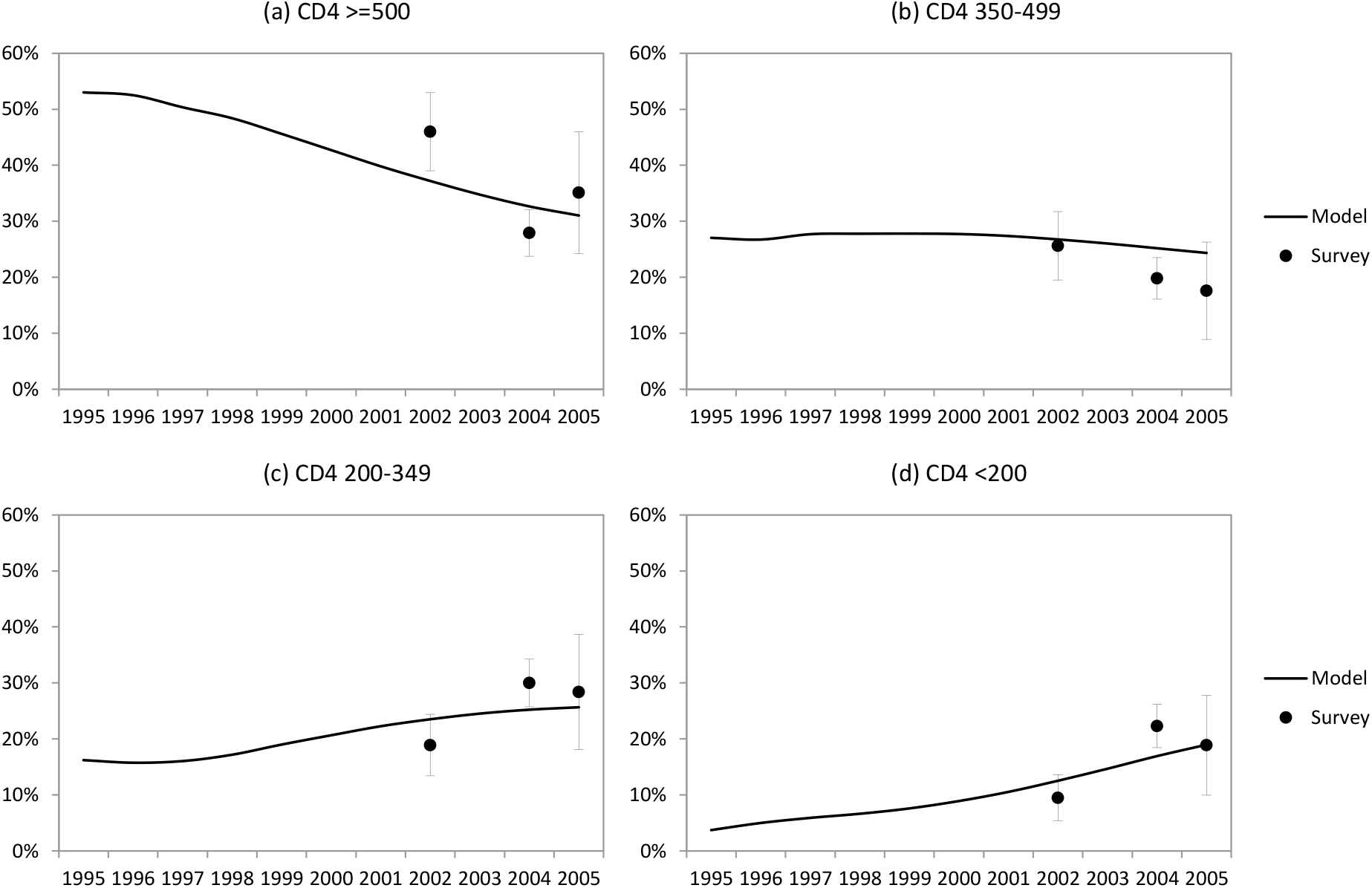
Proportion of HIV-positive adults in different CD4 categories. Model estimates are calculated as the average from 100 simulations, assuming no ART rollout. Vertical lines represent 95% confidence intervals around survey estimates.

To further validate the model, we examined the HIV survival times of all adults who acquired HIV over the course of three model simulations (in each model simulation we projected the change in the epidemic up to the middle of 2010, assuming no ART rollout). In these three simulations a total of 13 999 new HIV infections occurred in adults. Figure 6.5.2 shows the survival rates in men, compared with the published survival rates in a South African study of HIV-positive goldminers, conducted prior to the availability of ART [420]. Results are reasonably consistent, although the model slightly under-estimates the fraction of goldminers surviving at 11 years after HIV acquisition.

**Figure 6.5.2:**
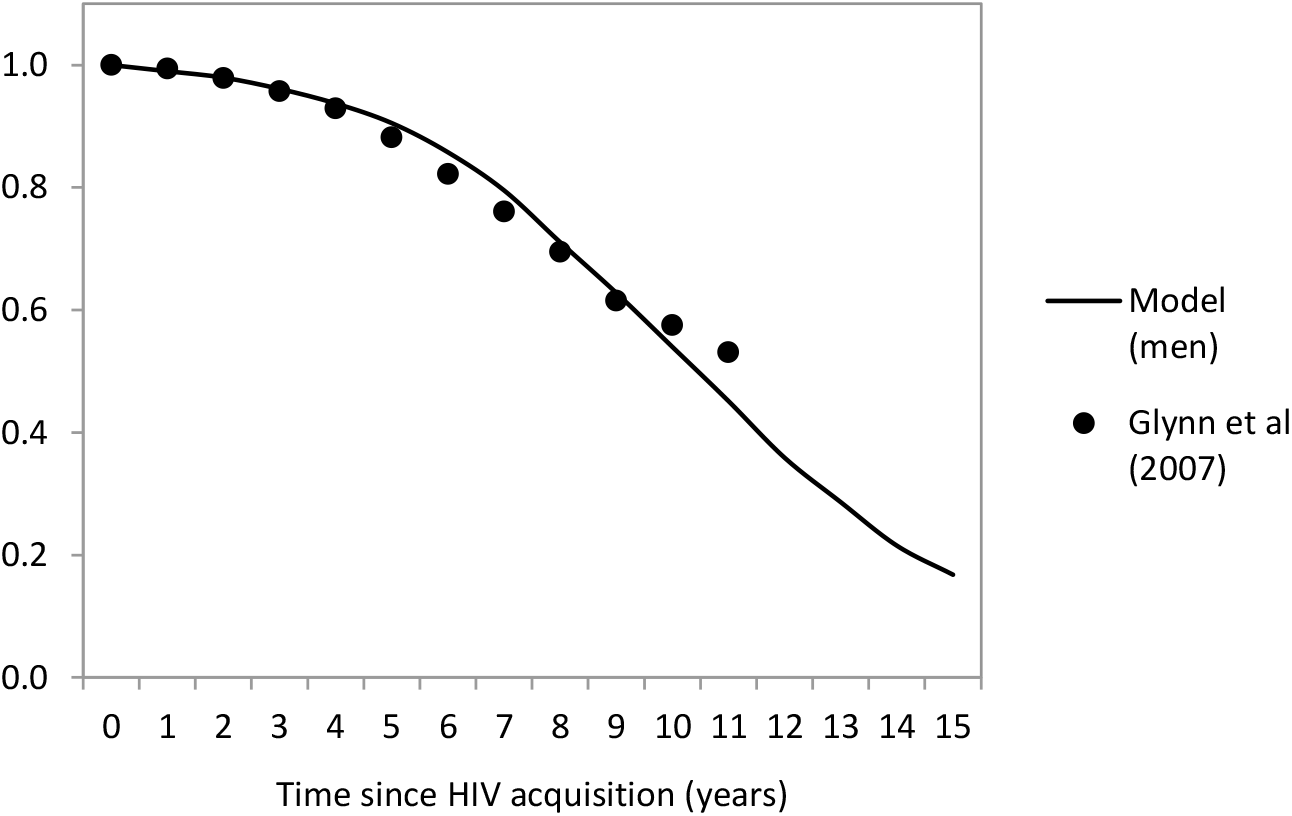
Survival rates in HIV-positive men in the absence of ART. Empirical estimates are taken from Glynn *et al* [420], after restricting the analysis to South African men, and after standardizing to ages 25-29 at seroconversion. Model estimates are obtained by calculating Kaplan-Meier curves from the simulated event histories generated for 5739 HIV-positive men.

From the same three simulations, we calculated the distribution of viral loads in all HIV-positive adults in 2005 (*n* = 7522). The median viral load (4.6) and interquartile range (4.0-5.1) was consistent with the results of a South African community survey conducted in 2002 (median 4.7, IQR: 4.0-5.2) [402]. The simulated distribution of viral loads within the subset of HIV-positive adults with CD4 counts <200 cells/μl (median 5.2, IQR: 4.8-5.6) was also consistent with that observed in the survey (median 5.2, IQR: 4.9-5.6). This suggests that the model assumptions about the baseline levels of viral load and rates of increase in viral load are plausible.

Studies have generally shown that after controlling for age, sex has relatively little influence on overall mortality rates. Two of the largest studies that have evaluated the effect of sex on HIV mortality report female-to-male hazard ratios of 0.91 (95% CI: 0.71-1.16) in a pooled analysis of data from developing countries [421] and 0.89 (95% CI: 0.74-1.07) in a pooled analysis of data from high income countries [422]. When a Cox proportional hazards model was fitted to the 13 999 simulated patient histories, controlling for age, the female-to-male hazard ratio was 0.88 (95% CI: 0.83-0.94), consistent the hazard ratios estimated in the literature.

Several studies have shown that age at seroconversion is positively associated with mortality. The increase in mortality, per 10-year increase in age, has been estimated at 1.20 (95% CI: 0.89-1.63) in Zimbabwe [423], 1.35 (95% CI: 1.25-1.46) in a pooled analysis of MSM survival data from high-income countries [422], 1.37 in Canada [424], 1.55 (95% CI: 1.31-1.83) in Italy [425] and 1.72 in Tanzania [426]. When a Cox proportional hazards model was fitted to the 13 999 simulated patient histories, each 10-year increase in age was associated with a hazard ratio of 1.34 (95% CI: 1.30-1.38), which is within the range of the empirical estimates quoted.

Several studies have shown that SPVL is strongly associated with the mortality rate and the rate of progression to AIDS. The factor by which the rate of mortality increases, per unit increase in the log of the SPVL, has been estimated at 1.8 (95% CI: 0.9-3.2) in the Gambia [427], 2.21 (95% CI: 1.36-3.59) in Kenya [397] and 2.66 (95% CI: 1.54-4.62) in the US [428]. When a Cox proportional hazards model was fitted to the 13 999 simulated patient histories, the increase in mortality per unit increase in the log of the SPVL was 1.98 (95% CI: 1.91-2.05), consistent with these empirical estimates. This suggests that the model assumptions about the effect of viral load on rates of CD4 decline are plausible.

## 7. HIV transmission

### 7.1 Baseline transmission probabilities per sex act

HIV transmission is modelled based on the assumption of a probability of HIV transmission per act of sex. This probability differs depending on the viral load and stage of HIV disease of the HIV-positive partner, as well as whether the female partner is using hormonal contraception, whether the male partner is circumcised, whether the HIV-positive partner is on antiretroviral treatment (ART), and whether either partner has a sexually transmitted infection (STI). The ‘base’ HIV transmission probabilities specified below are assumed to apply when

- The HIV-positive partner is untreated, has a viral load of 4.5 log_10_ copies/ml and a CD4 count ≥200 cells/μl, and is not in the acute stage of HIV infection;
- The male partner is uncircumcised;
- The female partner is not using hormonal contraception and is not pregnant;
- No condom or other HIV prevention method (e.g. pre-exposure prophylaxis) is used;
- Neither partner has an STI; and
- The female partner is aged 25 or older.

In addition, the base transmission probabilities are assumed to depend on the type of relationship (short-term, long-term or sex worker-client relationship). This assumption is made because HIV transmission probabilities per sex act tend to be lower in individuals who have been exposed to HIV-positive partners for longer durations than in individuals who have only recently been exposed to HIV-positive partners [429–431]. Model simulations suggest that this is explained by heterogeneity in HIV susceptibility and infectivity [432], and most of these sources of heterogeneity are already accounted for in our model. However, to the extent that there are sources of heterogeneity that we have not modelled (such as HLA class I discordance [433, 434] and acquired immunity [435, 436]), it remains important to allow for differences in transmission probabilities by relationship type, as a crude approach to capturing the effect of unmodeled heterogeneity in HIV transmission risk.

Setting the base HIV transmission probabilities is challenging because most empirical estimates of HIV transmission probabilities aggregate data from several different covariate categories. Asymptomatic STIs are very common, and this makes it difficult to obtain estimates of the HIV transmission probabilities that would apply in the absence of STIs. In a meta-analysis of studies conducted in the pre-ART era, Boily *et al* [437] estimated the average male-to-female transmission probability per sex act with an HIV-positive partner to be 0.0008 in high-income countries and 0.0030 in low-income countries. These estimates could be over-estimates of the transmission probabilities that would be expected in the absence of STIs, but they could also be under-estimates of the transmission probabilities that would be expected in short-term relationships, since longitudinal studies of serodiscordant couples tend to be biased towards individuals in more stable relationships. To represent the uncertainty around the male-to-female transmission probability per sex act in short-term relationships, we assign a beta prior with a mean of 0.0025 and a standard deviation of 0.0005. In the same meta-analysis, Boily *et al* estimated the female-to-male transmission probability per sex act with an HIV-positive partner to be 0.0004 in high-income countries and 0.0038 in low-income countries. As before, this could be an over-estimate or an under-estimate, but because female-to-male transmission of HIV is generally considered to be less efficient than male-to-female transmission, we represent the uncertainty around the female-to-male transmission probability using a beta prior with a lower mean (0.00125) than assumed for male-to-female transmission and a standard deviation of 0.00025.

As noted before, model simulations suggest that heterogeneity in HIV transmissibility and susceptibility imply a lower HIV transmission risk in individuals who have been cumulatively exposed to an HIV-positive partner many times than is the case for an individual who is newly exposed to an HIV-positive partner [432]. This suggests that the transmission probability should be lower in long-term relationships than in short-term relationships, but only to the extent that there are important sources of heterogeneity in infectivity/susceptibility that are not captured in our model. As it is difficult to quantify the significance of these factors, we assign a uniform prior to represent the uncertainty around the ratio of the HIV transmission probability per sex act in long-term relationships to that in short-term relationships (a separate prior is assigned for males and females). Recognizing that sex workers also have high levels of cumulative HIV exposure, we also assign a separate prior to the male-to-female transmission probability in sex worker-client contacts: this prior distribution has a mean of 0.0010 and a standard deviation of 0.0003. This is supported by client-to-sex worker transmission probabilities in Kenya and Senegal of around 0.0003-0.0006 [438, 439], substantially lower than the average of 0.0030 estimated in a meta-analysis of the male-to-female transmission probabilities in low-income settings, which did not involve commercial sex exposure [437].

In the case of male-to-male transmission of HIV, there are relatively few studies to inform estimates, and all are from high-income settings. In a systematic review of these studies, Baggaley *et al* [440] estimated the average transmission probability, per act of unprotected receptive anal intercourse, to be 0.014 (95% CI: 0.002-0.025). This is probably an over-estimate of the transmission probability that would be expected in the absence of STIs, but it could be an under-estimate if the data are biased towards MSM in stable relationships, or if transmission probabilities are substantially higher in South Africa than in high-income countries. The uncertainty around the transmission probability in short-term relationships is represented by a beta distribution with a mean of 0.0050 and a standard deviation of 0.0010. As in the case of heterosexual transmission, the uncertainty regarding the relative risk of transmission per sex act in long-term relationships (relative to short-term relationships) is represented by a uniform (0, 1) prior distribution. Studies suggest that the probability of transmission per act of insertive anal intercourse is roughly a quarter of that in receptive anal intercourse [441, 442]. We therefore assign priors only to the transmission probabilities per act of receptive anal intercourse, and assume the insertive transmission probabilities are one quarter of the receptive probabilities.

### 7.2 Effect of viral load, HIV stage and ART

The average transmission probability per act of unprotected sex, which is specified as an input parameter, is assumed to apply when the HIV-positive partner is untreated and has a log viral load of 4.5 copies/ml. The choice of 4.5 as the log viral load to which the average applies is based on the observation that the median viral load in untreated adults in 2005 was 4.6 (see section 6.5).

For HIV-positive adults who are not in the acute stage of HIV infection and who are not yet on ART, the probability that they transmit HIV in an act of unprotected sex is assumed to be a function of their log viral load, *x*. The factor by which the average transmission probability is multiplied, to take into account the individual’s viral load, is

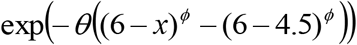

for *x* ≤ 6 and exp(*θ*×1.5^*ϕ*^) for *x* > 6. This is based on the Thembisa model of HIV in South Africa [443], in which the *θ* and *ϕ* parameters were set at 0.432 and 1.5 respectively. We have used the same parameters as proposed in this model. Using a *ϕ* parameter greater than one ensures that the although infectivity is an increasing function of viral load, the relative increases in infectivity become smaller with each unit increase in viral load, until infectivity becomes constant with respect to viral load above 6 log copies/ml [444]. Although the assumption of constant infectivity at viral loads above 6 log copies/ml may seem artificial, empirical data suggest that the assumption is reasonable [445]. The value of *θ* was set at 0.432 to ensure that at viral load levels of 4 log10 copies per ml, each unit increase in log viral load leads to a 2.5-fold increase in infectivity, in line with data from various studies of serodiscordant couples [446–448] (the average of 4 is consistent with the average observed in these serodiscordant couple studies).

For adults in the acute stage of HIV infection, there is an increased risk of HIV transmission that is independent of the viral load [449]. Rather than attempt to model the viral load in the acute infection phase, we assume that average infectivity in this stage is some multiple of the infectivity that applies after the individual has progressed to the chronic phase of HIV infection. Mathematically, the adjustment to the average transmission probability is

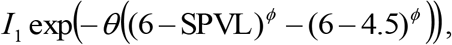

where *I*_1_ is the ratio of acute infectivity to infectivity immediately after progression to chronic infection, and SPVL is the individual’s set point viral load (the viral load immediately after the individual has progressed from the acute phase to the chronic phase). There is substantial uncertainty regarding the true value of the *I*_1_ parameter, and we have therefore assigned a prior distribution to represent the uncertainty around this parameter. We have chosen a gamma prior with a mean of 16 and a standard deviation of 6, based on previous studies that have estimated the relative increase in infectivity during acute HIV infection [449–452]. This is the same prior as assumed previously when the microsimulation model was fitted to South African HIV prevalence data [18].

There is also some evidence to suggest that at CD4 counts of <200 cells/μl or in advanced HIV disease, HIV infectivity may increase, independently of viral load (i.e. similar to the effect seen in acute infection). Wawer *et al* [450] found that HIV transmission rates in discordant couples were increased by a factor of 3.49 (95% CI: 1.76-6.92) in the 6-25 months before the death of the HIV-positive partner, after adjusting for the HIV-positive partner’s viral load. In another study of discordant couples in East Africa, Donnell *et al* [453] found that the HIV transmission risk was 4.98 times greater when the HIV-positive partner’s CD4 count was <200 cells/μl than when it was ≥350 cells/μl. Although the authors did not report the adjusted effect after controlling for viral load, a pooled analysis of viral load data from the East African region estimated an average difference in log viral load of approximately 0.9 when comparing patients with CD4 counts <200 cells/μl and those with CD4 counts ≥350 cells/μl [394]. If HIV infectivity increases by a factor of 2.5 per unit increase in log viral load [446–448], this suggests that after controlling for viral load differences, the infectivity of partners with CD4 counts <200 cells/μl is approximately 2.2 times (4.98/(2.5^0.9^)) that of partners with CD4 counts ≥350 cells/μl. There is also evidence of heightened mother-to-child transmission risk at low CD4 counts, after controlling for viral load [454]. It is not clear why CD4 count or advanced HIV disease would influence the HIV transmission risk independently of viral load, but it is possible that reductions in CD8-associated selection pressure in late disease may allow for reversion of HIV mutants to more stable epitopes that are more transmissible [455, 456]. We model this effect by assuming that in untreated HIV-positive adults with CD4 counts of <200 cells/μl, the adjustment to the average transmission probability is

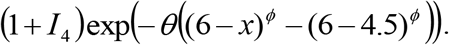

The uncertainty regarding the *I*_4_ parameter is represented by assigning a gamma prior with a mean of 1.8 and a standard deviation of 0.6. The prior mean corresponds to a multiple of 2.8, the average of the 3.49 and 2.2 multiples cited previously.

For adults who have started ART, changes in viral load levels after ART initiation are not modelled. Instead we assume that infectivity after ART initiation is some multiple (*I*_5_) of the infectivity that applied just prior to ART initiation. Expressing this mathematically, if *x* was the log viral load just prior to ART initiation, and the CD4 count prior to ART initiation was ≥200 cells/μl, the transmission probability that applies after ART initiation is the average transmission probability multiplied by a factor of

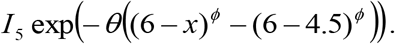

(The expression is also multiplied by (1 + I4) if the baseline CD4 count was <200 cells/μl.) The I5 parameter has been set at 0.2, which implies an 80% reduction in infectivity after ART initiation, roughly consistent with the 77% reduction in infectivity estimated in a recent study in KwaZulu-Natal [457]. The relative risk of 0.2 is somewhat higher than the relative risk estimates of 0.04-0.08 estimated from randomized controlled trials [453, 458], but lower than the relative risk of 0.36 estimated in a recent meta-analysis of observational studies [459].

### 7.3 Effect of hormonal contraception

There is evidence to suggest that the use of hormonal contraception increases the risk of HIV transmission in two ways: firstly, by increasing HIV-negative women’s susceptibility to HIV infection, and secondly, by increasing HIV-positive women’s infectiousness.

Considering firstly the risk of transmission from HIV-positive men to HIV-negative women, there are a number of biological reasons why the use of injectable contraception may increase HIV transmission risk. A study of women in KwaZulu-Natal found that the concentration of cervical HIV target cells in women using injectable contraceptives was almost 4 times that in women not using contraception [460]. It has also been noted that the use of injectable contraceptives often leads to thinning of the vaginal epithelium and increased frequency of vaginal bleeding [461], which could increase HIV transmission risk. Meta-analytic reviews have shown that the use of injectable contraception is associated with an increased HIV acquisition risk in women. In the most recent of these reviews, Polis *et al* [462] found that HIV incidence was significantly increased in users of DMPA (HR 1.40, 95% CI: 1.23-1.59) but not in users of NET-EN or in users of oral contraceptives (OC). Similar findings were reported in previous meta-analyses [463, 464]. It is important to note, however, that none of the studies included in the systematic reviews randomly allocated women to their contraceptive methods, and it is therefore possible that the observed association between DMPA use and HIV acquisition risk could be attributable to unmeasured confounding.

In South Africa, 59% of injectable contraceptive use is DMPA rather than NET-EN, although NET-EN appears to be more commonly used than DMPA among young women [36]. Since our model does not distinguish DMPA and NET-EN use, we use a weighted average of the hazard ratios estimated for these two methods (1.40 and 1.10 respectively, based on the meta-analysis of Ralph *et al* [463]), with weights of 0.59 and 0.41 respectively, to arrive at an assumed susceptibility multiplier of 1.28. No effect of OC use on HIV transmission risk is assumed.

The evidence of an effect of hormonal contraceptive use on women’s HIV infectiousness is less specific to DMPA. Mostad *et al* [465] found that the frequency of HIV-1 shedding was increased both in users of DMPA (OR 2.9, 95% CI: 1.5-5.7) and in OC users (OR 3.8, 95% CI: 1.4-9.9) in Mombasa. More dramatic effects were reported by Clemetson *et al* [466] in the case of OC, among women in Nairobi (OR 11.6, 95% CI: 1.7-77.6). On the other hand, a study among women in Mombasa found that the increase in HIV-1 shedding associated with recent initiation of OC or DMPA use was not significant [467]. More recently, in a study of HIV transmission rates in serodiscordant couples in seven African countries, Heffron *et al* [468] found that in those couples in which the female partner was HIV-positive, the female-to-male transmission rate was doubled if the woman was using hormonal contraception (aHR 1.97, 95% CI: 1.12-3.45). The effect appeared to be much the same in women using injectable methods and women using OC. Similar increases were observed in a Ugandan study of HIV-negative men whose partners were HIV-positive, although the effects of OC and DMPA were not statistically significant due to the small study size [469]. On the other hand, a recent study in Zambian serodiscordant couples found no increase in female-to-male transmission risk associated with hormonal contraceptive use (either OC or injectable methods) [470]. Pooling the results from the three studies that directly measured the effect of hormonal contraception on female-to-male contraception gives a pooled relative risk of 1.10 (95% CI: 0.67-1.81), with marginally stronger evidence of an effect in the case of oral contraception (1.46, 95% CI: 0.65-3.31) than in the case of DMPA (0.93, 95% CI: 0.49-1.75). Because the pooled relative risks are non-significant, we assume HIV-positive women’s use of hormonal contraception has no effect on their probability of transmitting HIV to their partners.

### 7.4 Effect of male circumcision

As described in section 5.1.3, men who are circumcised are assumed to have a 60% lower rate of HIV acquisition from HIV-positive female partners, per act of sex [264]. However, male circumcision is assumed to have no effect on male-to-female transmission of HIV [265] or male-to-male transmission of HIV (as there is only limited evidence of a protective effect in MSM [233]).

### 7.5 Effect of other sexually transmitted infections

In addition to HIV, five other STIs are modelled: genital herpes, syphilis, gonorrhoea, chlamydia and trichomoniasis. For each of these STIs, parameters are specified to determine the proportion of infections that become symptomatic, the average duration of infection in the absence of treatment, the duration of immunity following recovery, the probability of transmission per sex act and other key variables. For the purpose of the present analysis, each of these parameter values has been fixed at the mean of the 100 best-fitting parameter combinations obtained when the microsimulation model was fitted to South African STI prevalence data [18]. A more detailed description of the modelling of these other STIs is provided elsewhere [18].

In addition, the model simulates the incidence of bacterial vaginosis and vaginal candidiasis in women. Although these infections are not conventionally thought of as STIs, they can cause symptoms of vaginal discharge and have been shown to increase the probability of HIV transmission [471]. The approach to modelling these two infections is the same as described previously [18, 318]. Parameter values have been fixed at the posterior means estimated when an earlier deterministic model was fitted to South African bacterial vaginosis and vaginal candidiasis prevalence data [472].

For the purpose of modelling the effect of STIs on HIV transmission probabilities, we adopt the same approach as described in a previous modelling study [473]. Briefly, individuals with STIs are classified according to whether they are (a) experiencing a genital ulcer, (b) experiencing a discharge but not an ulcer, or (c) completely asymptomatic. The HIV transmission probabilities specified previously (which are assumed to apply to couples in which neither partner has an STI) are then multiplied by an STI cofactor. The magnitude of this STI cofactor is assumed to depend on the type of symptoms, whether the partner with the STI is HIV-negative or HIV-positive, and the sex of the partner with the STI. In the event that both partners have an STI, two cofactors are applied (one for each partner). Table 7.5.1 shows the STI cofactors that are assumed; these are based on systematic reviews of the effects of STIs on HIV transmission probabilities [471, 474, 475].

**Table 7.5.1:** Assumed STI cofactors

| Symptoms | Susceptibility cofactor<br>(STI in HIV-negative partner) |  | Infectiousness cofactor<br>(STI in HIV-positive partner) |  |
| --- | --- | --- | --- | --- |
|  | Male STI | Female STI | Male STI | Female STI |
| Ulcerative symptoms | 4.4 | 2.8 | 3.5 | 3.5 |
| Discharge/dysuria symptoms | 2.6 | 1.6 | 3.1 | 1.7 |
| Asymptomatic infection | 2.0 | 1.5 | 1.2 | 1.2 |

### 7.6 Effect of age

Data suggest that HIV transmission probabilities per act of sex are higher in young women than in older women [439, 476, 477]. Young women may be at a biologically increased risk of HIV acquisition due to the high prevalence of cervical ectopy in adolescence and young adulthood [478–480], and relatively low levels of protective lactobacilli [481]. The model makes allowance for this heightened susceptibility by assuming that the HIV transmission risk per act of sex is increased by a factor of 2.5 in girls aged 10-19 and by a factor of 1.5 in women aged 20-24 (relative to women aged 25 and older).

### 7.7 Effect of pregnancy

Several studies suggest that pregnancy increases women’s risk of HIV acquisition, even after controlling for behavioural factors. Mugo *et al* [482] found that in serodiscordant couples across Africa, male-to-female HIV transmission risk was increased by a factor of 1.71 (95% CI: 0.93-3.12) after controlling for confounding factors. Another analysis of discordant couples in Uganda [483] found a very similar effect of pregnancy on the HIV transmission risk after controlling for confounding factors (IRR 1.76, 95% CI: 0.62-4.03). A more recent study, which controlled for the number of unprotected sex acts, found that women’s susceptibility to HIV per unprotected sex act increased by a factor of 2.07 (95% CI: 0.78-5.49) during the first 13 weeks of pregnancy, 2.82 (95% CI: 1.29-6.15) at pregnancy durations >13 weeks, and 3.97 (95% CI: 1.50-10.51) during the first 24 weeks postpartum [484]. In Malawian women, the incidence of HIV during pregnancy was observed to be 2.19 times that in the postpartum period [485], though this study did not control for confounding factors and was not limited to discordant couples. Other studies have found no increase in susceptibility to HIV in pregnant women, although these studies were not conducted in discordant couples and did not control for behavioural confounders [486, 487]. Based on the results of the three studies conducted in discordant couples, we assume that the rate of male-to-female transmission of HIV is increased by a factor of 1.75 during pregnancy.

There is also strong evidence to suggest that pregnancy increases the frequency of HIV shedding in the genital tracts of HIV-positive women [466, 488], and as a result female-to-male HIV transmission risk is increased during pregnancy [482]. Mugo *et al* [482] found that after controlling for confounding factors, the female-to-male transmission risk was increased by a factor of 2.47 (95% CI: 1.26-4.85) during pregnancy. Similar or greater multiples have been observed for the effect of pregnancy on the odds of HIV shedding in the genital tract [488]. It is therefore assumed that HIV-positive pregnant women have a transmission risk 2.5 times that of HIV-positive women who are not pregnant.

### 7.8 Initial HIV prevalence

The HIV epidemic is seeded by randomly assigning an HIV status to each individual in 1990 (although the simulation begins in 1985, it is convenient to start the simulation of HIV transmission in 1990 because stochastic variation in HIV trajectories is less substantial when the HIV epidemic is initialized using a higher initial HIV prevalence). A prior distribution is assigned to represent the uncertainty around parameter *V*_0_, the initial HIV prevalence in black, high risk females aged 15-49 in 1990. Since the observed antenatal HIV prevalence in 1990 was 0.76% [489], and since HIV prevalence in the general population of women aged 15-49 tends to be lower than the antenatal prevalence [490], it is likely that the overall HIV prevalence in women aged 15-49 would not have been greater than 0.76%. Since high risk females are assumed to comprise 25% of the sexually experienced female population, and since the initial prevalence of HIV is assumed to be concentrated only in the high risk group, this implies that the prevalence of HIV in high risk females in 1990 could not have been greater than 0.76%/0.25 = 3.04%. We therefore set the prior on the *V*_0_ parameter to be uniform on the range [0.01, 0.03). This initial HIV prevalence is adjusted by a set of scaling factors to determine the initial HIV prevalence by sex and by 5-year age group, based on the relative levels of HIV prevalence in males and females in different age groups in a 1991 survey in KwaZulu-Natal [491]. Suppose that *s_g_*(*x*) represents the scaling factor for high risk individuals of age *x* and sex *g*, and that *v_g,r_*(*x*) represents the initial HIV prevalence (in 1990) in heterosexuals of sex *g* and risk group *r*, who are aged *x*. We calculate *v_g,r_*(*x*) as

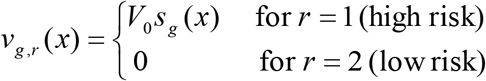

The values assumed for the *s_g_*(*x*) scaling factors, based on the KwaZulu-Natal survey, are summarized in Table 7.8.1. In the case of MSM, the initial HIV prevalence in 1990 is assumed to be 3.37 times the corresponding HIV prevalence (by age and risk group) in heterosexual men. This multiple of 3.37 is based on the 100 best-fitting parameters identified when the model was previously fitted to HIV prevalence data from South African MSM [160], and it reflects the early evidence of a severe epidemic in MSM preceding the start of the heterosexual HIV epidemic in South Africa [492, 493].

**Table 7.8.1:** Assumed ratios of initial HIV prevalence to average prevalence in females aged 15-49 (high risk group)

|  | 15-19 | 20-24 | 25-29 | 30-34 | 35-39 | 40-44 | 45-49 |
| --- | --- | --- | --- | --- | --- | --- | --- |
| Males ($g = 0$) | 0.04 | 0.44 | 1.00 | 0.69 | 0.58 | 0.48 | 0.29 |
| Females ($g = 1$) | 1.11 | 1.24 | 1.24 | 0.95 | 0.68 | 0.54 | 0.38 |

In addition, the initial HIV prevalence is adjusted by a set of race-specific scaling factors, which are determined based on the relative levels of HIV prevalence by race in the 1990 antenatal survey [494]. The race-specific adjustment factors are set to 1 for black South Africans (since the baseline *v_g,r_*(*x*) rates apply to black South Africans), 0.18 for coloured South Africans and 0.07 for white South Africans.

The above assumptions are used to calculate the expected initial number of HIV infections in 1990. The initial number of infections is then randomly allocated across the adult population using the same weighting factors, but in addition assigning zero weight to individuals living in rural areas. This means that the epidemic is seeded in urban areas, as discussed in section 3.6.6.

## 8. Model calibration procedure

In previous sections we have specified prior distributions to represent the uncertainty regarding a number of the model parameters. Table 8.1 summarizes these prior distributions. Our approach to model calibration is to

a. draw a sample of 48 000 parameter combinations from these prior distributions;
b. for each parameter combination, enter the parameters into the model, run the model and assess how well the model outputs compare with observed levels of HIV prevalence in South Africa; and
c. select the 100 parameter combinations that yield model outputs most consistent with observed HIV prevalence.

**Table 8.1:**
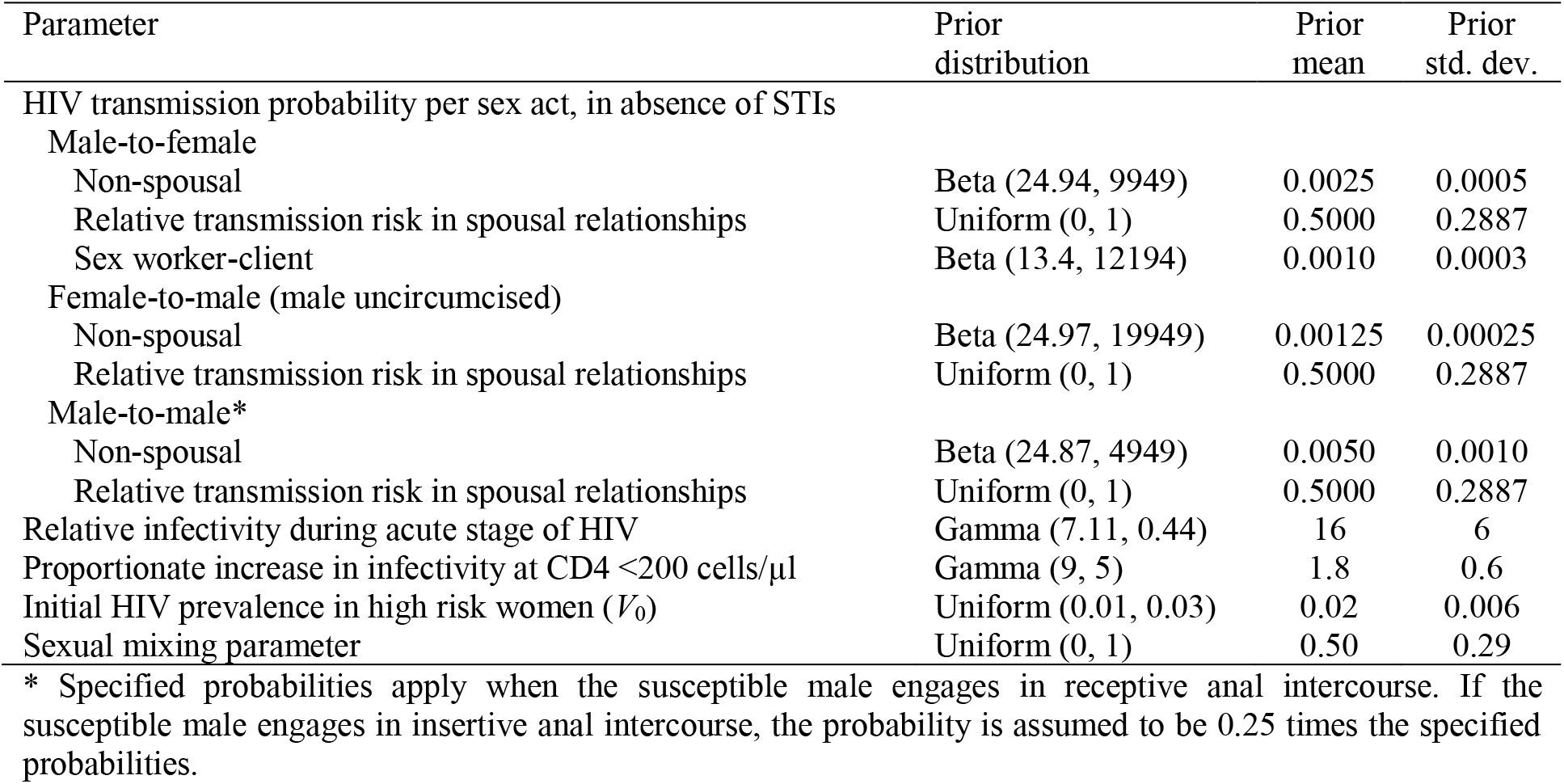
Summary of prior distributions

These 100 parameter combinations are then used in showing the model calibration to the HIV prevalence data; all results are calculated for each of the 100 parameter combinations, and the results presented are the means and 95% confidence intervals around the means (or in some cases the medians and interquartile ranges).

The measure of consistency used in steps (b) and (c) is a likelihood function. This likelihood function is calculated with respect to four HIV prevalence data sources: antenatal surveys, household surveys, surveys conducted in sex workers and surveys conducted in MSM. The sections that follow explain how the likelihood is calculated for each data source. The total likelihood is the product of the likelihood values calculated for each data source.

### 8.1 Antenatal survey data

The approach adopted in defining the likelihood for the antenatal HIV data is similar to that described previously [80], but has been updated to include more recent HIV prevalence data. Antenatal survey HIV prevalence data [495] are included for the period 1997-2015 and for 5-year age groups 15-19,…, 35-39. Suppose that *H_x,t_*(**θ**) is the model estimate of HIV prevalence in pregnant women aged *x* to *x* + 4, at the start of year *t*, where the vector **θ** represents the values of the model input parameters. The corresponding prevalence of HIV actually measured in the antenatal survey is represented by *y_x,t_*. It is assumed that if **θ** is the true set of parameter values, then the difference between the logit-transformed model estimate and the logit-transformed observed prevalence is normally distributed with zero mean. The variance of the distribution is assumed to be composed of a ‘model error’ term (representing error due to the model approximation of HIV prevalence in pregnant women), and a ‘survey error’ term (representing the uncertainty around the survey estimate due to binomial variation and cluster variation in the survey). More formally, it is assumed that

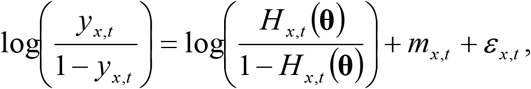

where 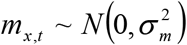 and 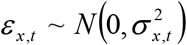. The latter two terms represent the model error and the survey error respectively. The logit transformations ensure that the error terms are closer to normality and that the model error terms are roughly independent of the level of HIV prevalence.

Antenatal data collected prior to 1997 are not included in the likelihood definition, in part because the antenatal sampling protocol prior to 1997 was not designed to be nationally representative, and in part because 95% confidence intervals were incorrectly calculated in the early antenatal surveys (not accounting for clustering of observations by clinic). The 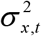 values are estimated from the 95% confidence intervals that have been published for the post-1996 survey estimates. The 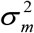 parameter is fixed at 0.005, the variance of model errors estimated when the Thembisa model was calibrated to antenatal survey data using a similar procedure [496]. The likelihood in respect of the antenatal data is calculated based on the assumption that the error terms are normally distributed:

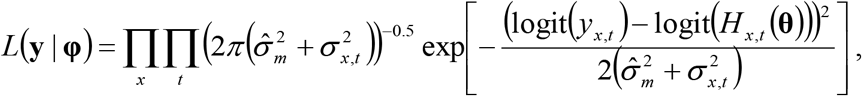

where **y** represents the matrix of *y_x,t_* values, across age bands 15-19 to 35-39, and across calendar years 1997 to 2015.

In the above equations, *H_x,t_*(**θ**) is calculated by weighting pregnant women according to their probability of being included in the antenatal survey and adjusting for likely false-positive reactions in antenatal surveys. Since the survey is only conducted in public antenatal clinics, and the racial profile of women attending public antenatal clinics is very different from that of women attending private antenatal clinics, we define *p*(*r*) to be the proportion of pregnant women of race *r* who use public antenatal clinics. Based on data from the 1998 DHS, this proportion has been set to 0.862 in black women, 0.771 in coloured women, and 0.110 in white and Asian women (Ria Laubscher, personal communication). It is also necessary to take into account that pregnant women have a greater risk of stillbirth if they are HIV-positive [497]. Since the model only simulates those pregnancies that lead to a live birth, it is necessary to weight pregnant women by the inverse of the probability of a live birth. Suppose *s*(*h*) is the probability of stillbirth in women of HIV status h. This parameter is set to 0.014 in HIV-negative women (*h* = 0), based on data from the 1998 DHS [54], and 0.056 in HIV-positive women (*h* = 1), based on a meta-analysis that estimated an odds ratio of 3.9 for the association between maternal HIV infection and stillbirth [497]. Finally, *Sp* is defined at the specificity of a single ELISA, as used in the South African antenatal surveys. This parameter has been set to 0.977, the average value estimated in a previous analysis of HIV data in South Africa [498]. Then the modelled HIV prevalence that would be expected in a survey of pregnant women attending public antenatal clinics at time *t* is

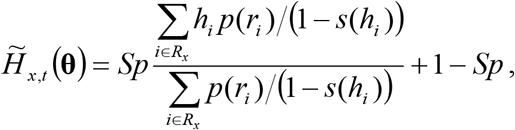

where *h_i_* is the HIV status of woman *i, r_i_* is the race of woman *i*, and *R_x_* is the set of pregnant women aged *x* to *x* + 4, at the start of year *t*. South Africa has relatively low fertility rates [54]. Because of the small number of women who are pregnant at any time, and the resulting stochastic variation in HIV prevalence, we apply a moving average to the modelled HIV prevalence so that the changes in HIV prevalence over time are relatively smooth. The ‘smoothed’ model HIV prevalence estimate is calculated as

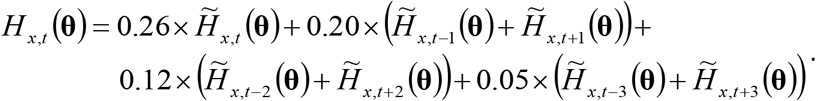

Because of the potential for stochastic variation to distort the results, and because the moving average adjustment may introduce a bias if the time trend is non-linear around time *t*, it is necessary to include some provision for model error when calculating the likelihood. This is represented by the *m_x,t_* factor described previously.

A limitation of this modelling approach is that it relies on relatively outdated data on proportions of pregnant women using public antenatal care. In addition, it does not take into account the high level of movement between public and private health services in South African pregnant women [499]. In reality there is probably greater socio-economic variation in antenatal public health sector utilization than is reflected here.

### 8.2 Household survey data

We define a likelihood in respect of the HSRC household survey HIV prevalence data [25, 68, 108] using a similar approach to that adopted for the antenatal survey data. The most important difference is that no moving average adjustment is applied to the model estimates of HIV prevalence, since the simulated population numbers in each age group are relatively large, and the stochastic variation is thus likely to be relatively small. This also means that the model error term is omitted in the variance calculation, since the model error was included in section 8.1 mainly to account for stochastic variation and the potential bias introduced by the moving average adjustment. The likelihood is calculated separately for 2005, 2008 and 2012, for males and females, and for each 5-year age band from 15-19 up to 55-59. Although a household survey was also conducted in 2002, these data have been excluded as they were obtained using a single saliva test with no confirmatory testing, and survey response rates were relatively low [109].

### 8.3 Sex worker data

Unlike the antenatal and household surveys, no HIV prevalence study in sex workers has used a nationally-representative sampling frame. The approach to defining the likelihood must therefore take into account that no individual study is nationally representative. We use a likelihood definition that includes a ‘random effect’ term to represent the difference in HIV prevalence between sex workers in the community surveyed and sex workers nationally. The statistical definition of the likelihood is described in more detail elsewhere [500]. Table 8.3.1 summarizes the HIV prevalence data that have been included in the likelihood calculation. As in section 8.1, a moving average is applied to the modelled estimate of HIV prevalence in sex workers in year *t*, in order to smooth out some of the stochastic variation in HIV prevalence, which exists because the size of the simulated sex worker population is relatively small.

**Table 8.3.1:** Studies of HIV prevalence in commercial sex workers

| Study | Location | Year | Sample size | Prevalence of HIV |
| --- | --- | --- | --- | --- |
| Ramjee *et al* [501] | Truck stops between Durban and Johannesburg | 1996 | 416 | 50% |
| Dunkle *et al* [250] | Johannesburg | 1996 | 295 | 46.4% |
| Leggett *et al* [502] | Johannesburg, Durban, Cape Town | 1998* | 249 | 42.6% |
| Williams *et al* [100] | Carletonville | 1998 | 121 | 68.6% |
| Ndhlovu *et al* [503] | Carletonville | 2001 | 101 | 78% |
| van Loggerenberg *et al* [252] | Durban | 2004 | 775 | 59.6% |
| Luseno & Wechsberg [274] | Pretoria | 2005 | 276 | 59.1% |
| Greener *et al* [504] | Durban | 2012 | 349 | 66.9% |
| UCSF [55] | Johannesburg | 2013 | 764 | 71.8% |
|  | Cape Town |  | 650 | 39.7% |
|  | Durban |  | 766 | 53.5% |

\* The study date was not stated, and has been assumed to be three years prior to the date of publication, based on average publication delays in other STI prevalence surveys [505].

### 8.4 MSM survey data

As with the sex worker data, none of the surveys conducted in MSM has used a nationally representative sampling frame, and it is therefore necessary to include a random effect term when specifying the likelihood, to represent the difference between the true HIV prevalence in MSM nationally and the prevalence that exists in the specific community being surveyed. A further complication to consider is that almost all of the HIV prevalence studies that have been conducted in South African MSM used respondent-driven sampling (RDS) to sample MSM and appropriately weight the MSM recruited into the study. Recent analysis suggests that these RDS surveys may be biased towards younger MSM, who typically have lower HIV prevalence than older MSM [160]. It is therefore necessary to adjust the modelled HIV prevalence in MSM to correspond to the age distribution in each RDS survey. A more detailed explanation of the method used to define the likelihood is provided elsewhere [160]. Table 8.4.1 summarizes the HIV prevalence survey data that are used in defining the likelihood.

**Table 8.4.1:** Studies of HIV prevalence in men who have sex with men

| Reference | Location | Year | % aged 25+ | Prevalence | Standard deviation |
| --- | --- | --- | --- | --- | --- |
| Cloete *et al* [131] | Cape Town | 2012 | 67.3% | 22.3% | 0.234 |
|  | Durban | 2012 | 27.0% | 48.2% | 0.181 |
|  | Johannesburg | 2012 | 52.1% | 26.8% | 0.196 |
| Lane *et al* [161] | Gert Sibande district | 2012 | 29.6% | 29.4% | 0.185 |
|  | Ehlanzeni district | 2012 | 28.0% | 15.9% | 0.233 |
| Lane *et al* [135] | Soweto | 2008 | 31.0% | 13.6% | 0.236 |
| Rispel *et al* [162] | Johannesburg | 2008 | 33.3% | 49.5% | 0.217 |
|  | Durban | 2008 | 33.3% | 27.5% | 0.416 |
| Sandfort *et al* [163] | Pretoria | 2011 | 41.0% | 30.1% | 0.154 |
| Kufa *et al* [506] | Johannesburg | 2015 | 54.8%* | 43.4%* | 0.133 |
|  | Bloemfontein | 2015 | 28.4%* | 17.3%* | 0.178 |
|  | Mafikeng | 2015 | 28.8%* | 14.6%* | 0.201 |
|  | Polokwane | 2015 | 40.3%* | 22.4%* | 0.196 |
| Dunkle *et al* [158] | 3 districts in Eastern Cape and KwaZulu-Natal | 2008 | 52.1% | 27.4% | 0.290 |

\* RDS-weighted estimates are not yet available (although an RDS design was used), so only the unweighted estimates are used.

## 9. Results

Table 9.1 summarizes the 100 parameter combinations that generated the best fits to the HIV prevalence data, and compares the distributions with the associated priors. For most of the parameters, the prior distribution and the distribution of best-fitting parameter values are similar. However, the best-fitting estimates of the transmission probabilities in short-term heterosexual relationships are somewhat lower than the prior medians. Conversely, the best-fitting estimates of the initial HIV prevalence level in 1990 tend to be higher than the prior median. The model also tends to fit the data better when the assumed effects of low CD4 count on the HIV transmission risk are more modest than those assumed *a priori*.

**Table 9.1:** Comparison of prior distribution and 100 best-fitting parameter values

| Parameter | Prior distribution<br>(median, IQR) | 100 best-fitting<br>parameter combinations<br>(median, IQR) |
| --- | --- | --- |
| HIV transmission probability per 1000 sex acts |  |  |
| Male-to-female |  |  |
| Non-spousal | 2.47 (2.15-2.82) | 2.14 (1.92-2.38) |
| Relative transmission risk in spousal relationships | 0.50 (0.25-0.75) | 0.62 (0.39-0.84) |
| Sex worker-client | 0.97 (0.78-1.18) | 1.02 (0.80-1.25) |
| Female-to-male (male uncircumcised) |  |  |
| Non-spousal | 1.23 (1.04-1.44) | 1.07 (0.93-1.22) |
| Relative transmission risk in spousal relationships | 0.50 (0.25-0.75) | 0.51 (0.28-0.76) |
| Male-to-male* |  |  |
| Non-spousal | 4.93 (4.29-5.63) | 4.94 (4.33-5.43) |
| Relative transmission risk in spousal relationships | 0.50 (0.25-0.75) | 0.53 (0.26-0.75) |
| Relative infectivity during acute stage of HIV | 15.3 (11.7, 19.5) | 16.3 (13.7-19.3) |
| Proportionate increase in infectivity at CD4 <200 cells/μl | 1.74 (1.37-2.16) | 1.52 (1.24-1.95) |
| Initial HIV prevalence in high risk women ($V_0$) | 2.00% (1.50-2.50%) | 2.33% (1.75-2.66%) |
| Sexual mixing parameter | 0.50 (0.25-0.75) | 0.53 (0.34-0.73) |

Figure 9.1 shows the model calibration to the antenatal HIV prevalence data. Although the model fits the overall antenatal HIV trend reasonably well, the model tends to slightly under-estimate HIV prevalence in women aged 35-39. Antenatal survey results prior to 1997 survey were not used in defining the likelihood function, but are shown for validation purposes: the model tends to over-estimate the pre-1997 prevalence at the young ages (15-24), but under-estimates the pre-1997 prevalence in the 30-39 age group.

**Figure 9.1:**
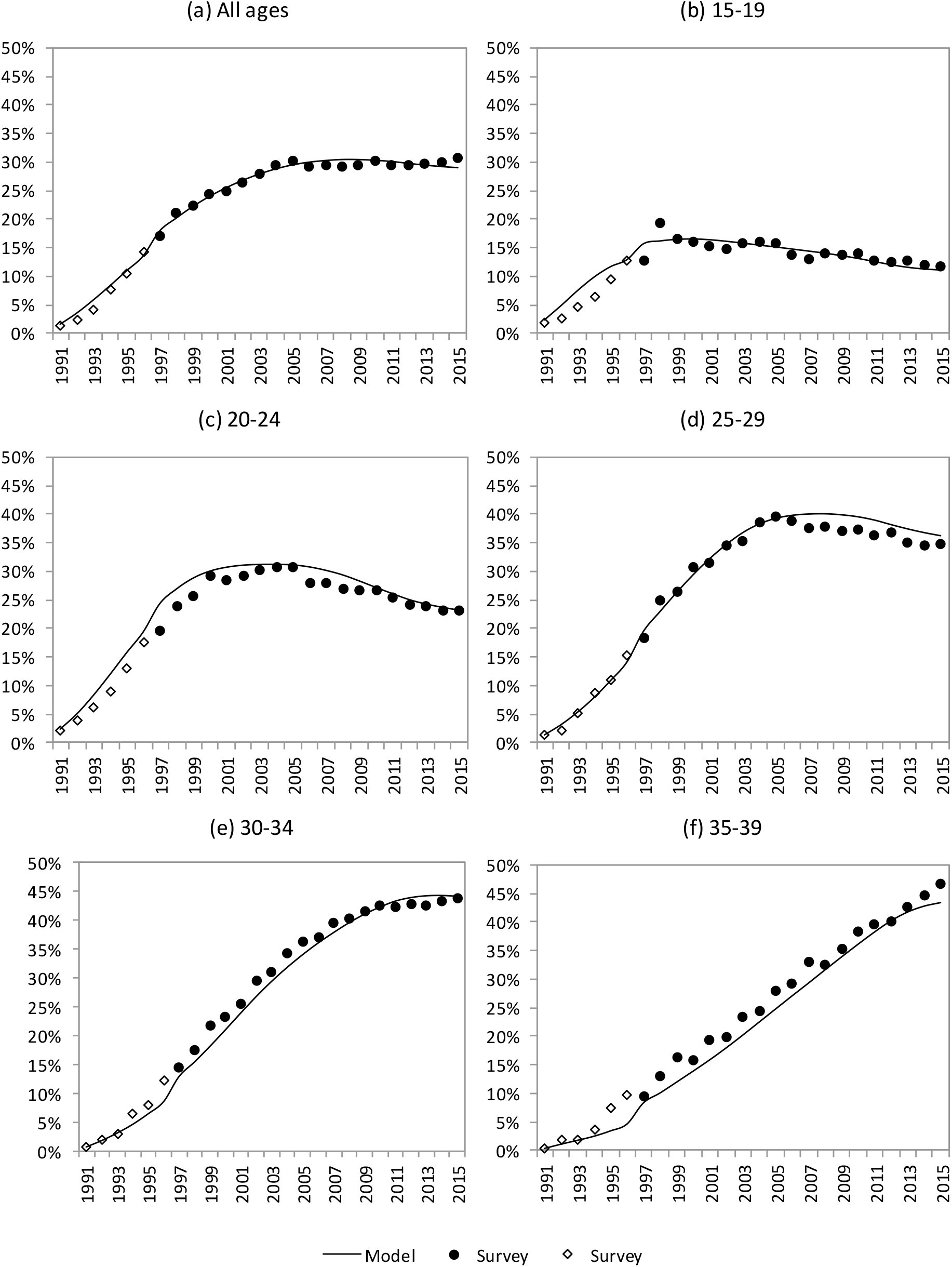
HIV prevalence in pregnant women attending public antenatal clinics. Model results have been adjusted to reflect the antenatal biases described previously (note that the antenatal bias changes in 1997 due to changes in antenatal testing protocols; surveys in 1997 and subsequent years did not include confirmatory testing and some exaggeration due to false positive test results is therefore expected). Model results are the average results generated using the 100 best-fitting parameter combinations.

Figure 9.2 compares the model estimates of HIV prevalence in the general population with those observed in the HSRC household surveys. The model provides a reasonable fit to the data. It is worth noting that the model tends to estimate a higher prevalence among women aged 35-39 than observed in the HSRC surveys, in contrast to the pattern of under-estimation seen in Figure 9.1.

**Figure 9.2:**
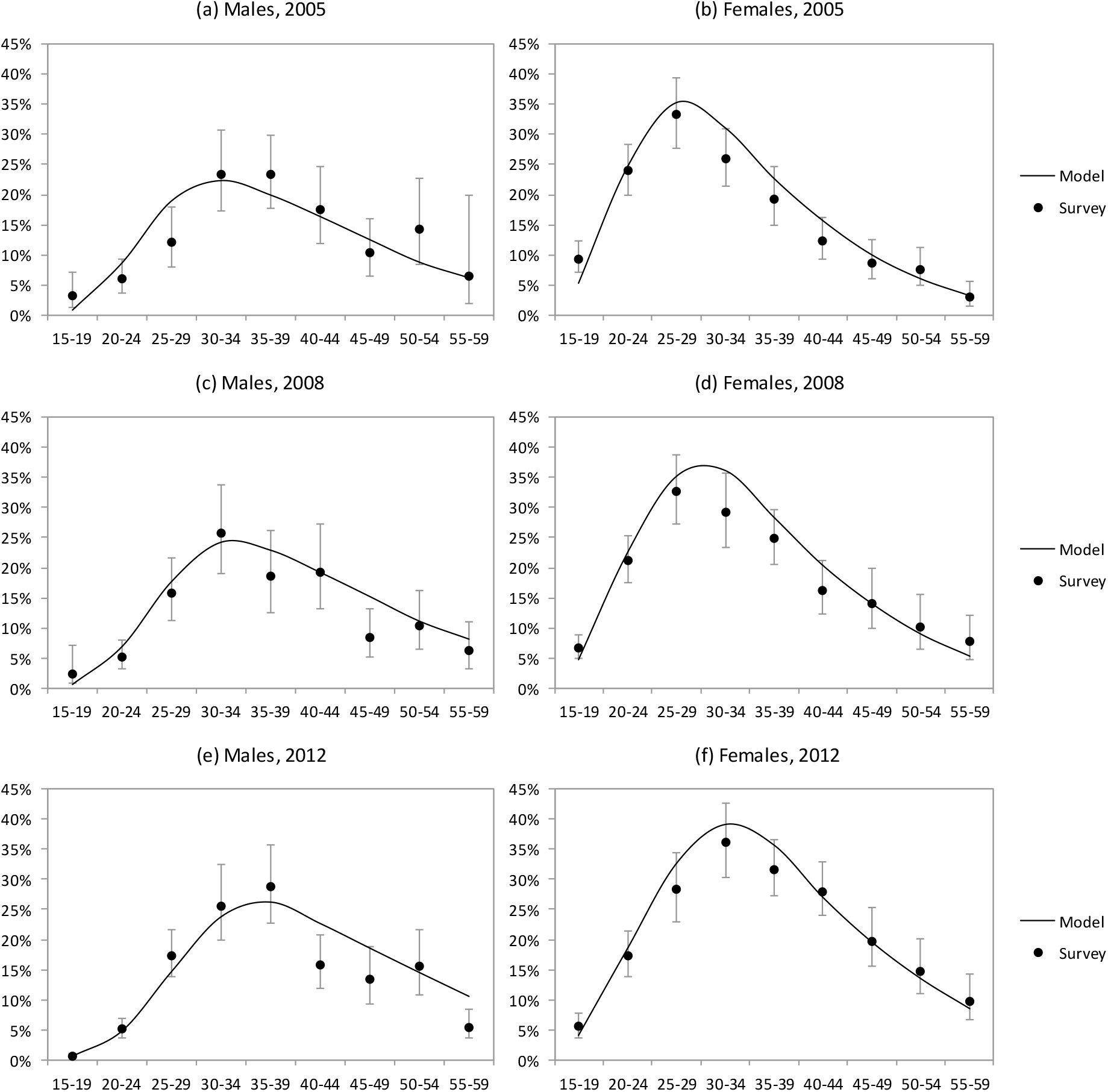
HIV prevalence in the general population. Model results are the average results generated using the 100 best-fitting parameter combinations.

Figure 9.3 compares the model estimates of HIV prevalence in key populations with the levels of HIV prevalence measured in local surveys. Although the model estimates of HIV prevalence in MSM appear higher than those observed in surveys, the surveys are probably biased towards younger MSM, who have lower HIV prevalence, and the model estimates are roughly consistent with those obtained in a similar recent analysis that controlled for age differences between modelled and observed MSM populations [160]. Model estimates of HIV prevalence in female sex workers appear roughly consistent with survey data, though slightly higher on average.

**Figure 9.3:**
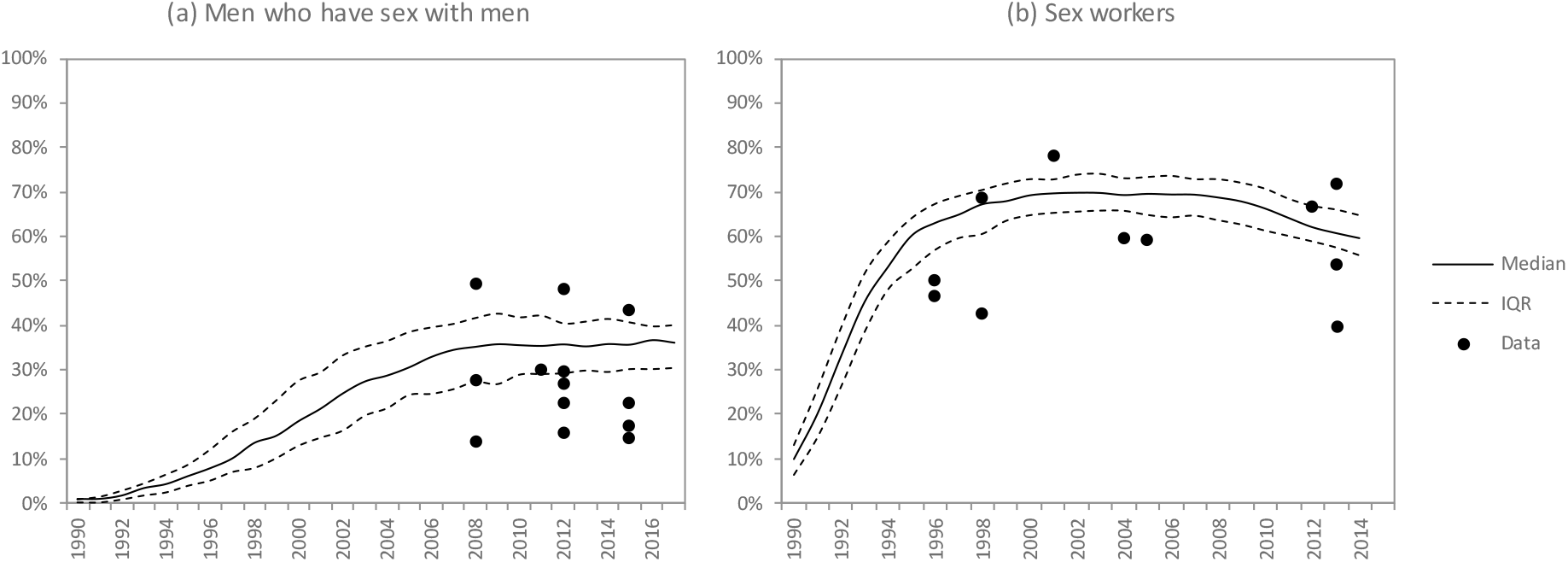
HIV prevalence in key populations. Model results are the median HIV prevalence estimates and interquartile ranges (IQRs) generated using the 100 best-fitting parameter combinations.

Although the 2002 HSRC household survey data have not been used in the calibration of the model, it is useful to compare the pattern of HIV prevalence by educational attainment, as estimated in the 2002 survey, with that simulated in 2002. Figure 9.4 shows the model estimates of HIV prevalence in 2002, in the population aged 15 and older, compared with the HIV prevalence measured in the 2002 HSRC household survey [507]. The model results are in close agreement with the survey results, except in the case of individuals with incomplete secondary education. The model and survey data both suggest that HIV prevalence is lowest among individuals with tertiary education or no education, while HIV prevalence is highest in individuals with secondary education.

**Figure 9.4:**
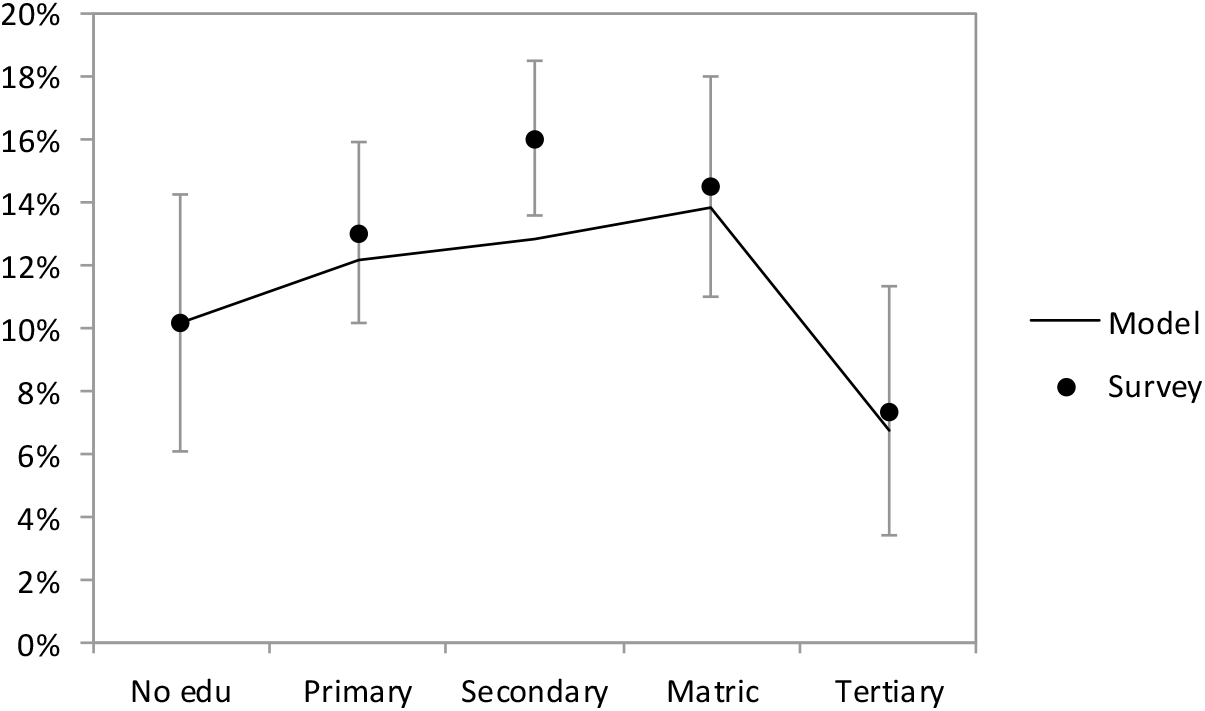
HIV prevalence in the population aged 15 and older, in 2002

## 10. Discussion

### 10.1 Strengths

The previous section shows that the MicroCOSM model matches reasonably closely the observed levels of HIV prevalence in South Africa by age and sex, as well as the observed changes in HIV prevalence over time. The model also matches observed patterns of HIV prevalence by educational attainment, by urban-rural location and by history of recent migration. Estimates of HIV prevalence in key populations (sex workers, MSM and prisoners) are roughly consistent with surveys, although none of the key population surveys are nationally representative, and surveys of MSM are likely to be biased towards younger MSM. Unlike previous models of HIV in Africa, MicroCOSM is calibrated to HIV prevalence data from antenatal clinics based on a fully mechanistic representation of fertility and the factors that affect public antenatal clinic attendance, rather than using an abstract or heuristic representation of antenatal bias. This allows us to understand why HIV prevalence in pregnant South African women differs so substantially from HIV prevalence in the general South African population, and why antenatal bias may be changing over time.

The model has also been calibrated to match total numbers of HIV tests and male circumcision operations performed in South Africa. The model estimates of levels of HIV diagnosis and ART coverage are consistent with the Thembisa model, an HIV model that has been calibrated to South African HIV testing and ART data. MicroCOSM simulates HIV testing in more detail than Thembisa, allowing for dynamics such as HIV testing in STI clinics, HIV testing in prisons and HIV testing following partner notification. This makes it possible to simulate a greater range of possible new HIV testing strategies.

### 10.2 Limitations and scope for further research

#### Contraception

A limitation of the model of contraception is that we have not modelled the recent introduction of hormonal implants [508]. This contraceptive method could potentially have a dramatic effect on fertility levels in South Africa, since it is one of the most effective non-permanent contraceptive methods [66]. Although there has been concern that its efficacy may be compromised in women who are on efavirenz-based antiretroviral treatment, it remains more effective than other hormonal methods in these women [63, 66]. In addition, its use is not associated with increased HIV transmission risk [462, 470].

Another limitation is that we have not modelled the potential association between hormonal contraceptive use and HIV diagnosis. Morroni *et al* [48] found that among South African women who had been diagnosed positive, many women reported adopting condoms and discontinuing hormonal contraceptive methods since diagnosis: 60% adopted condoms and 38% discontinued a contraceptive method (mostly injectable or oral contraceptives). This is consistent with the previously-noted reductions in HIV risk behaviours after HIV diagnosis and the previously-noted negative association between use of barrier methods and use of hormonal methods. However, as the Morroni study is the only study that showed direct evidence of an effect of HIV diagnosis on hormonal contraceptive use, we have not modelled this effect.

Although this analysis models the effect of hormonal contraception on the risk of HIV transmission, we have not modelled the effects of hormonal contraception on other STIs. For example, the risk of chlamydia is significantly increased in women who use oral contraceptives [509] and injectable contraceptives [510]. The frequency of HSV-2 shedding has also been found to increase in women who are using hormonal contraception [511, 512], and the incidence of vaginal candidiasis appears to be high in women using oral contraceptives [513–515]. On the other hand, the use of hormonal contraceptive methods is associated with a significantly reduced risk of bacterial vaginosis [516]. Further work is required to model these causal pathways, as the mediating effect of other STIs may be important in understanding the relationship between hormonal contraceptive use and HIV transmission risk.

#### Pregnancy

A limitation of our model is that it calculates rates of conception in such a way that rates of fertility in the model – by age, race and calendar year – are the same as those estimated by the ASSA2008 model. Although this has the advantage of ensuring that the model estimates of annual numbers of births to women in different age and race categories are realistic, it limits our ability to explain observed patterns of fertility. It would serve as an important validation of the model assumptions about sexual behaviour, contraception and fecundability if the model could roughly match observed patterns of fertility – by age, race, HIV status and calendar period – without the inclusion of the adjustment factors. Further work is required to model the determinants of fertility more realistically. This may include modelling of fertility goals, modelling of spontaneous and induced abortions, and modelling of the effect of STIs on fecundability. Modelling of pregnancy loss is important in understanding heterogeneity in birth intervals [73], and may therefore be necessary if we aim to validate the model using birth interval data [54]. HIV is known to increase the risk of spontaneous abortion and stillbirth [497], which is likely to be important if we wish to develop a more detailed model explaining the reduction in fertility in HIV-positive women. It is also possible that changes in access to abortion services and quality of antenatal care may change the fraction of pregnancies that lead to live birth, and this might be important to characterize if we intend to find explanations for observed changes in fertility over time.

The model currently assumes that the age at which women become fecund is independent of the age at which women start sexual activity. This might not be realistic; one would expect that girls with earlier onset of menarche would be likely to start sexual activity earlier than those with a later onset of menarche [517, 518], although evidence does not always support this [519]. Early menstrual cycles are often anovulatory, and hence the age at which a woman becomes fecund will typically be later than the age at which she starts menstruating; this may weaken any association between age at which women become fecund and age at sexual debut.

The assumption that variation in fecundability is entirely determined by female attributes is also not realistic; male partner characteristics can also play a role in determining fecundability. We have modelled the variation in fecundability in this way because most previously-published analyses are female-centric and do not quantify the variation that may be attributable to partner factors. Further work will be required to differentiate the relative contribution of male factors and female factors in determining couple-level variations in fecundability.

Although the model allows for the effect of pregnancy on HIV transmission, the model does not consider the potential effect of pregnancy on the transmission of other STIs. Female-to-male transmission of HSV-2 may be increased during pregnancy [511]. Pregnancy appears to increase the risk of vaginal candidiasis [520, 521], but it reduces the risk of bacterial vaginosis [522–525]. Modelling these effects may be important in understanding variation in HIV transmission risk as well as variation in STI prevalence across different settings.

#### Condom use

A limitation of our model of condom use is that it assumes condom discontinuation occurs only at the point at which a relationship switches from short-term to cohabiting or marital. In reality, condom discontinuation could occur at any point in a relationship, and it may be more realistic to assume a constant rate of discontinuation over the course of a relationship. A related limitation is that the model assumes condoms are initiated only at the start of a new relationship, and are not initiated in couples who have previously had unprotected sex (except in the case were one partner has disclosed that they are HIV-positive). This might be unrealistic, particularly if there are substantial numbers of couples who decide to adopt condoms for contraceptive purposes. Although it would be possible to modify the model to allow for more flexibility in the timing of condom adoption and discontinuation, this would require several new parameters and would make the model of condom use substantially more complex.

The assumption that couples use condoms either all the time or never may also be simplistic. For example, a study that measured condom use in HIV-discordant couples in Zambia over four consecutive 3-month intervals found that although overall levels of condom usage were high, only 23% of couples reported using condoms consistently in every interval, 26% reported unprotected sex in 1 interval and 24% reported unprotected sex in 2 of the 4 intervals [340]. On the other hand, a study in Zimbabwe found that couples tended to report using condoms either all the time or never; only about 7% of couples reported a level of condom use that was more than 0% but less than 100% [526].

#### Rape and intimate partner violence

Although we have modelled the effect of intimate partner violence, and have not found it to be a major driver of HIV transmission in South Africa [21], we have not modelled the potential effect of non-partner rape on the transmission of HIV and other STIs. South Africa has unusually high levels of rape. In a study of young women in Cape Town, for example, 11% reported having ever been raped [527]. In an Eastern Cape study of young men, 16% reported having ever raped a non-partner [528]. Perpetration of rape was in turn strongly related to gang membership and drug and alcohol use, which are additional social drivers that we have not considered. Further work is required to model the role of these factors in driving HIV transmission in South Africa.

#### Male circumcision

A limitation of this analysis is that we do not consider the effect of male circumcision on the transmission of STIs other than HIV. A systematic review of the evidence suggests that male circumcision significantly reduces the risk of ulcerative STIs (genital herpes, syphilis and chancroid) [529]. There appears to be little effect of male circumcision on gonorrhoea and chlamydia, but male circumcision may be partially protective against trichomoniasis in men [530, 531]. In addition, women whose partners are circumcised may be at a reduced risk of bacterial vaginosis [532].

The effect of male circumcision on HIV transmission risk may be mediated to some extent by male genital hygiene. Part of the reason why male circumcision is protective is that it is easier to keep the penis clean if it is circumcised, and lack of cleaning may cause inflammation of the foreskin, which increases HIV transmission risk [533]. Lack of cleaning might also increase the risk of other STIs [534, 535], which in turn are a risk factor for HIV. Male HIV status is correlated with poor genital hygiene in some studies [532, 536], and poor genital hygiene is more common in men of lower socioeconomic status, especially those who do not have access to indoor tap water [537]. However, due to the lack of data on male genital hygiene in South Africa and its effect on HIV transmission, we have not attempted to model this dynamic.

#### Sexually transmitted infections

A limitation of our model is that it does not consider the role of socio-economic factors in determining access to STI treatment. We have implicitly assumed that the rate of health-seeking and the choice of healthcare provider are independent of the individual’s race, educational attainment and location, which is probably unrealistic. Even when considering a given provider type, treatment practices may vary considerably depending on the patient’s socioeconomic characteristics. For example, Schneider *et al* [315] found that private practitioners often varied their prescriptions according to what they thought the STI patient could afford. Within public sector clinics, Bachmann *et al* [538] found that staff attitudes and provision of STI information to patients varied significantly according to the racial profile of the surrounding community, with staff attitudes to STI patients and provision of STI information both being poorer in clinics serving mainly black and coloured communities. Given the high proportion of HIV transmission that may be attributable to STIs in South Africa [473], it is important to better understand these socio-economic determinants of STI treatment. However, this is an under-researched area, and few local data exist to inform model assumptions.

Another limitation of our modelling approach is that it does not consider possible differences between STIs in heterosexuals and MSM. The assumptions about STI transmission probabilities and durations of infection that were previously estimated were produced prior to the inclusion of MSM in the model, and are thus representative of heterosexual STIs [18]. It is possible that these parameters may differ for MSM, for example because anal intercourse may be associated with different transmission risks (which probably depend on which partner is insertive), or because rectal STI symptoms may be less likely to be diagnosed and treated. However, this is an under-researched area, and most mathematical models of STIs in MSM make assumptions similar to those typically made in heterosexual STI models, often without distinguishing urethral and rectal STIs [539–542]. In the absence of good data, we have adopted a similarly simplistic approach, assuming that the average durations of STIs in MSM are the same as in heterosexual men, and that the probabilities of male-to-male STI transmission are the same as the probabilities of male-to-female transmission.

#### Mother-to-child transmission and paediatric HIV

A general limitation is that the model of mother-to-child transmission remains simplistic. Further work is required to extend the model so that the rate of mother-to-child transmission depends on the mother’s viral load [543–546], her stage of HIV disease [547], receipt of triple-drug ART [548], and duration and type of breastfeeding. The model of paediatric HIV also needs to be updated to include paediatric HIV testing and diagnosis, and to allow for differences in HIV survival depending on when ART is initiated [549]. To be consistent with the modelling of the natural history of HIV in adults, it will be important to extend the model to simulate changes in CD4 count and viral load in HIV-positive children.

#### Gender inequality

Studies suggest that condom use is lower in relationships in which women have less relationship power [550, 551]. Women who report lower ability to refuse sex or insist on condom use are also more likely to report being in concurrent partnerships [207, 208], which suggests an association between relationship power and concurrency. Relationship power in turn depends on factors such as partner age differences, which have also been shown to affect the odds of condom use [107]. Relationship power may also depend on factors such as women’s educational attainment and employment status, and how this compares with that of their partner. A limitation of our model is that we do not explicitly model these power dynamics; we make the naïve assumption that both partners in a relationship have equal influence in determining the probability of condom use. Further work is required to incorporate measures of relationship power into the model, and to link these to variables that can be influenced by government policies. For example, cash transfer interventions and job creation programmes focused on young women could reduce gender inequalities in socio-economic status, which may in turn increase women’s relationship power.

#### Wealth, income and employment status

Although our model simulates the effect of educational attainment on HIV risk behaviours, a limitation of the model is that it does not consider the potential effect of other economic measures on the risk of HIV acquisition (wealth, income and employment status). However, the evidence regarding the effect of these other economic measures is very mixed. Fraser-Hurt *et al* [177] analysed the 2008 HSRC household survey data and found that among African adults (aged 15 and older), HIV prevalence was relatively constant across income brackets of less than R4000 per month, but at higher income levels there was a strong negative association between monthly income and HIV risk. It was also found that among Africans, those who were informally employed had the highest risk of HIV (32%, compared to 28% in those who were unemployed and 20% among those who were formally employed). Among youth in rural Eastern Cape, it was found that socio-economic status (defined in terms of household goods and access to cash in emergency) was not significantly associated with HIV risk, either in men or in women [136, 552], and a similar lack of association was observed among youth in rural Limpopo [553]. Studies from the Africa Centre in rural KwaZulu-Natal have generally found that individuals in the middle quintiles of the household wealth index have the highest risk of HIV acquisition, after controlling for education and other HIV risk factors [457, 554]. In a synthesis of data from surveys in South Africa workforces, Colvin *et al* [555] found that among African employees, HIV prevalence levels were lower among managerial and technical workers than among unskilled labour. In Khutsong, an informal settlement near a major gold mining complex, Williams *et al* [100] found no significant association between employment status and HIV risk.

Taken together, the South African evidence suggest that after controlling for race, wealth and income have relatively little effect on HIV risk in the low-to middle-income categories – in fact, HIV risk may even be *positively* associated with greater wealth/income. However, at higher income levels, income and wealth may be protective. It is possible that the latter association is confounded by educational attainment, since the studies that did find significantly reduced HIV risk at the highest income levels [177, 555] did not control for educational attainment – which we have previously shown to be strongly associated with condom use. Many South Africa studies, such as those conducted in rural areas and informal settlements [100, 136, 457, 552–554], sample relatively poor communities, and may fail to identify a protective effect of higher wealth/income because they include relatively few individuals from the higher income groups.

These South African studies are consistent with data from other African countries. Forston [556] analysed DHS data from five African countries and found that in only one country (Tanzania) was there a significant effect of wealth on individual risk of infection; the relationship between wealth and HIV in Tanzania was non-linear, with HIV risk being highest around the median wealth category. In a similar analysis of DHS data from eight different African countries, Mishra *et al* [557] found that in many countries HIV prevalence in men was highest in the richest wealth quintile; it was also found that men in the richer wealth quintiles were more likely to report multiple partnerships and non-regular partnerships. However, the effect of wealth ceased to be significant after controlling for many of the other known HIV risk factors. As average GDP per capita in most other African countries is substantially lower than that in South Africa, one might expect higher income groups to be less well represented in these DHSs, and this might explain why these studies have generally not found a protective effect of high income or wealth.

#### Other social drivers

A limitation of this analysis is that we have not modelled many of the more complex structural drivers of HIV risk behaviour. For example, in an analysis of predictors of HIV risk in South African adolescents, Cluver *et al* [558] found that sexual risk behaviour was affected by a host of factors such as abuse (physical, emotional and sexual), mental health distress, and behavioural problems (e.g. aggressive behaviour, attention problems, social withdrawal), which in turn were strongly influenced by levels of structural deprivation (e.g. food insecurity, informal housing, community violence exposure), positive parenting and teacher social support. As another example, Nkosi *et al* [551], found that the frequency of unprotected sex in bar patrons in North West province was strongly affected by sexual relationship power, employment status and alcohol consumption. These are factors that are not currently modelled, but which could be added to the model in future.

### 10.3 Conclusion

Reducing HIV incidence in South Africa requires a marriage of the biomedical approach to HIV prevention and a broader developmental approach to address the social contexts that promote the spread of HIV and other STIs. Unfortunately, the social context has often been neglected, often because the links between social drivers and HIV are less direct and thus more difficult to quantify. This work lays the foundation for a more quantitative approach to understanding the role of social and behavioural factors that drive the spread of HIV in South Africa. In so doing, it also provides policymakers with insights into which sub-populations are currently at the greatest HIV risk. These key populations need to be the focus of new HIV prevention methods such as PrEP, cash transfers and vaccines.

## Acknowledgements

This research was funded primarily by the South African National AIDS Council. The HIV testing component of the model was funded by the HIV Modelling Consortium. We are grateful to the Centre for High Performance Computing for providing us with computing facilities to run our model, and we thank Cari van Schalkwyk for assistance in the use of these computing facilities.

